# A comprehensive benchmarking of WGS-based structural variant callers

**DOI:** 10.1101/2020.04.16.045120

**Authors:** Varuni Sarwal, Sebastian Niehus, Ram Ayyala, Sei Chang, Angela Lu, Nicholas Darci-Maher, Russell Littman, Karishma Chhugani, Arda Soylev, Zoia Comarova, Emily Wesel, Jacqueline Castellanos, Rahul Chikka, Margaret G. Distler, Eleazar Eskin, Jonathan Flint, Serghei Mangul

## Abstract

Advances in whole genome sequencing promise to enable the accurate and comprehensive structural variant (SV) discovery. Dissecting SVs from whole genome sequencing (WGS) data presents a substantial number of challenges and a plethora of SV-detection methods have been developed. Currently, there is a paucity of evidence which investigators can use to select appropriate SV-detection tools. In this paper, we evaluated the performance of SV-detection tools using a comprehensive PCR-confirmed gold standard set of SVs. In contrast to the previous benchmarking studies, our gold standard dataset included a complete set of SVs allowing us to report both precision and sensitivity rates of SV-detection methods. Our study investigates the ability of the methods to detect deletions, thus providing an optimistic estimate of SV detection performance, as the SV-detection methods that fail to detect deletions are likely to miss more complex SVs. We found that SV-detection tools varied widely in their performance, with several methods providing a good balance between sensitivity and precision. Additionally, we have determined the SV callers best suited for low and ultra-low pass sequencing data.

## Introduction

Structural variants (SVs) are genomic regions that contain an altered DNA sequence due to deletion, duplication, insertion, or inversion^1^. SVs are present in approximately 1.5% of the human genome^1,2^, but this small subset of genetic variation has been implicated in the pathogenesis of psoriasis^3^, Crohn’s disease^4^ and other autoimmune disorders^5^, autism spectrum and other neurodevelopmental disorders^6–9^, and schizophrenia^10–13^. Specialized computational methods—often referred to as SV callers—are capable of detecting structural variants directly from sequencing data. At present, the reliability, sensitivity, and precision of SV callers has not been systematically assessed. We benchmarked currently-available WGS-based SV callers in order to determine the efficacy of available tools and find methods with a good balance between sensitivity and precision.

Substantial differences exist in the number of identified variants in SV catalogs published during the past decade. The 1000 Genomes Project SV dataset identified over 68,000 SVs^14^, a genome-wide survey of 769 Dutch individuals identified approximately 1.9 million structural variants^15^, and a survey based on profiled whole genomes of 14,891 individuals across diverse global populations identified 498,257 SVs^16^. In addition, discrepancies in the number of SVs reported by these methods suggest that SV callers may fail to detect SVs and may report false positives (i.e., SVs that do not actually exist).

Lack of comprehensive benchmarking makes it impossible to adequately compare the performance of SV callers. In the absence of benchmarking, biomedical studies rely on the consensus of several SV callers^16,17^. In order to compare SV callers given the current lack of a comprehensive gold standard dataset, a recent study^18^ used long read technologies to define a ground truth in order to evaluate a large number of currently available tools. However, a comprehensive gold standard dataset is still needed; current long read technologies are prone to producing high error rates, which confounds efforts to detect SVs at single-base pair resolution. In response to the pressing need for a comprehensive gold standard dataset, our paper presents a rigorous assessment of sensitivity and precision of SV-detection tools when applied to mouse data.

## Results

### Preparing the gold standard data and WGS data

Over the last decade, a plethora of SV-detection methods have been developed (Table 1 and Supplemental Table 1), but the relative performance of these tools is unknown^19–25^. In order to assess the precision and accuracy of currently available SV callers, we simplified the problem presented to the detectors by using a set of homozygous deletions present in inbred mouse chromosomes. Methods failing to detect deletions are likely unreliable for the more challenging task of identifying other SV categories (e.g., insertions, inversions, translocations). We manually curated the mouse deletions used in this benchmarking study, and we used targeted PCR amplification of the breakpoints and sequencing to resolve the ends of each deletion to the base pair^26^. The same read alignment file which was used for manual curation of the deletions was used as an input to the structural variant callers, making our gold standard complete, containing all possible true deletions (true positives). We only used deletions since we could not confidently determine that other forms of SVs could be comprehensively detected with today’s SV callers.

**Table 1.**
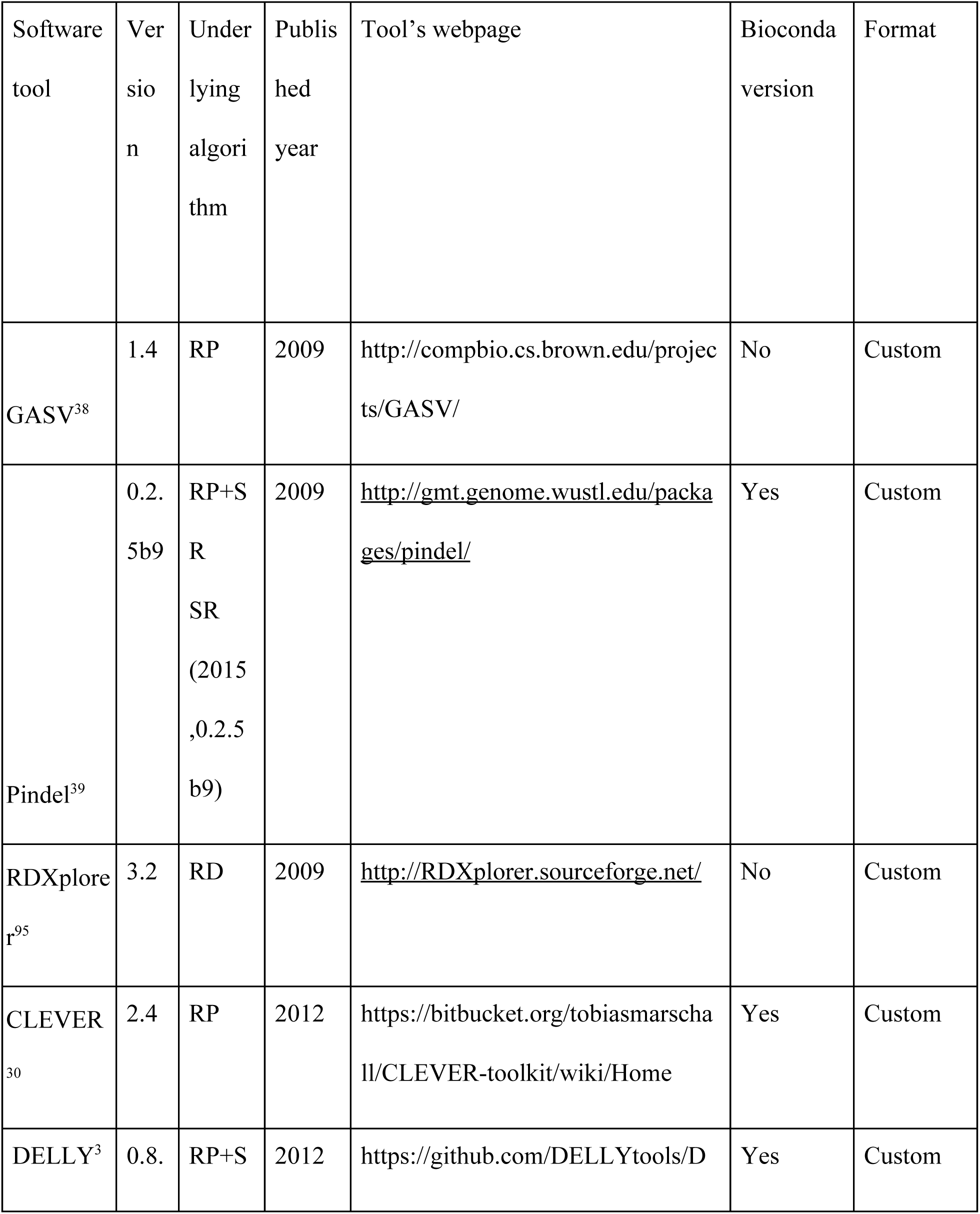
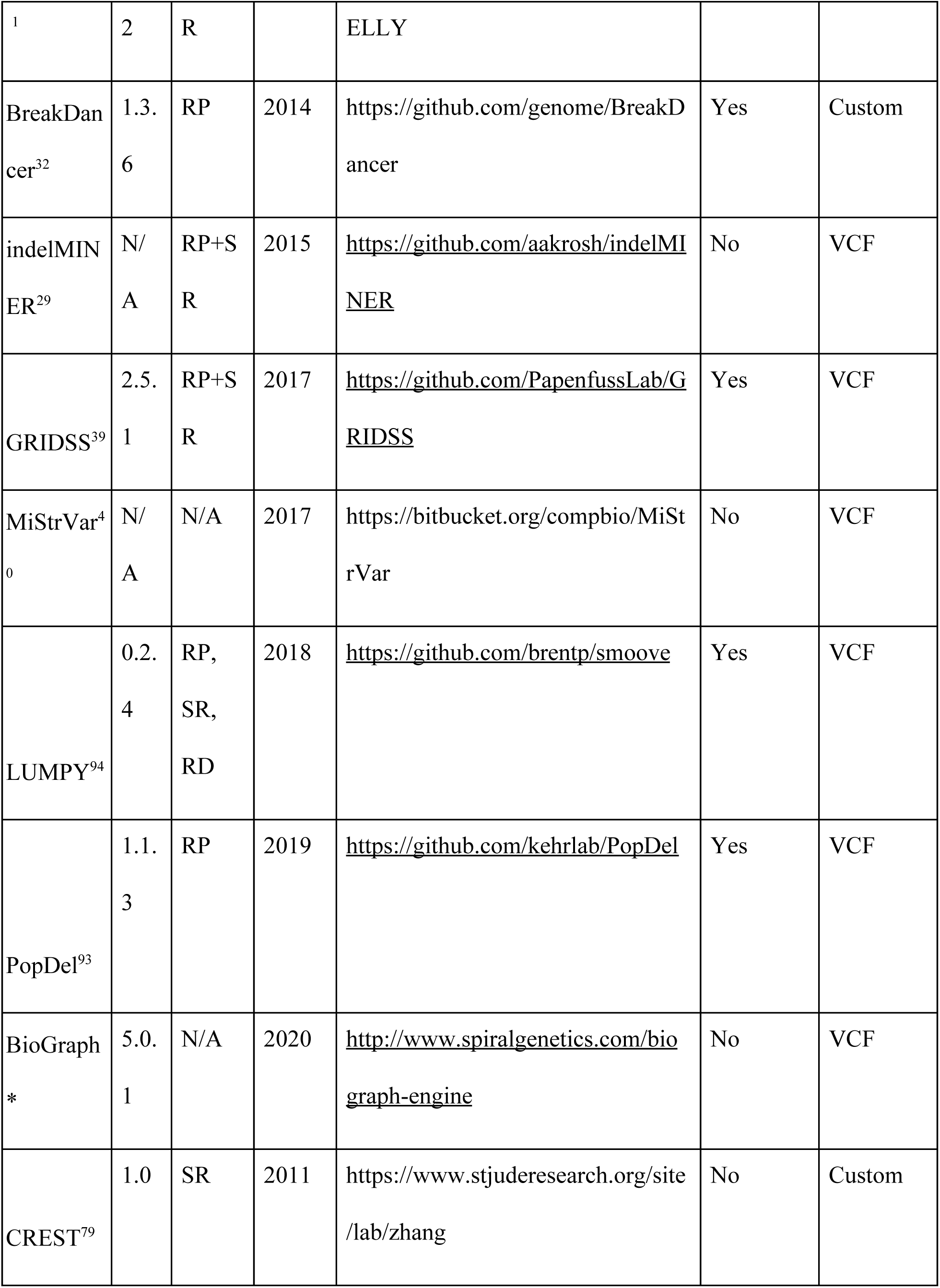
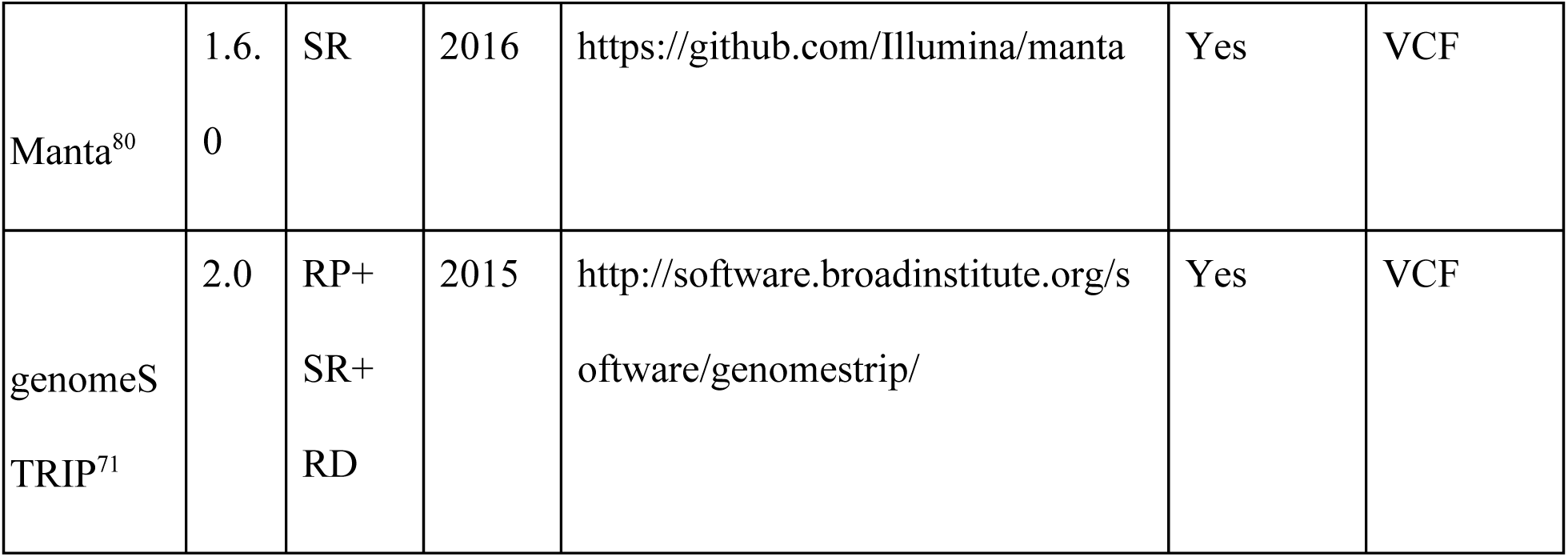
Overview of SV-detection methods included in this study. Surveyed SV-detection methods sorted by their year of publication from 2009 to 2018 are listed along with their underlying algorithm: Read-depth (RC), Read-Pair Algorithms (RP), Split-Read Approaches (SR), Discordant Pairs (DP), or a combination of algorithms. We documented the version of the software tool used in the study (‘Version’), the year the software tool was published (‘Published year’), the webpage where each SV-detection method is hosted (‘Tool’s webpage’), and whether or not the Bioconda package of the software was available (‘Bioconda version’). Asterisk (*) denotes that the method was proprietary.

The set of deletions we used among seven inbred strains, called with reference to C57BL/6J, is illustrated in Figure 1a and listed in Supplemental Table 2^26^. We filtered out deletions shorter than 50 bp, as such genomic events that are usually detected by indel callers rather than SV callers. In total, we obtained 3,710 deletions with lengths ranging from 50 to 239,572 base pairs (Supplemental Figure 1 and Supplemental Table 2^26^). Almost half of the deletions were in the range of 100-500 bp. Almost 30% of deletions were larger than 1000 bp (Supplemental Figure 1). High coverage sequence data was used as an input to the SV callers in the form of aligned reads. Reads were mapped to the mouse genome (GRCm38 Mouse Build) using BWA with -a option. In total, we obtained 5.2 billion 2×100 bp paired end reads across seven mouse strains. The average depth of coverage was 50.75x (Supplemental Table 3). Details regarding the gold standard and raw data preparation and analysis are presented in the Supplementary Materials.

**Figure 1.**
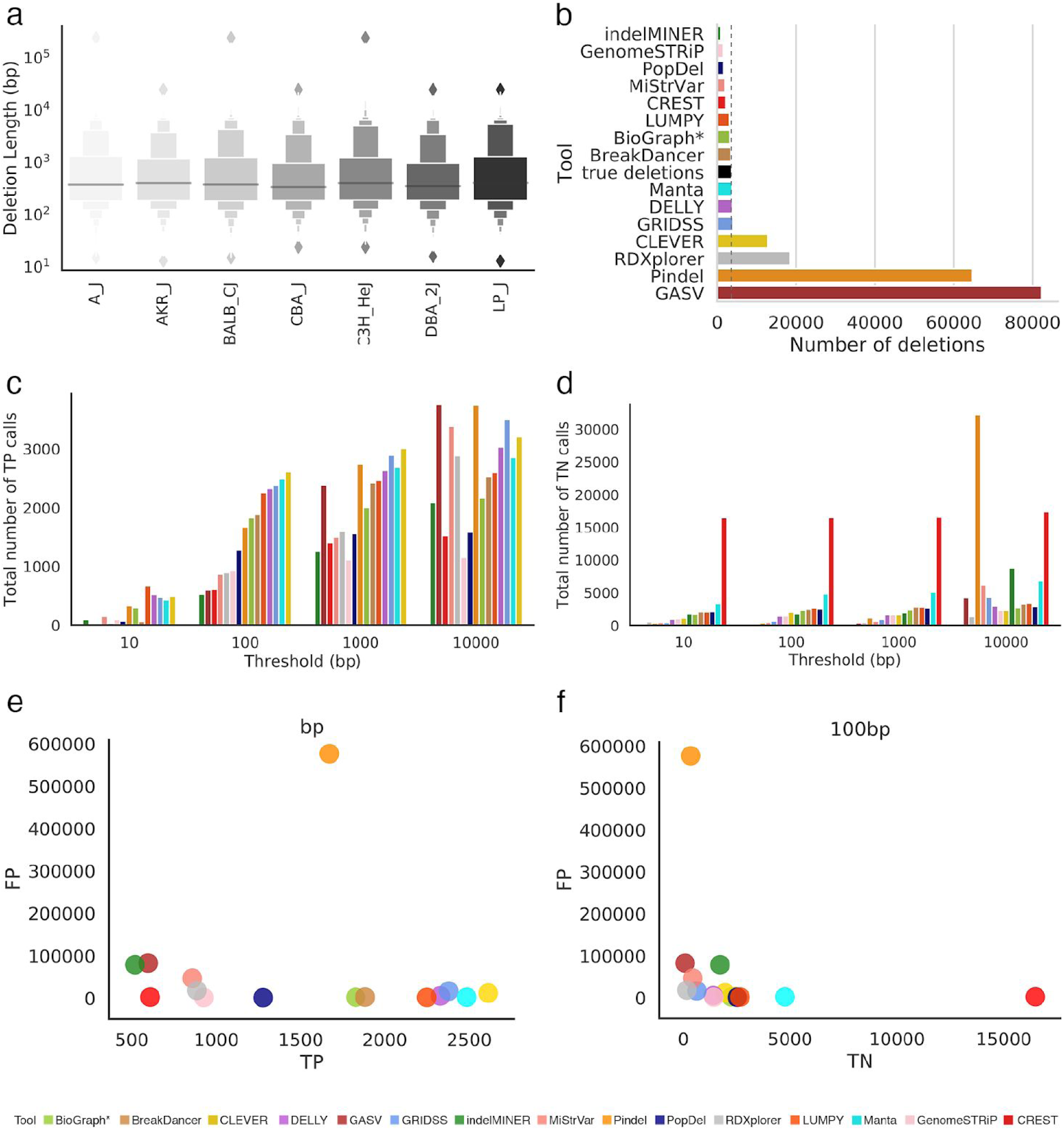
Comparison of inferred deletions across SV callers on mouse data. (a) Length distribution of molecularly-confirmed deletions from chromosome 19 across seven strains of mice. (b) Number of molecularly-confirmed deletions (‘true deletions’ black color) and number of deletions detected by SV callers. (c) Barplot depicting the total number of true positive calls across all error thresholds for each SV caller. (d) Barplot depicting the total number of true negative calls across all error thresholds for each SV caller. (e) Scatter plot depicting number of correctly detected deletions (true positives - ‘TP’) by number of incorrectly detected deletions (false positives - ‘FP’) at the 100 bp threshold. Deletion is considered to be correctly predicted if the distance of right and left coordinates are within the given threshold from the coordinates of true deletion. (f) Scatter plot depicting number of correctly detected non-deletions (true negatives - ‘TN’) by number of incorrectly detected deletions (false positives - ‘FP’) at the 100 bp threshold. An SV caller was considered to detect a given non-deletion if no deletions were reported in a given region.

### Choice of SV callers

For this benchmarking study, we selected methods capable of detecting SVs from aligned WGS reads. SV detection algorithms typically use information about coverage profile in addition to the alignment patterns of abnormal reads. We excluded tools that were designed to detect SVs in tumor-normal samples (e.g., Patchwork^86^, COPS^87^, rSW-seq^88^, bic-seq^63^, seqCBS^89^) and tools designed to detect only small (less than 50 bp in length) SVs (e.g., GATK^91^, Platypus^92^, Varscan^93^). Some tools were not suitable for inclusion in our dataset as they were unable to process aligned WGS data (e.g., Magnolya^27^). Other tools were designed solely for long reads (e.g., Sniffles^28^). The complete list of tools excluded from our analysis are provided in Supplemental Table 4. In total, we identified 55 suitable SV methods capable of detecting deletions from WGS data (Table 1 and Supplemental Table 1).

Our benchmarking study produced an analysis of the results generated by 15 SV-detection tools (Table 1). We were able to internally install and run all tools except Biograph, which was run by the developers of the tool. The remaining 40 tools could not be installed and were not included in this study. Supplemental Table 4 presents detailed information about the issues that prevented us from installing these software tools. Commands to install the tools and details of the installation process are provided in the Supplementary Materials.

### Comparing the performance of SV callers on mouse WGS data

We compared the performance of 15 SV callers in terms of inferring deletions. The number of deletions detected varied from 899 (indelMINER^29^) to 82,225 (GASV^38^). 53% of the methods reported fewer deletions than are known to be present in the sample (Figure 1b). We allowed deviation in the coordinates of the detected deletions and compared deviations to the coordinates of the true deletions. Even at relaxed stringency, the best method correctly detected the breakpoints of only 20% of known deletions in our curated dataset.

The majority of SV callers typically detect deletions whose coordinates differ from the correct positions by up to 100 bp. Figure 1c and 1d show the true positive (TP) and true negative (TN) rates for the SV callers at four different resolution values. It is notable that some tools with high TP rates also have decreased TN rates. For example, at the 100 bp threshold, the highest TP rate was achieved by CLEVER^30^ followed by Manta^80^, GRIDSS^39^ and DELLY^31^ (Figure 1c). However, for the same threshold, GRIDSS^39^ and DELLY^31^ underperform in the number of correctly detected non-deletions (TNs) compared to Manta^80^ (Figure 1d-f). The total number of false negative (FN) and false positive (FP) calls decreased with increase in threshold (Supplemental Figure 2). The FP rate for pindel Popdel^93^ was more susceptible to changes in the threshold as compared to Pindel^39^, GASV^38^. In general, the length distribution of detected deletions varied across tools and was substantially different from the distribution of true deletions across multiple SV detection methods (Figure 2 and Supplemental Table 2). Deletions detected by BreakDancer^32^ were the closest to the true median deletion length, while seven out of 15 SV callers overestimated deletion lengths (Figure 2).

**Figure 2.**
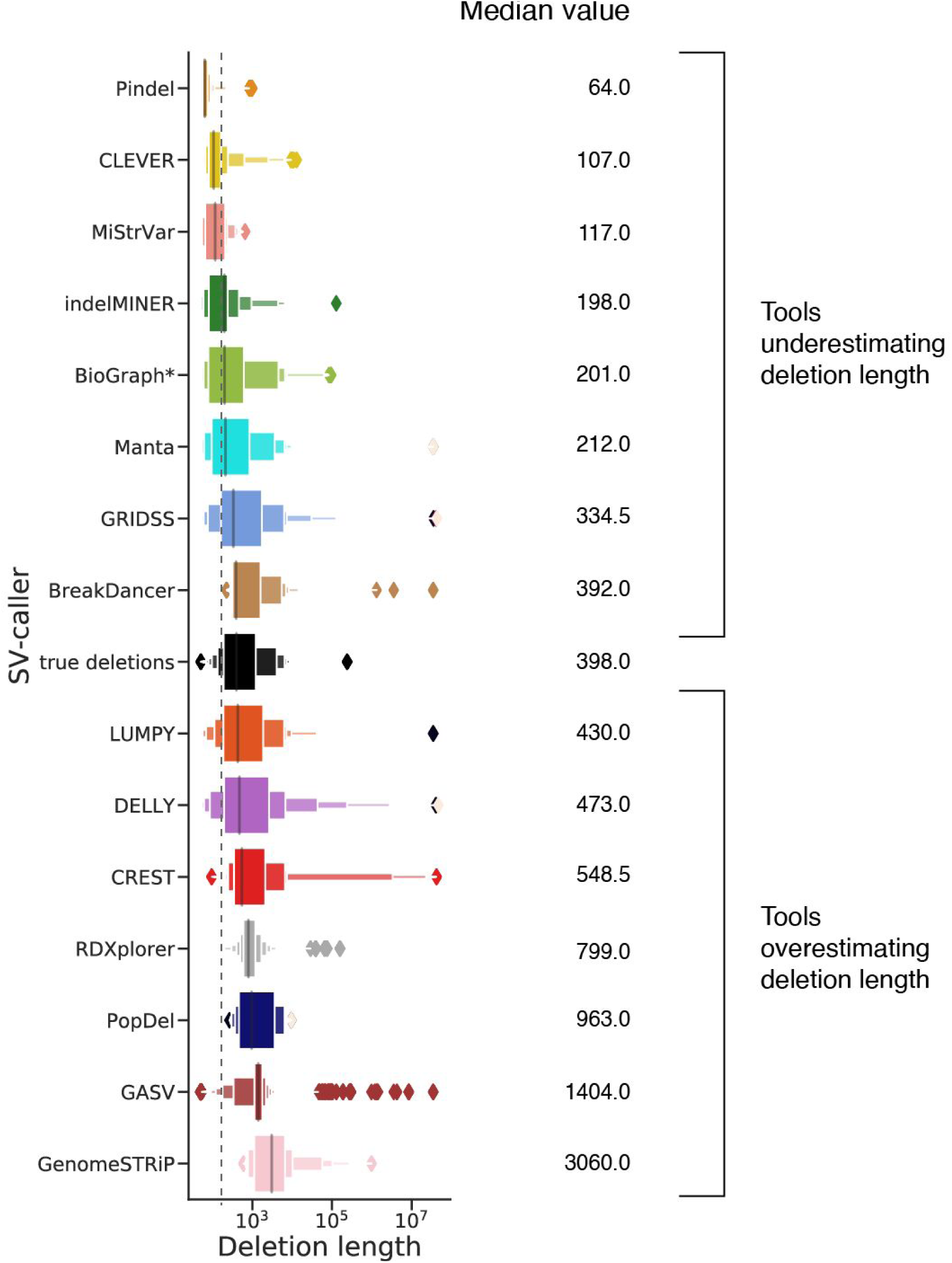
Length distribution of deletions detected by each SV caller. True deletions indicated in black. Tools were sorted in increasing order based on their median deletion length. The vertical dashed line corresponds to the median value of true deletions

Increasing the resolution threshold increases the number of deletions detected by the SV callers (Figure 1c). Several methods detected all deletions in the sample at 10,000 bp resolution but with precision close to zero (Figure 3b). We used the harmonic mean between precision and sensitivity (F-score) rates to determine the method with the best balance between sensitivity and precision. Several methods (e.g., Manta^80^, LUMPY^94^) offered the highest F-score for deletion detection—consistently between 100-10,000 bp resolution across all the mouse strains (Supplemental Figure 3). For a resolution of 10 bp, the method with the best performance for all the samples was LUMPY^94^ while at higher resolutions the best performing method was Manta^80^(Supplemental Figure 3). The method with the best precision for a threshold of 100-1,000 bp was PopDel^93^, but the sensitivity rate of PopDel^93^ did not exceed 50% (Figure 3a, Supplemental Figure 4 and 5).

**Figure 3.**
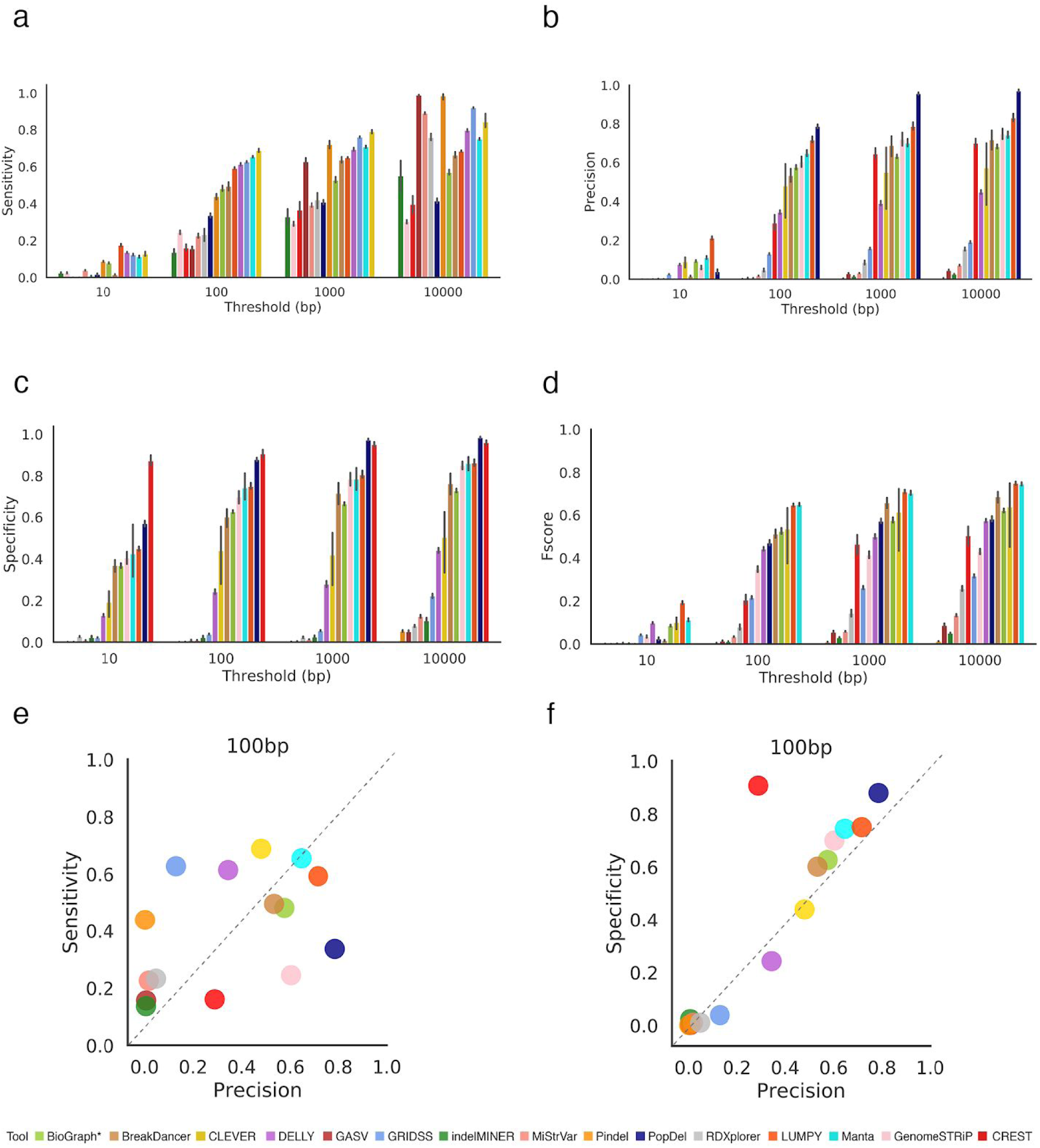
Comparing the performance of SV callers based on whole genome (WGS) data across seven inbred mouse strains. A deletion is considered to be correctly predicted if the distance of right and left coordinates are within the threshold τ from the coordinates of a true deletion. (a) Sensitivity of SV callers at different thresholds. (b) Precision of SV callers at different thresholds. (c) Specificity of SV callers at different thresholds. (d) F-score of SV callers at different thresholds. (e) Scatter plot depicting the Precision (x-axis) and Sensitivity (y-axis) for 100 bp threshold. (f) Scatter plot depicting the Precision(x-axis) and Specificity (y-axis) 100 bp threshold. Figures (a-d) are sorted in increasing order based on their performance at 100bp threshold. Results for other thresholds are presented in Supplemental Figure 6.

While specificity is often used to compare the deletions detected by tools, we use specificity to provide insights on the tools’ ability to predict diploid regions of the genome. Methods that produced a higher F-score tend to also have significantly higher specificity rates with Spearman’s correlations greater than 0.66 for thresholds greater than 100 bp (Figure 3d and Supplemental Figure 3 and 6) and are the most balanced in precision and sensitivity; few methods skewed towards just one of the metrics (Figure 3e and Supplemental Figure 7.)

Specificity rate was generally lower for the majority of the methods when compared with sensitivity rate. All methods except CREST^79^ with a high precision tend to also have significantly higher specificity rates, with Spearman’s correlation greater than 0.84 for thresholds greater than 10 bp (p-value<0.0005) (Supplemental Figure 8). Several tools, such as PopDel^93^ and LUMPY^94^, were able to balance precision and specificity, with rates exceeding 70% for each metric (Figure 3f). Manta^80^, LUMPY^94^ and CLEVER^30^ were the only methods able to successfully balance precision and sensitivity, with rates above 50% for each metric (Figure 3e and Supplemental Figure 7). CLEVER^30^ was able to achieve the highest sensitivity rate at the majority of thresholds (Figure 3a and Supplemental Figure 5). The most precise method we observed was PopDel^93^, with rates exceeding 80% for thresholds 1000 bp onwards, but the sensitivity of this method was two times lower than the majority of other tools (Figure 3b).

We examined whether the SV callers included in this study maintained similar SV detection accuracy across the different mouse strains. We compared results from each tool to study how consistent the results were across the samples. Among the tools with a sensitivity rate above 10%, LUMPY^94^ maintained the most consistent sensitivity rate across samples, with the highest rate of 60% when applied to both C3H/HeJ and CBA/J strains. The lowest sensitivity rate achieved by LUMPY^94^ was 58% for A/J and DBA/2J strains. Several tools, such as CREST^79^ and PopDel^93^ maintained a consistent specificity across mouse strains (Figure 3c and Supplemental Figure 6). Sensitivity rates were the most stable across the 7 different strains (Supplemental Figure 5). Specificity showed the highest variability among strains compared to other measures (Supplemental Figure 6). Precision shows the second highest variability across the strains, with the most stable results provided by Pindel^39^ and indelMINER^29^ (Supplemental Figure 4).

We have also compared CPU time and the maximum amount of RAM used by each of the tools. Across all of the tools, GASV^38^ required the highest amount of RAM while PopDel^93^ required the lowest amount of RAM to run the analysis. CREST^79^ required the longest amount of time to perform the analysis. Breakdancer^32^ was the fastest tool. We have also compared the computational resources and speed of SV callers based on datasets with full coverage and those with ultra-low coverage (Supplemental Figure 9).

### Performance of SV-detection tools on low and ultra-low coverage data

We assessed the performance of SV callers at different coverage depths generated by down-sampling the original WGS data. The simulated coverage ranged from 32x to 0.1x, and ten subsamples were generated for each coverage range. For each method, the number of correctly detected deletions generally decreased as the coverage depth decreased (Supplemental Figure 10). Some of the methods were able to call deletions from ultra low coverage (<=0.5x) data. While tools like Manta^80^ reached a precision of 82%, the overall sensitivity, specificity, and F-score values were less than 8% for all tools. None of the methods were able to detect deletions from 0.1x coverage.

As suggested by other studies^89^, most tools reached a maximum precision and specificity at an intermediate coverage (Figure 4b-c). Both the sensitivity rate and the F-score improved as the coverage increased (Figure 4a,d). Overall, DELLY^31^ showed the highest F-score for coverage below 4x (Figure 4d). For coverage between 8 and 32x, Manta^80^ showed the best performance. LUMPY^94^ was the only tool to attain precision above 90% for coverages 1x to 4x. However, a decreased sensitivity in coverages below 4x led to a decreased F-score when compared to DELLY^31^. Precision in results from DELLY^31^ for ultra-low coverage data was above 90% when the threshold was set at 1000 bp, but changing the threshold had no effect on LUMPY^94^ (Supplemental Figure 11).

**Figure 4.**
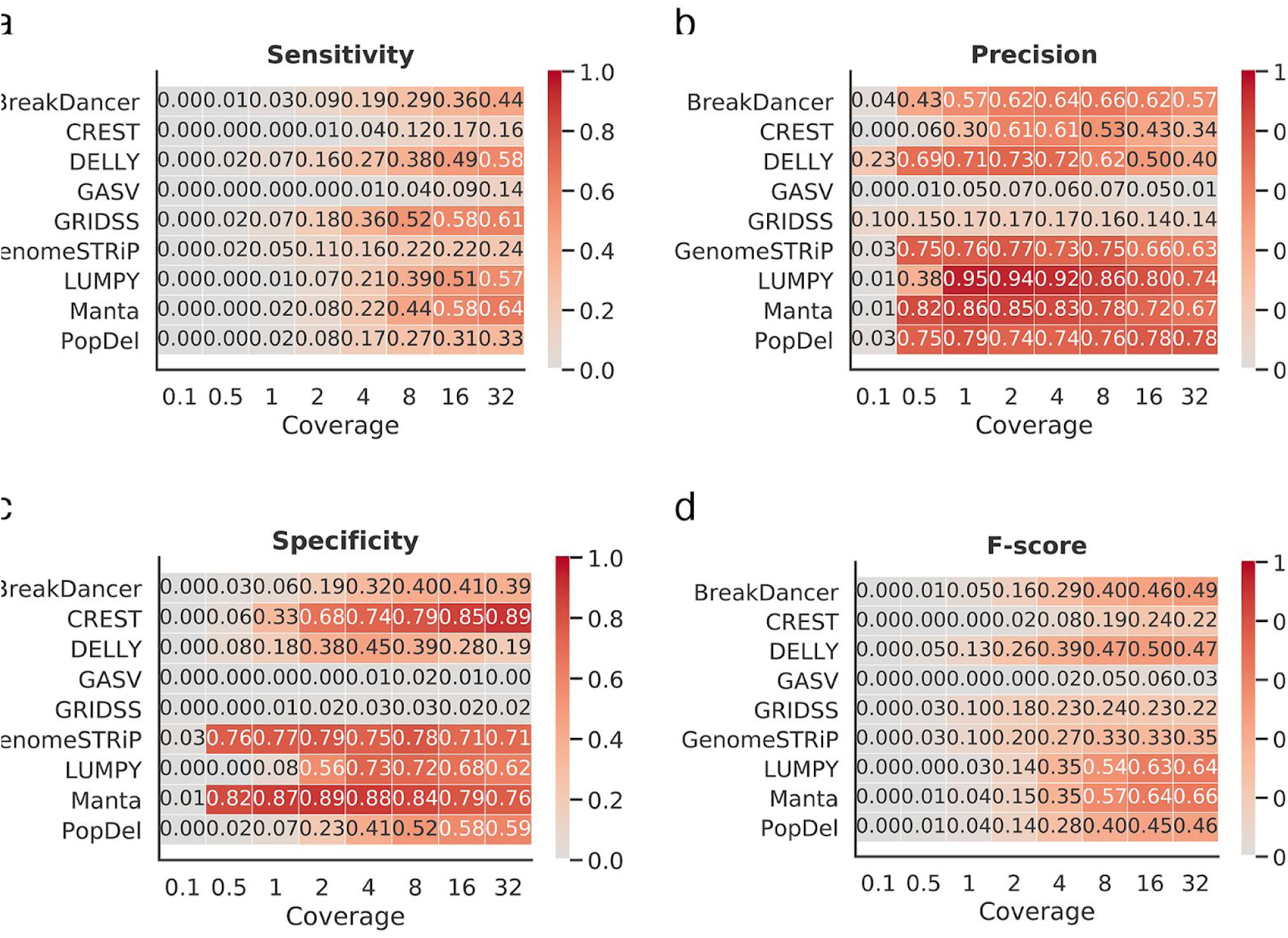
Performance of SV-detection tools on low and ultra-low coverage data. (a) Heatmap depicting the sensitivity based on 100 bp threshold across various levels of coverage. (b) Heatmap depicting the precision based on 100 bp threshold across various levels of coverage. (c) Heatmap depicting the specificity based on 100 bp threshold across various levels of coverage. (d) Heatmap depicting the F-score based on 100 bp threshold across various levels of coverage.

### Length of deletions impacts the performance of the SV callers

We separately assessed the effect of deletion length on the accuracy of detection for four categories of deletions (Figure 5). The performance of the SV callers was significantly affected by deletion length. For example, for deletions shorter than 100 bp, precision, specificity, and F-score values were typically below 40% regardless of the tool (Figure 5b,c,d and Supplemental Figures 12, 14, 15), while sensitivity values were above 50% for several tools (Figure 5a) (Supplemental Figure 13). For deletions longer than 100 bp, the best performing tool in terms of sensitivity and precision significantly varied depending on the deletion length (Figure 5a,b). CLEVER^30^ provided a sensitivity of above 60% for deletions less than 500 bp, however DELLY^31^ provided the highest sensitivity for deletions longer than 500 bp (Figure 5a and Supplemental Figures 17, 21, 25). LUMPY^94^ delivered the best precision for deletion lengths from 50-500 bp, and CLEVER^30^ performed well for longer deletion lengths (Figure 5b and Supplemental Figure 14, 18, 22, 26). indelMINER^29^ provided the high precision rate of detection of deletions in the range of 100 bp-500 bp and when longer than 1000 bp, but the precision of detecting deletion in the 500 bp-1000 bp range was lower than that of other tools (Figure 5b). In general, Manta^80^ and LUMPY^94^ were the only methods able to deliver an F-score above 30% across all categories (Figure 5d and Supplemental Figure 15, 19, 23, 27). Manta^80^ and CREST^79^ were the only tools with high specificity for deletions shorter than 500bp. For all the other tools, specificity was low across all the categories, except for deletions with lengths higher than 1,000bp (Figure 5c and Supplemental Figure 12, 16, 20, 24).

**Figure 5.**
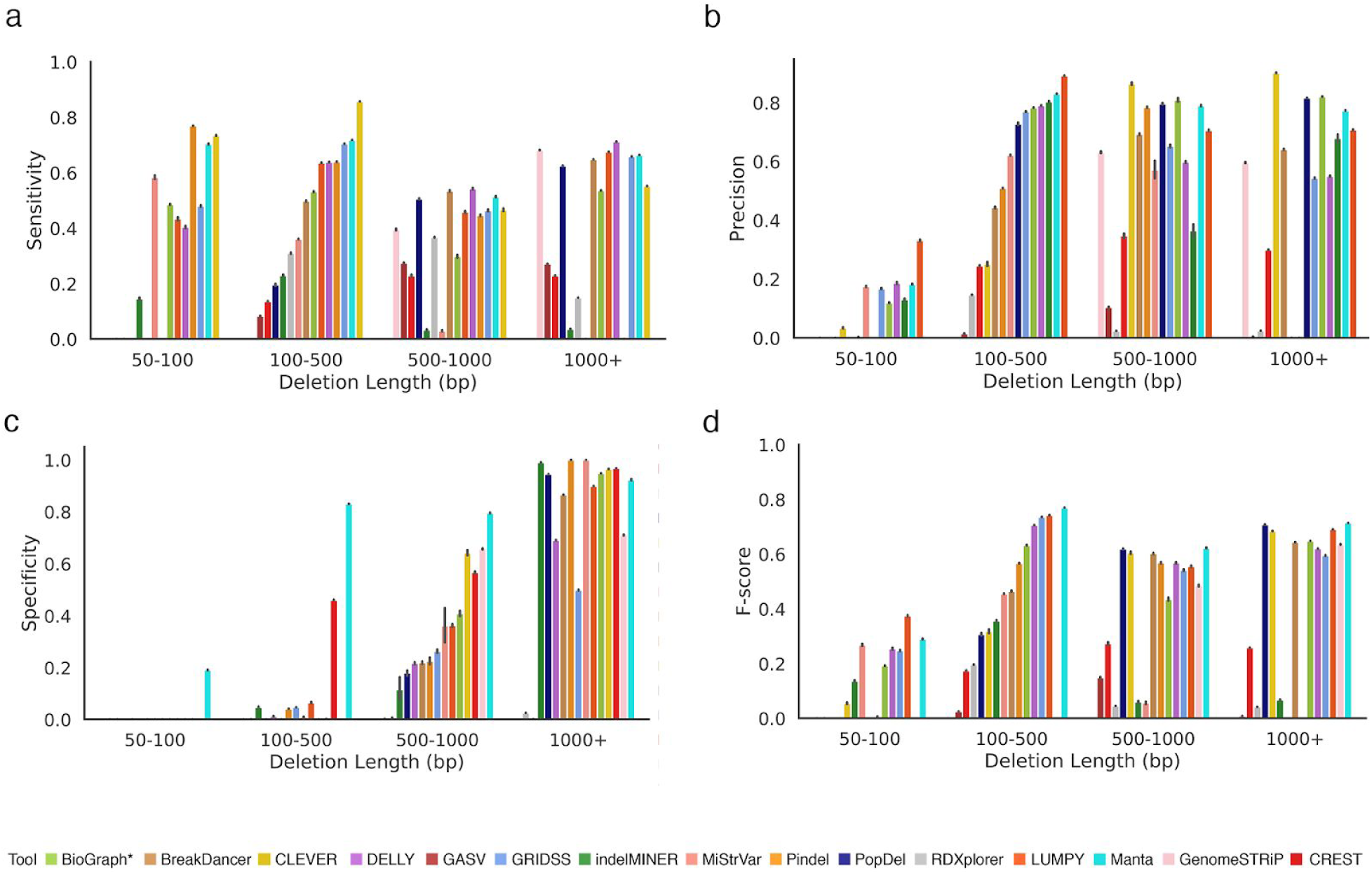
Comparing the performance of SV callers across various deletions lengths. (a) Sensitivity of SV callers at 100 bp thresholds across deletion length categories. (b) Precision of SV callers at 100 bp thresholds across deletion length categories. (c) Specificity of SV callers at 100 bp thresholds across deletion length categories. (d) F-score of SV callers at 100 bp thresholds across deletion length categories.

## Discussion

In this paper, we performed a systematic benchmarking of algorithms to identify structural variants (SVs) from whole-genome sequencing data. In contrast to methods which are used to identify single nucleotide polymorphisms and have coalesced around a small number of approaches, there is currently no consensus on the best way to detect SVs in mammalian genomes. Indeed, we were able to find 56 different methods, each claiming relatively high specificity and sensitivity rates in the original publication. Upon applying the tools to our curated datasets, many did not perform as reported in the original publication. This discrepancy may be because molecular data were not used in the analyses performed for the original publication. Instead, authors often solely derive conclusions from simulated data that may fail to capture the full complexity of real sequencing data^33^.

In comparison to previous benchmarking efforts based on simulated data ^19,21,25,34,35^, we obtained and employed a set of molecularly defined deletions for which breakpoints are known at base pair resolution. Other benchmarking studies have employed long-read-based gold standard datasets with approximate coordinates of deletions^18^. Long read technologies are often unable to cover the entire genome at sufficient resolution for precise SV characterization. In addition, long-read technologies carry a high error rate that limits their ability to detect SVs at single-base pair resolution. Our benchmarking method, using a gold standard set of molecular-defined deletions, overcomes the limitations of simulated data and incomplete characterization. Thus, our benchmarking study represents a robust assessment of the performance of currently-available SV detection methods when applied to a representative data set.

When installing the majority of SV callers, we noticed significant difficulties due to inadequate software implementation and technical factors^36^. Deprecated dependencies and segmentation faults were the most common reasons preventing successful tool installation^37^. The majority of the tools have a consensus on the output format to be used (Table S6), but the requirements for the format varied among tools. Lack of documentation about format requirements may further limit the use of SV callers.

We identified a series of factors that determined the performance of SV-caller methods. The most important factors were the size of deletions and the coverage of WGS data. For example, BreakDancer^32^ only detected deletions larger than 100 bp. Some tools achieved excellent sensitivity, with the caveat that their precision was close to zero. For example, Pindel^32^ achieved the highest sensitivity rate among all the tools, with a precision rate of less than 0.1%. Other tools (e.g., PopDel^93^) employ a more conservative SV detection approach, resulting in higher precision at the cost of decreased sensitivity for smaller deletion events. Few tools were able to maintain a good balance between precision and sensitivity. For example, Manta^80^, CLEVER^30^, LUMPY^94^, BreakDancer^32^, and BioGraph maintained both precision and sensitivity rates above 40%. In addition to differences in the accuracy of SV detection, we observed significant differences in run times and required computational resources (Supplemental Figure 9).

We envision that future SV-caller methods should enable detection of deletions with precise coordinates. The inability of current methods to precisely detect breakpoints was coupled with the issue of 61.5% tools underestimating the true size of SVs. A limitation of our benchmarking study is that our gold standard used inbred homozygous mouse genomes, which potentially poses as an easier target for assessment when compared to heterozygous human genomes. Additionally, the human genomes, for which most SV callers were designed, contain a higher number of repetitive regions than does the mouse, posing an additional challenge which is not reflected in mouse-based gold standard datasets.

## Data availability

WGS mouse samples used for benchmarking of SV-callers are available under the A/J (ERP000038), AKR/J (ERP000037), BALB/cJ (ERP000039), C3H/HeJ (ERP000040), DBA/2J (ERP000044), LP/J (ERP000045) accession numbers in the European Nucleotide Archive. VCF file with true deletions from gold standard, and the output VCF’s produced by the tools, the gold standard VCF’s, the analysis scripts, and figures are available at https://github.com/Mangul-Lab-USC/benchmarking-sv-callers-paper/

## Code availability

Source code to compare SV detection methods and to produce the figures contained within this text is open source, free to use under the MIT license, and available at https://github.com/Mangul-Lab-USC/benchmarking-sv-callers-paper/

## Acknowledgments

We thank Dr. Lana Martin for the helpful discussions and comments on the manuscript. We thank our colleagues Niranjan Shekar, Lisa Herta, and Adam English at Spiral Genetics for developing and running their tool, BioGraph software (http://www.spiralgenetics.com/biograph-engine) on mouse WGS data. We thank the authors of the tools surveyed in this work—Cenk Sahinalp, Ryan Layer, Ira Hall, Tony Papenfuss, Gerton Lunter, Michael Schatz, Alexander Schoenhuth, Ken Chen, Aakrosh Ratan, and Tobias Rausch—for providing helpful feedback and verifying information related to their tool.

## Author contributions

V.S. created scripts for running and evaluating the software tools. E.W., J.C., M.G.D., N.D.-M., R.A., R.C., R.L., S.C., and V.S. contributed to installing, running, and evaluating software tools. S.N. applied GRIDSS^39^, LUMPY^94^, Manta^80^, and PopDel^93^ to the mouse data and discussed evaluation metrics. R.A. curated data and prepared mouse data for evaluation with the software tools. A.L., R.A., and V.S. generated the figures; R.A. and V.S. generating the tables. E.E., J.F., M.G.D., S.M., S.N., and V.S. wrote, reviewed, and edited the manuscript. J.F. and S.M. led the project.

## Methods

### Running SV-detection tool

Commands required to run each of the tools and the installation details are available in Supplemental Table S5. The tools were run based on the recommended settings. LUMPY^94^ was run using smoove, as was recommended by developers of smoove. The diploidSV vcf files were used for Manta, based on the recommendation of the developers. BioGraph was run by developers of the tool, based on the provided BAM files.

### Convert the output of the SV-detection tool to a universal format

We have adopted the VCF format proposed by VCFv4.2, as the universal format used in this study. Custom format of SV-detection tools were converted to VCFv4.2. Description of custom formats is provided in Supplemental Table S6. Scripts to convert custom formats of SV-detection tools to VCFv4.2 are available at https://github.com/Mangul-Lab-USC/benchmarking-sv-callers-paper

### Compare deletion inferred from WGS data with the gold standard

We compared the deletions inferred from SV callers from WGS data (inferred deletions) with the molecular-based gold standard (true deletions). Start and end position of the deletion were considered when comparing true deletions and inferred deletions. Inferred deletion was considered correctly predicted if the distance of right and left coordinates are within the resolution threshold τ from the coordinates of true deletion. We consider the following values for resolution threshold τ: 0 bp, 10 bp, 100 bp, 1000 bp,10000 bp. Correctly predicted deletions are defined as true positives (TP). In case an inferred deletion matches several true deletions, we randomly choose one of them. Similarly, in case true deletion matches several inferred deletions we choose the first deletion that matches. randomly choose one of them. The deletion predicted by the SV caller but not present in the golden standard was defined as false positives (FP). Similarly, each deletion present in the gold standard was matched with only one deletion predicted by the software. The SV that was not predicted by the SV caller were defined as false negatives (FN). SV detection accuracy was assessed using various detection thresholds (τ). The accuracy at threshold τ is defined as the percentage of SVs with an absolute error of deletion coordinates smaller or equal to τ. To compute specificity, we have defined non-deletions calls as regions of the genome not containing deletions. We have generated the true set of non-deletions based on the gold standard. Start and end position of the non-deletion were considered when comparing true non-deletions and inferred non-deletions. Inferred non-deletion was considered correctly predicted if the distance of right and left coordinates are within the resolution threshold τ from the coordinates of true non-deletion. We consider the following values for resolution threshold τ: 0 bp, 10 bp, 100 bp, 1000 bp,10000 bp. Correctly predicted non-deletions are defined as true positives (TN).

We have used the following measured to compare the accuracy of SV-allers:

- Sensitivity=TP/(TP+FN)
- Precision=TP/(TP+FP)
- F-score=2*Sensitivity*Precision/(Sensitivity+Precision)
- Specificity= TN/(TN+FP)

The scripts to compare the deletions inferred by the SV caller versus the true deletions is available at https://github.com/Mangul-Lab-USC/benchmarking-sv-callers-paper.

### Compare computational performance of SV callers

The CPU time and RAM of each tool was measured to determine it’s computational performance. The statistics were measured for 1x coverage and full coverage bam files, with samples A/J and BALB/cJ for mouse data. The CPU time was computed using either the GNU time program that is inbuilt in make bash terminals or the Hoffman2 Cluster qsub command. For GNU time, we used this specific command /usr/bin/time -f “%e\t%U\t%S\t%M” which we either had to run manually on an interactive qsub session or through another method that wasn’t a qsub. This GNU time command would output one line containing Wallclock time in seconds, user-time in seconds, kernel-space time in seconds, and peak memory consumption of the process in kilobytes. CPU-time was calculated by adding user-time and kernel-space time. RAM usage was equivalent to peak memory consumption in the case of this command. For qsubs on the Hoffman2 Cluster, we used the command qsub -m e which would email the user a full list of records when the tool finished running. This list included CPU-time and Max Vmem which was designated as RAM usage for each tool.

### Downsampling the WGS samples

We have used custom script to downsample the full coverage BAM file to desired coverage. Existing tools (e.g., samtools) are not suitable for this purpose as they treat each read form a read pair independently, resulting in singletons reads in the downsample BAM file.

## Supplementary Tables

**Table S1.**
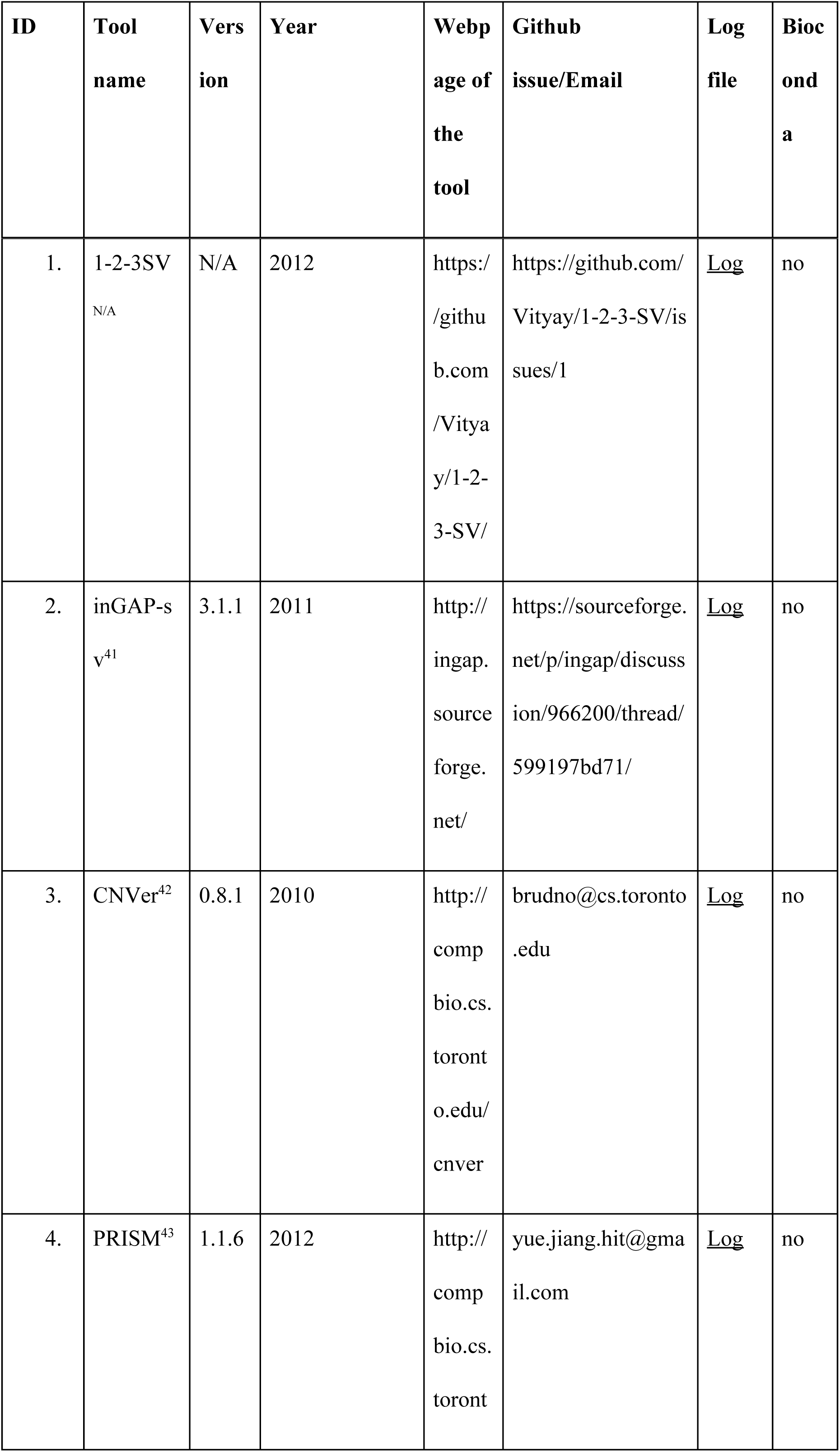
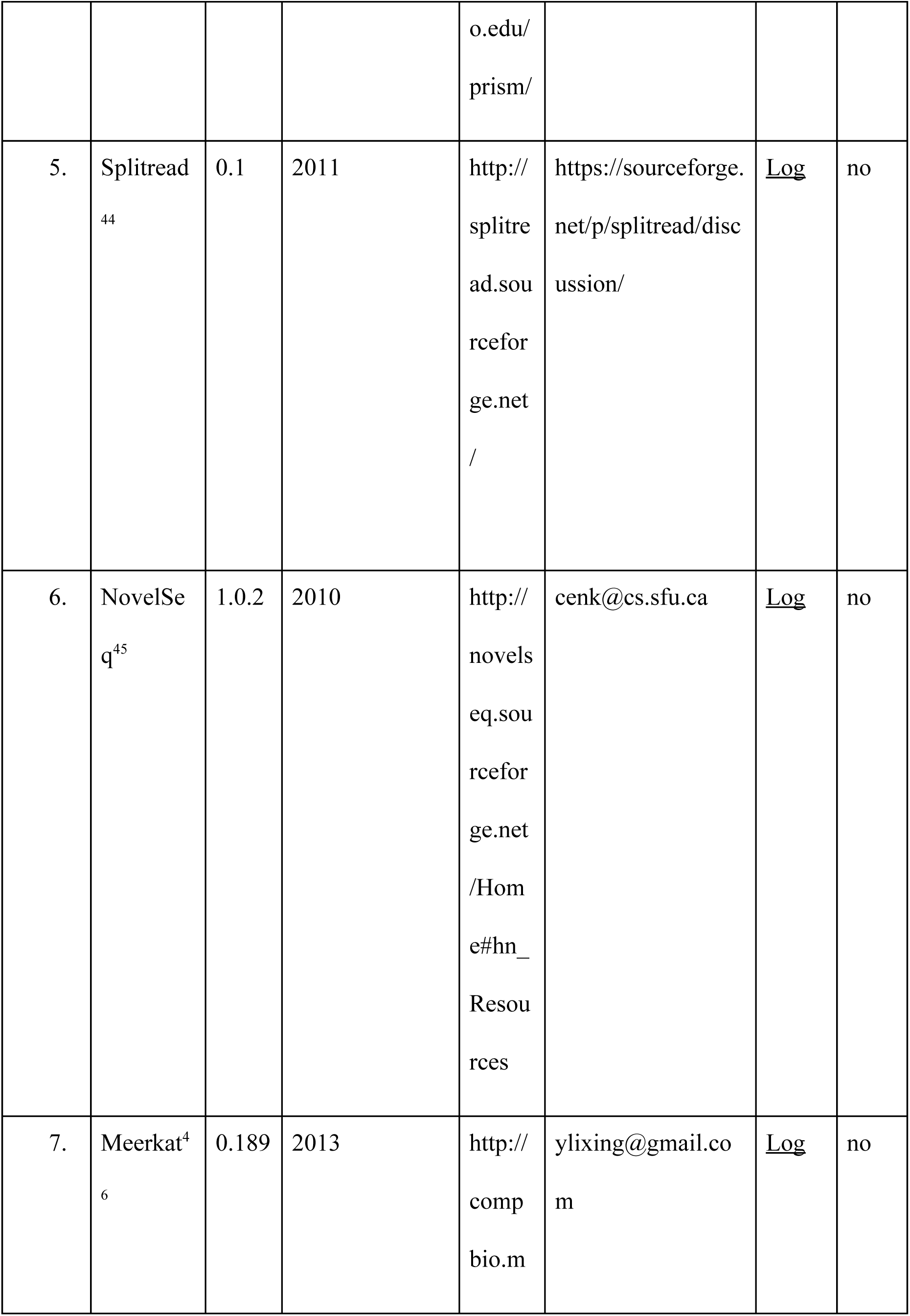
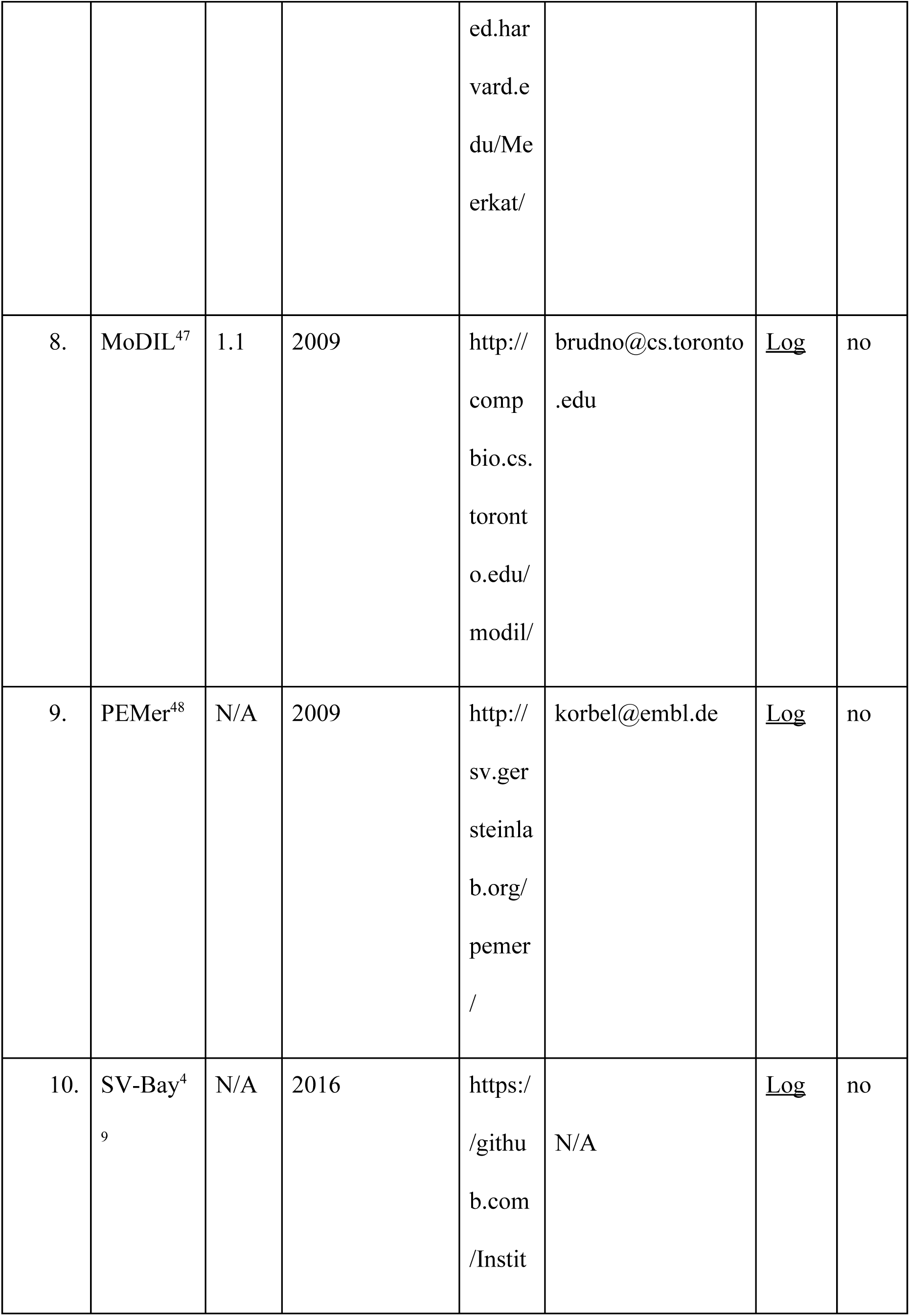
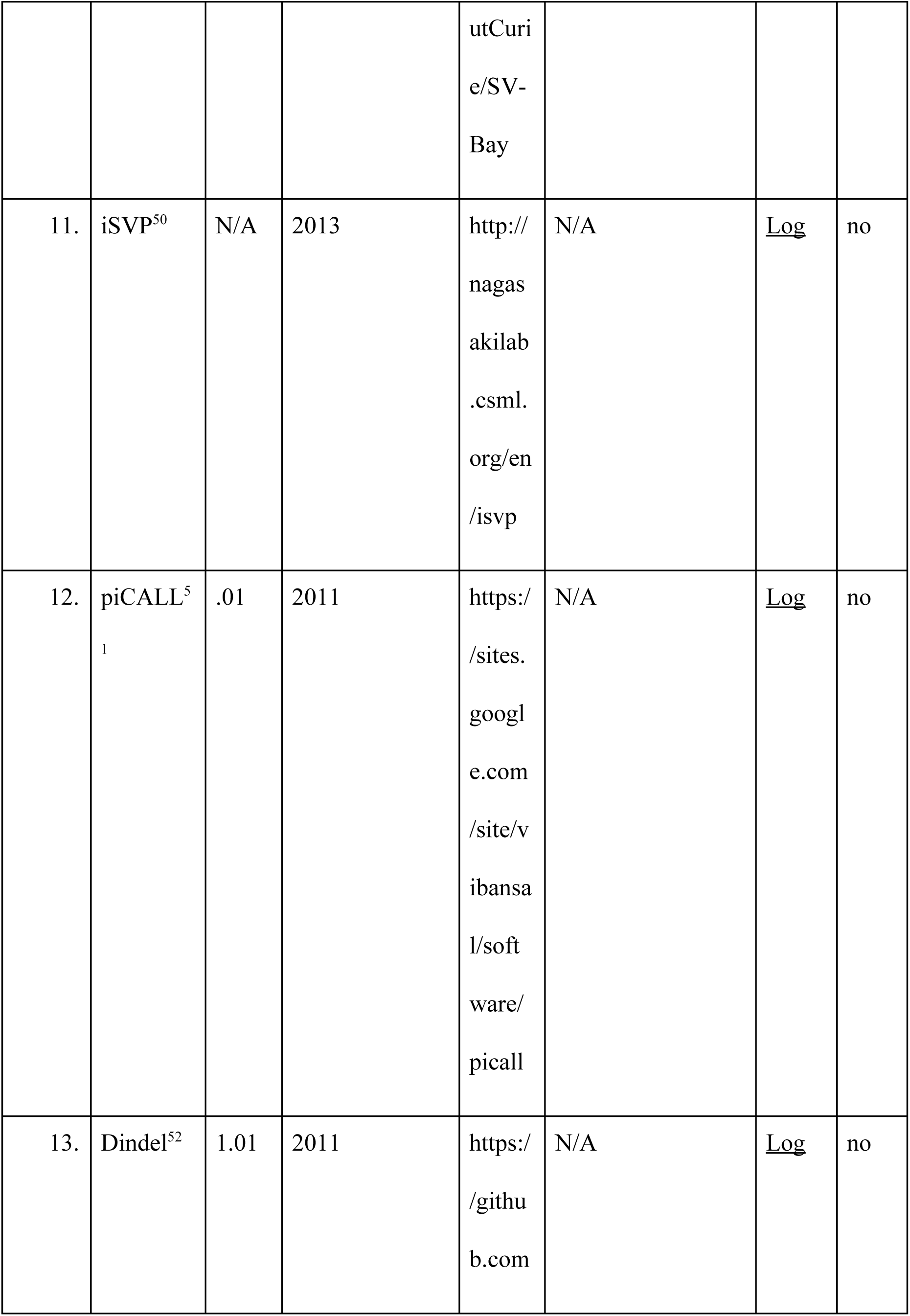
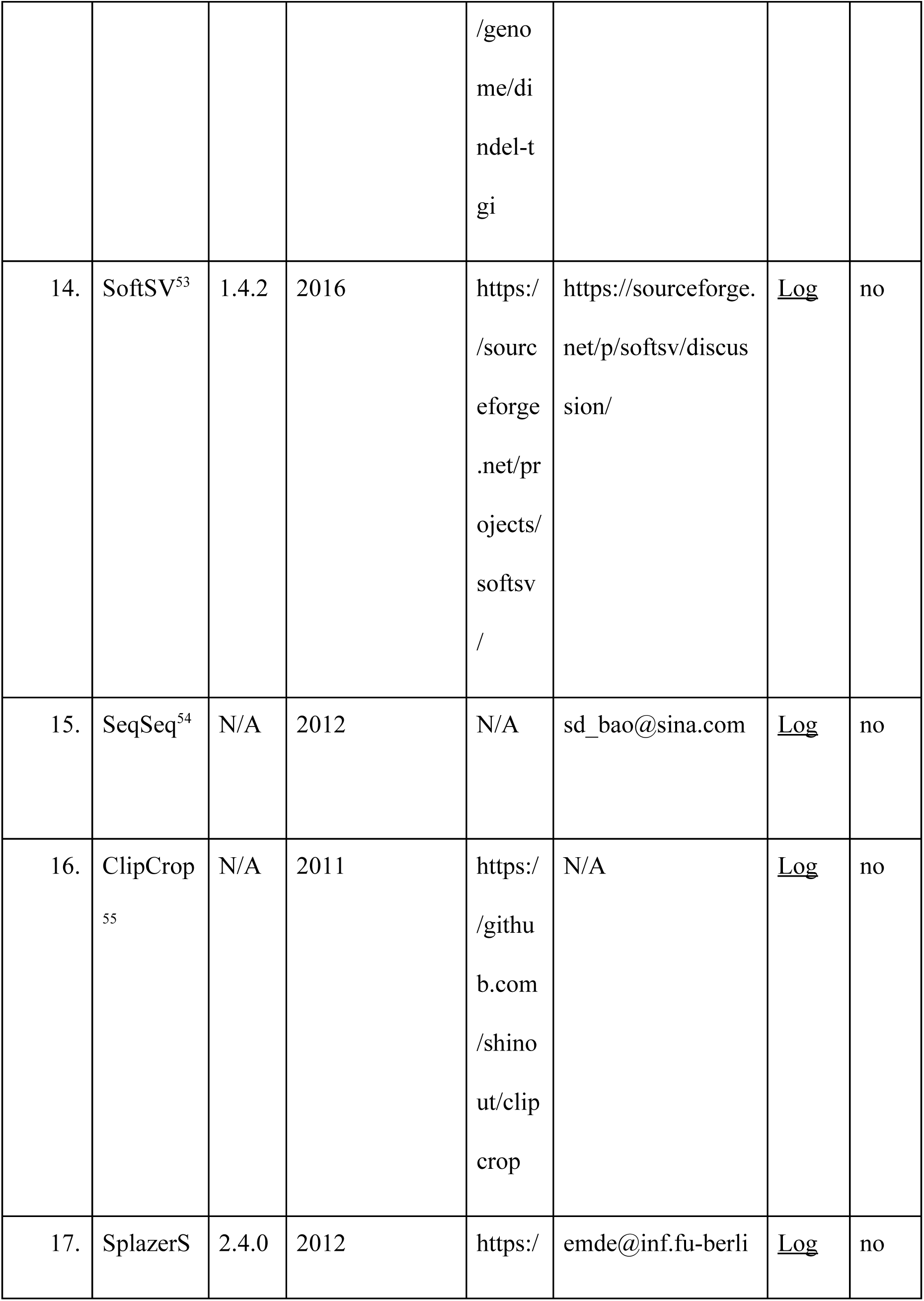
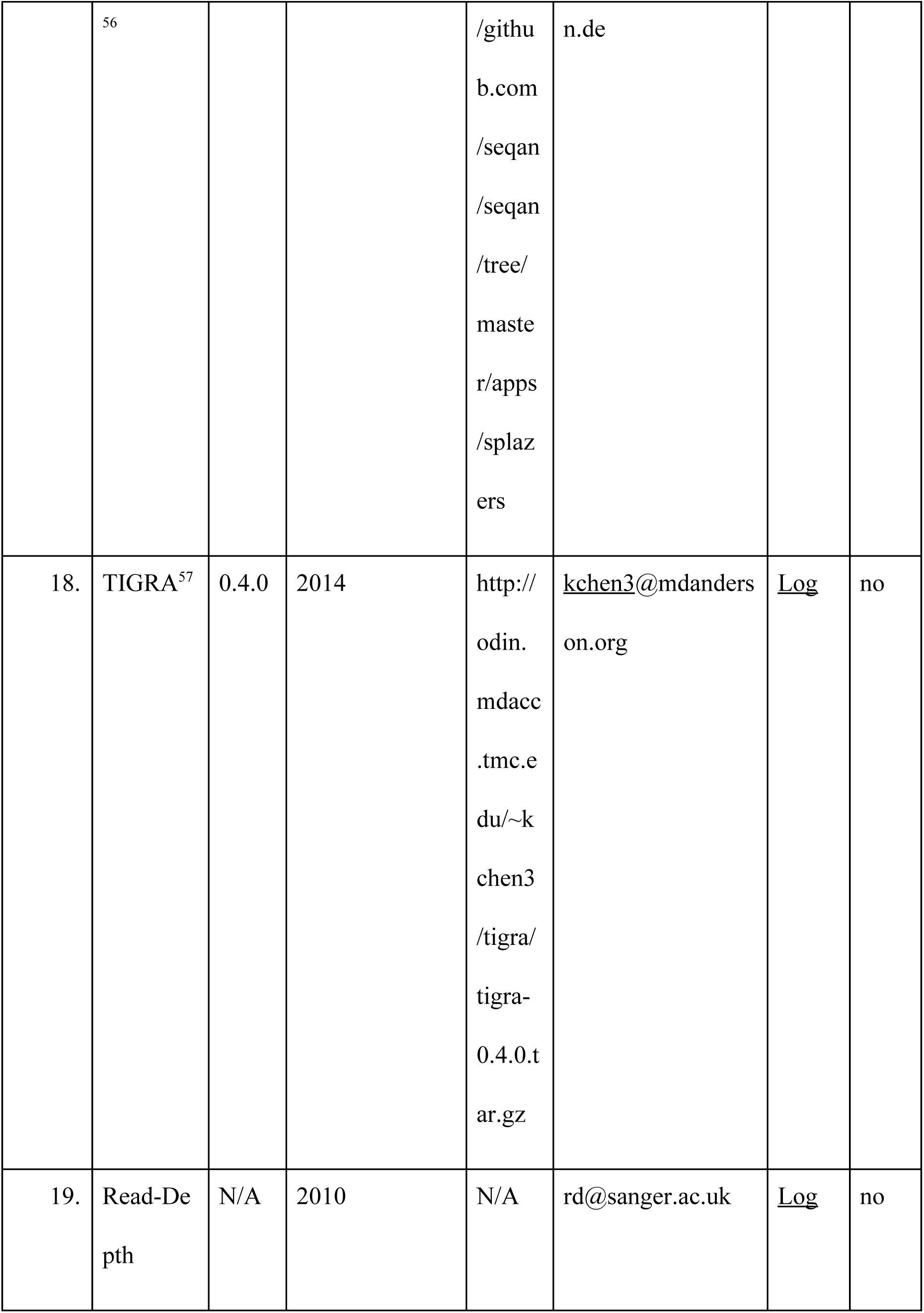
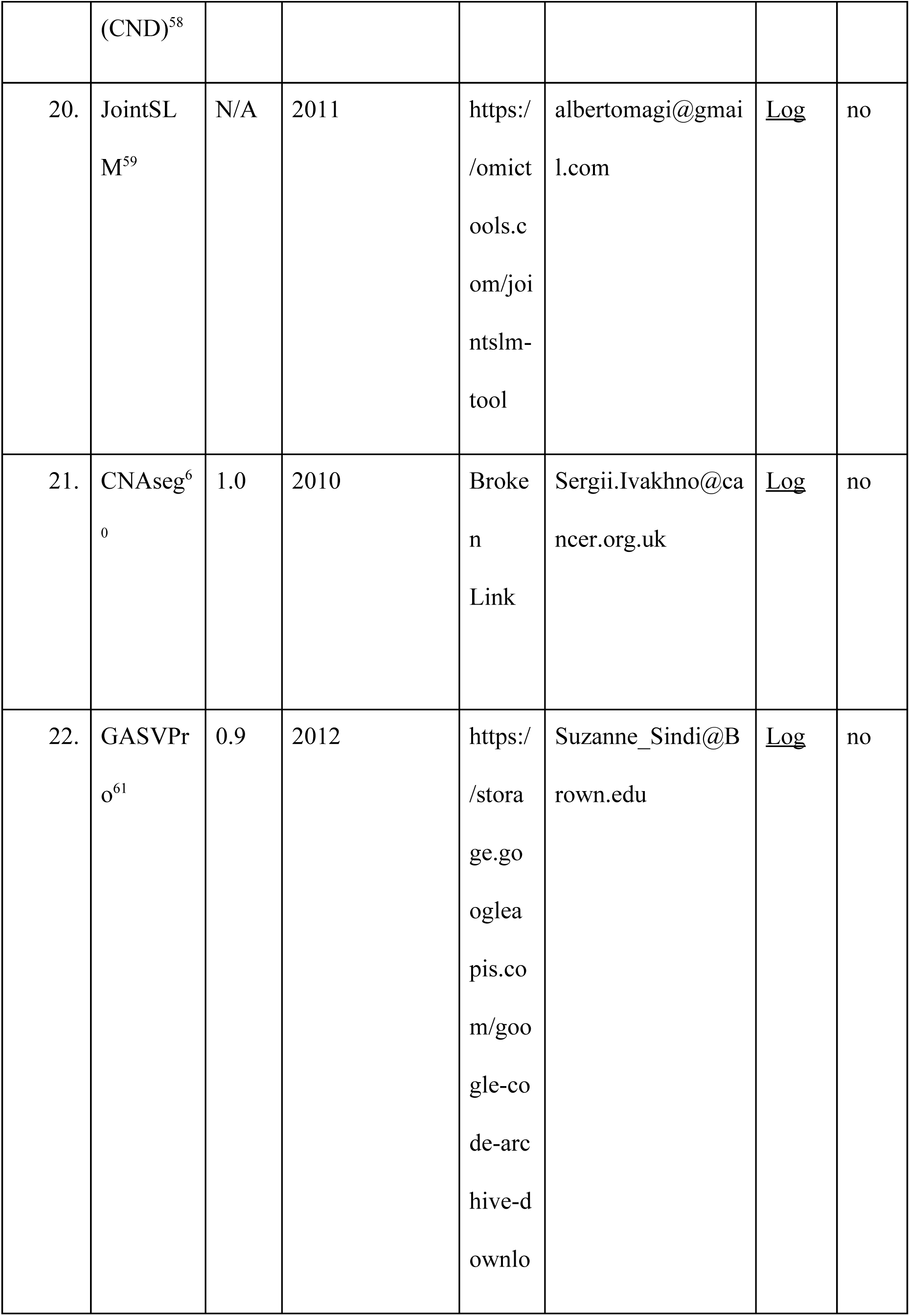
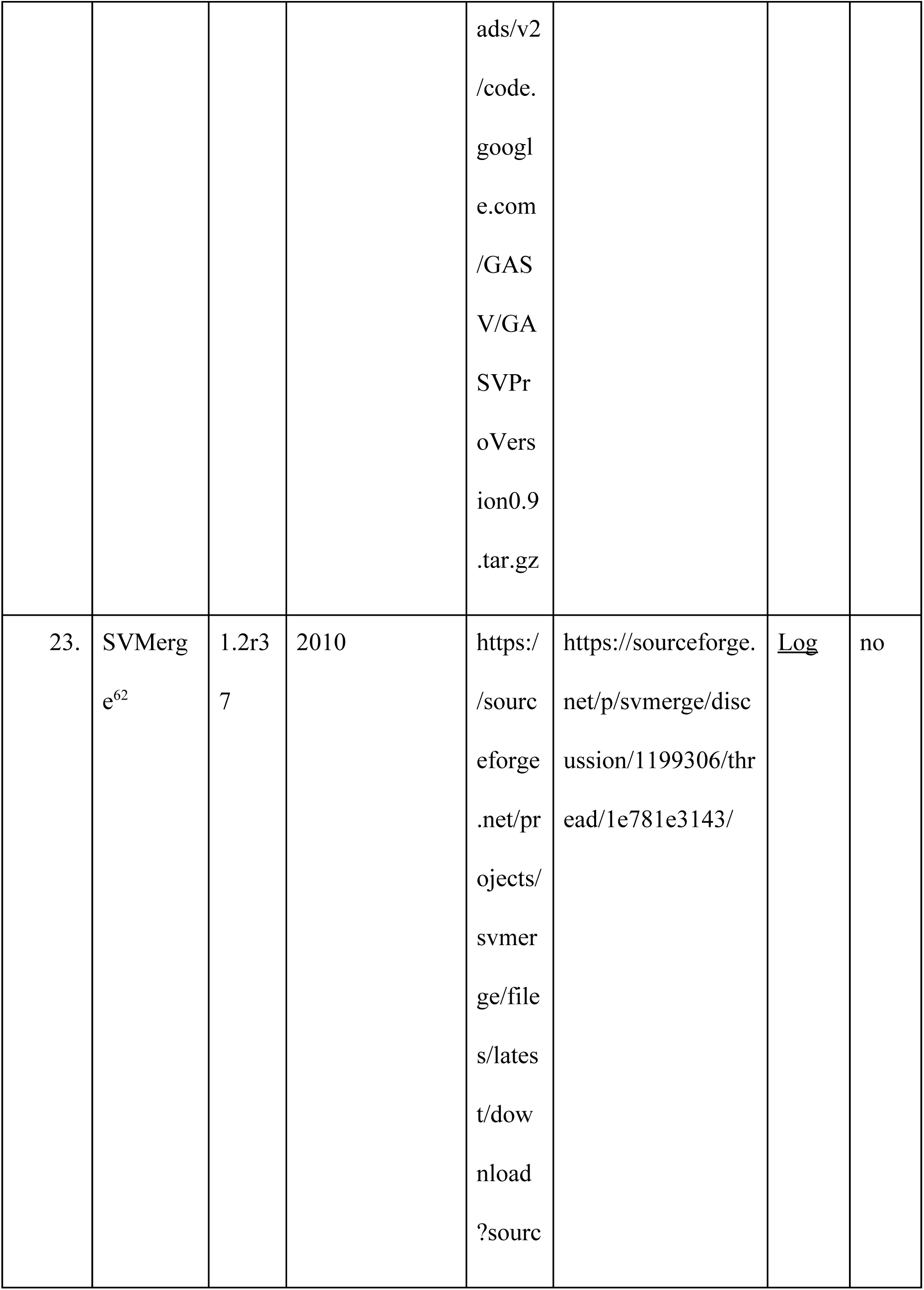
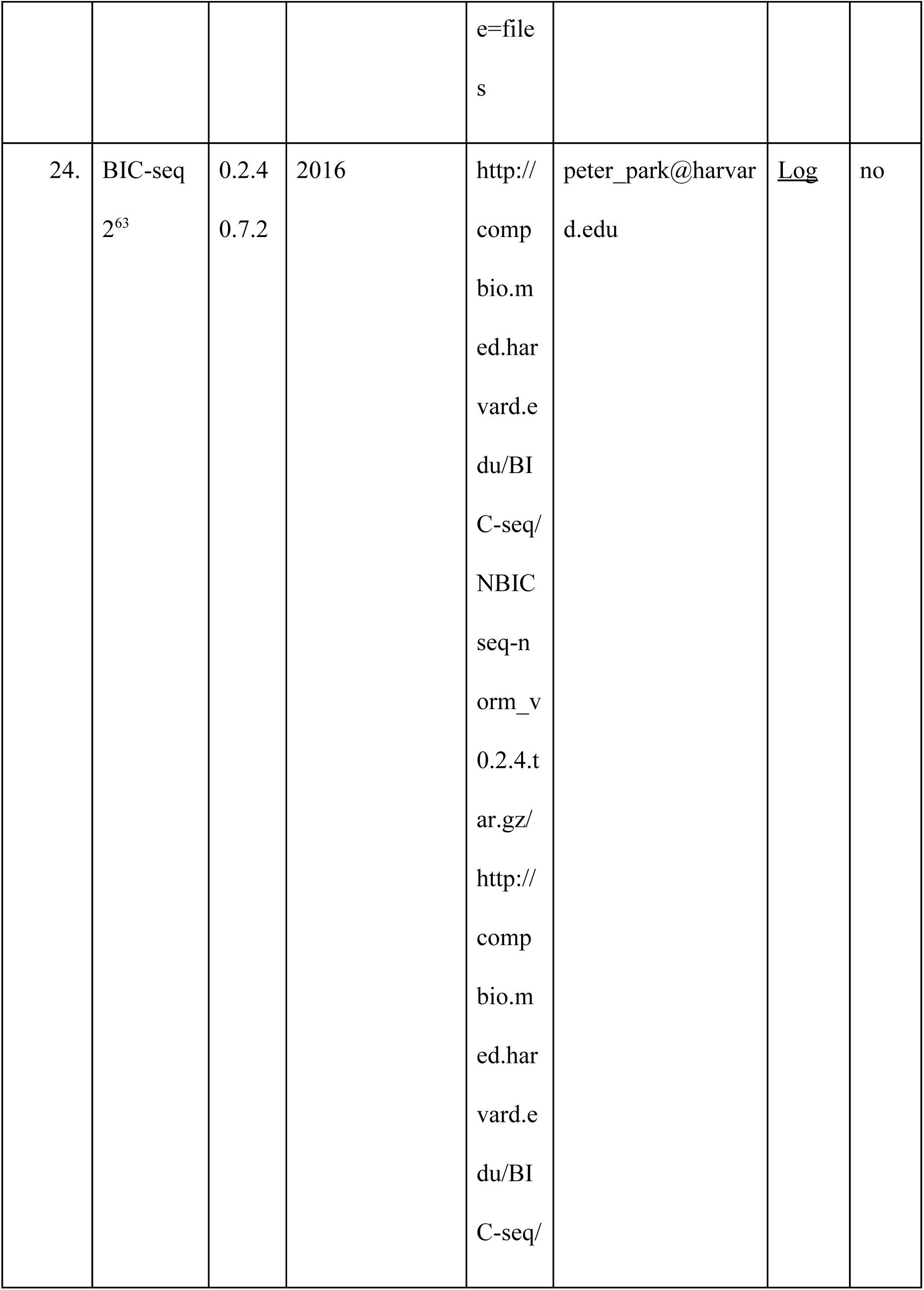
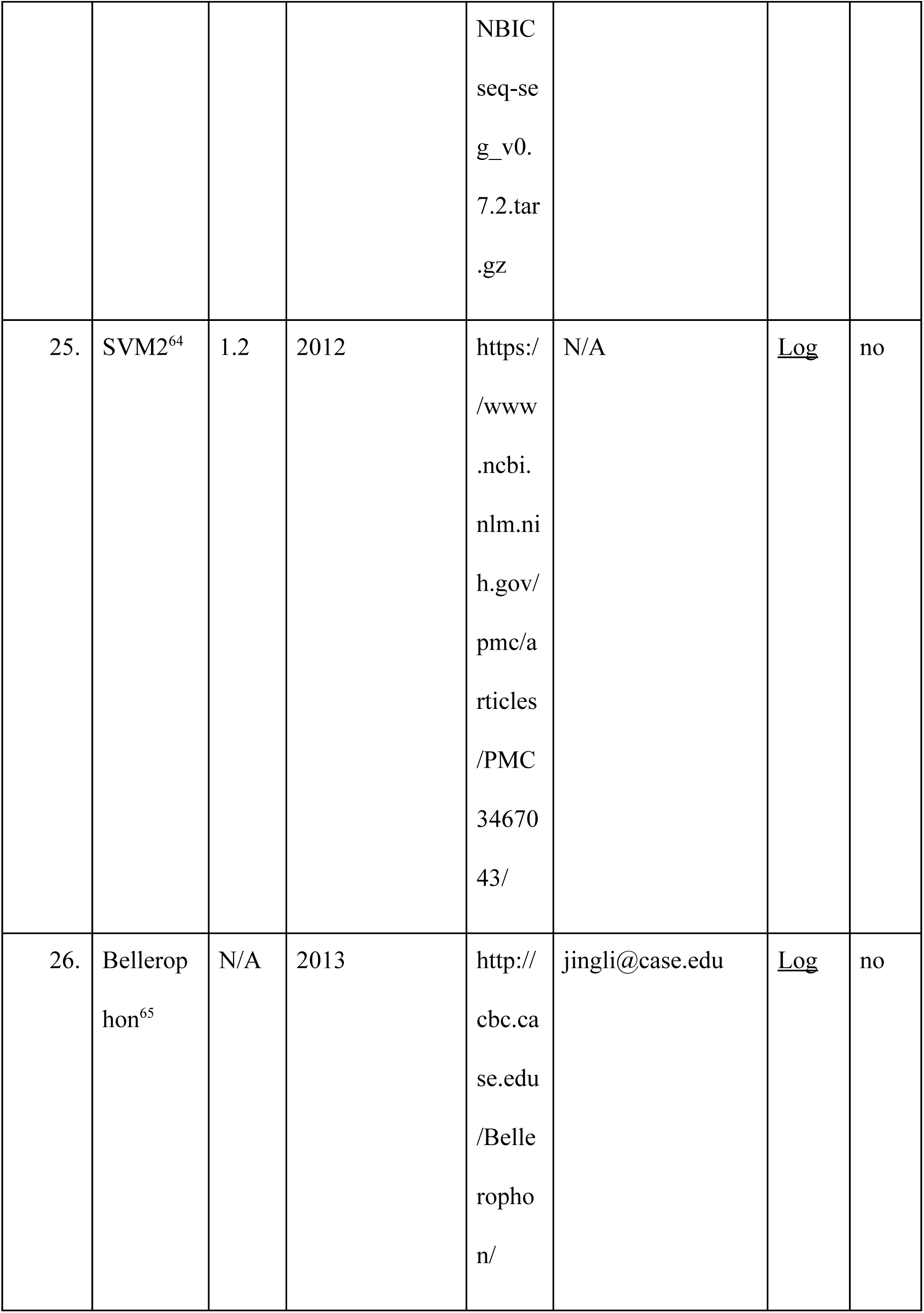
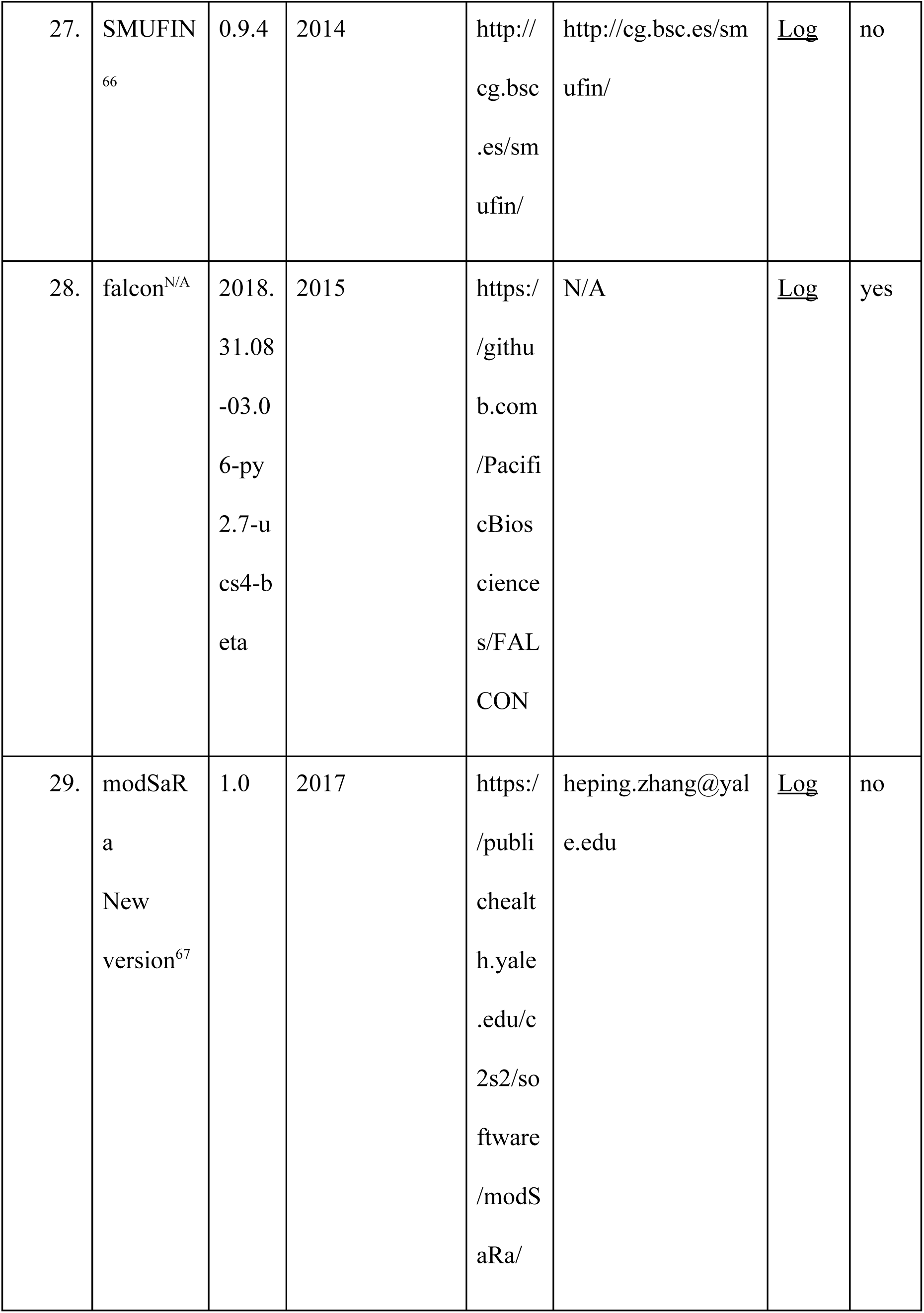
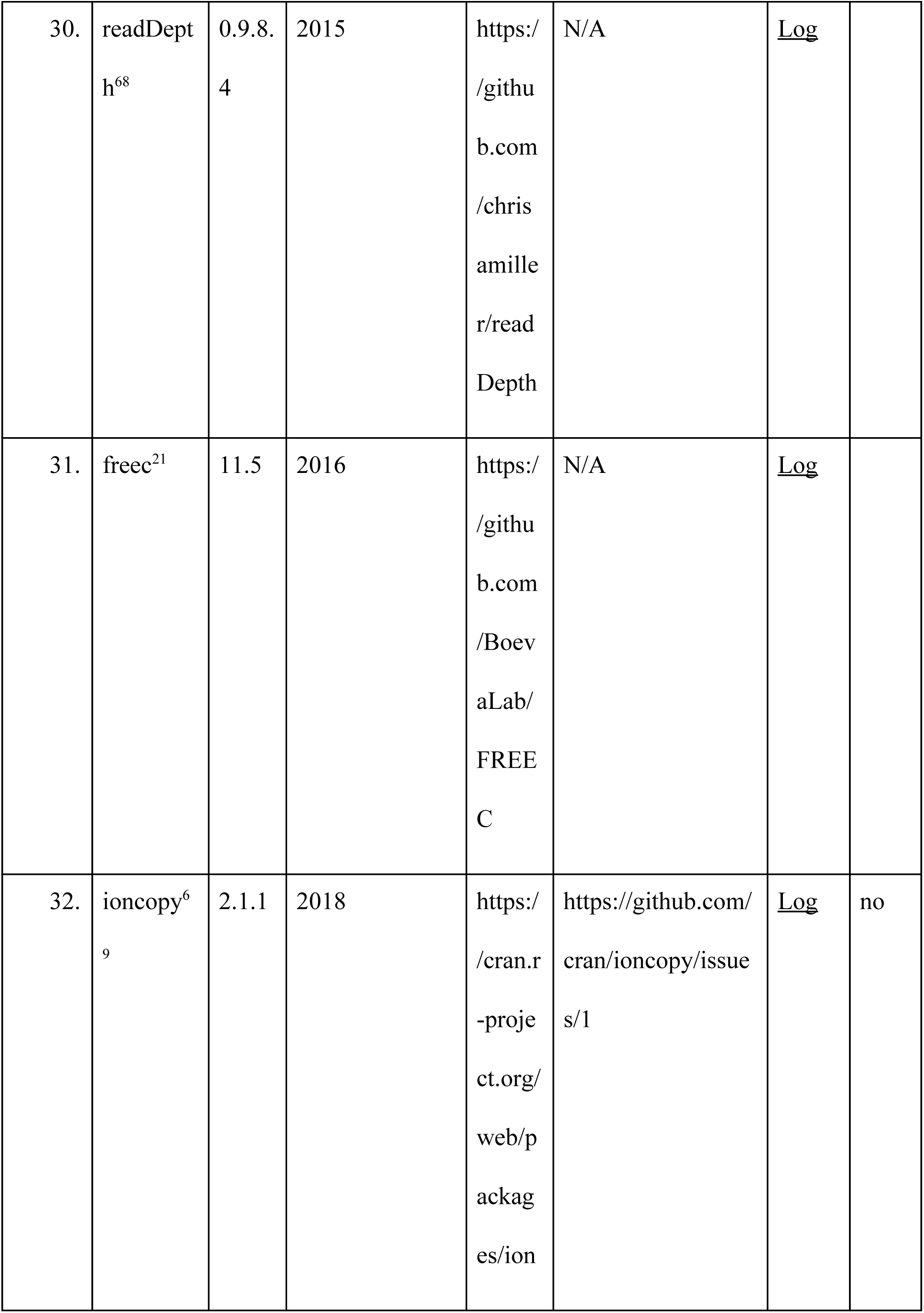
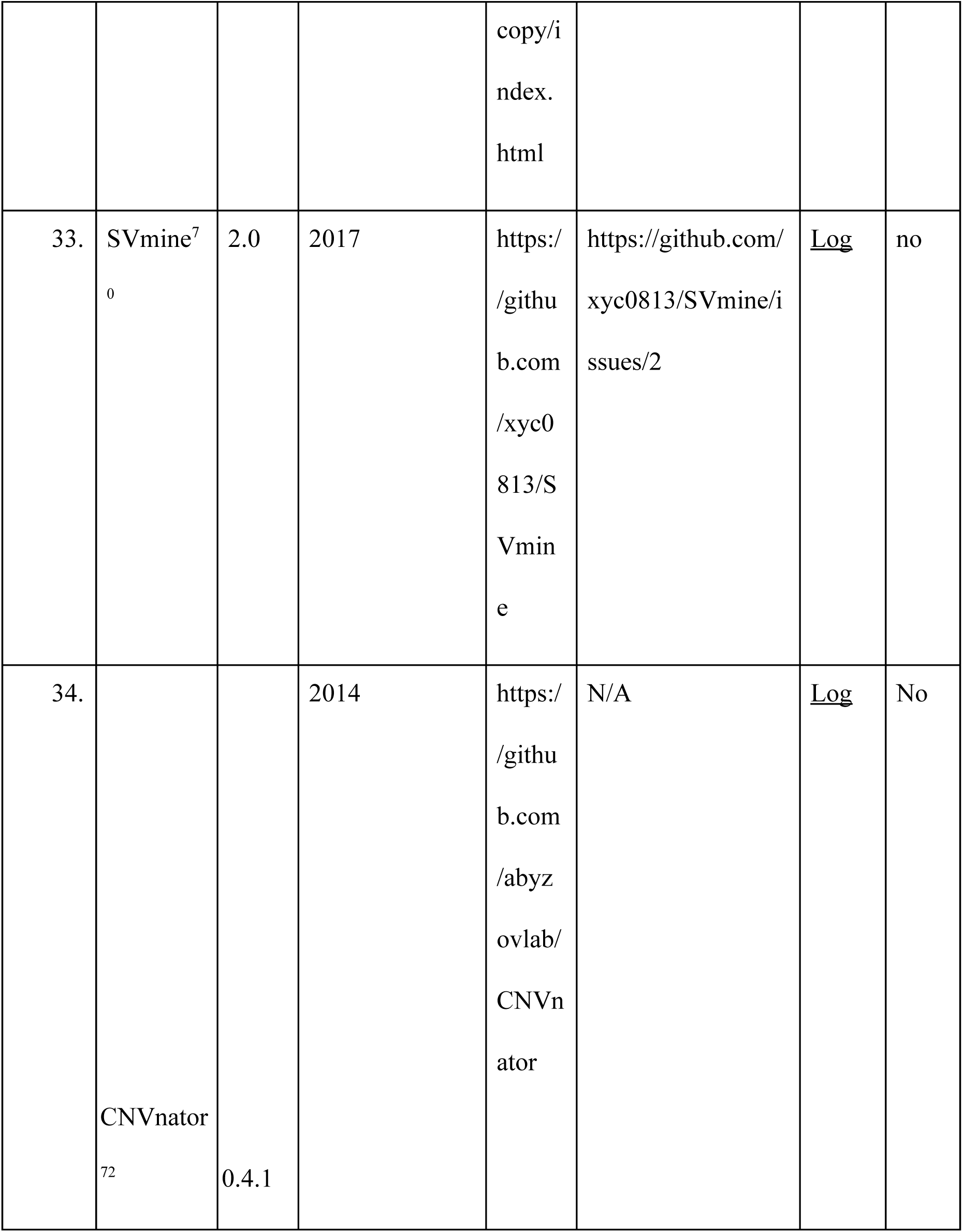
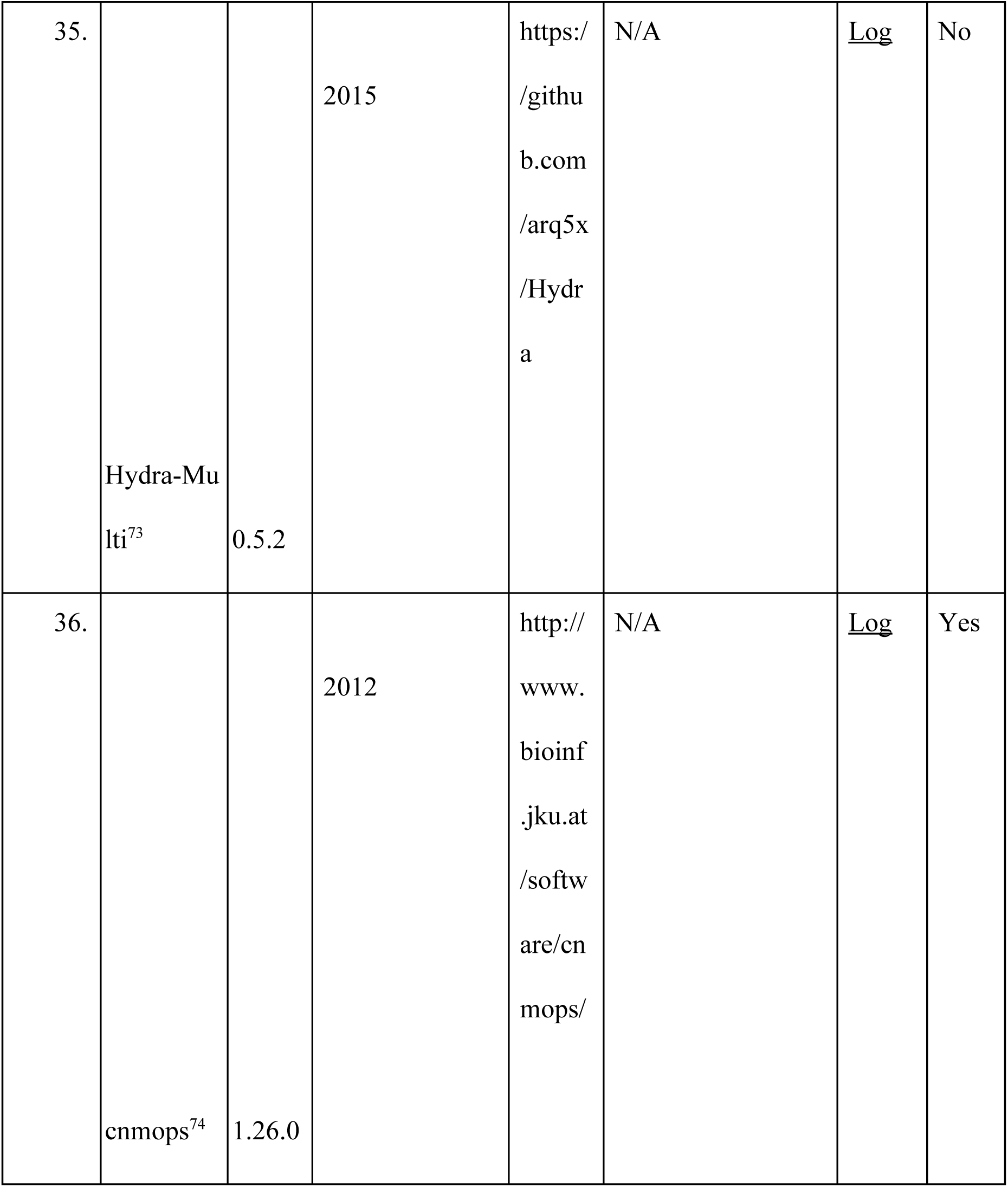
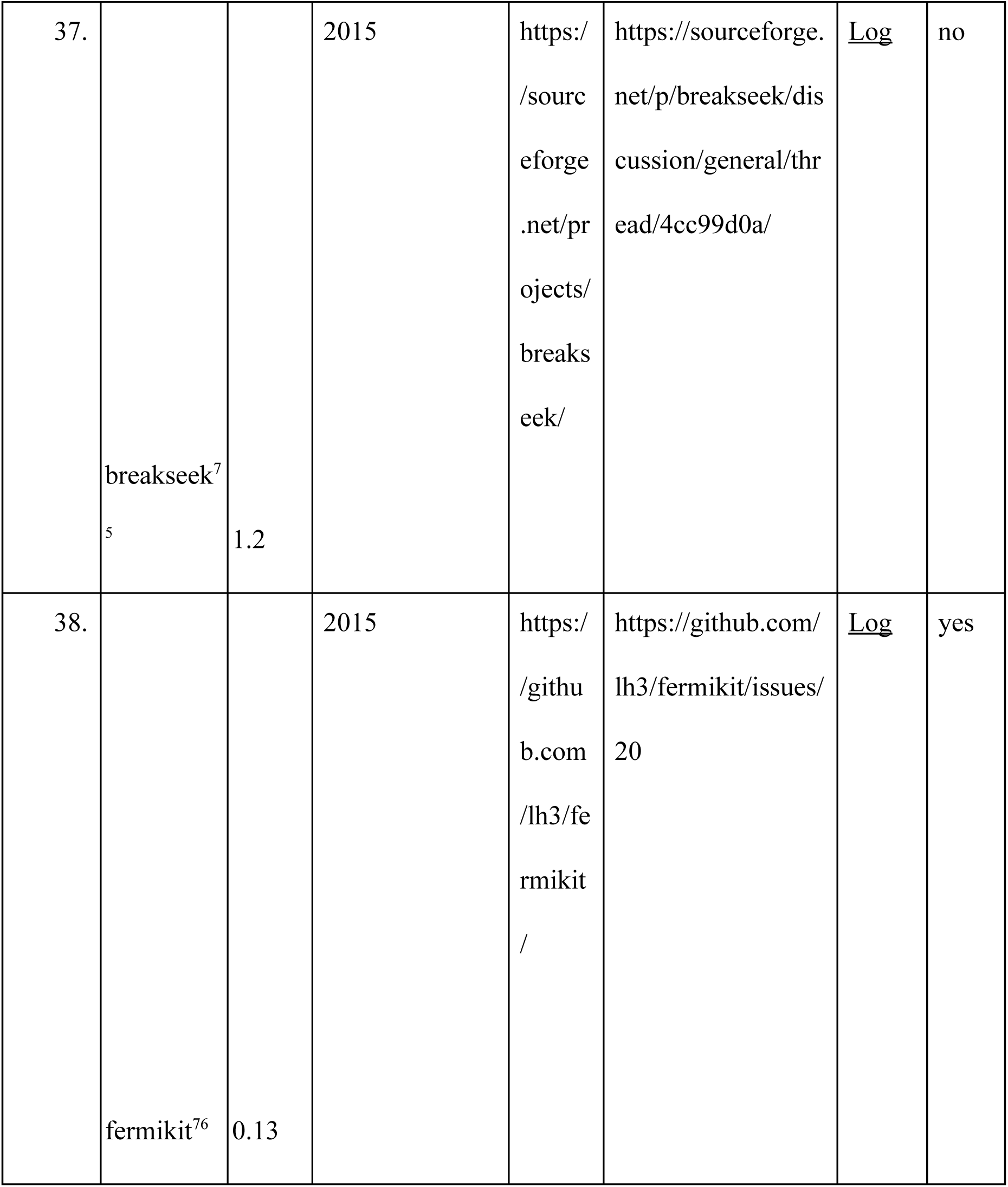
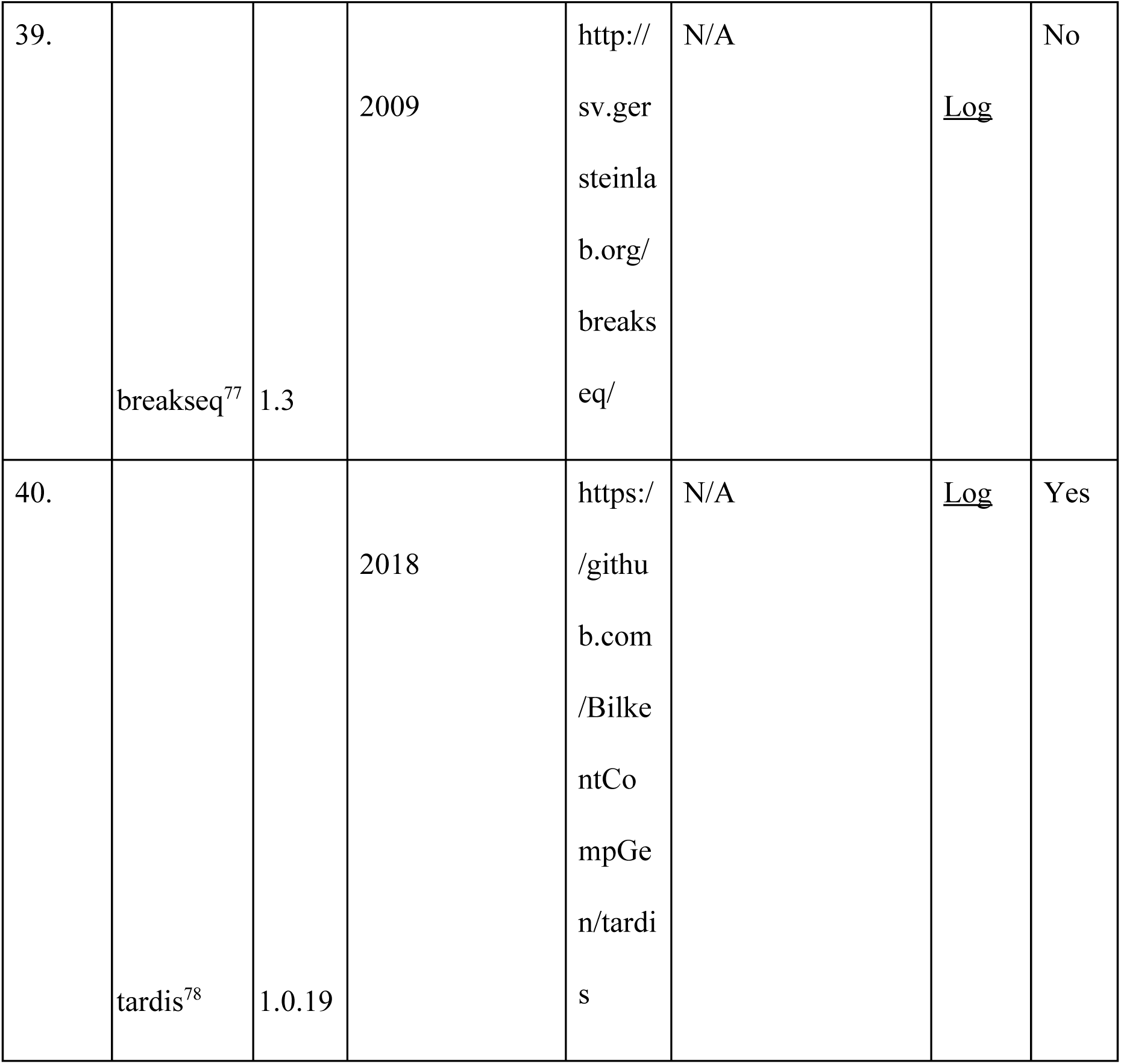
Existing methods to detect structural variants which failed to be installed. If github was available, we have opened the issue. Otherwise, email was sent to the corresponding authors of the paper.

**Table S2.** Gold standard deletion calls from chromosome 19 from 7 inbred mouse strains.

| Strain name | Total number | Average | Minimum | Maximum |
| --- | --- | --- | --- | --- |

|  | **of gold**<br>**standard**<br>**deletions** | **deletion length** | **deletion length** | **deletion length** |
| --- | --- | --- | --- | --- |
| A/J | 533 | 1968 | 15 | 239572 |
| AKR/J | 504 | 1400 | 13 | 24301 |
| BALB/cJ | 545 | 1947 | 15 | 239572 |
| C3H/HeJ | 539 | 1993 | 15 | 239572 |
| DBA/2J | 609 | 1301 | 16 | 24301 |
| LP/J | 483 | 1534 | 13 | 24301 |
| CBA/J | 586 | 1330 | 24 | 24301 |

**Table S3.** Information about WGS data obtained for seven inbred mouse strains.

| **Strain**<br>**name** | **Total**<br>**number**<br>**PE reads** | **Number**<br>**of reads**<br>**mapped** | **Number**<br>**of reads**<br>**mapped to** | **Number**<br>**of**<br>**discordant** | **Number**<br>**of**<br>**multi-map** | **Average**<br>**coverage**<br>**(chr19)** |
| --- | --- | --- | --- | --- | --- | --- |

|  |  | **reads** | **chr 19** | **reads**<br>**(chr19)** | **ped reads**<br>**(chr19)** |  |
| --- | --- | --- | --- | --- | --- | --- |
| A/J | 691798369 | 15097330 | 14903814 | 97469 | 14765316 | 48.52 |
| AKR/J | 758992189 | 16205276 | 15814814 | 66938 | 16205276 | 51.49 |
| BALB/cJ | 828502183 | 17292403 | 17093309 | 106915 | 17292403 | 55.65 |
| C3H/HeJ | 828898555 | 17293104 | 17093572 | 77254 | 17293104 | 55.65 |
| CBA/J | 736613128 | 15517147 | 15343318 | 67076 | 15517147 | 49.95 |
| DBA/2J | 738506916 | 14860907 | 14690187 | 72075 | 14860907 | 47.83 |
| LP/J | 703108275 | 14331024 | 14176319 | 60233 | 14331024 | 46.15 |
| Total | 528641961<br>5 | 110597191 | 109,115,33<br>3 | 547960 | 110265177 | 50.75 |

**Table S4.** Tools excluded from analysis.

| **S.N**<br>**o** | **Tool name** | **Version** | **Year** | **Webpage** | **Reason the**<br>**tools was**<br>**excluded** |
| --- | --- | --- | --- | --- | --- |
| 1. | GATK<sup>90</sup> | 4.1.6.0 | 2010 | https://github.com/broadinstitute/gatk | designed to detect only small SVs (less than 50 bp in length) |
| 2. | rSW-seq<sup>88</sup> | N/A | 2010 | https://bioinformaticshomepage.com/tools/cnv/descriptions/rSW-seq.html | designed to detect SVs in tumor-normal samples |
| 3. | Variation Hunter<sup>83</sup> | 0.04 | 2010 | http://variationhunter.sourceforge.net/ | replaced with TARDIS |
| 4. | inGAP-SV<sup>41</sup> | 3.1.1 | 2011 | http://ingap.sourceforge.net/ | utilizes a GUI, hence command line tasks<br>with many<br>datasets<br>would not be<br>viable for<br>benchmarkin<br>g |
| 5. | COPS<sup>87</sup> | N/A | 2012 | ftp://115.119.160.213<br>username:<br>cops<br>password:cop<br>s | designed to<br>detect SVs in<br>tumor-normal<br>samples |
| 6. | Magnolya<sup>27</sup> | 0.15 | 2012 | https://source<br>forge.net/proj<br>ects/magnoly<br>a/files/ | unable to<br>process<br>aligned WGS<br>data |
| 7. | Varscan<sup>92</sup> | 2.3.9 | 2012 | https://source<br>forge.net/proj<br>ects/varscan/f<br>iles/ | designed to<br>detect only<br>small SVs (less than 50 bp in length) |
| 8. | Patchwork<sup>86</sup> | 2.4 | 2013 | http://patchwork.r-forge.r-project.org/ | designed to detect SVs in tumor-normal samples |
| 9. | PBHoney-NGM<sup>82</sup> | 13.10 | 2014 | https://sourceforge.net/projects/pb-jelly/ | uses long-read sequencing data |
| 10. | Platypus<sup>91</sup> | 0.8.1 | 2014 | https://github.com/andyrimmer/Platypus | designed to detect only small SVs (less than 50 bp in length) |
| 11. | seqCBS<sup>89</sup> | 1.2.1 | 2014 | https://cran.r-project.org/web/packages/seqCBS/index html | designed to detect SVs in tumor-normal samples |
| 12. | Socrates<sup>85</sup> | 1.1 | 2014 | http://bioinf.<br>wehi.edu.au/s<br>ocrates/<br>https://github.<br>com/jibsch/so<br>crates | replaced with GRIDSS, as the toolmaker who developed both suggested GRIDSS for SV detection |
| 13. | Hydra<sup>84</sup> | N/A | 2015 | http://web.en<br>gr.illinois.edu<br>/~mkim158/H<br>yDRA.zip | replaced with LUMPY |
| 14. | Bic-seq<sup>63</sup> | 0.2.4 | 2016 | http://compbi<br>o.med.harvar<br>d.edu/BIC-se<br>q/ | designed to detect SVs in tumor-normal samples |
| 15. | Control-FREEC<sup>21</sup> | 11.5 | 2016 | https://github.com/BoevaLab/FREEC | It only accepts only GZ format and no longer supports bam files (starting with from version 8.0) |
| 16. | NanoSV<sup>81</sup> | 1.2.4 | 2017 | https://github.com/mroosmalen/nanosv | uses long-read sequencing data |
| 17. | Sniffles<sup>96</sup> | 1.0.8 | 2018 | https://github.com/fritzsedlazeck/Sniffles | Designed for long reads |

**Table S5.**
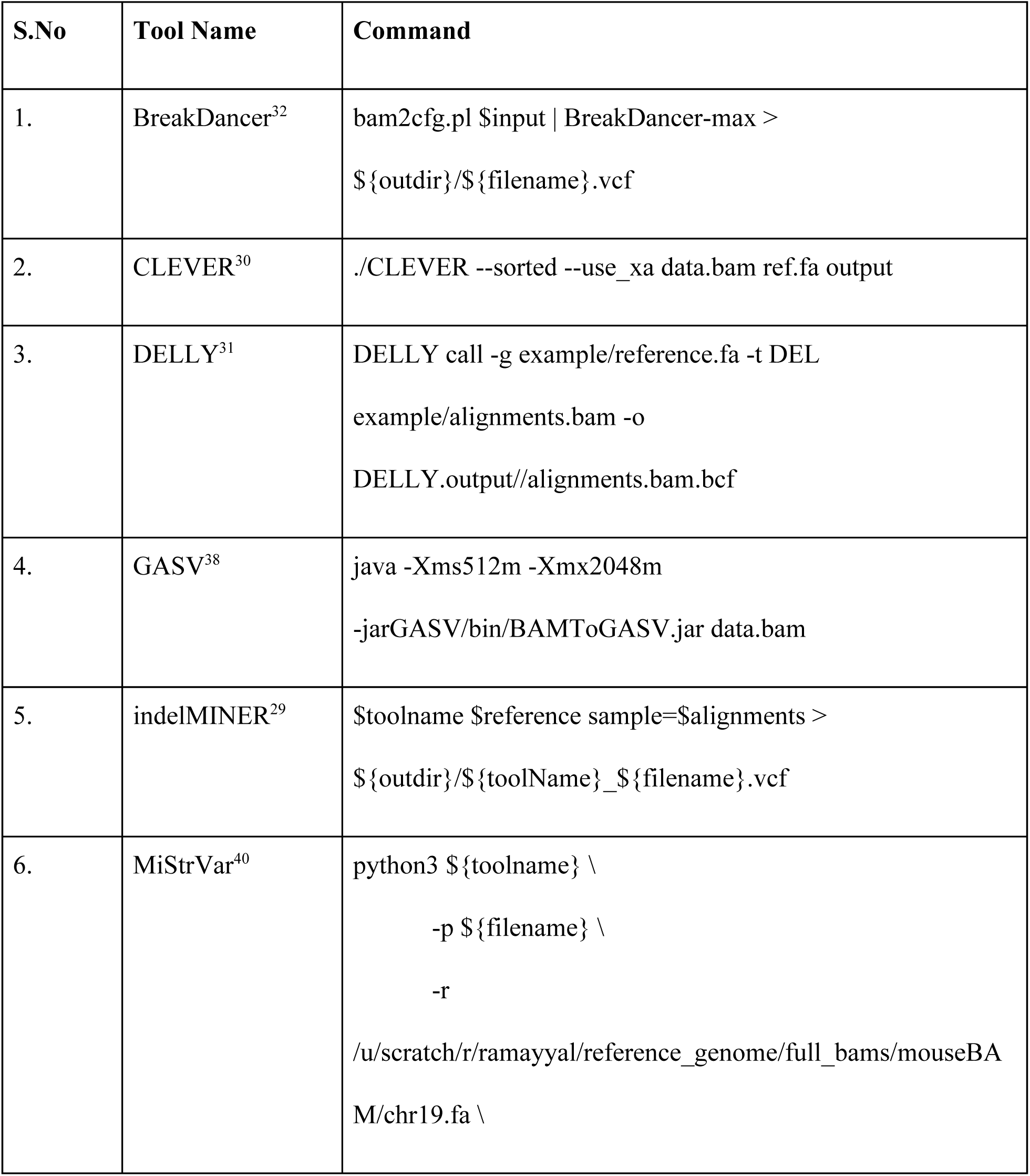
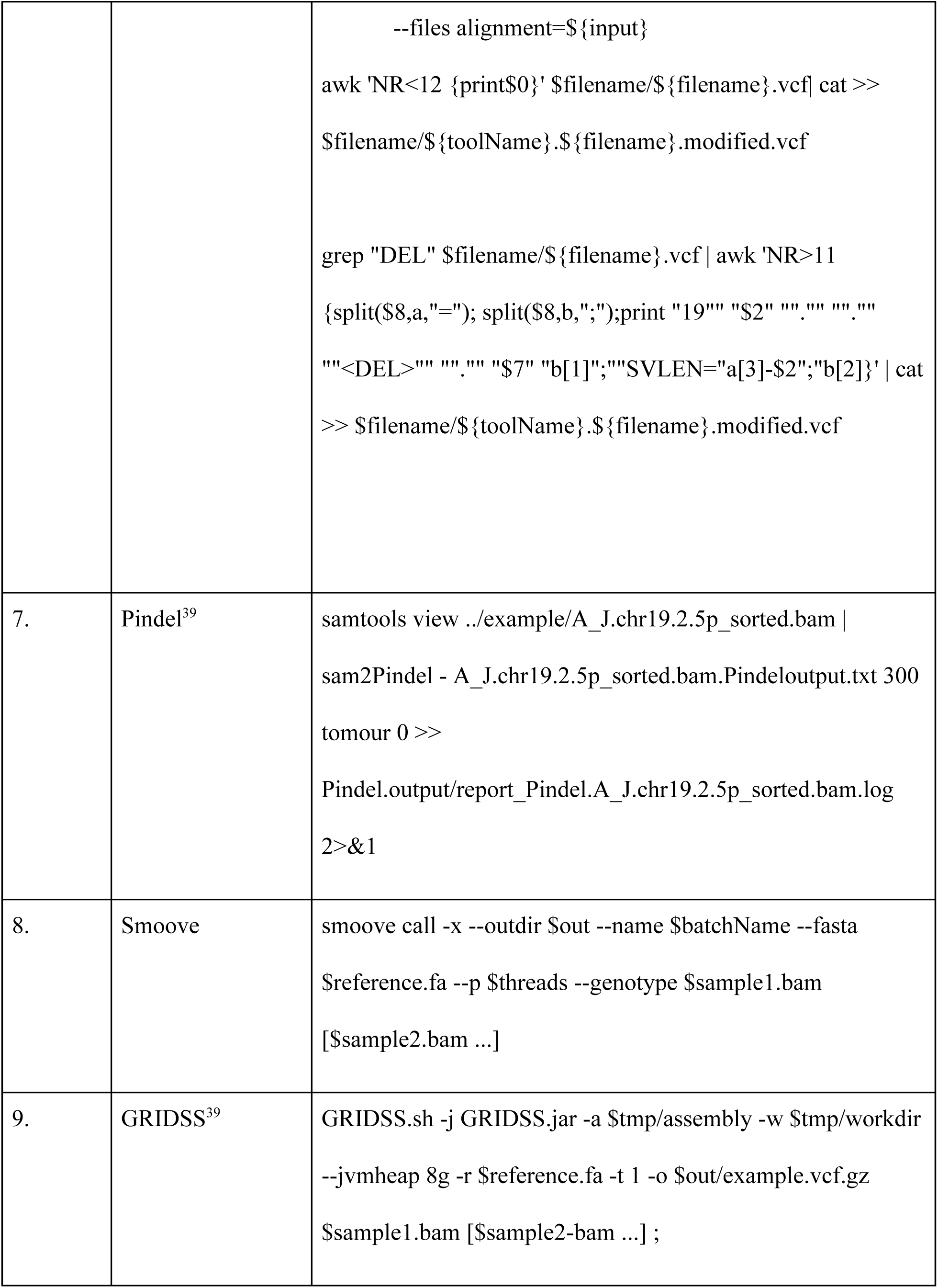
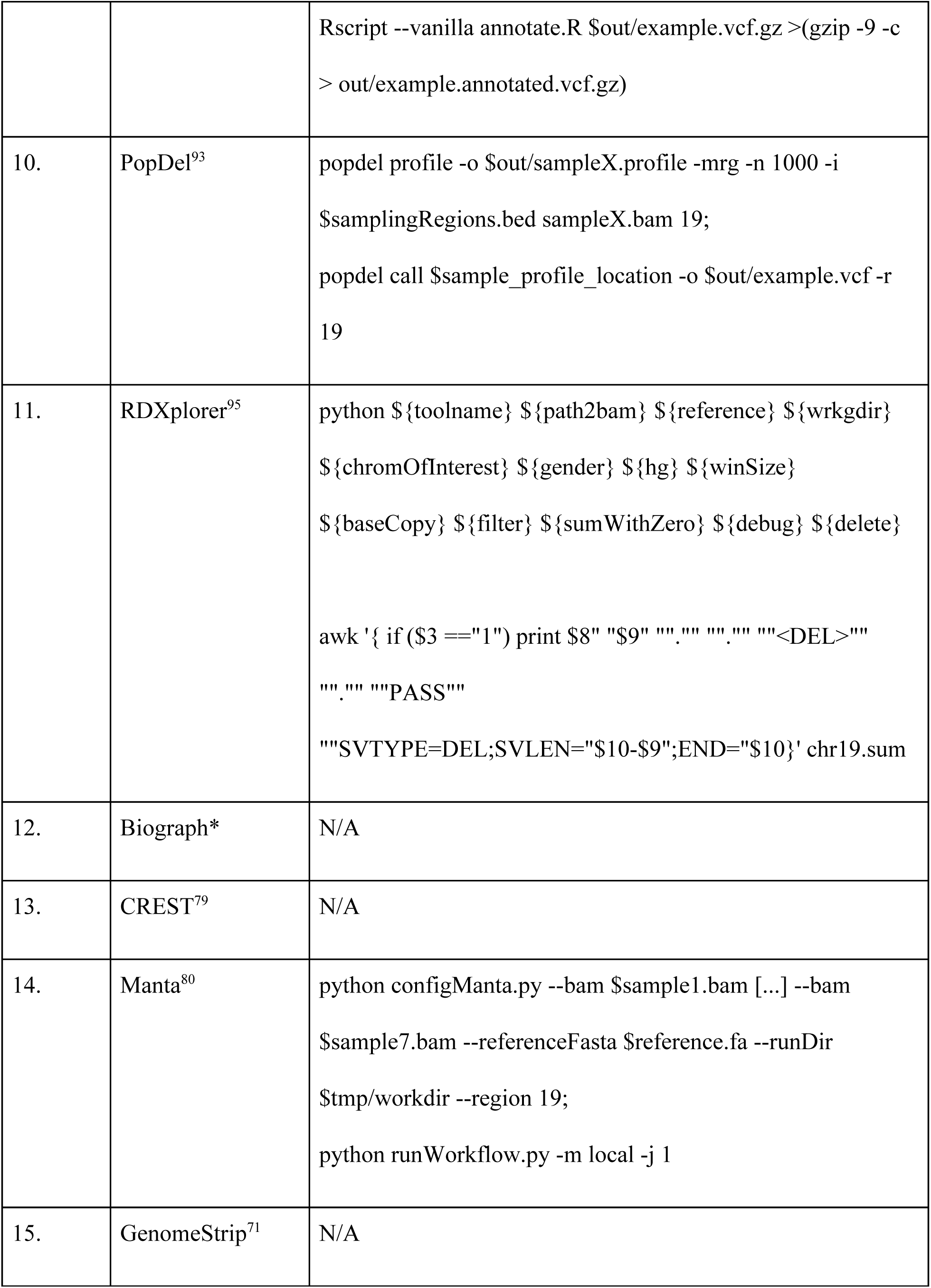
Commands used to run the tools and produce an output vcf file with predicted deletions using a reference (FASTA format) and aligned reads (BAM format) file as input arguments.

**Table S6.** Output format of different tools

| S.No | Tool Name | Output type | Description of the format |
| --- | --- | --- | --- |
| 1. | BreakDancer<sup>32</sup> | Custom | Start, end, type, length |
| 2. | CLEVER<sup>30</sup> | Custom | VCFv4.1 |
| 3. | DELLY<sup>31</sup> | Custom | VCFv4.2 |
| 4. | GASV<sup>38</sup> | Custom | Start, end, type |
| 5. | indelMINER<sup>29</sup> | VCF | VCFv4.1 |
| 6. | LUMPY<sup>94</sup> | VCF | VCFv4.2 |
| 7. | MiStrVar<sup>40</sup> | VCF | VCFv4.2 |
| 8. | Pindel<sup>39</sup> | Custom | Start,end |
| 9. | BioGraph* | VCF | VCFv4.2 |
| 10. | GRIDSS<sup>39</sup> | VCF | VCFv4.2 |
| 11. | Popdel<sup>93</sup> | VCF | VCFv4.3 |
| 12. | RDXplorer<sup>95</sup> | Custom | Start, end, length |
| 13. | CREST<sup>79</sup> | Custom | Start, end, type |
| 14. | Manta<sup>80</sup> | VCF | VCFv4.1 |
| 15. | genomeSTRIP<sup>71</sup> | VCF | VCFv4.2 |

## Supplementary Figures

**Figure S1.**
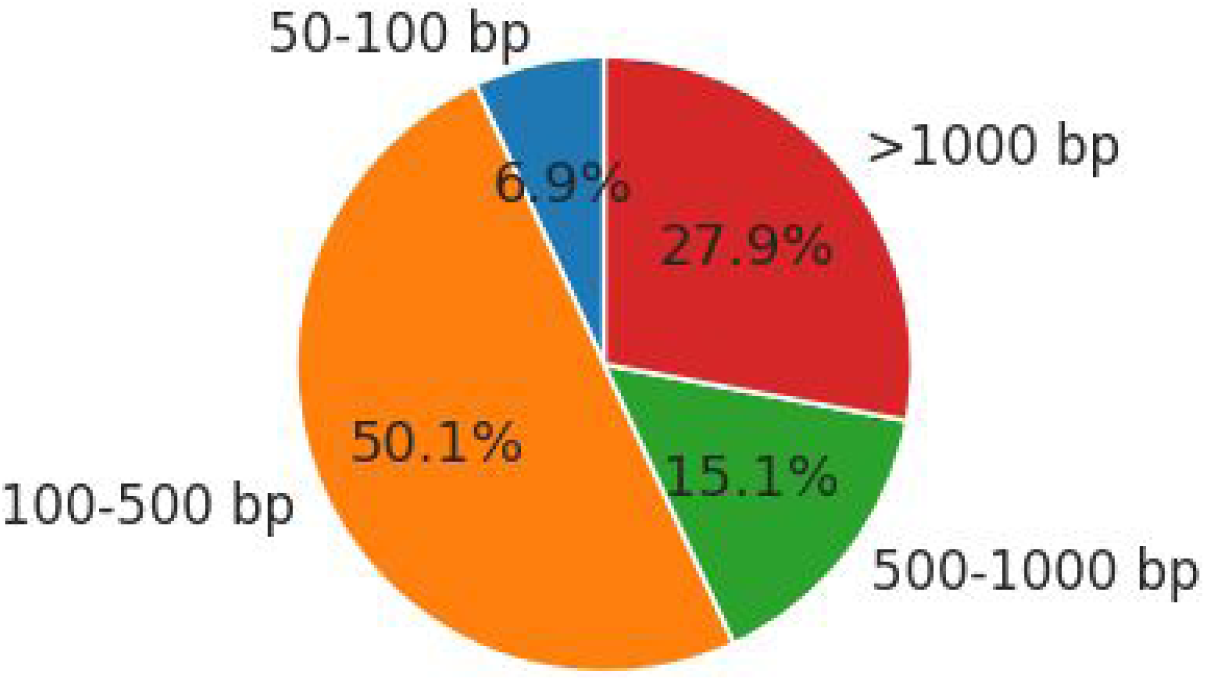
Total number of deletions across seven mouse strains split across categories defined based on deletion length

**Figure S2.**
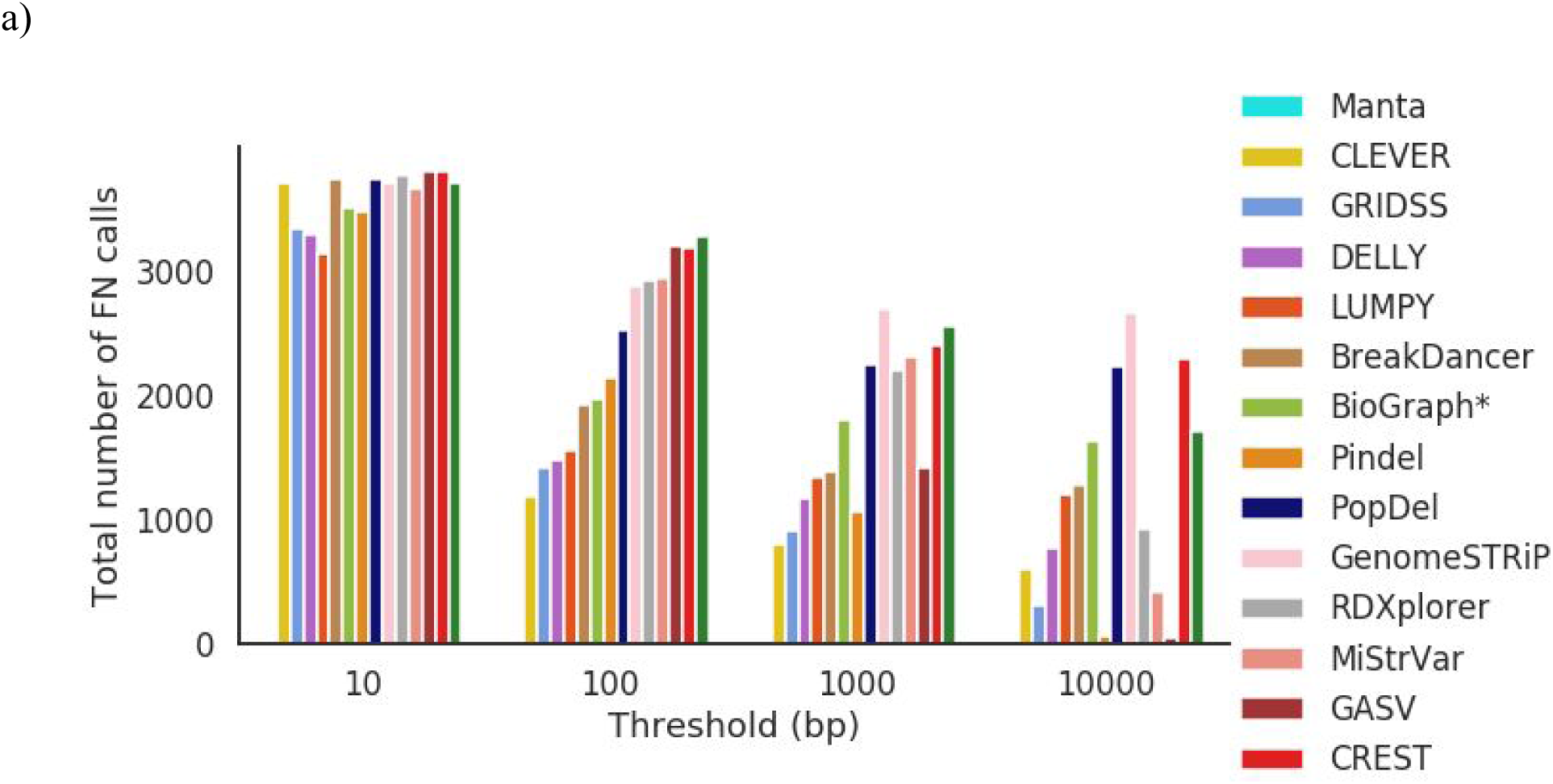
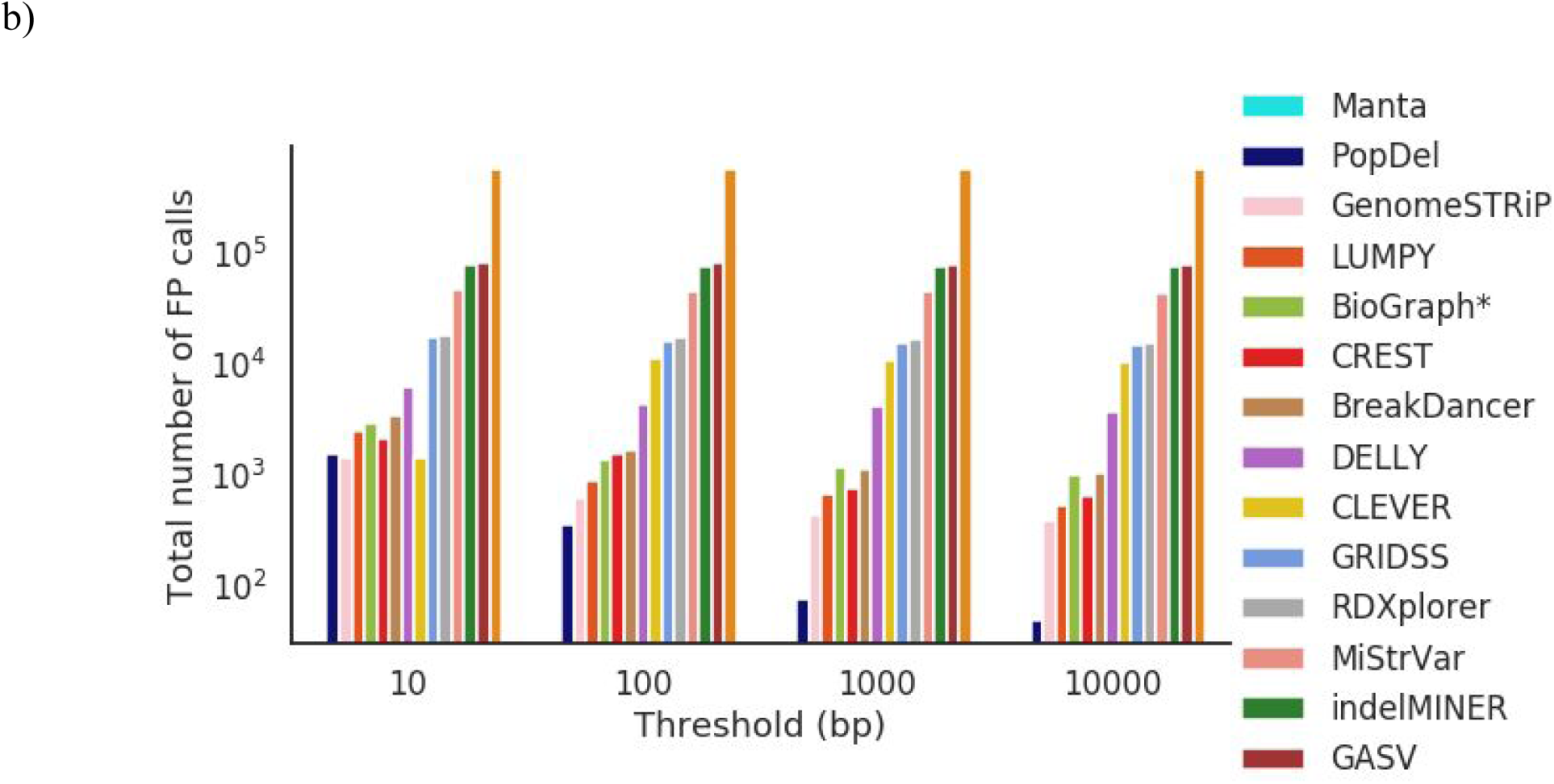
(a) Total number of FN calls for different error thresholds across 7 different mouse strains (b) Total number of FP calls for different error thresholds across 7 different mouse strains

**Figure S3.**
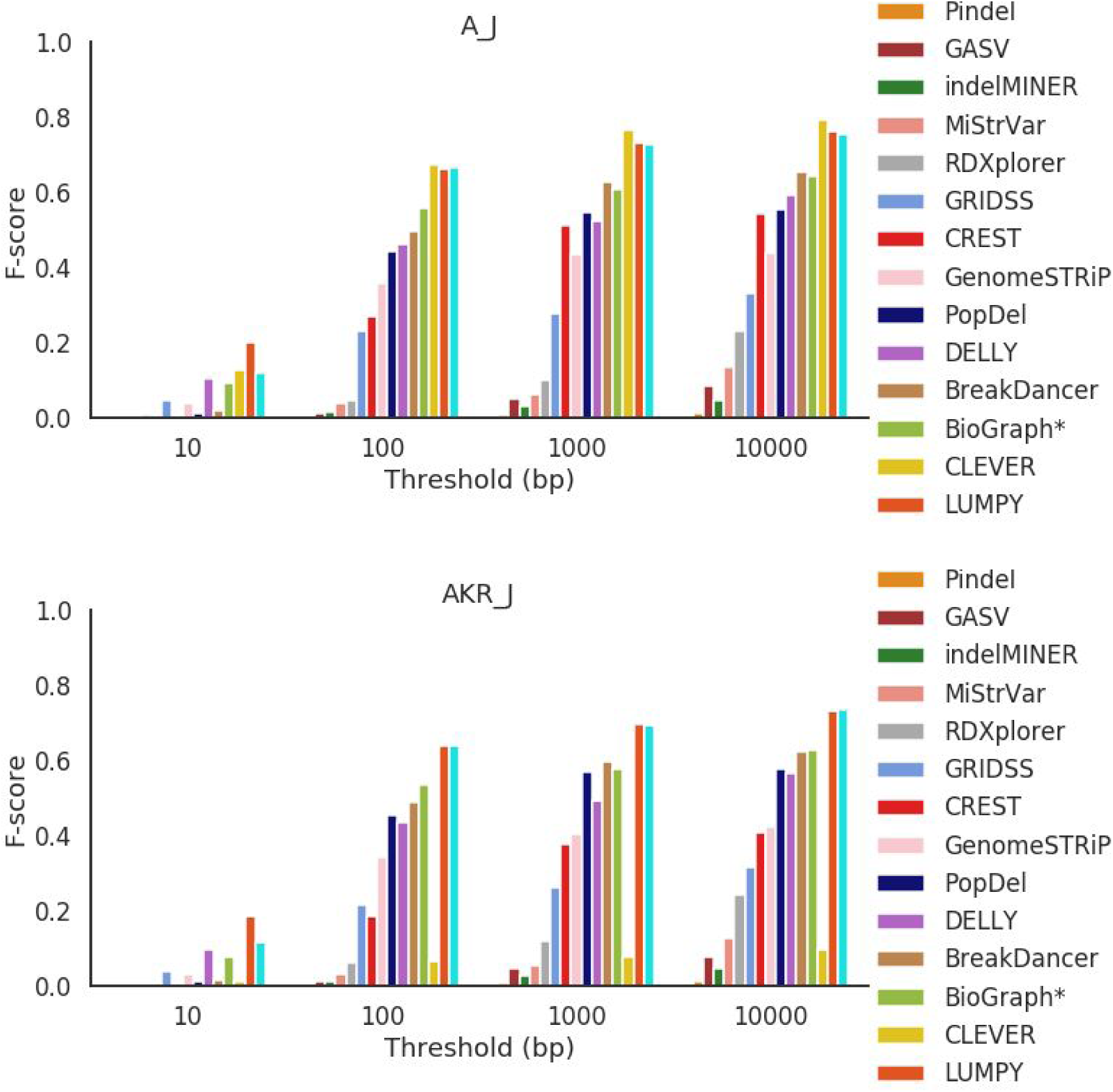
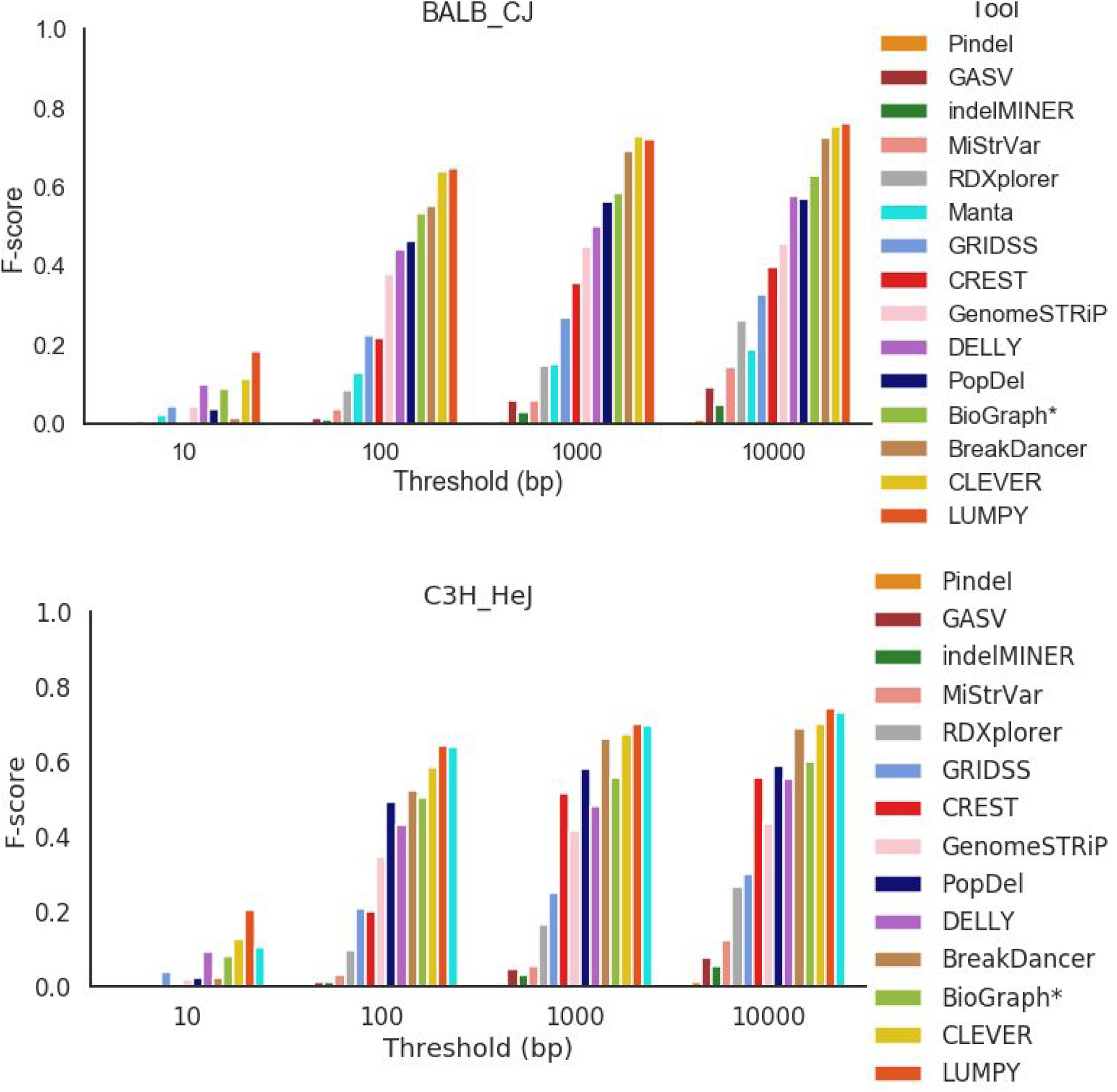
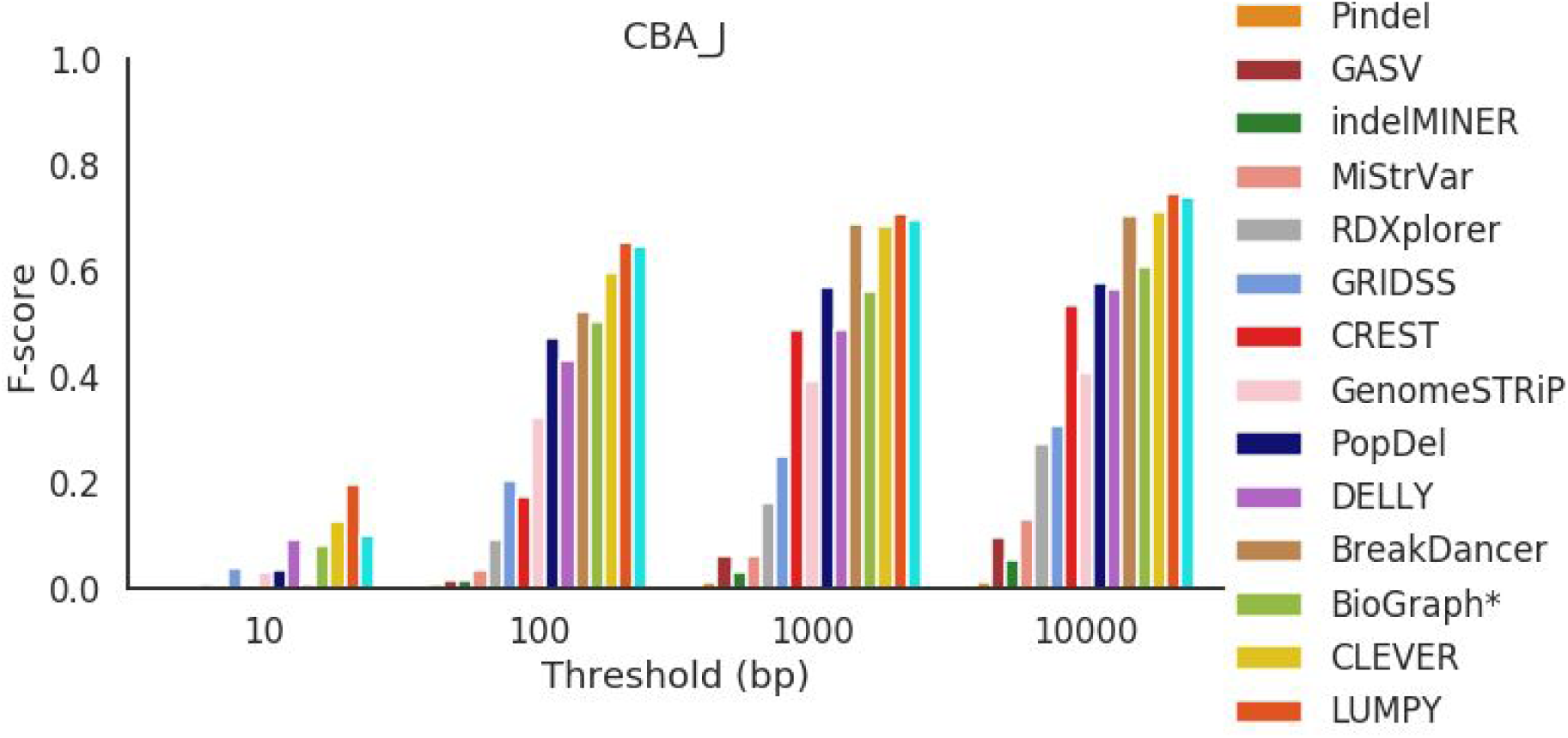
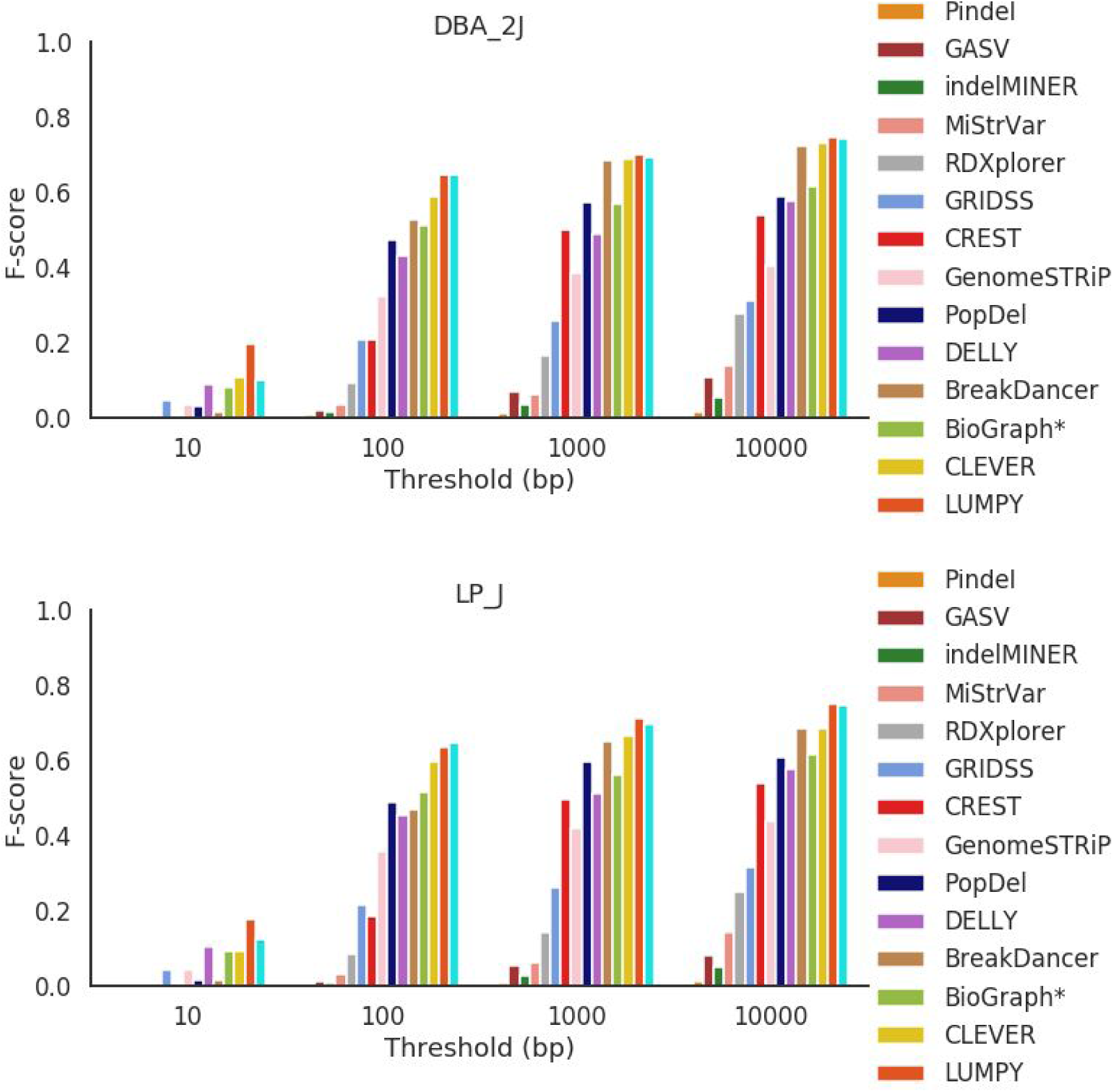
F-score presented for each mouse strain across all error thresholds.

**Figure S4.**
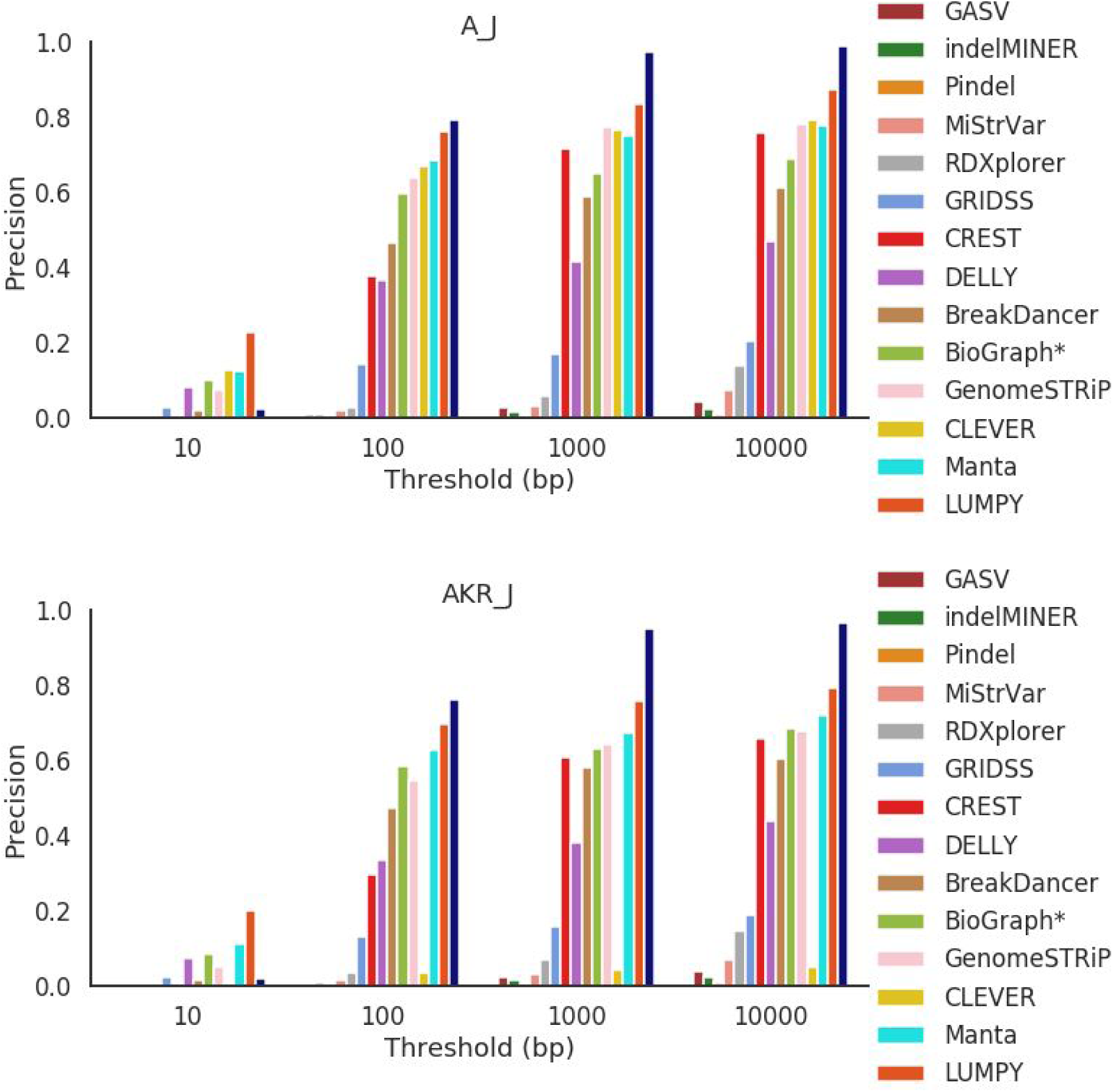
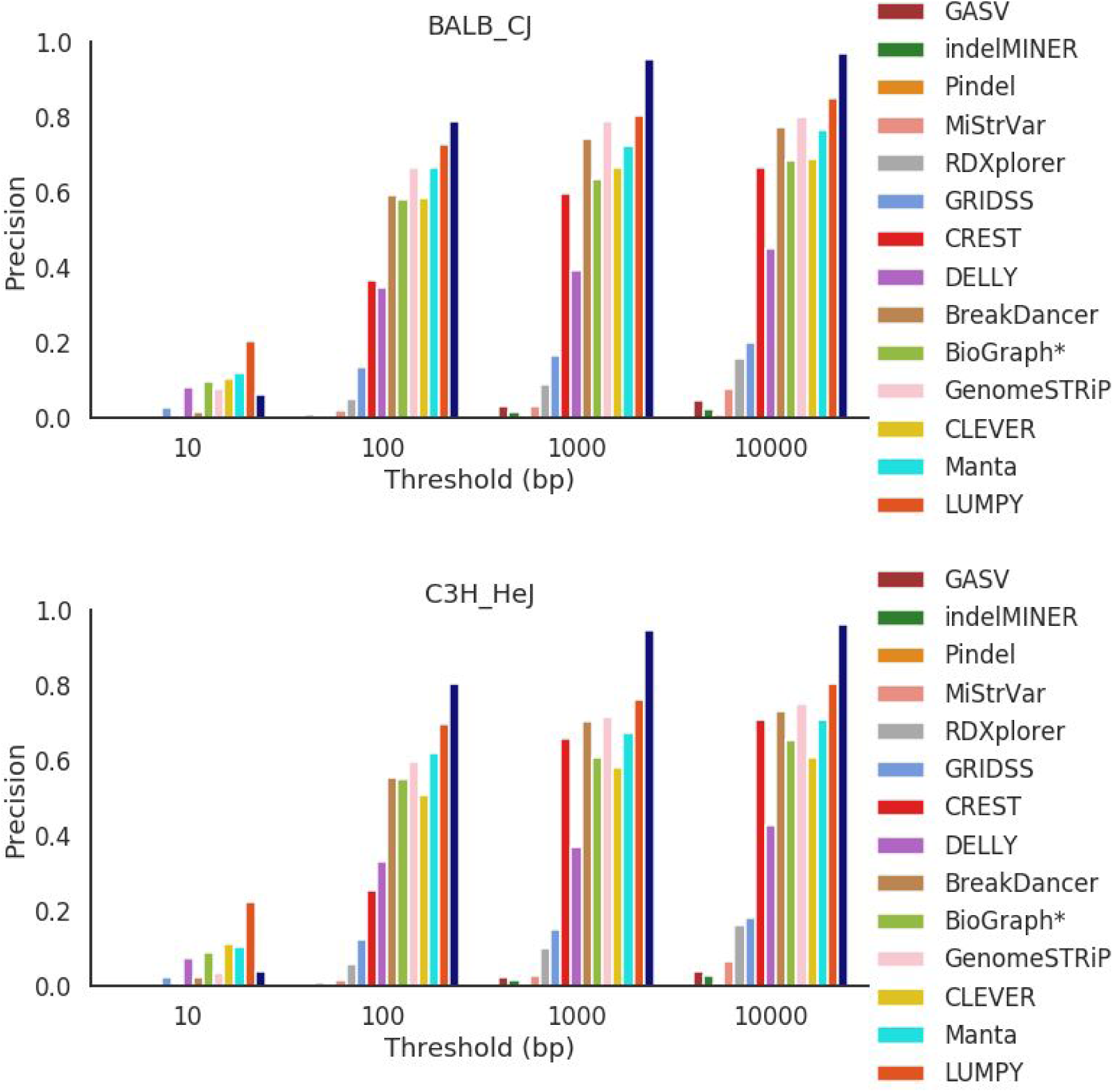
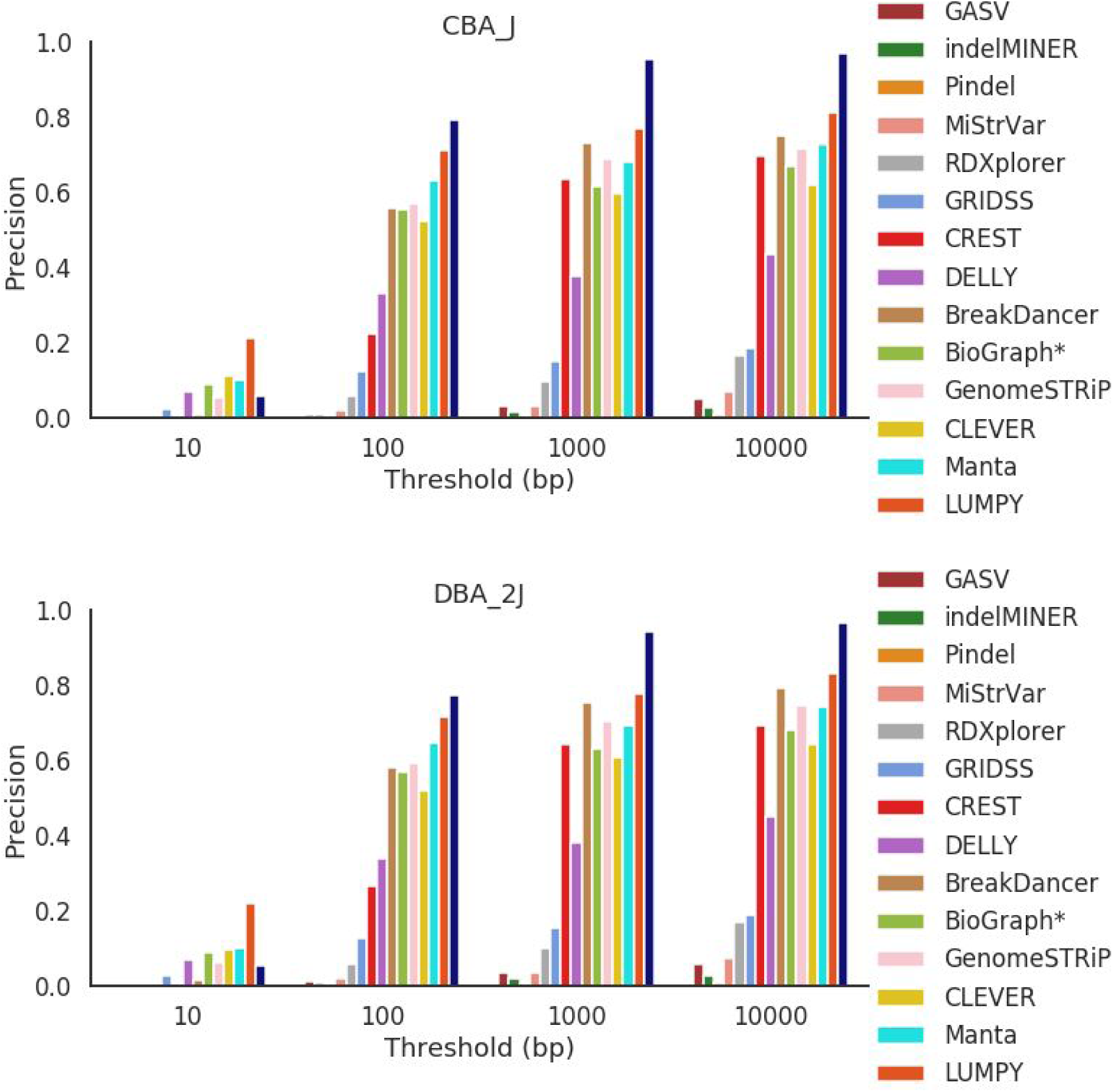
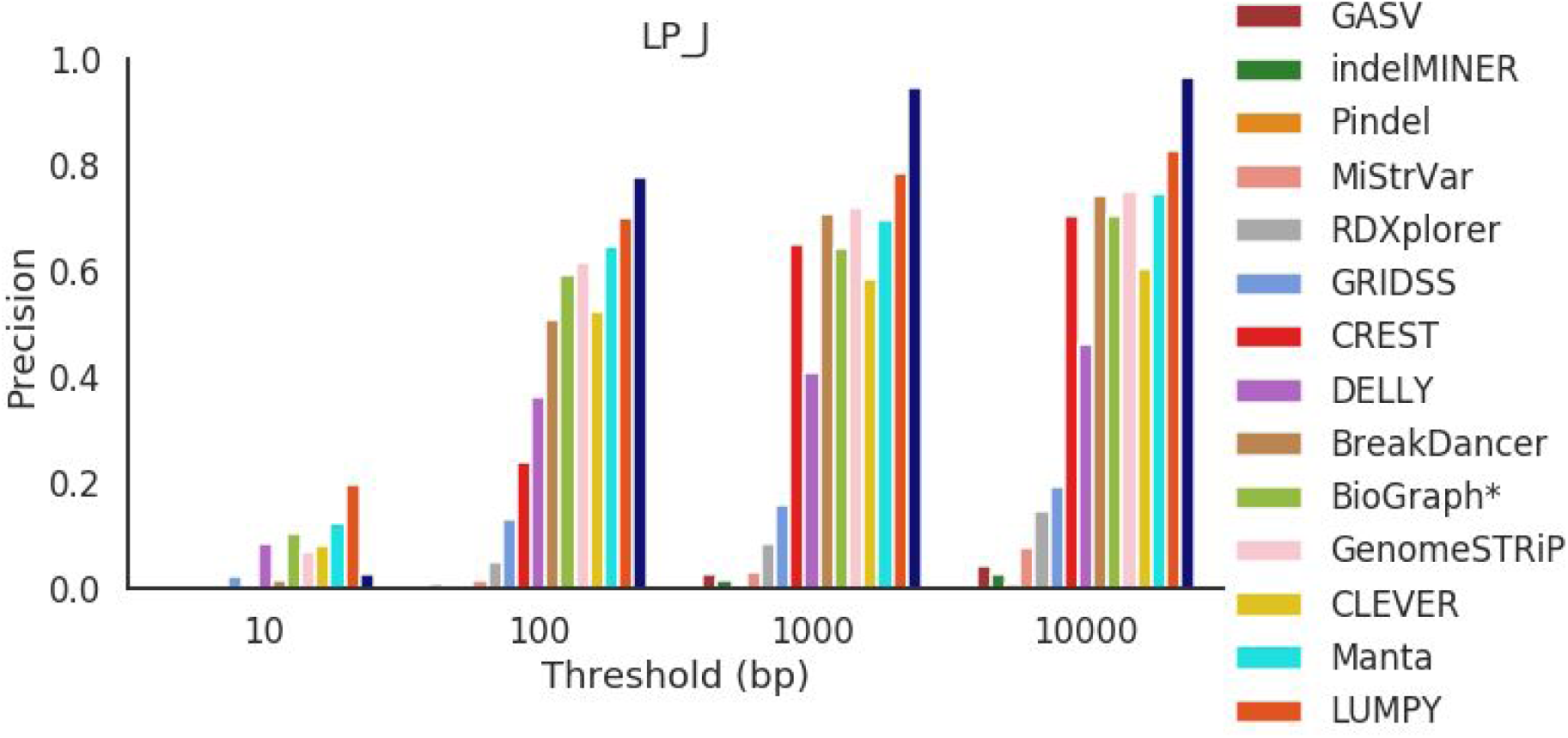
Precision presented for each mouse strain across all error thresholds.

**Figure S5.**
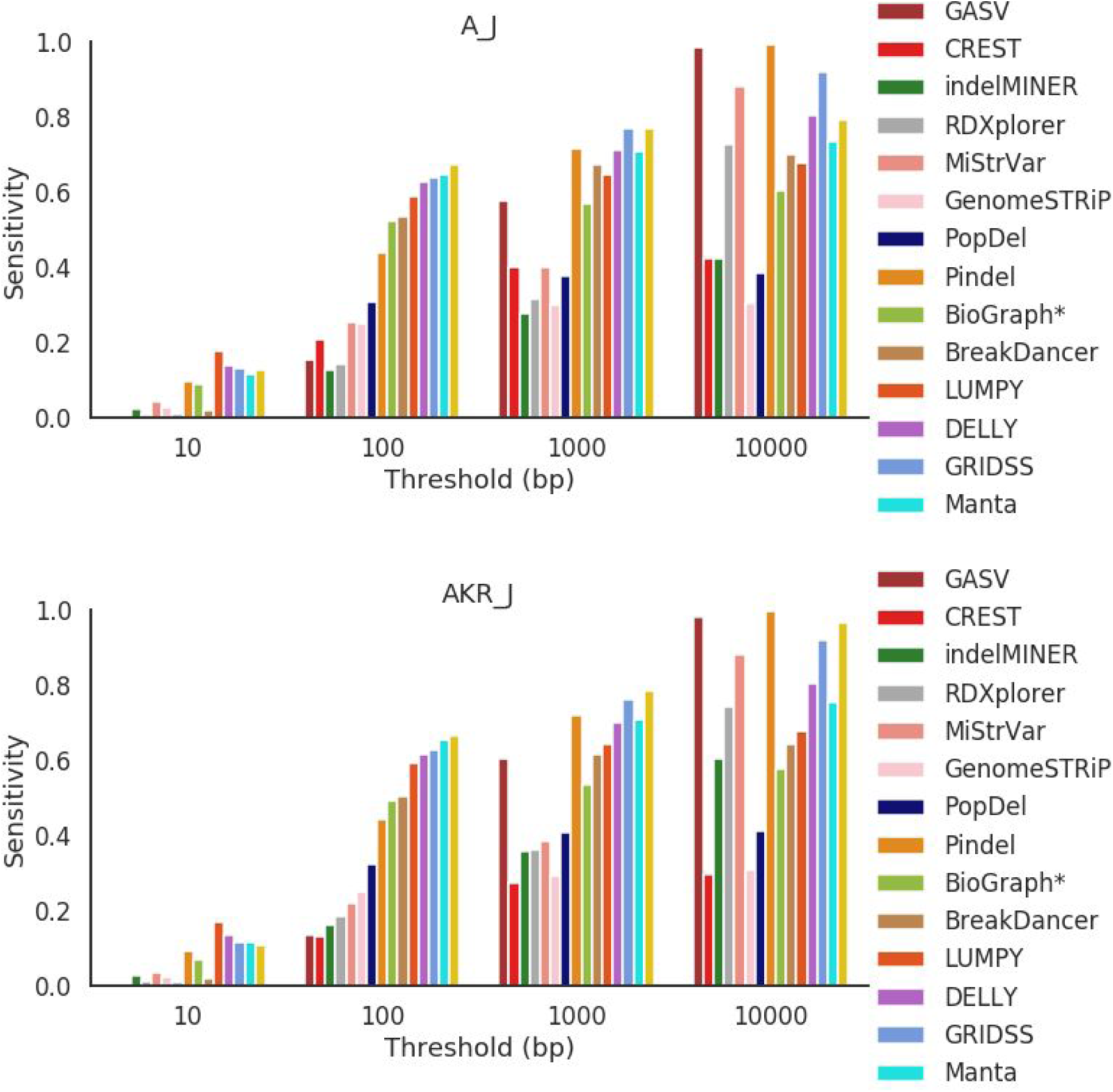
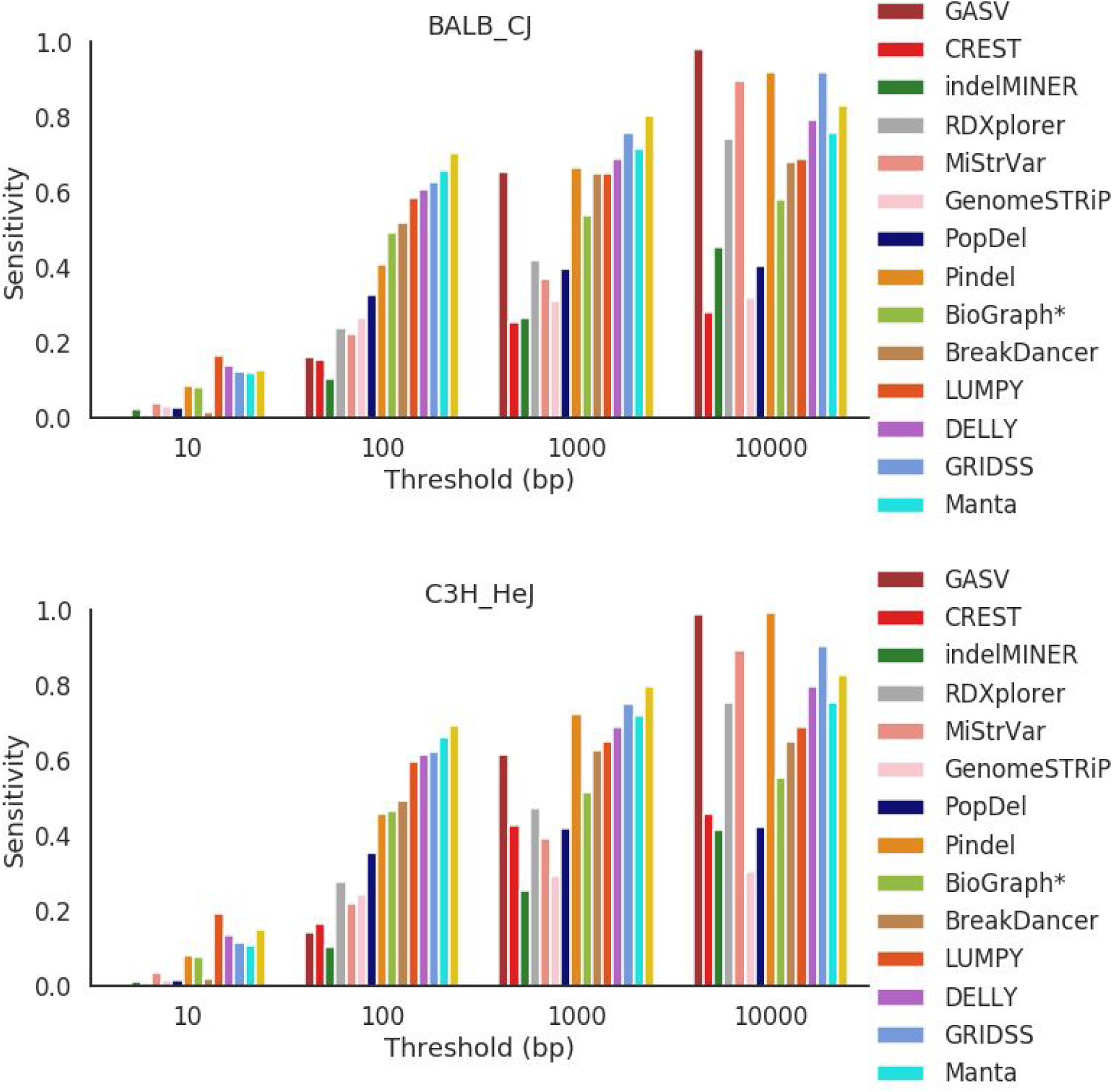
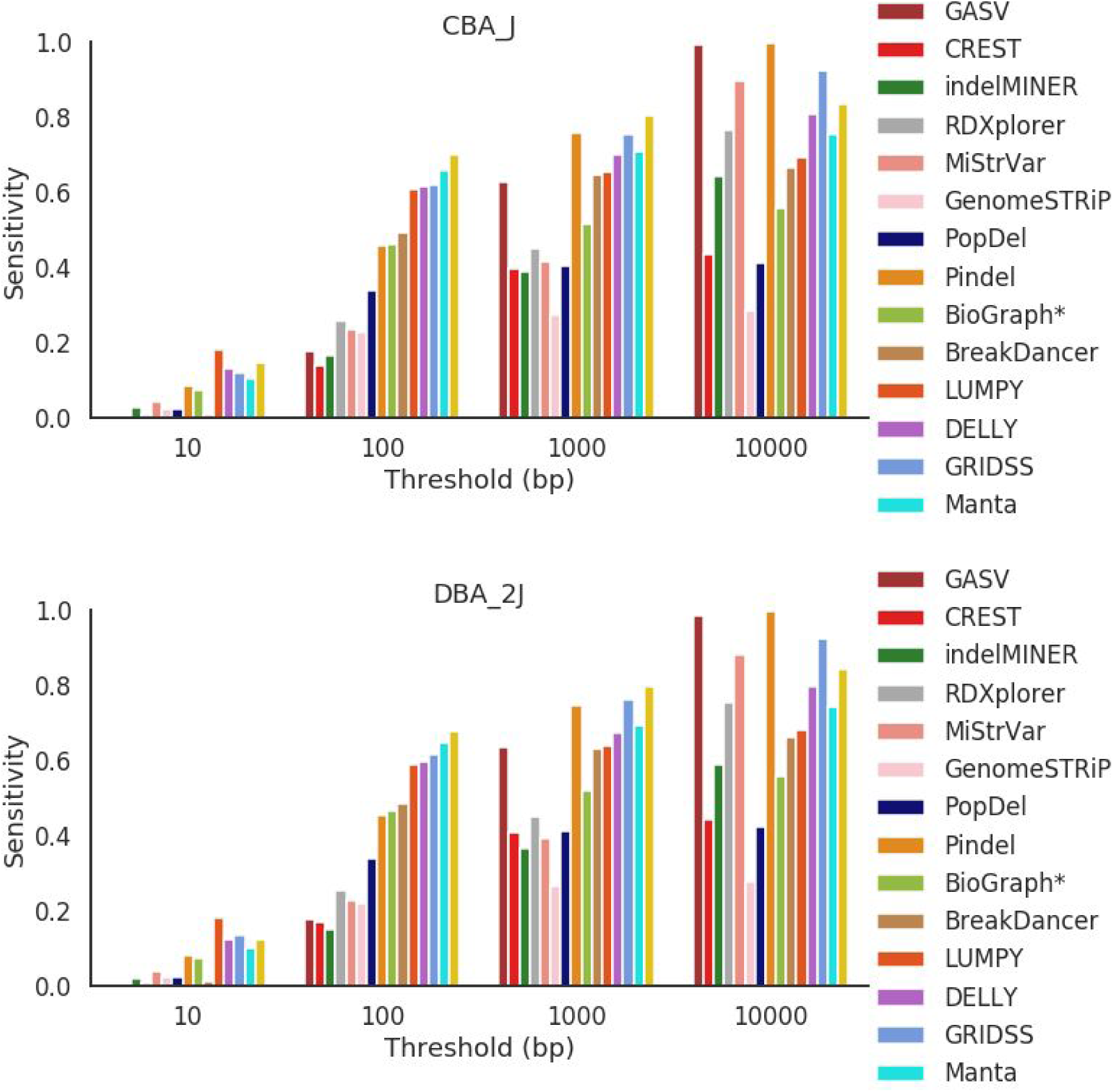
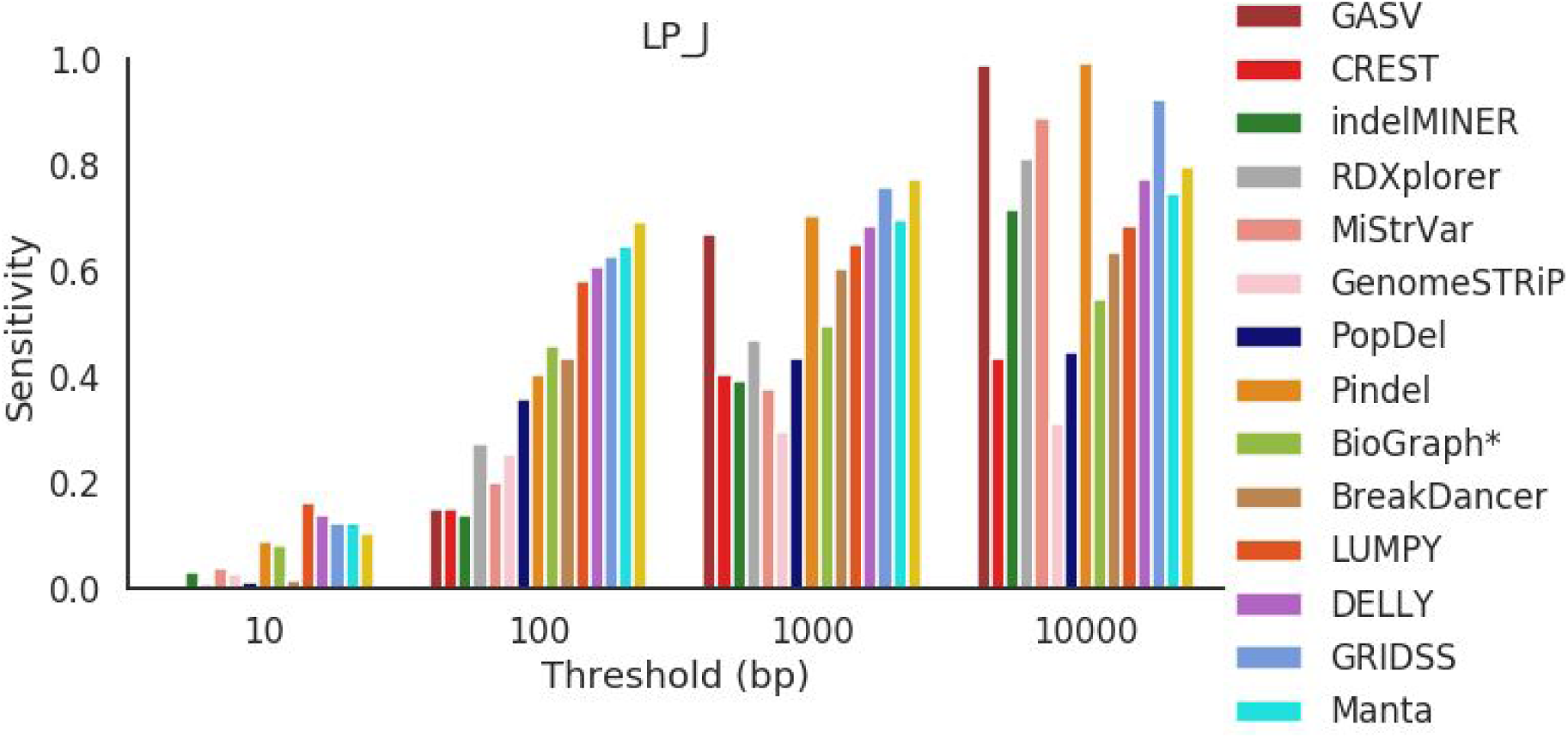
Sensitivity presented for each mouse strain across all error thresholds

**Figure S6.**
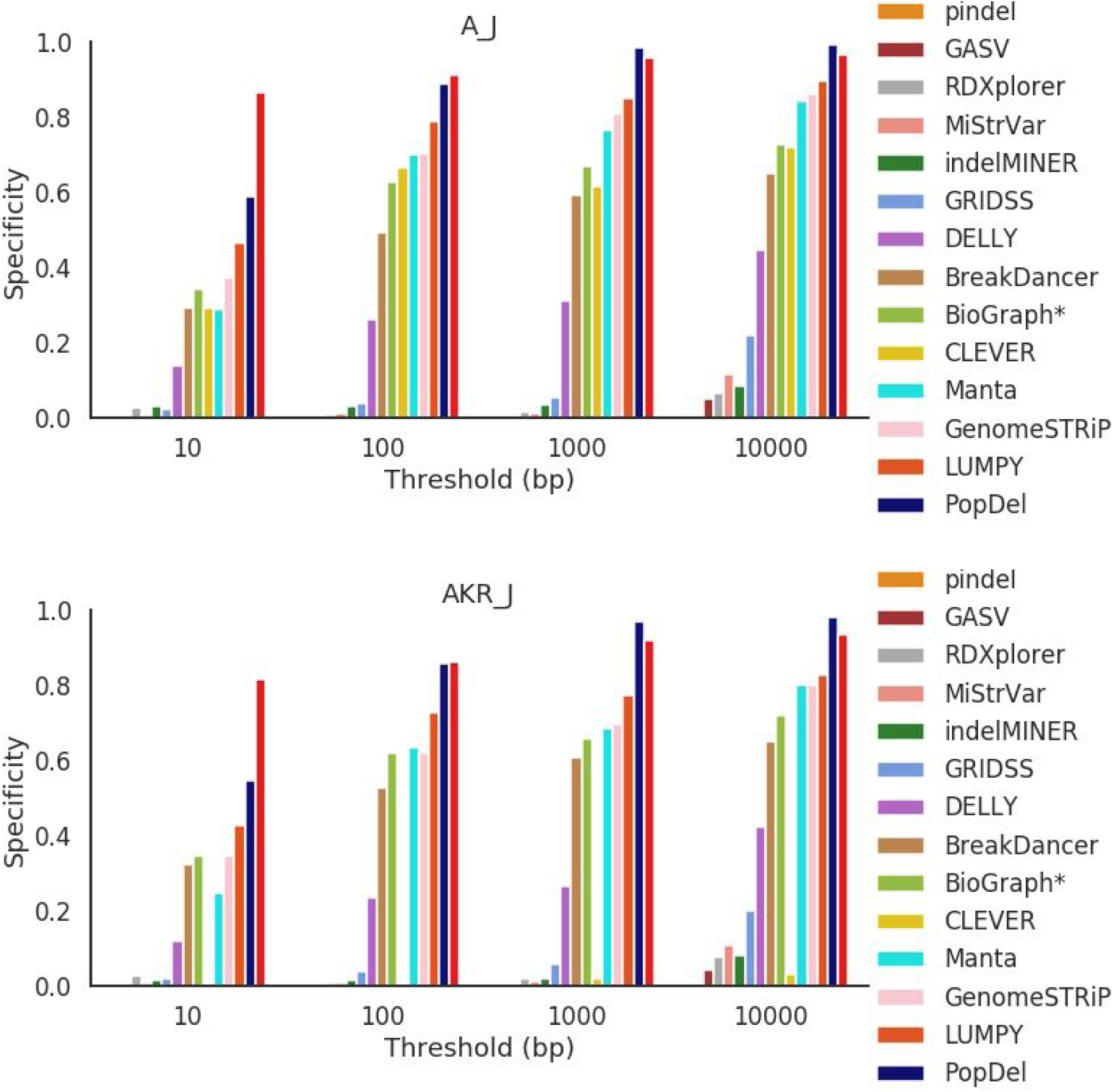
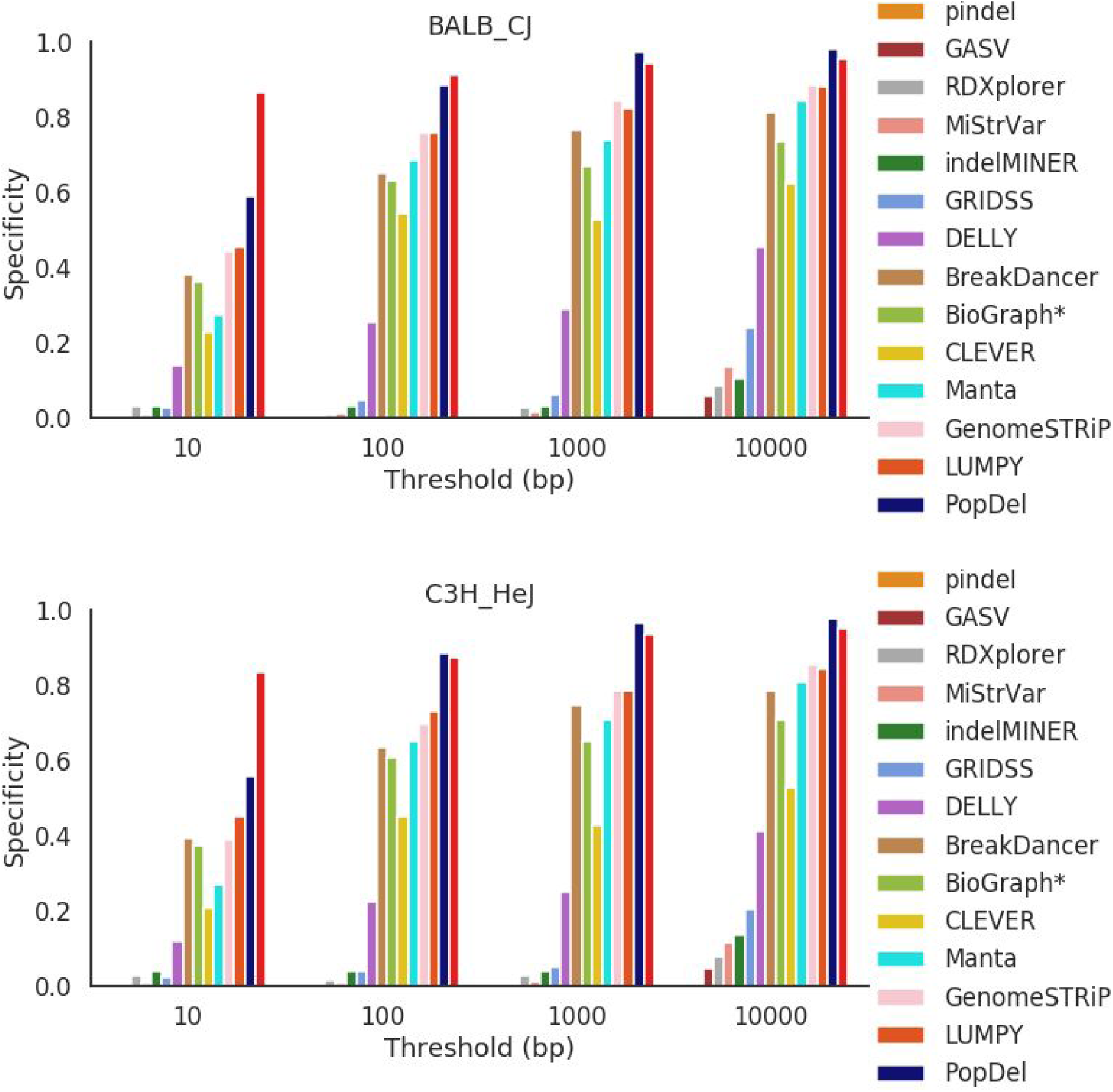
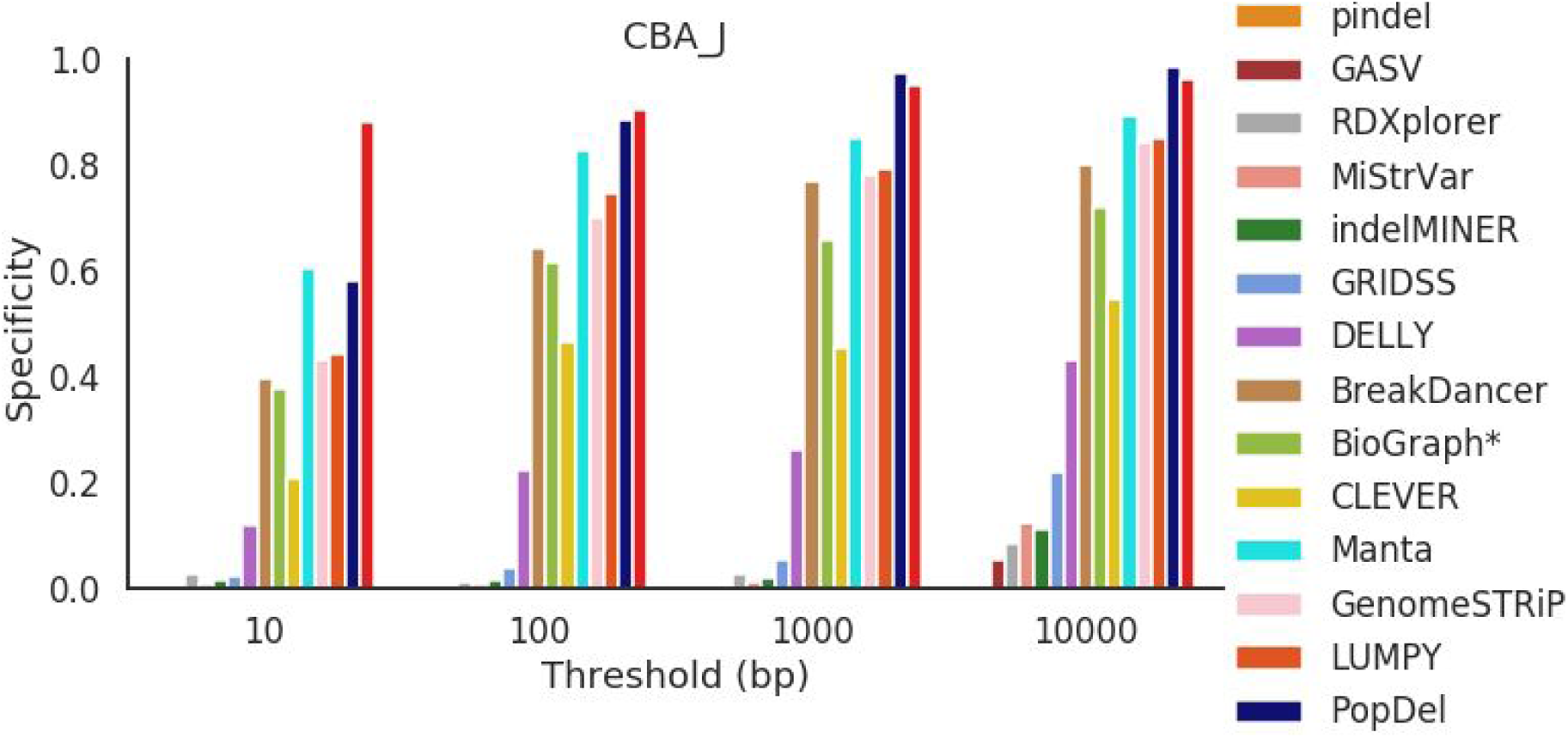
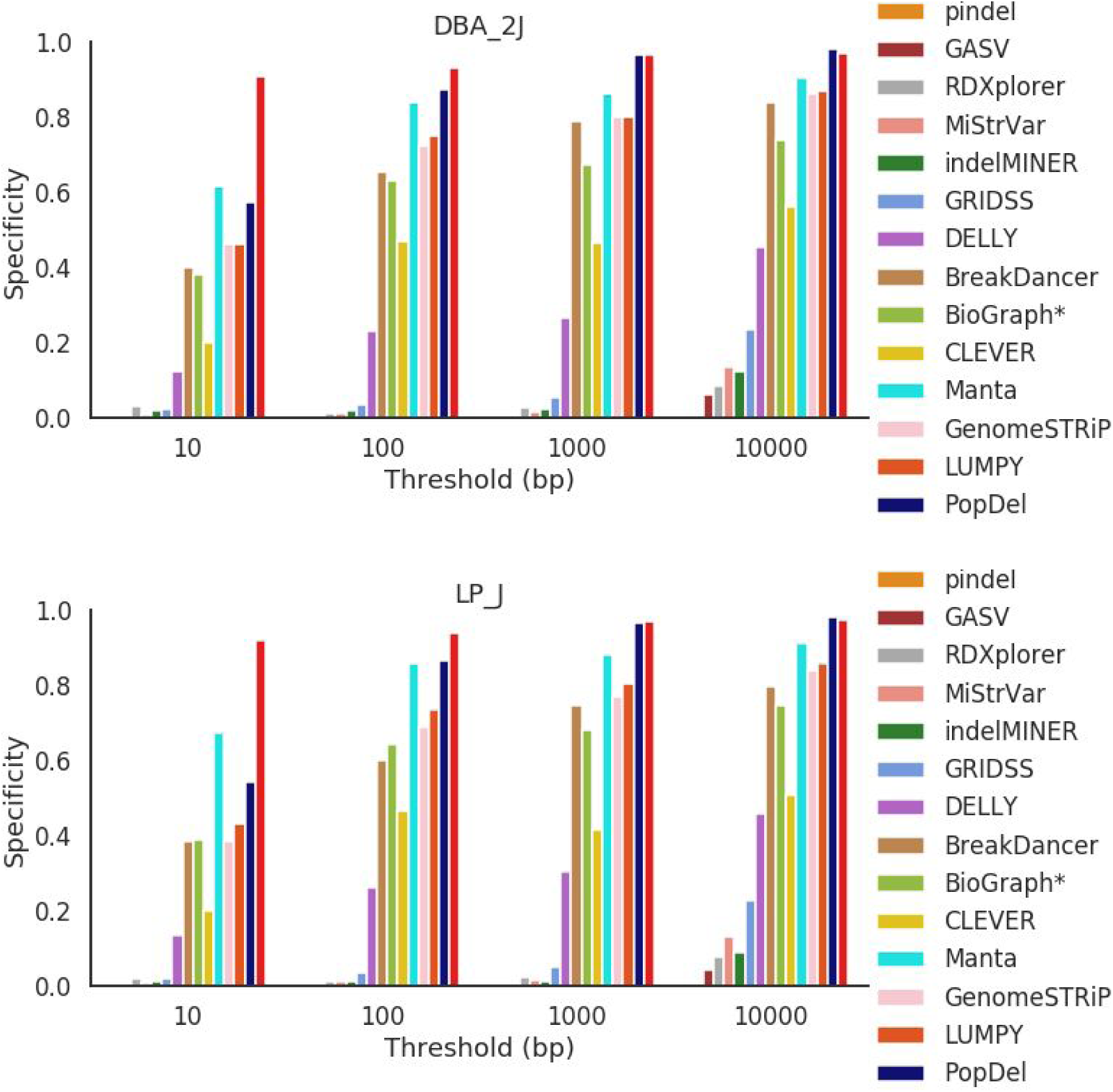
Specificity presented for each mouse strain across all error threshold

**Figure S7.**
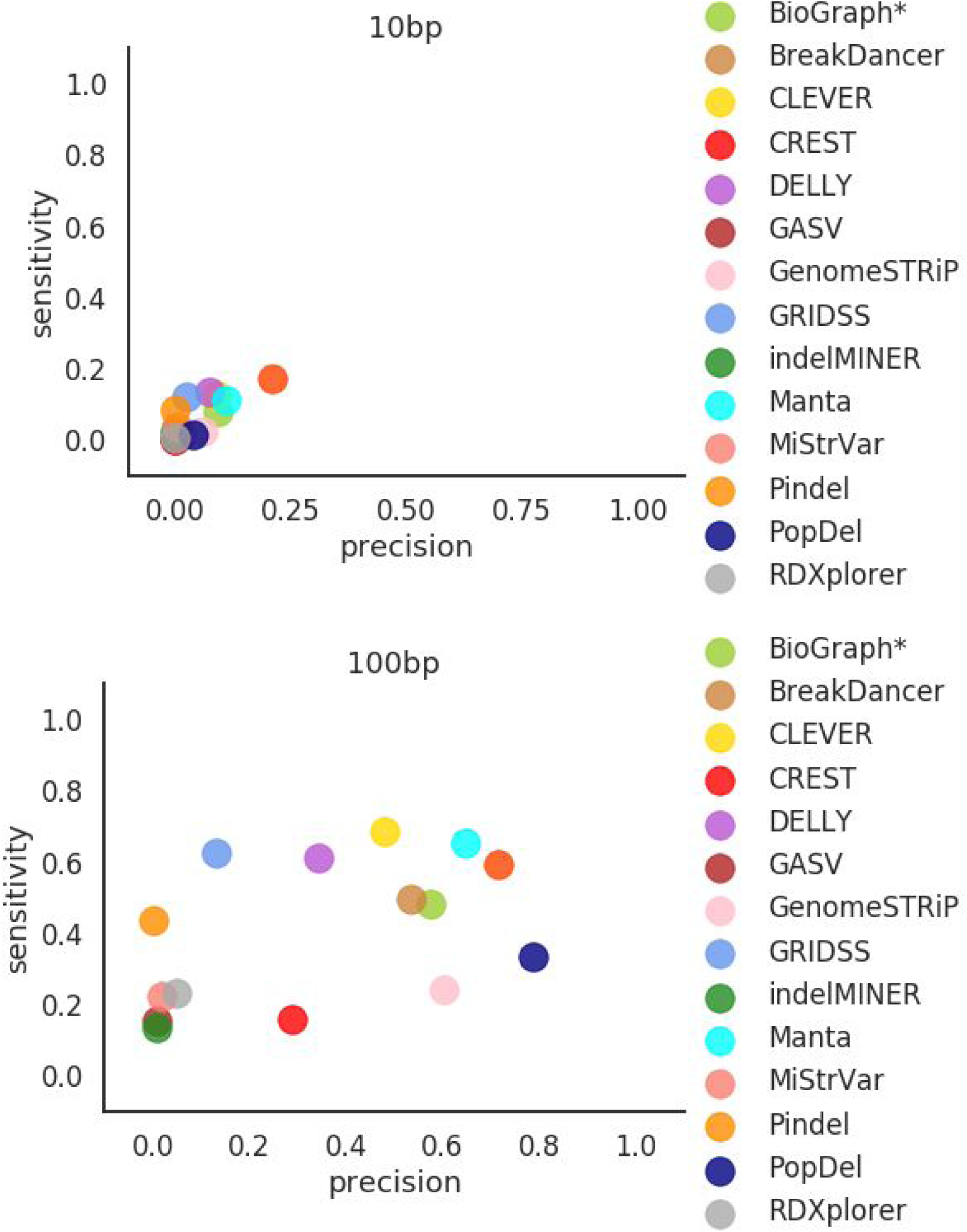
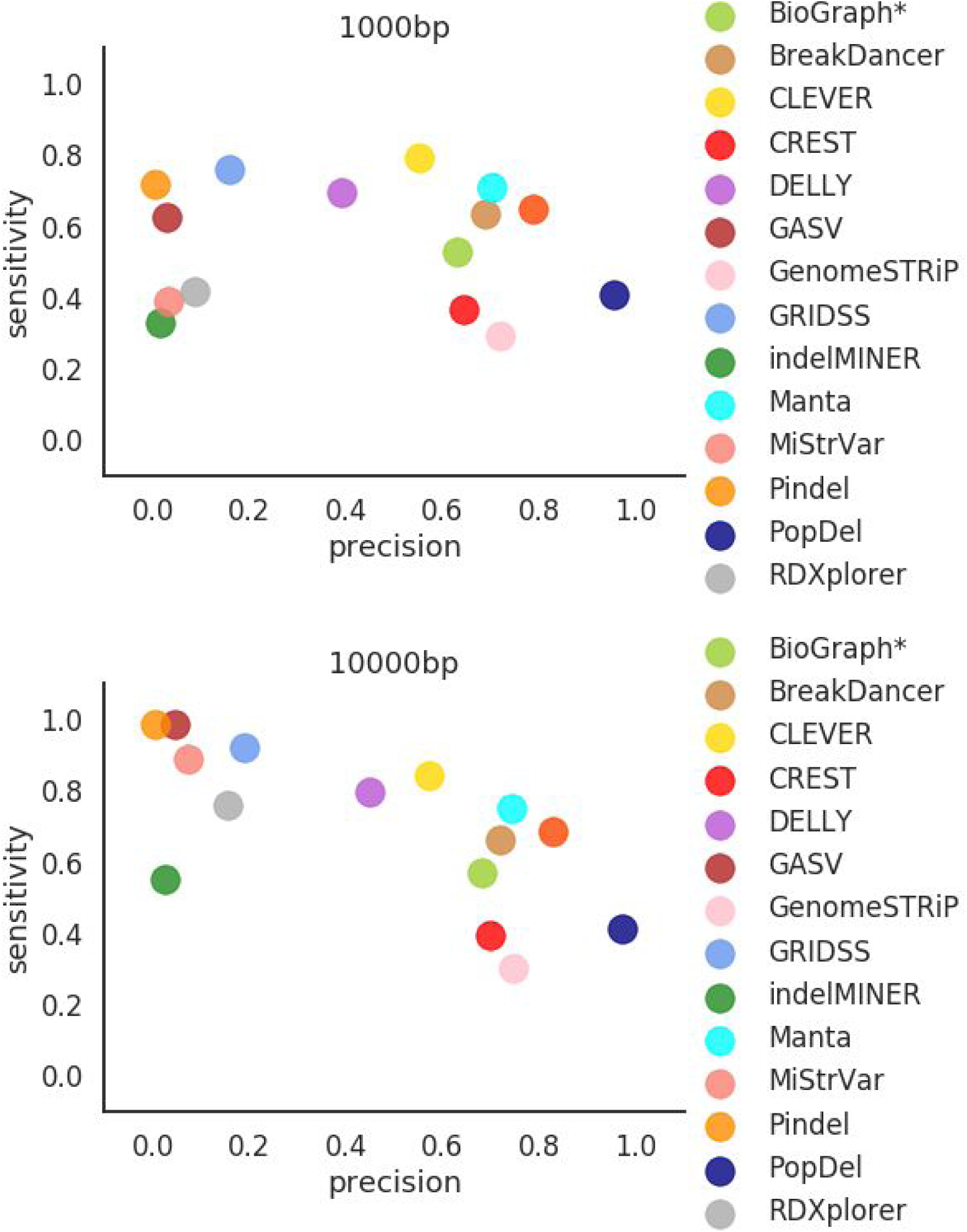
Scatter plot depicting the Precision (x-axis) and Sensitivity (y-axis) for 10 bp, 100 bp, 1000 bp, and 10000 bp thresholds.

**Figure S8.**
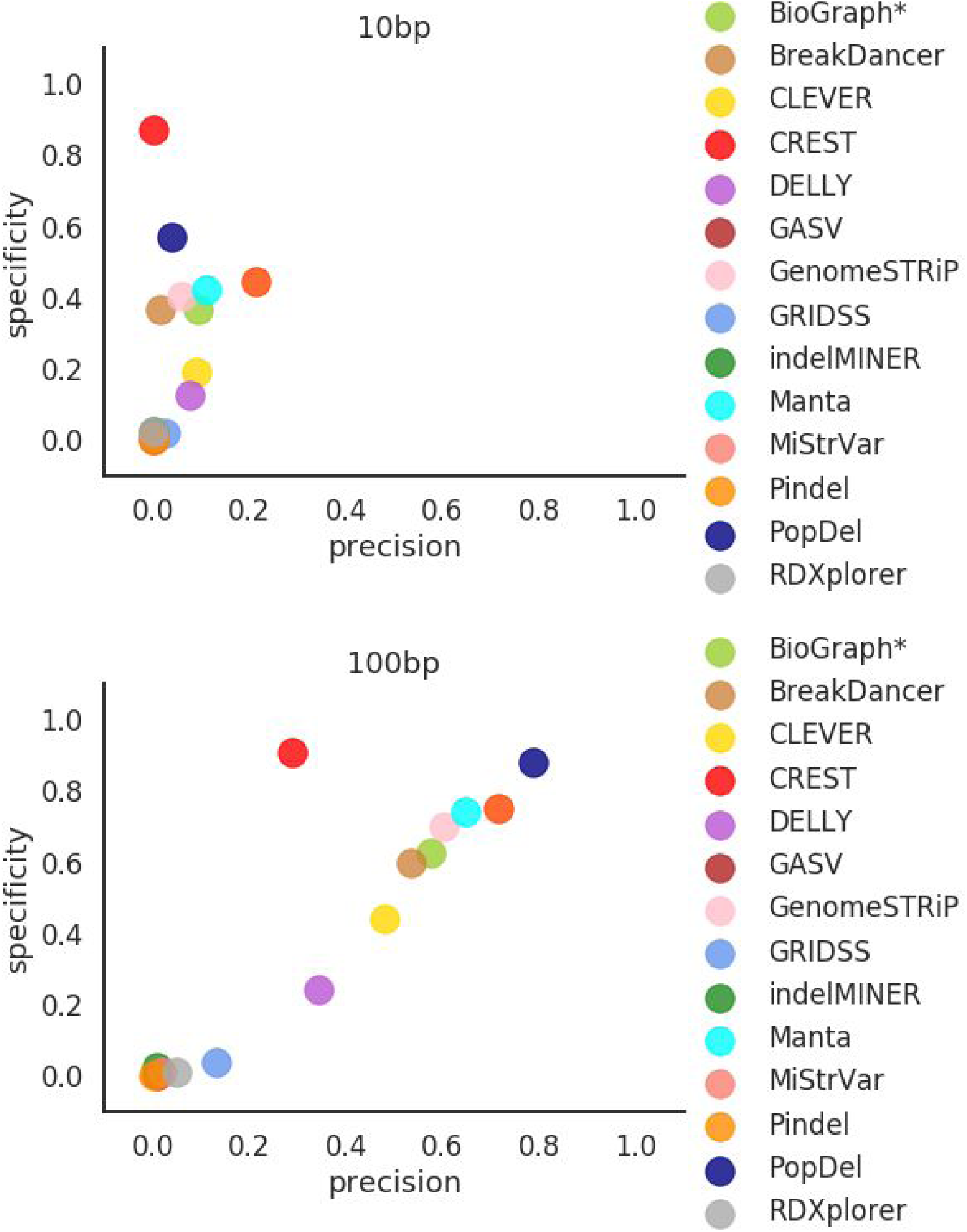
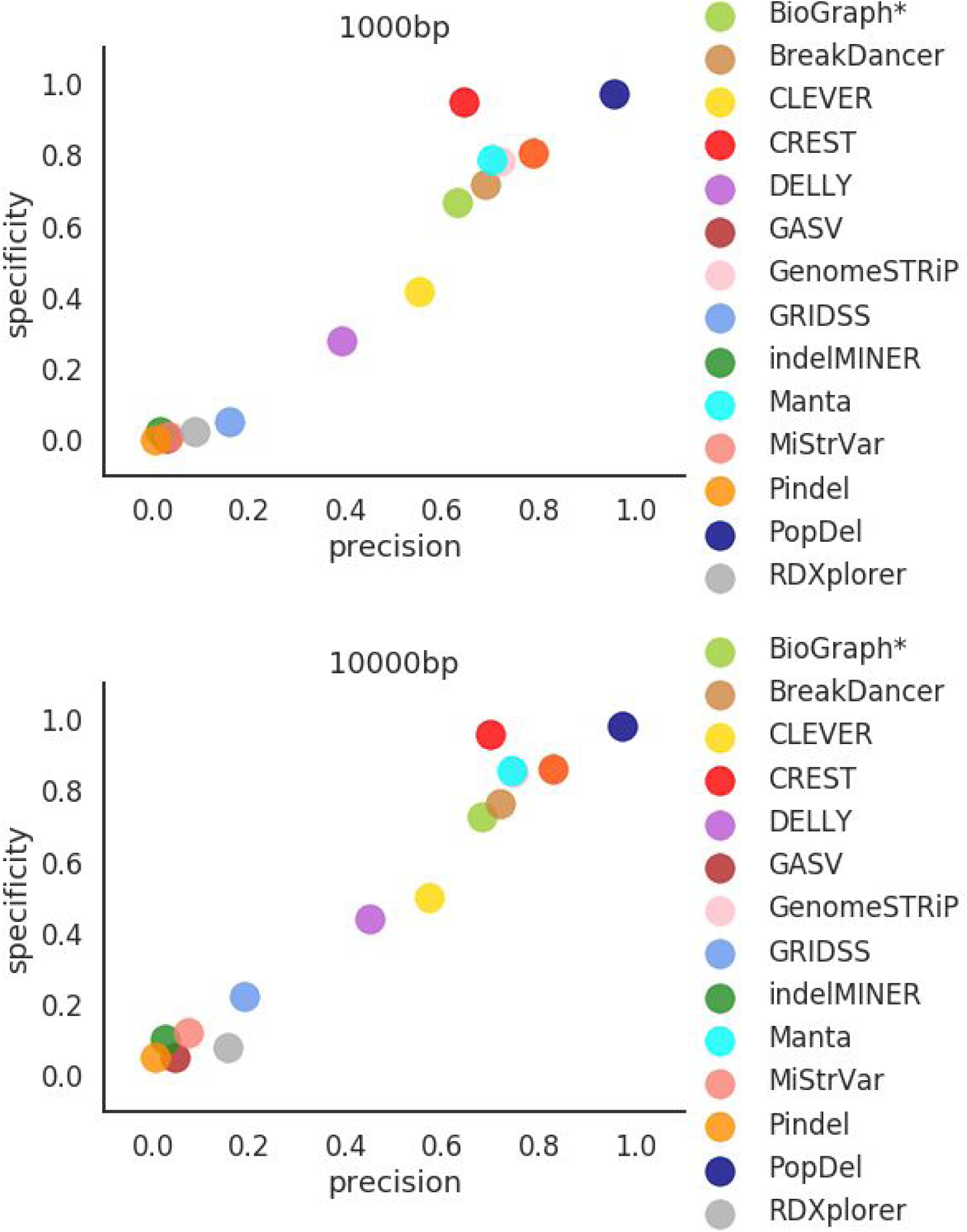
Scatter plot depicting the Precision (x-axis) vs Specificity (y-axis) for 10 bp, 100 bp, 1000 bp, and 10000 bp thresholds with Spearman’s correlation and p-values (0.53, 0.027; 0.84, 2.37e-05, 0.92,1.18e-07; 0.93,5.64e-08) respectively

**Figure S9.**
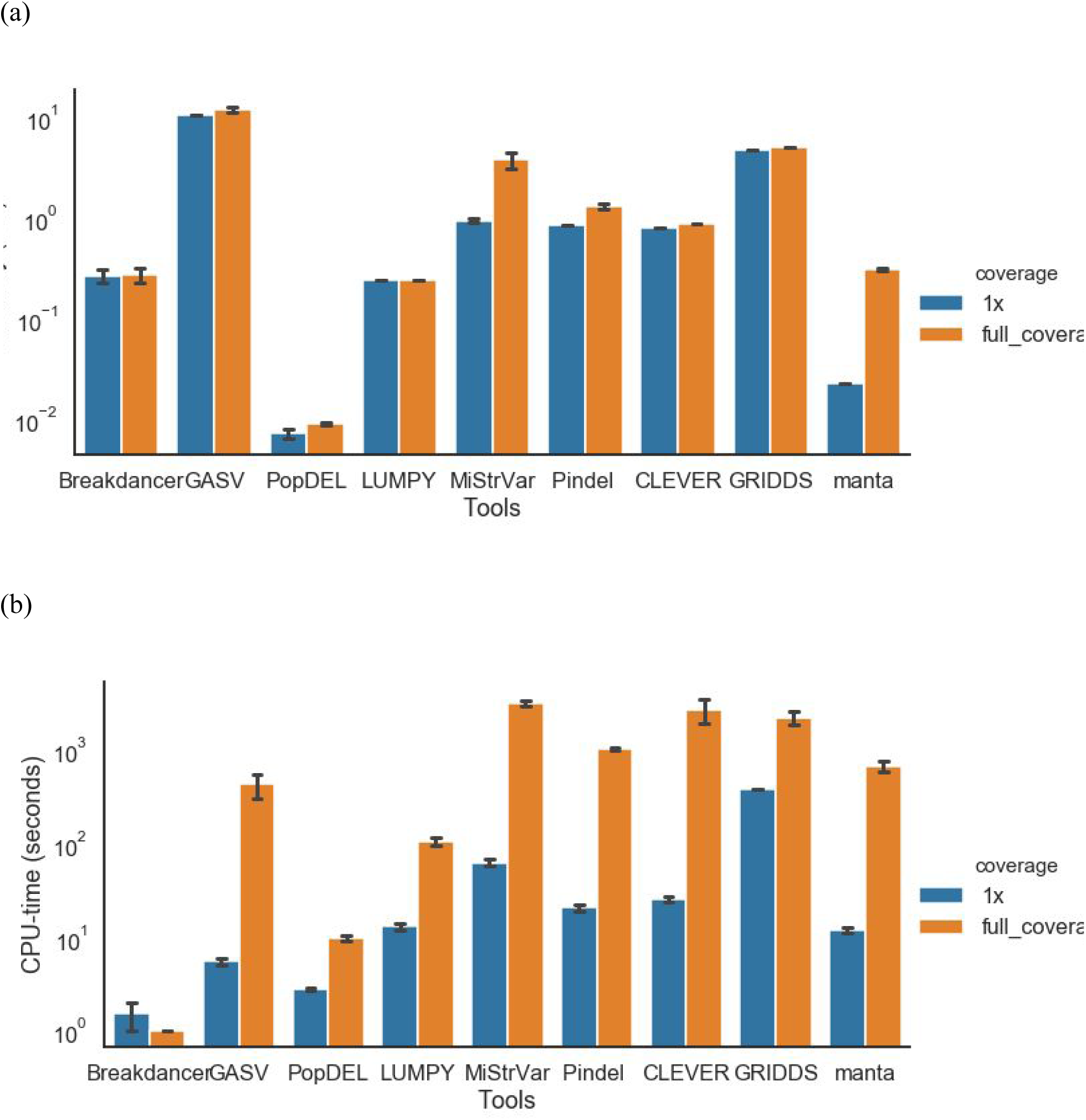
Comparison of computational performance of SV callers. (a) A bar-plot depicting the RAM usage across all of the tools in Table 1. (b) A bar-plot depicting the CPU-time across all of the tools in Table 1.

**Figure S10.**
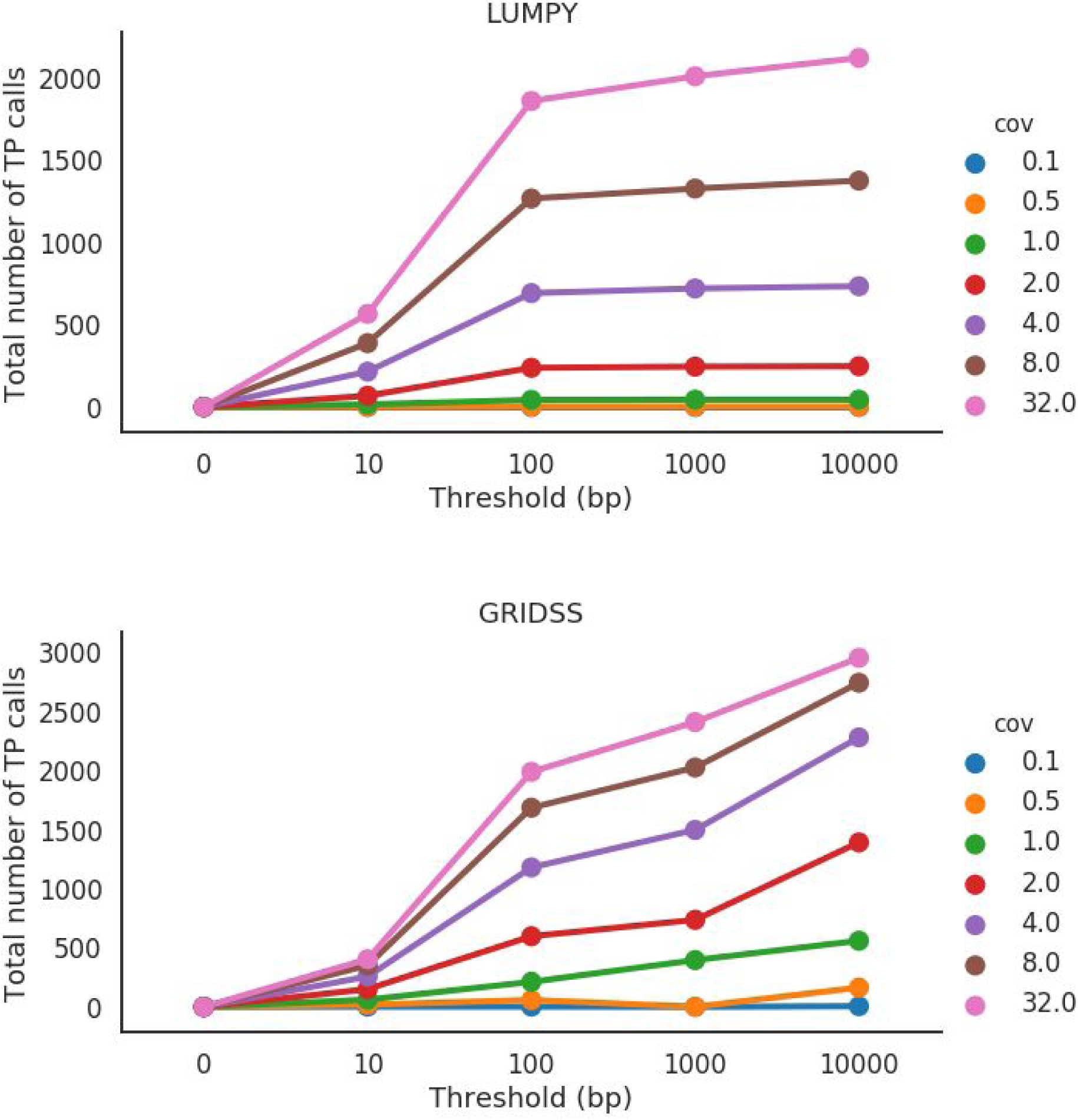
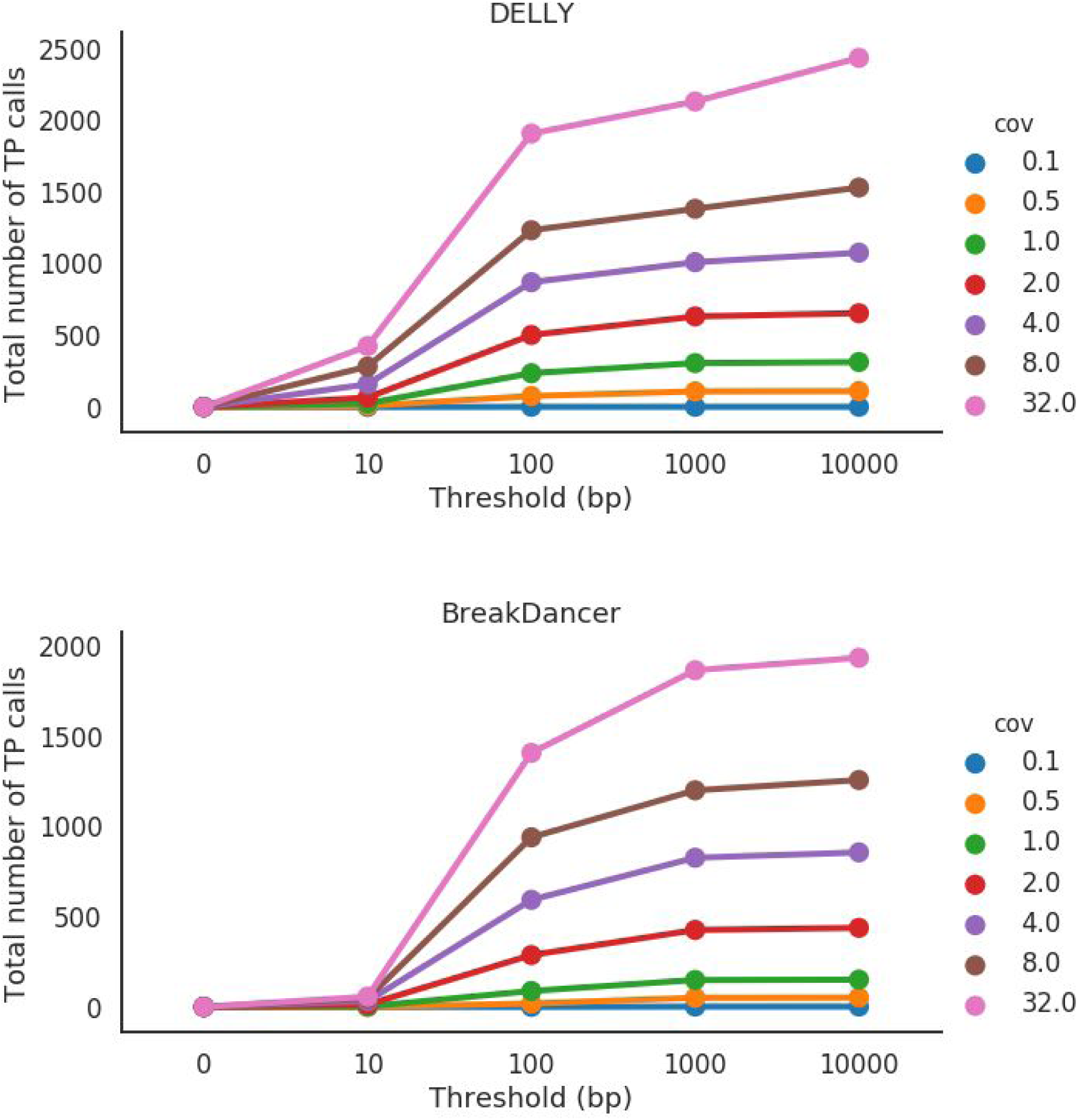
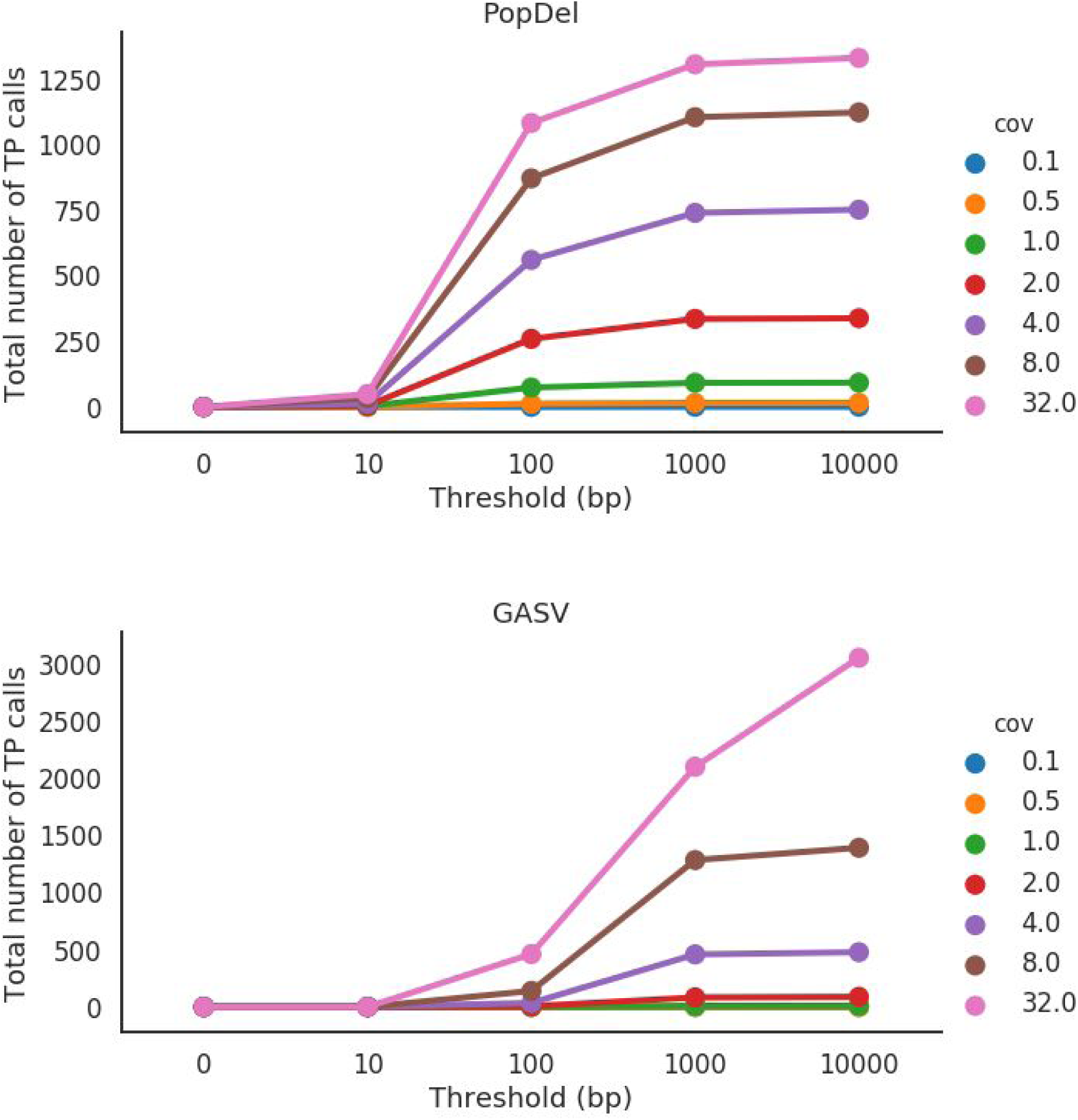
Number of correctly detected deletions (true positives - ‘TP’) by SV callers across various thresholds and genome coverages. Deletion is considered to be correctly predicted if the distance of right and left coordinates are within the given threshold from the coordinates of true deletion.

**Figure S11.**
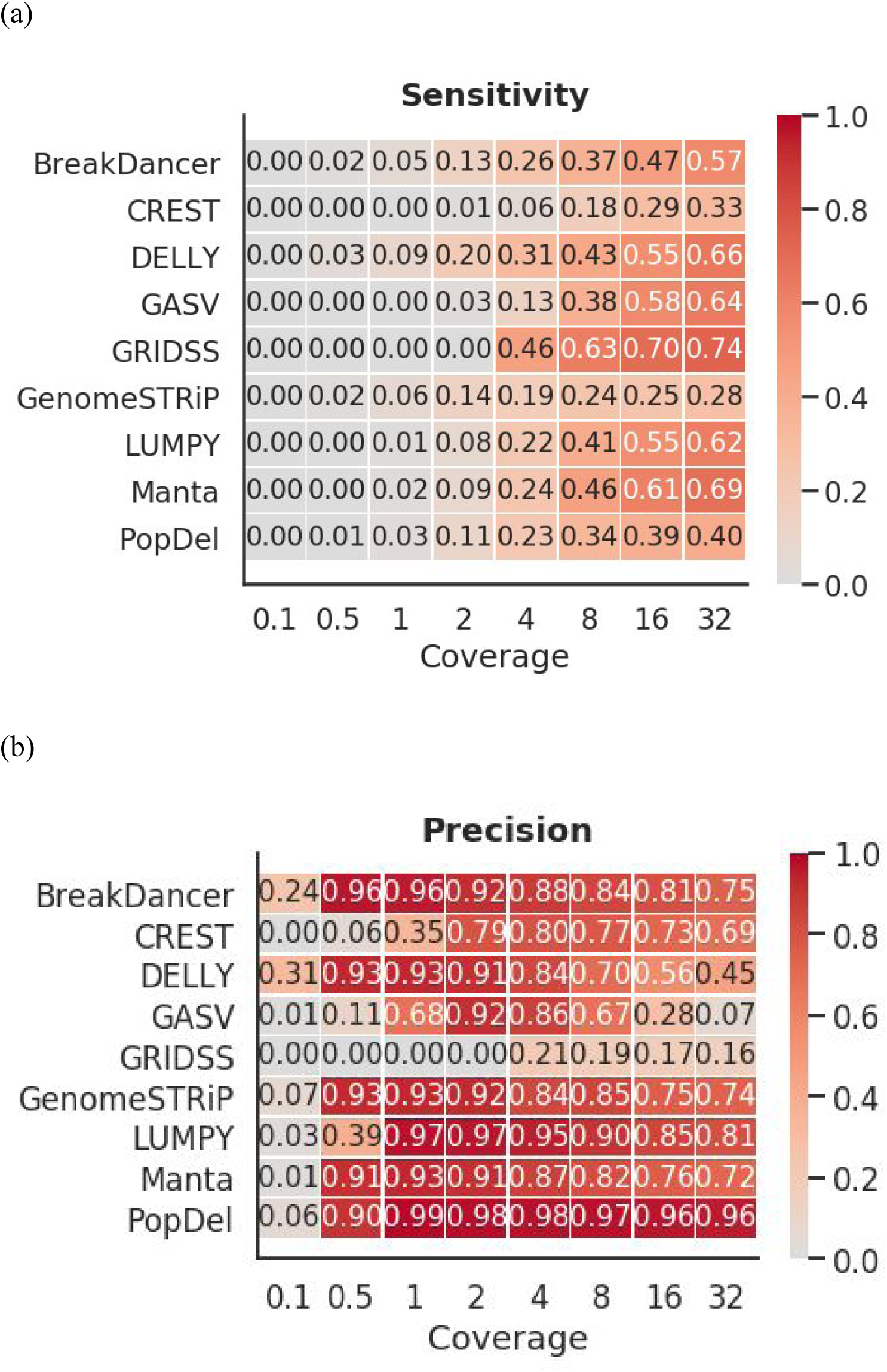
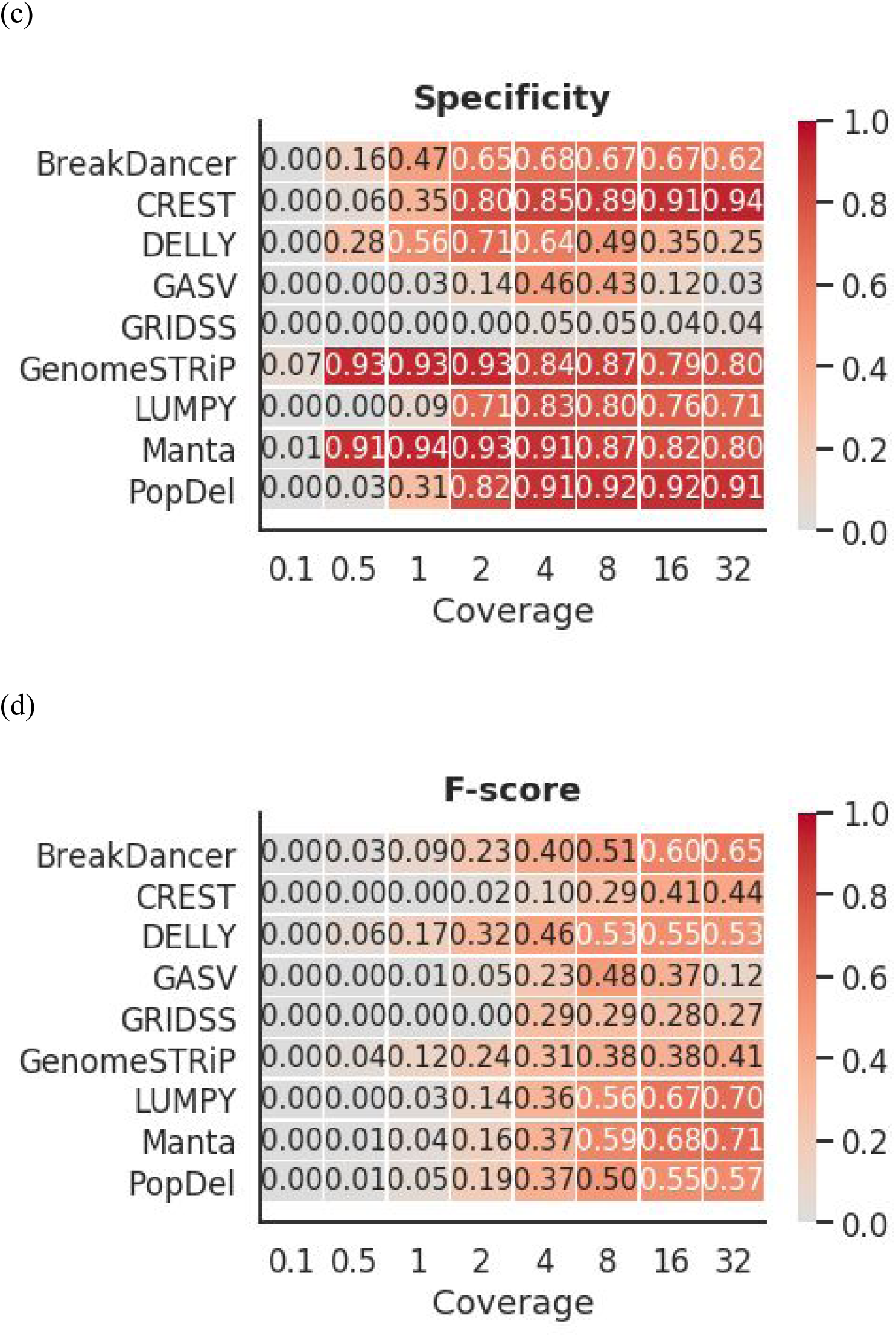
(a) Heatmap depicting the sensitivity based on 1000 bp threshold across various levels of coverage. (b) Heatmap depicting the precision based on 1000 bp threshold across various levels of coverage. (c) Heatmap depicting the specificity based on 1000 bp threshold across various levels of coverage. (d) Heatmap depicting the F-score based on 1000 bp threshold across various levels of coverage.

**Figure S12.**
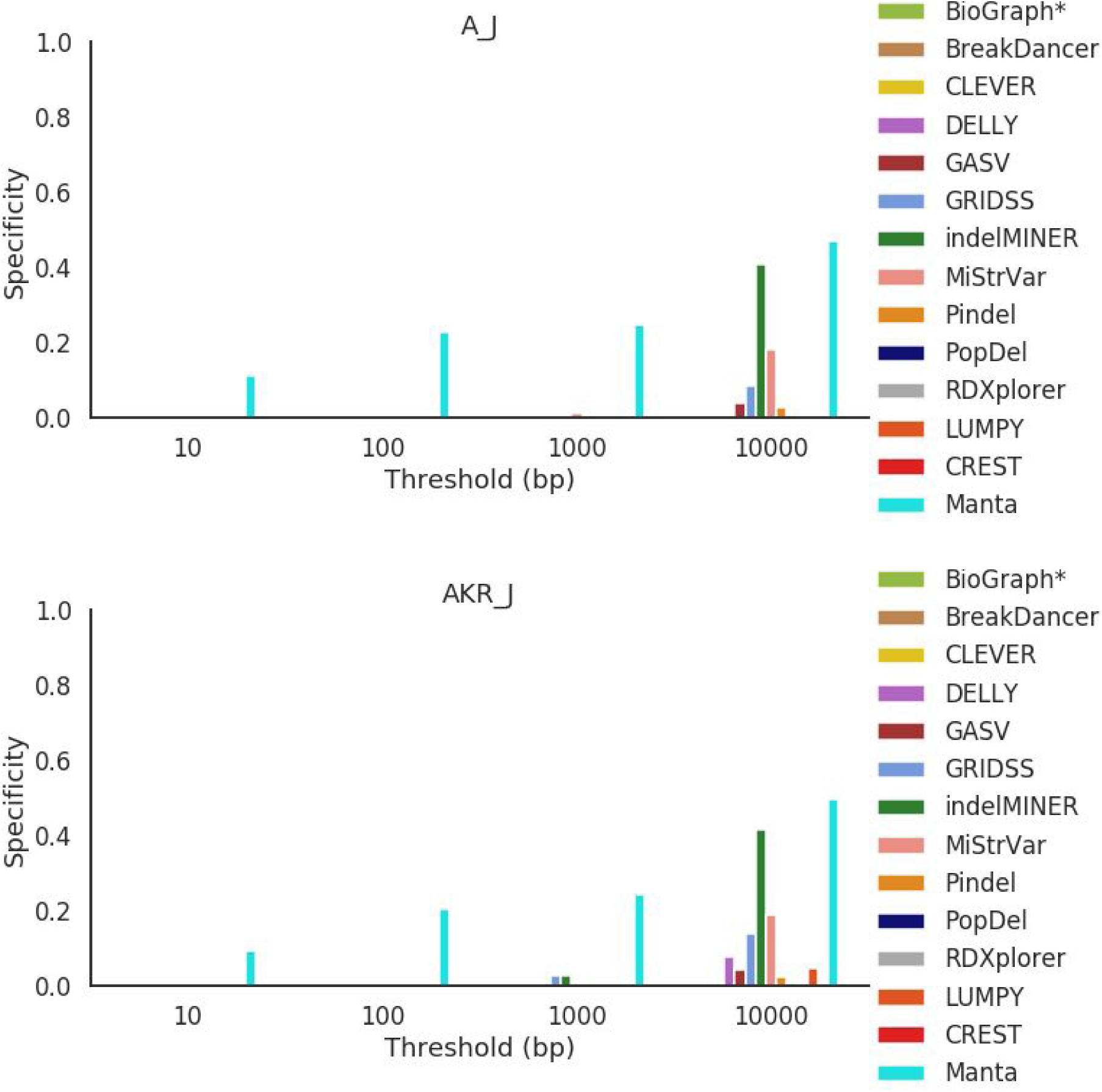
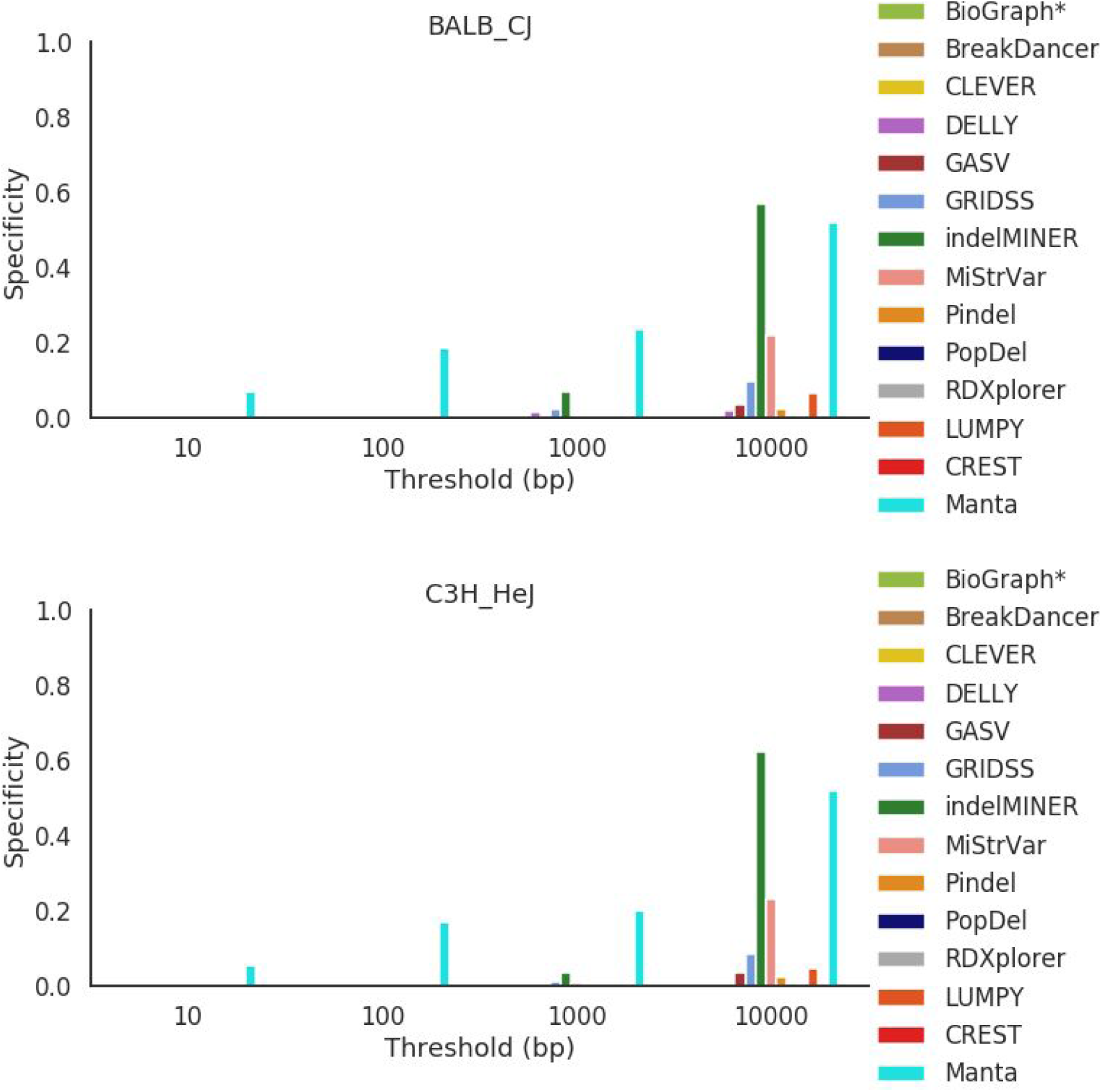
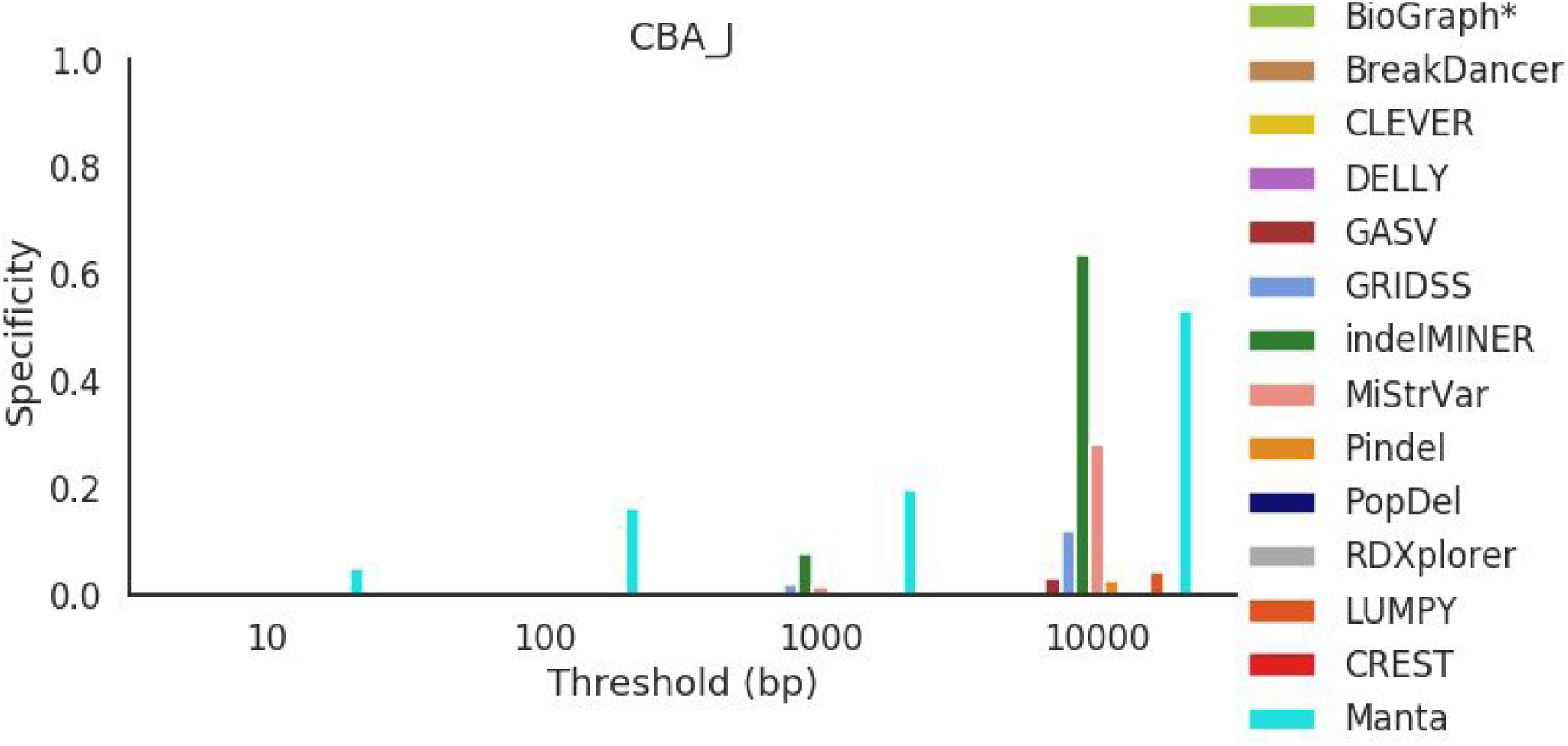
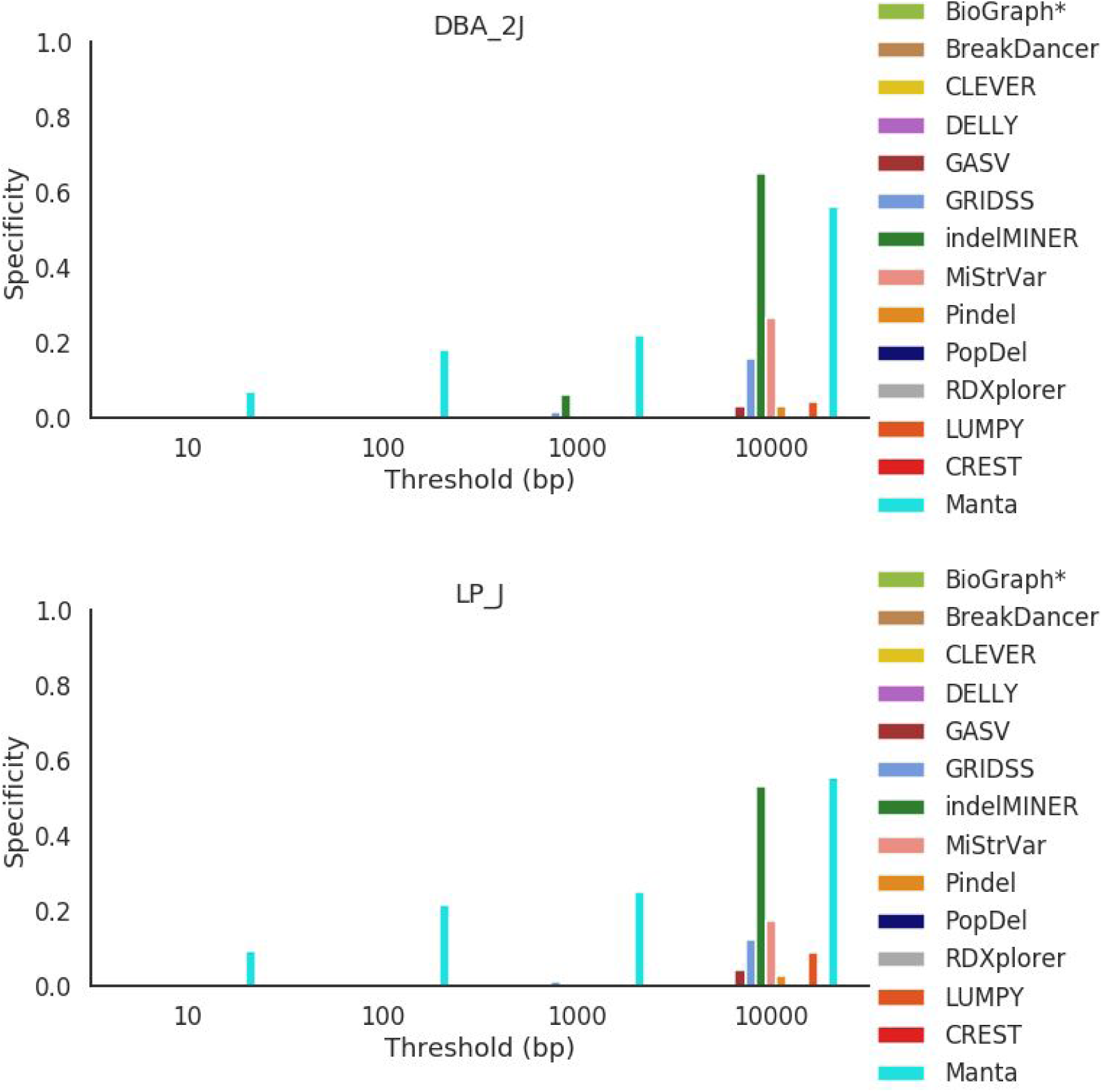
Specificity across all mouse strains for deletions between 50 bp and 100 bp in length.

**Figure S13.**
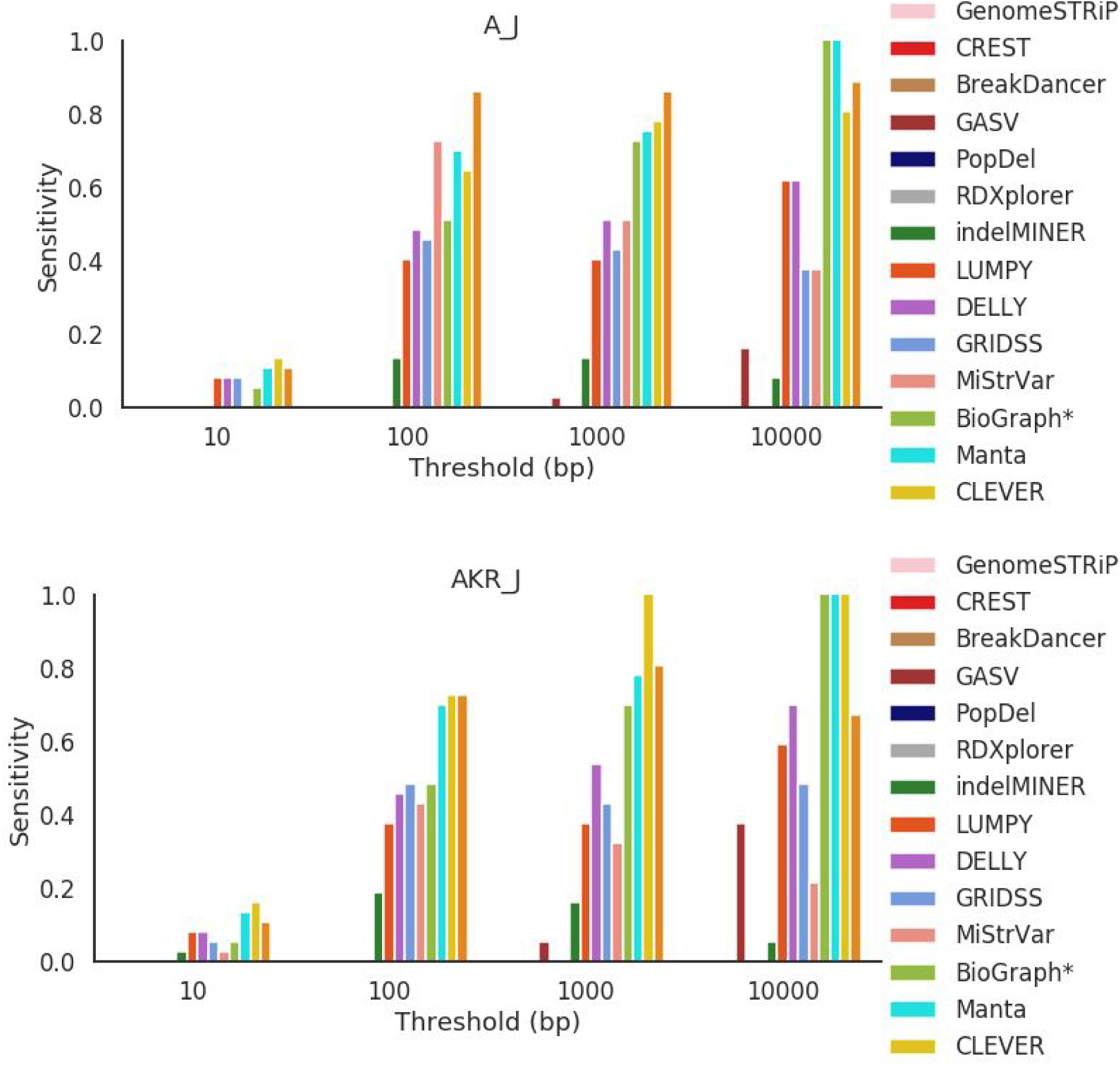
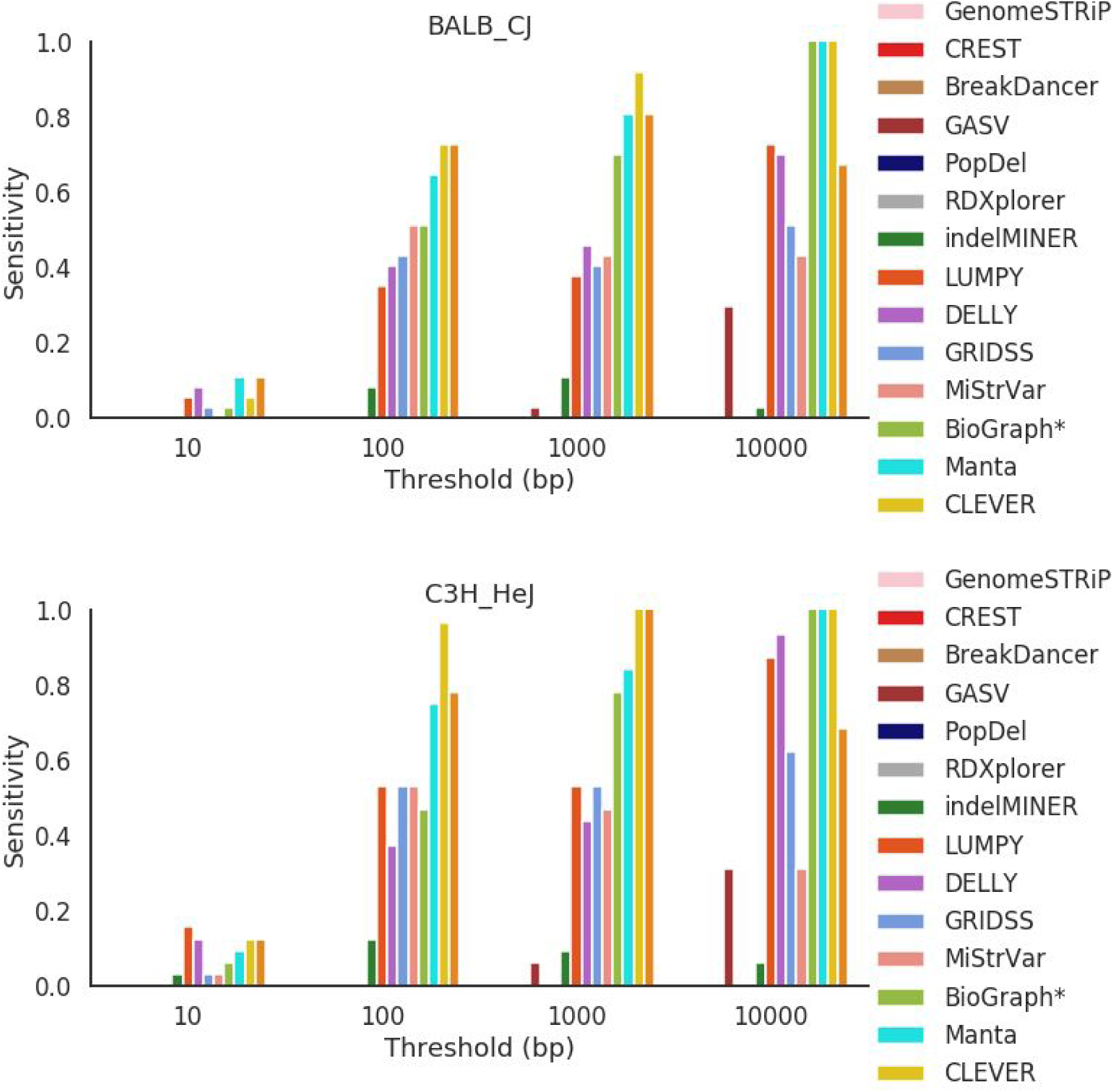
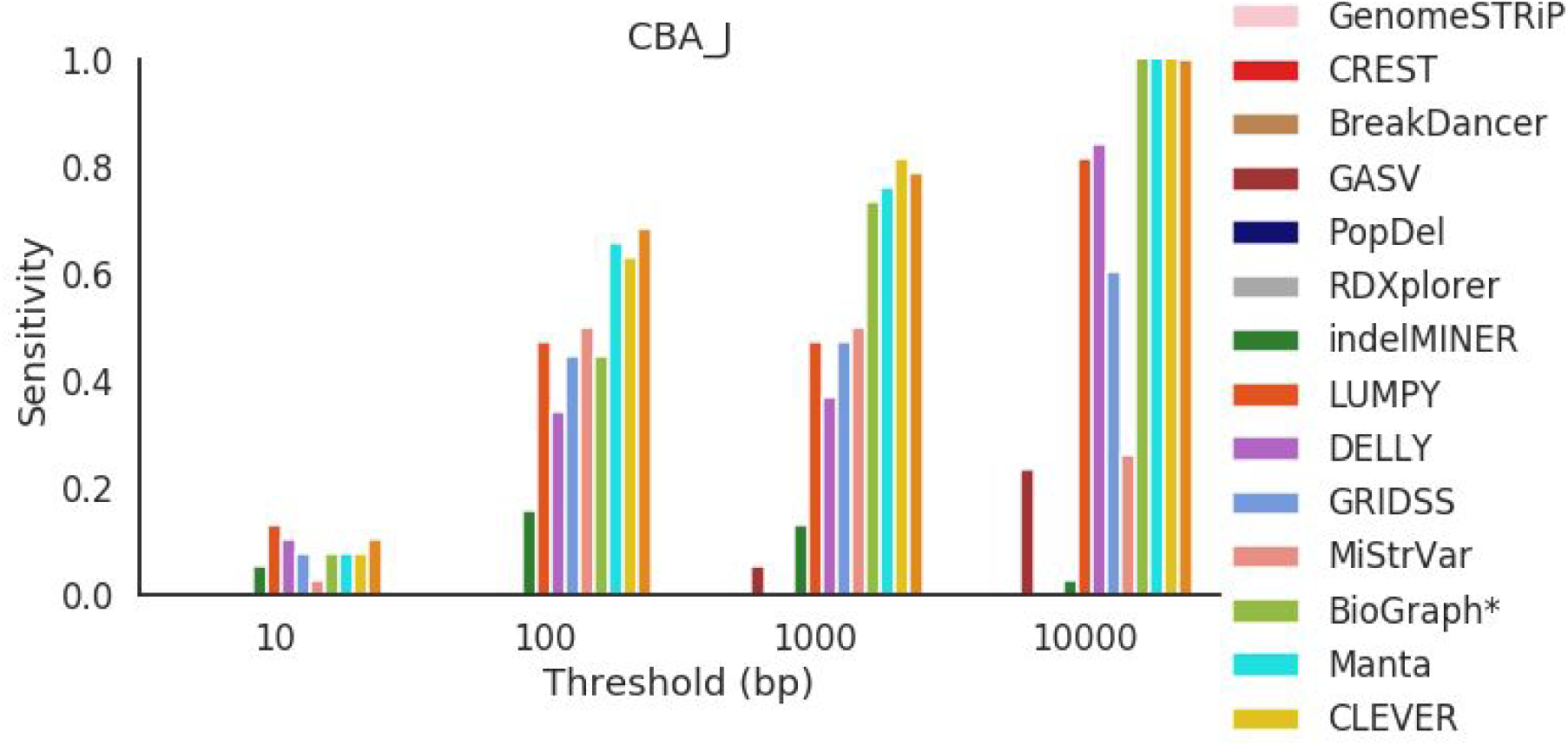
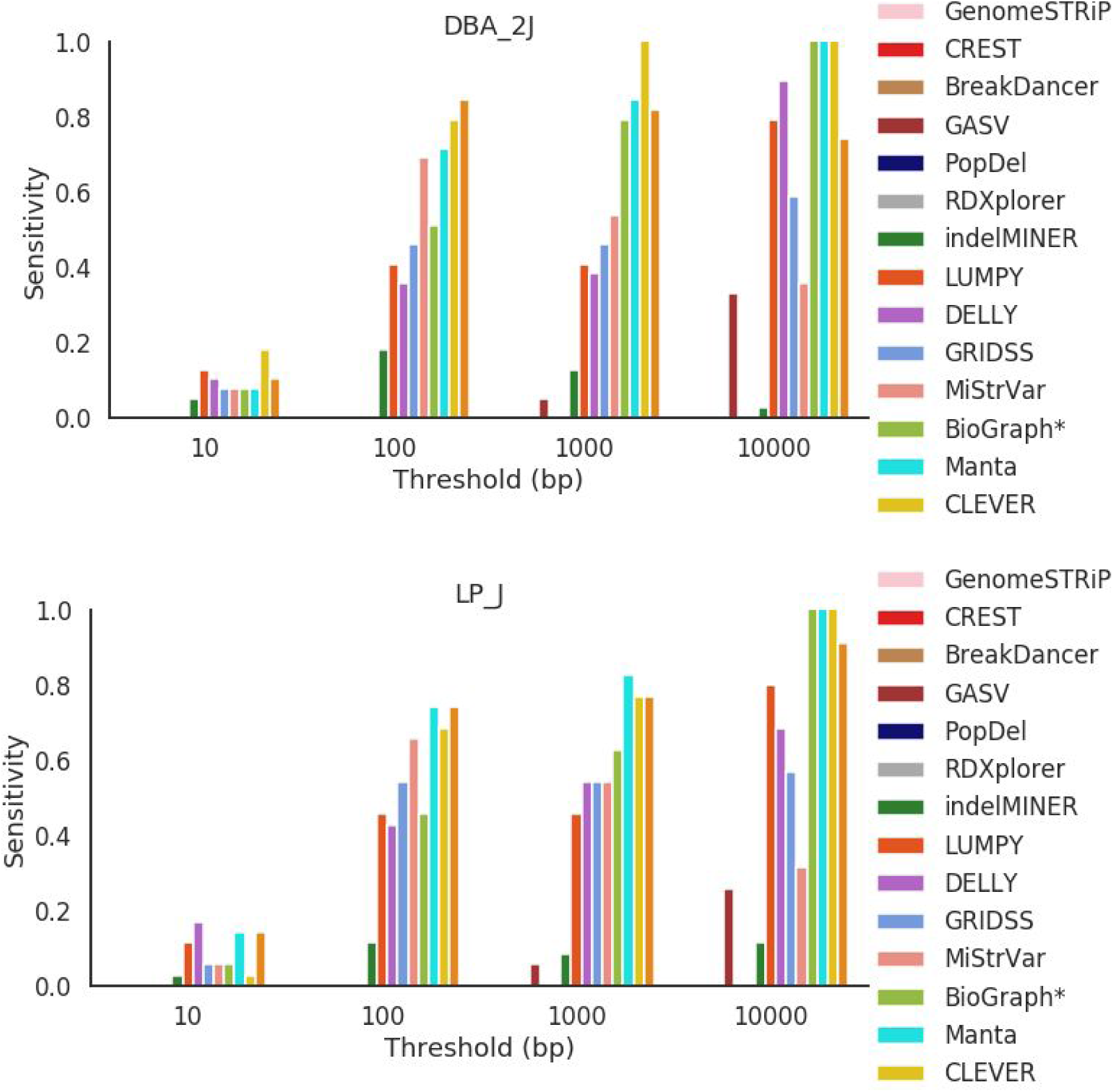
Sensitivity across all mouse strains for deletions between 50 bp and 100 bp in length.

**Figure S14.**
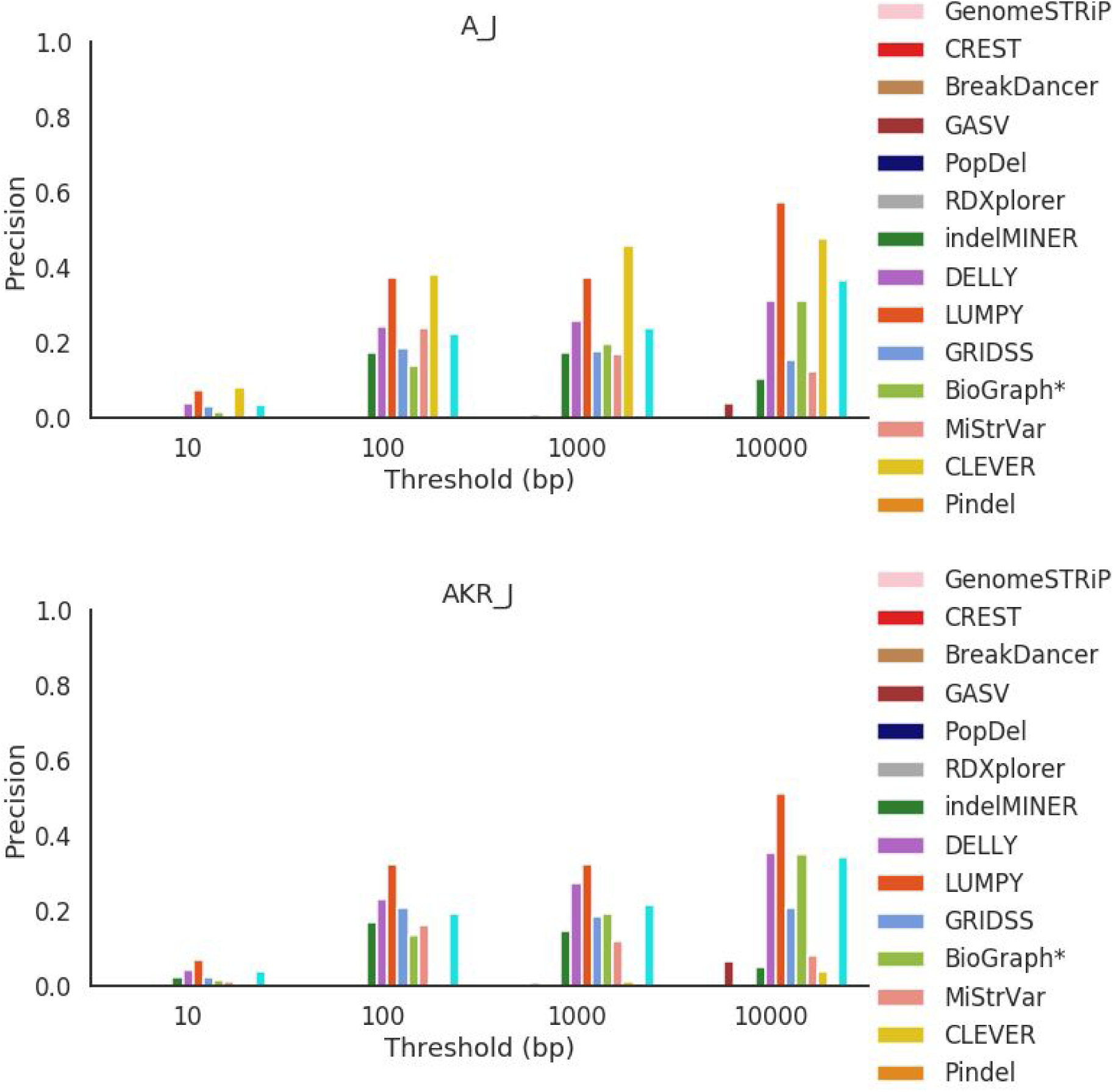
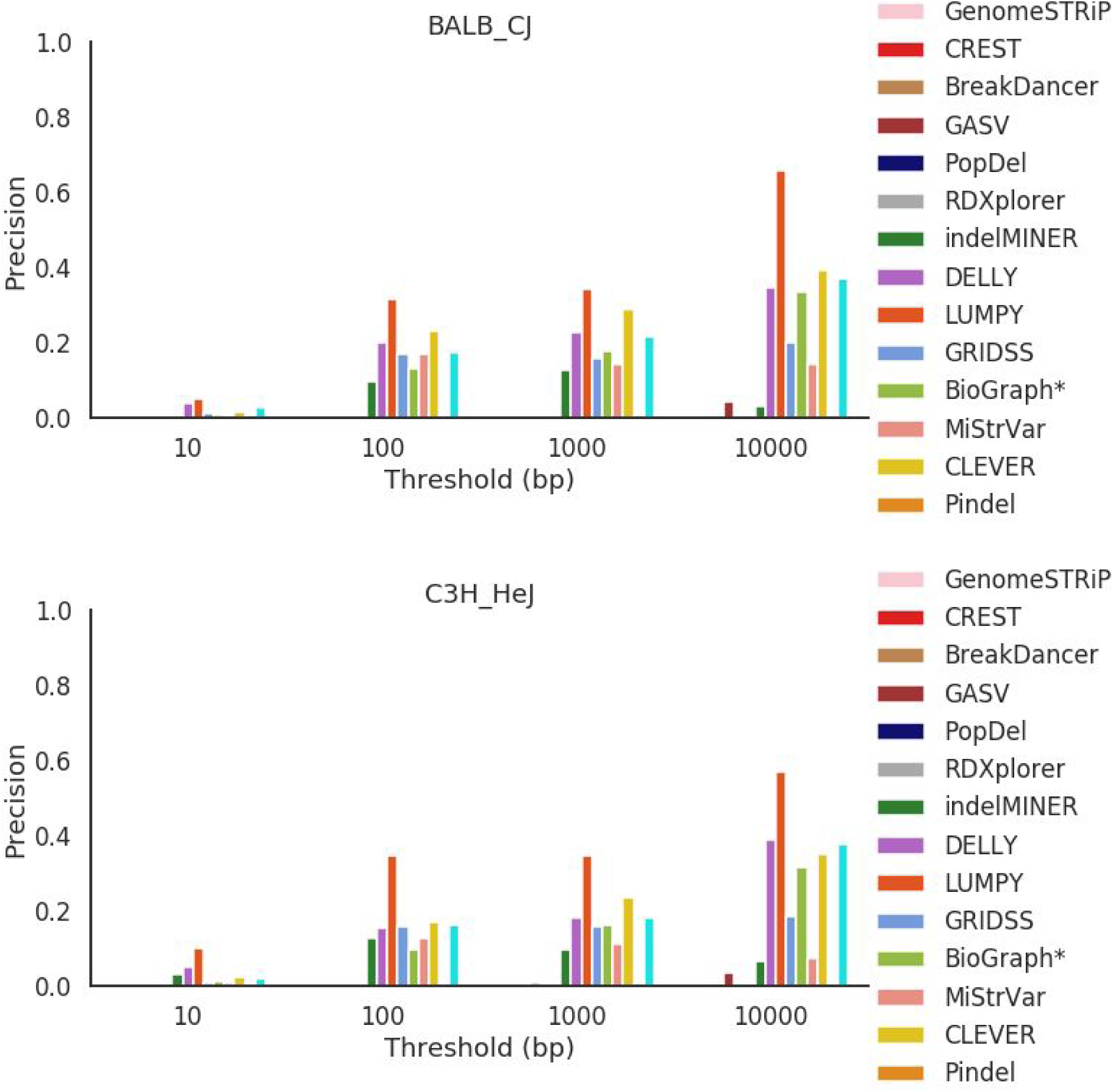
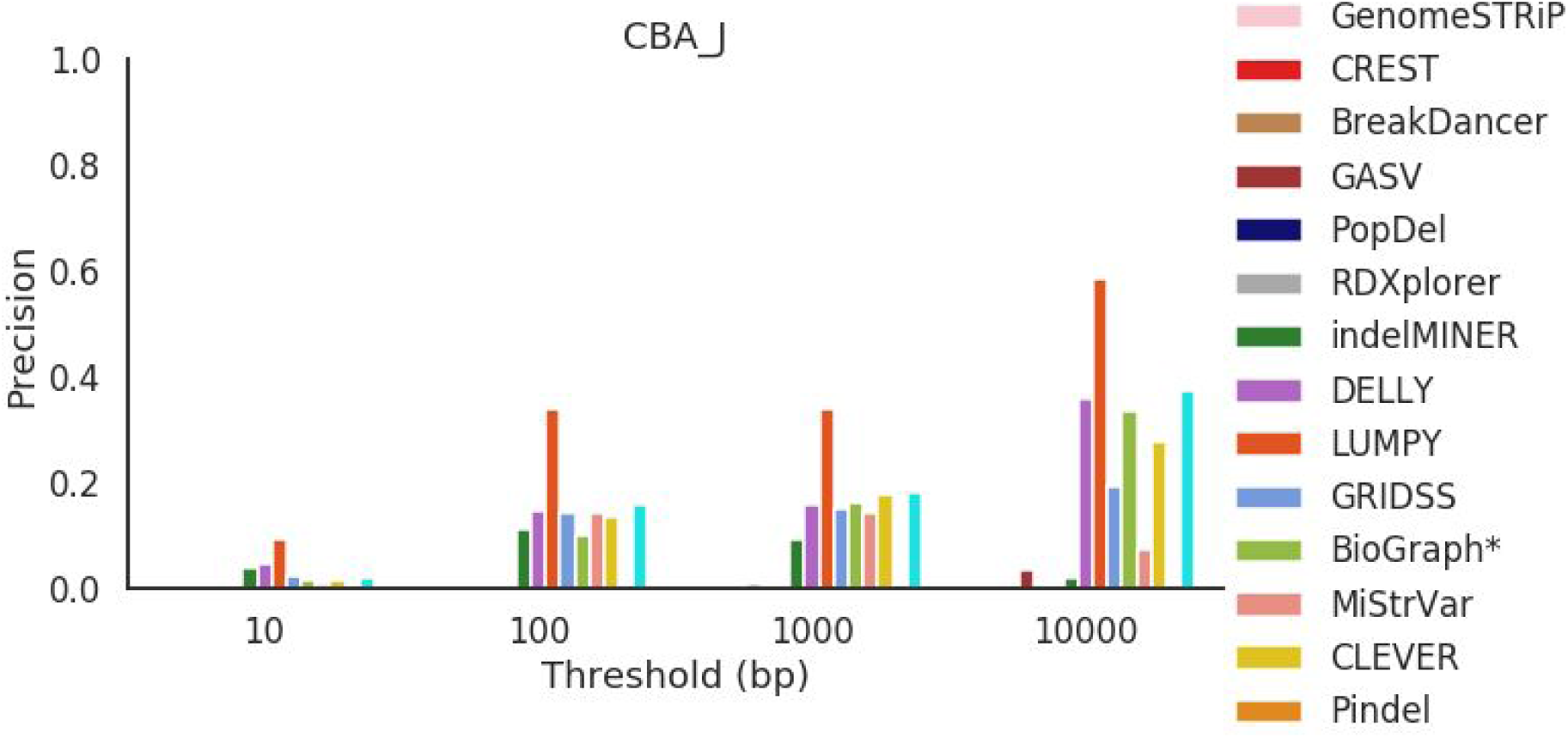
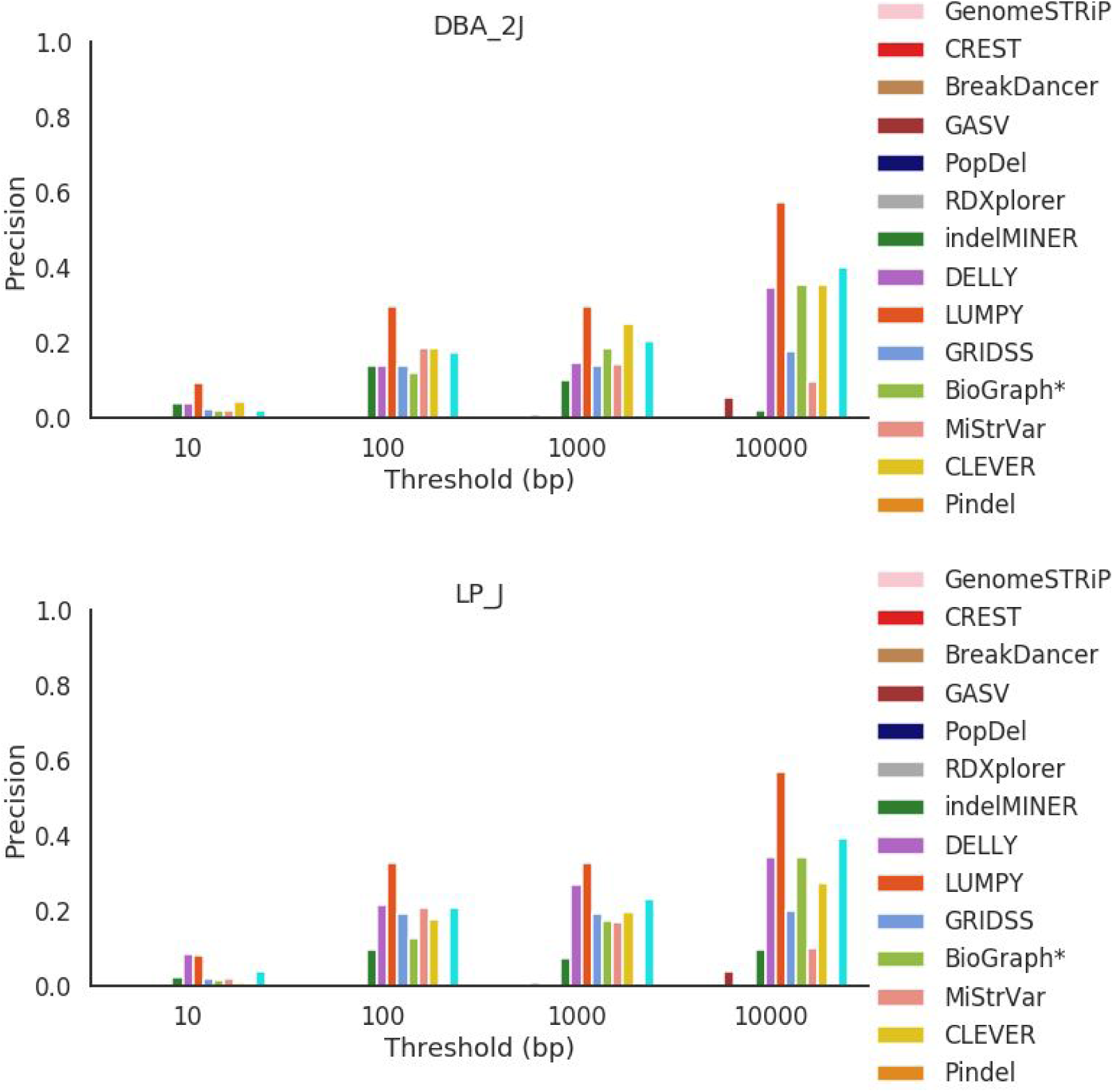
Precision across all mouse strains for deletions between 50 bp and 100 bp in length.

**Figure S15.**
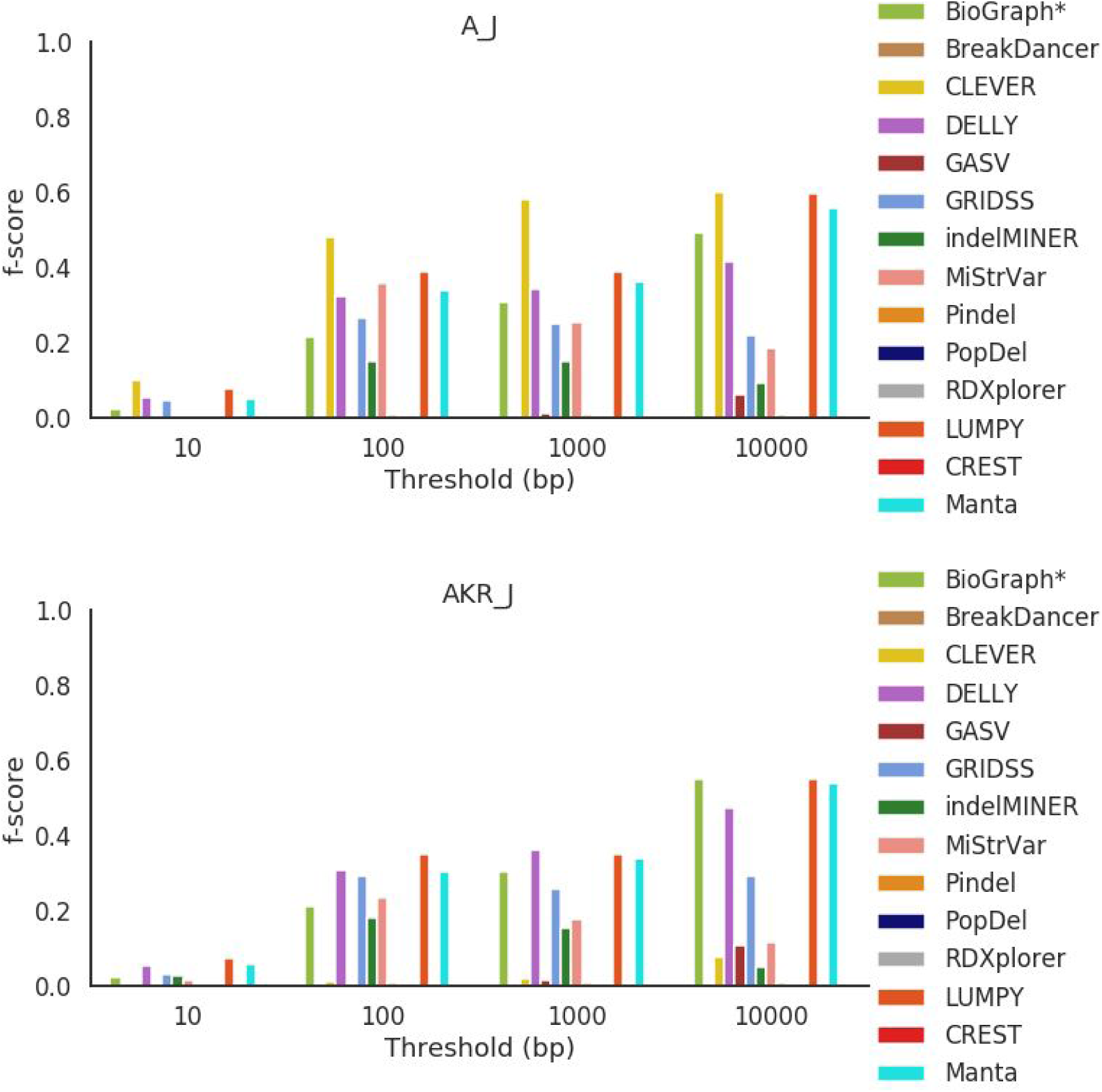
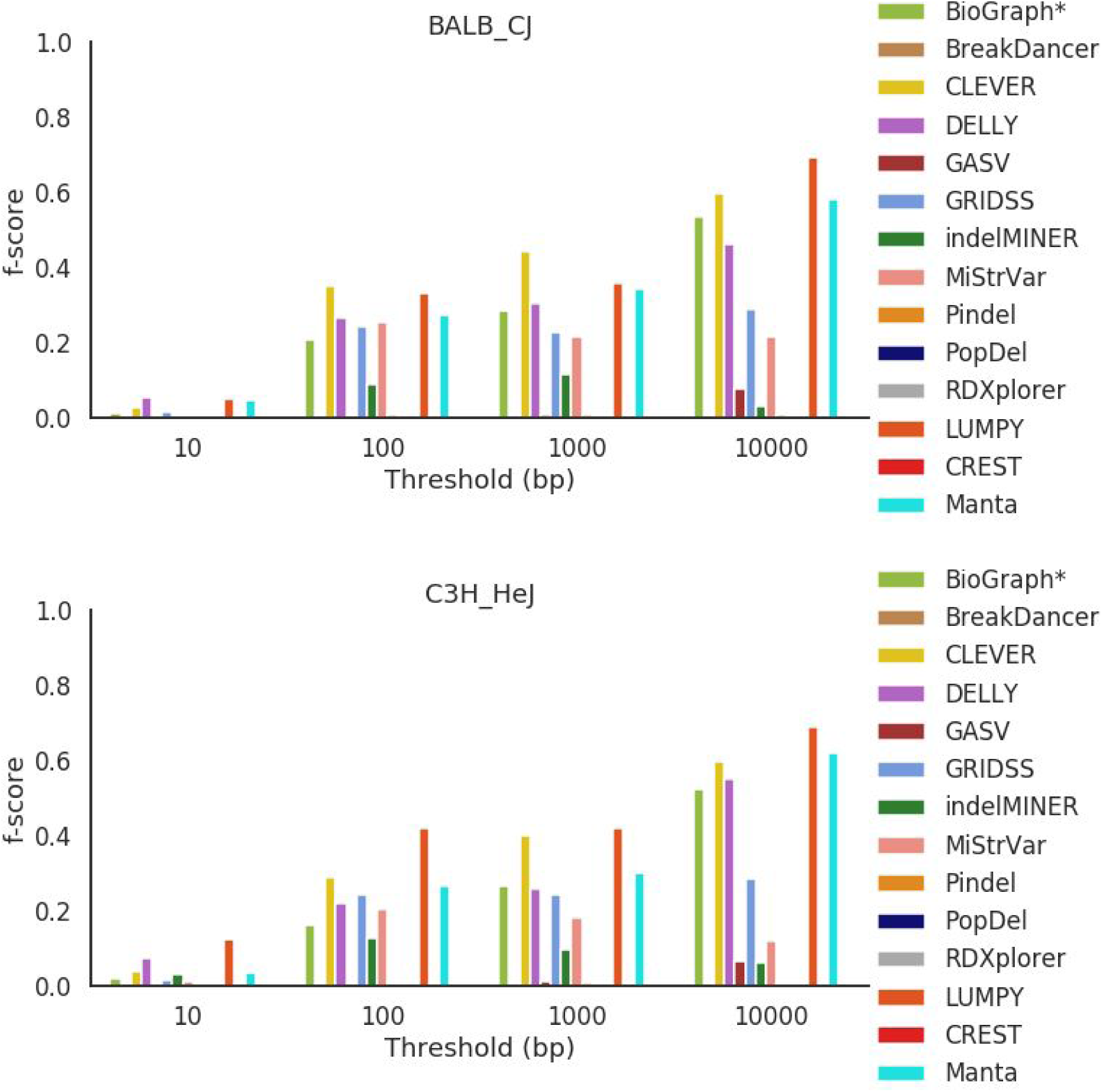
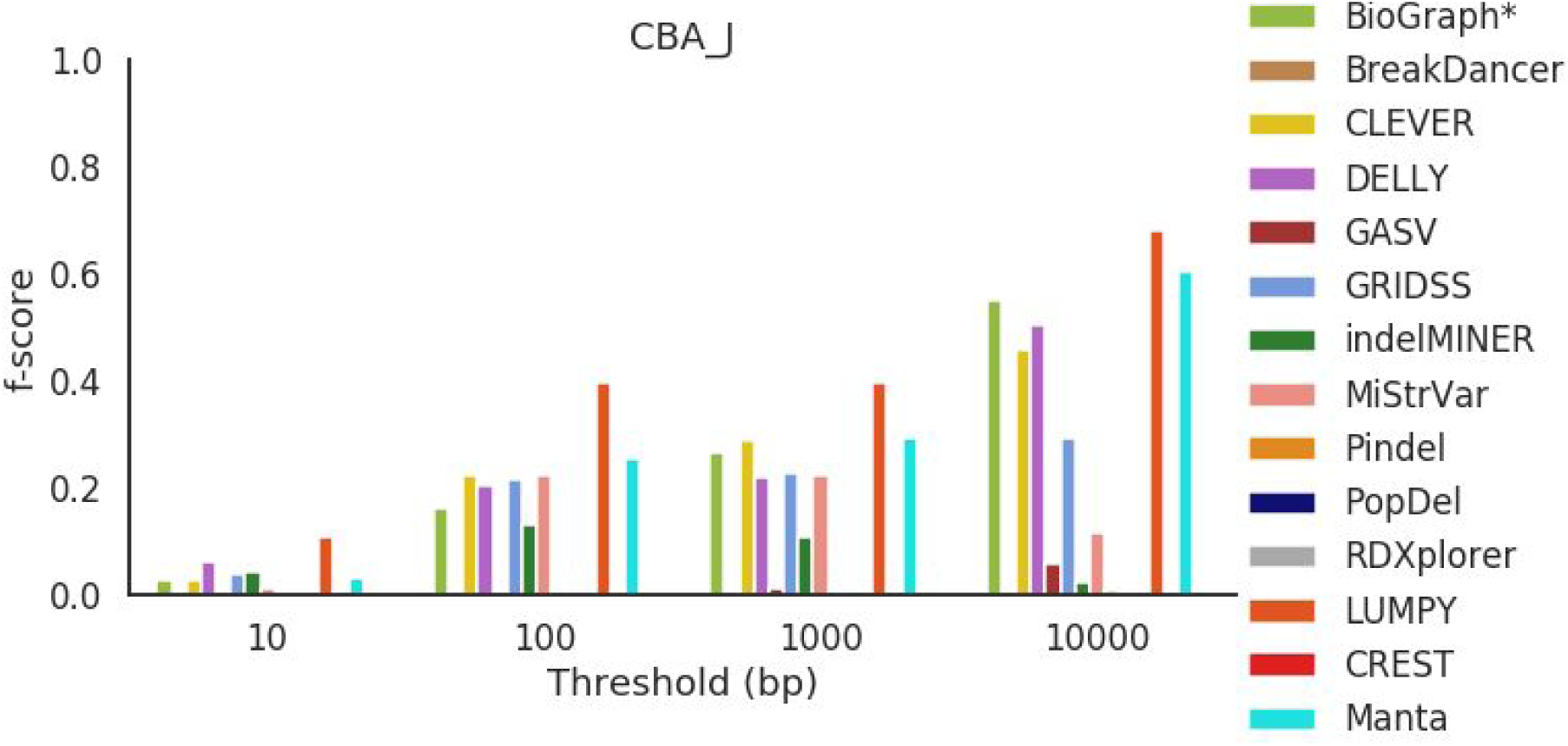
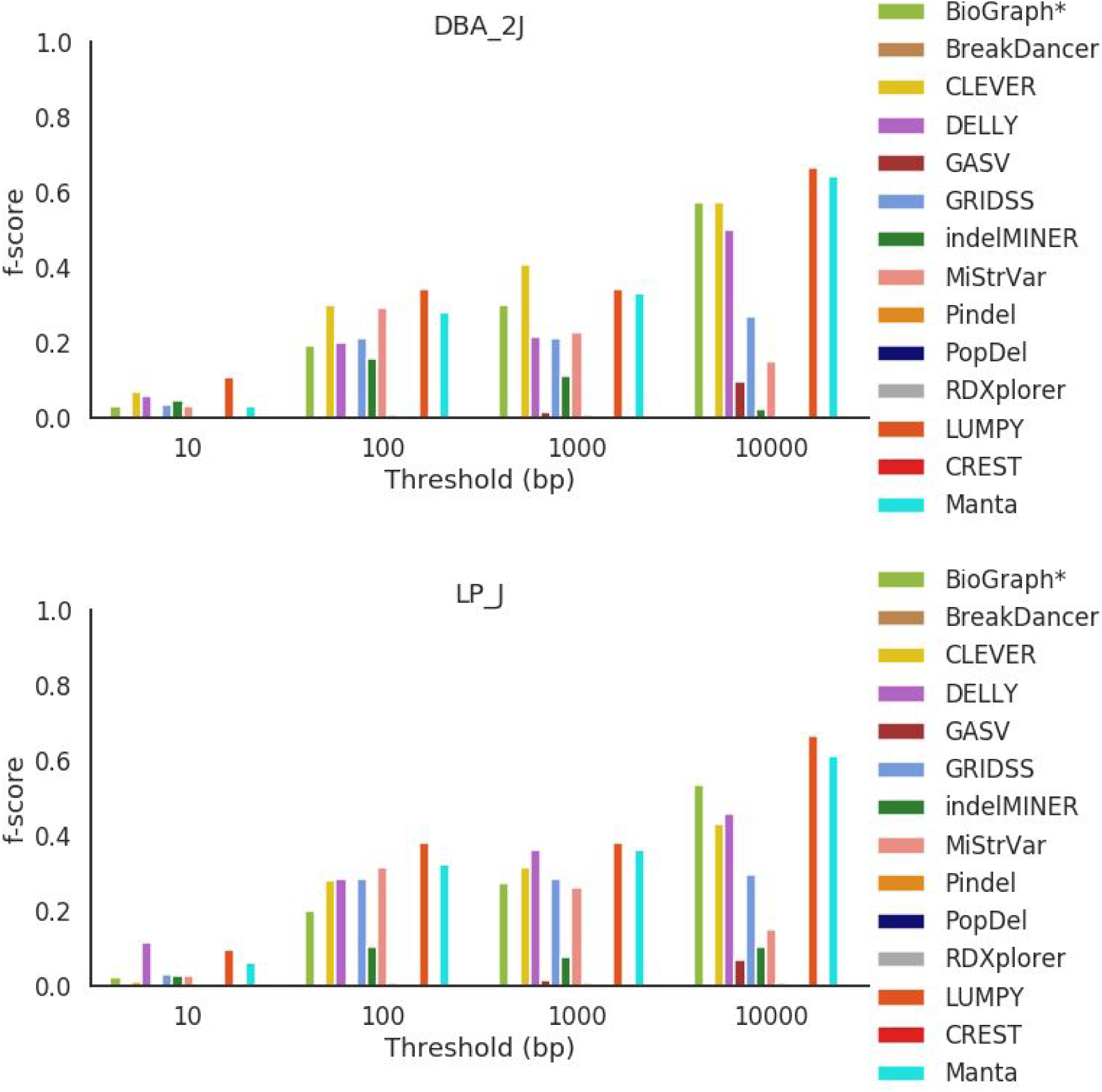
F-score across all mouse strains for deletions between 50 bp and 100 bp in length.

**Figure S16.**
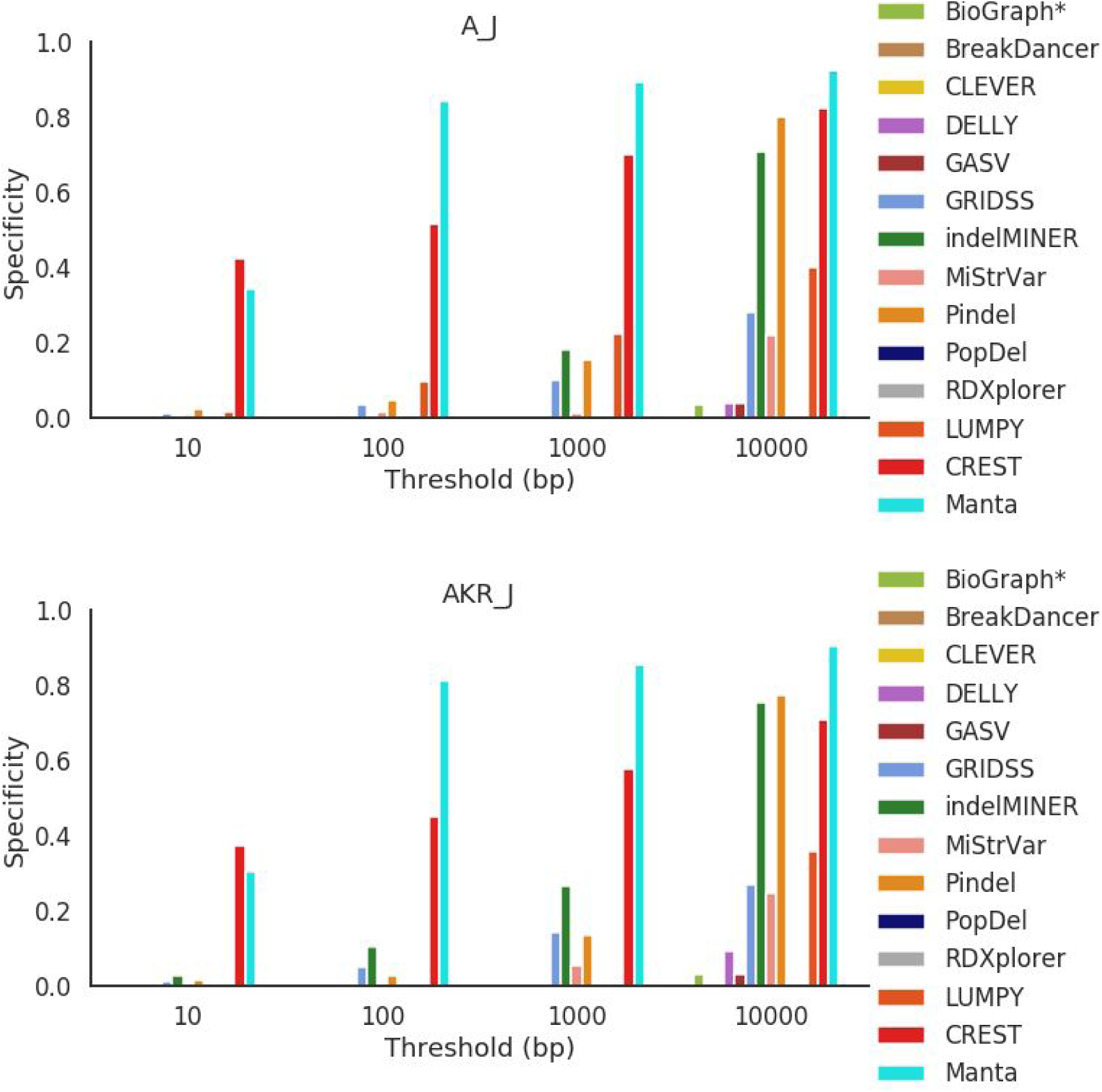
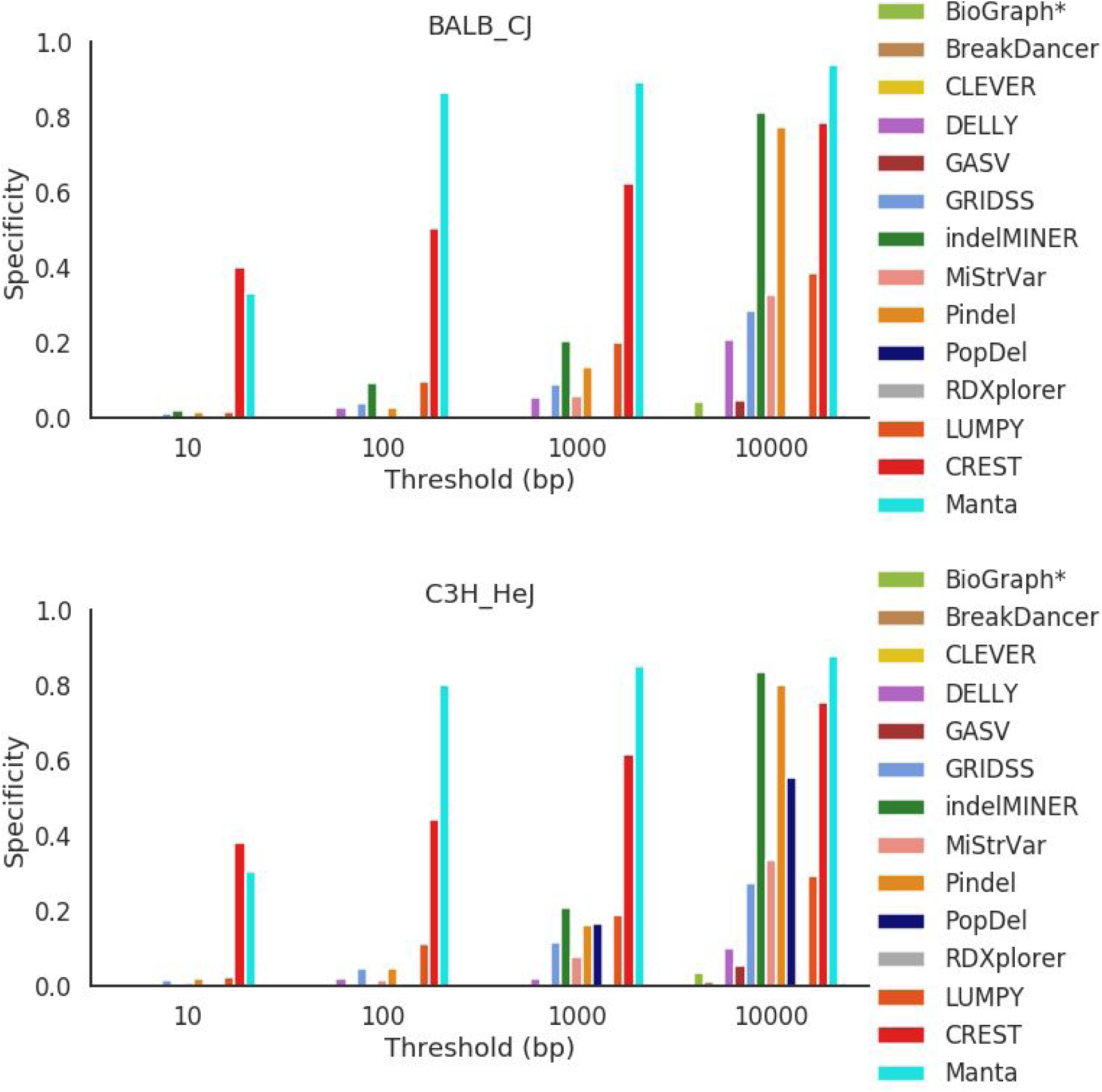
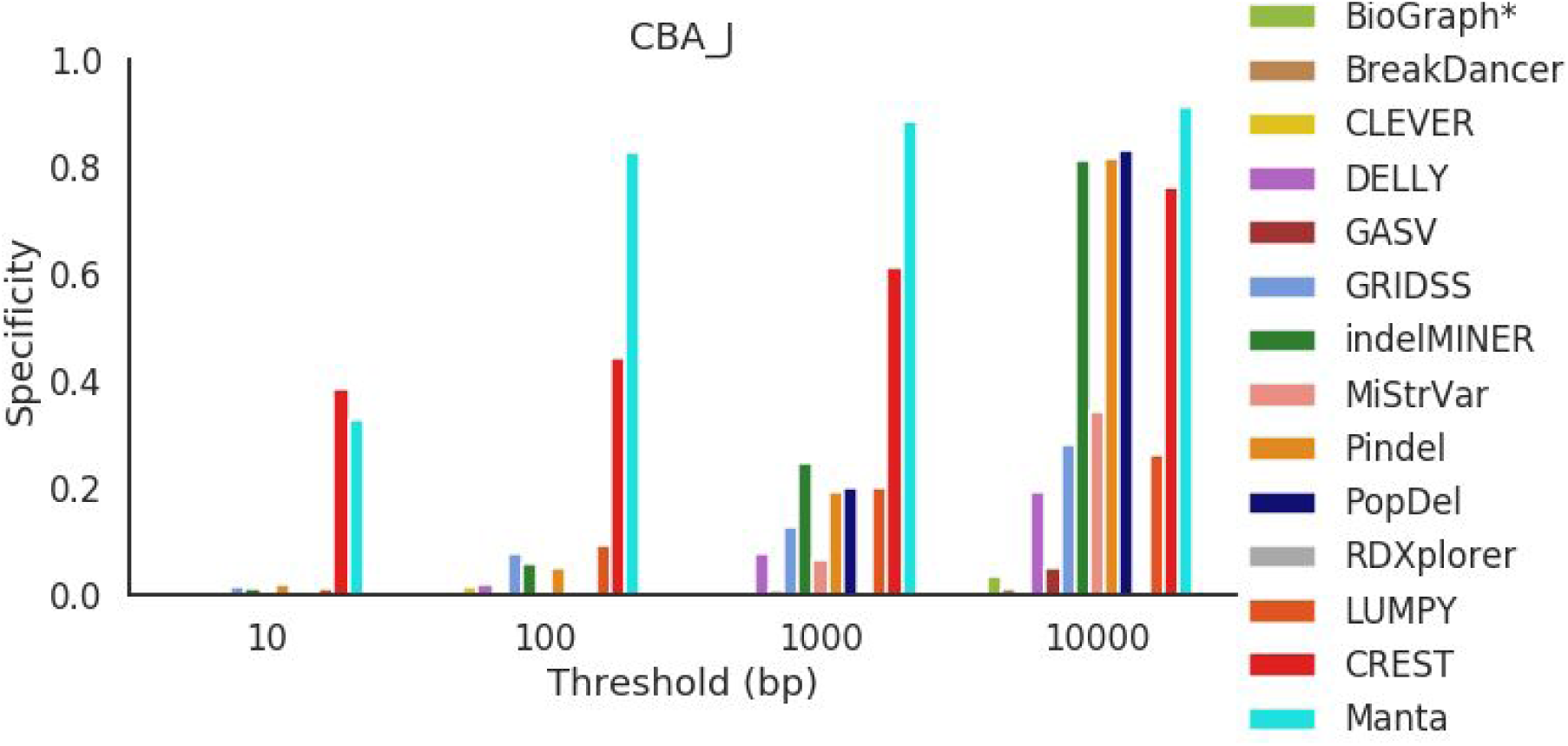
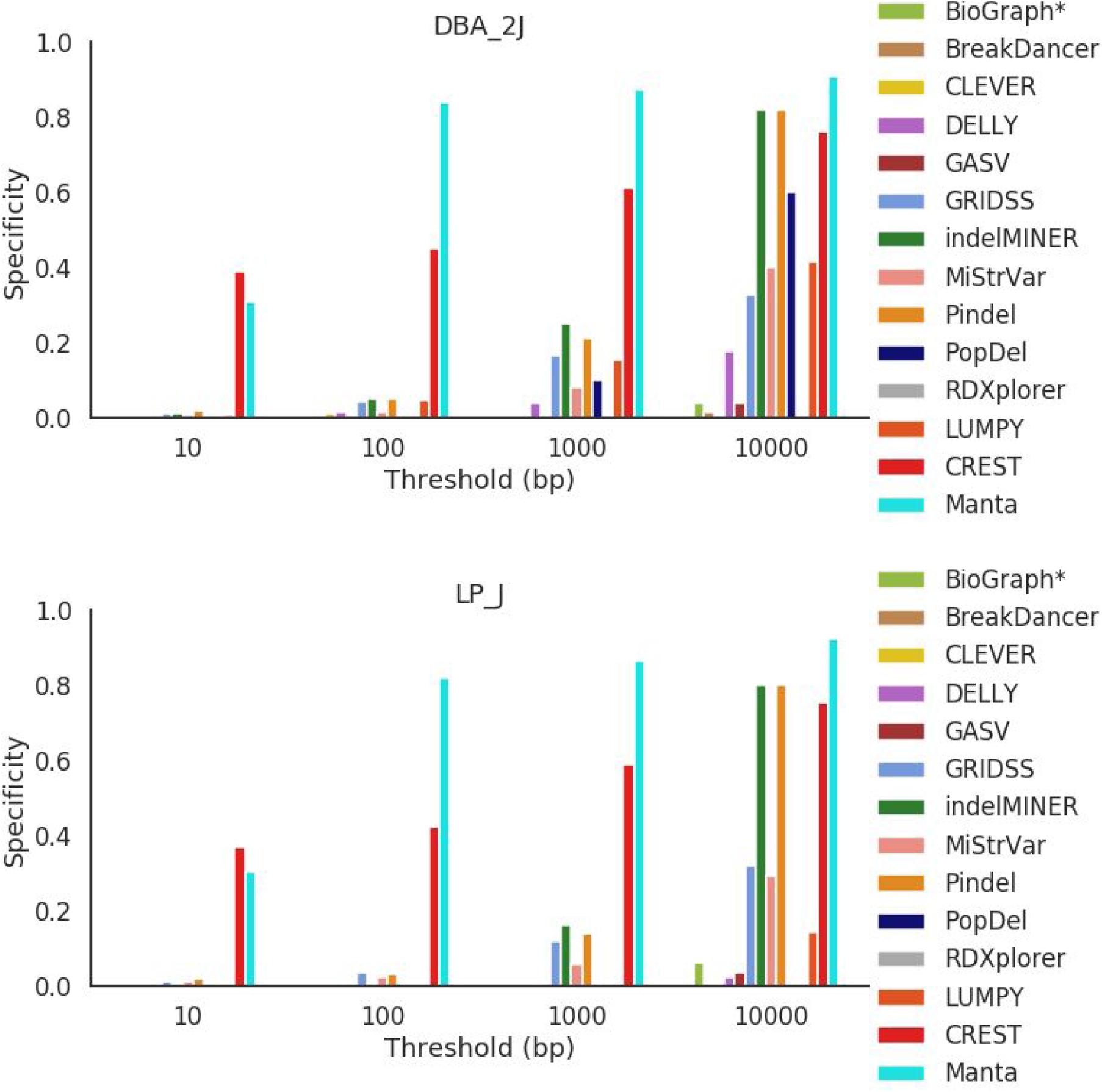
Specificity across all mouse strains for deletions between 100 bp and 500 bp in length.

**Figure S17.**
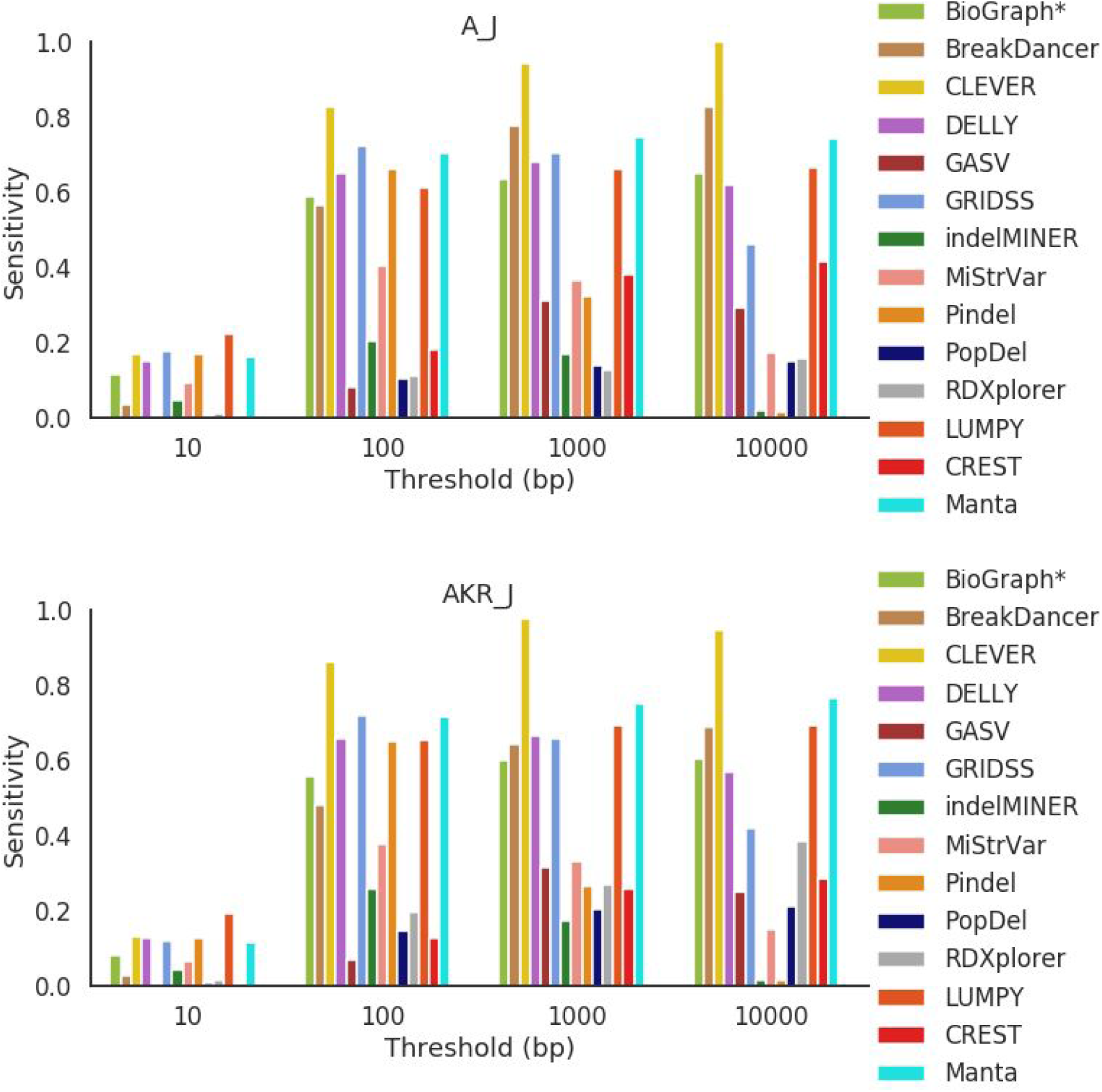
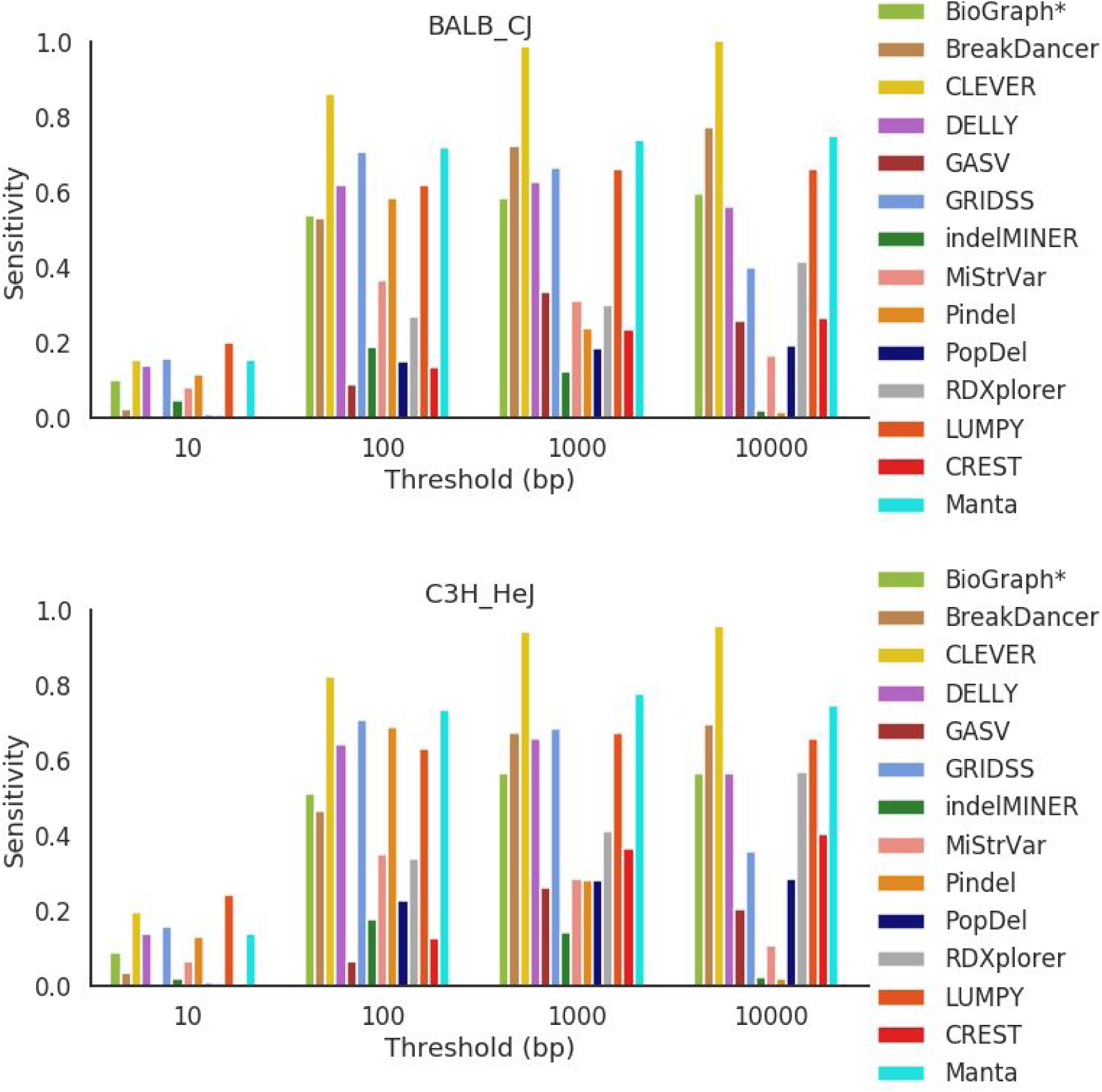
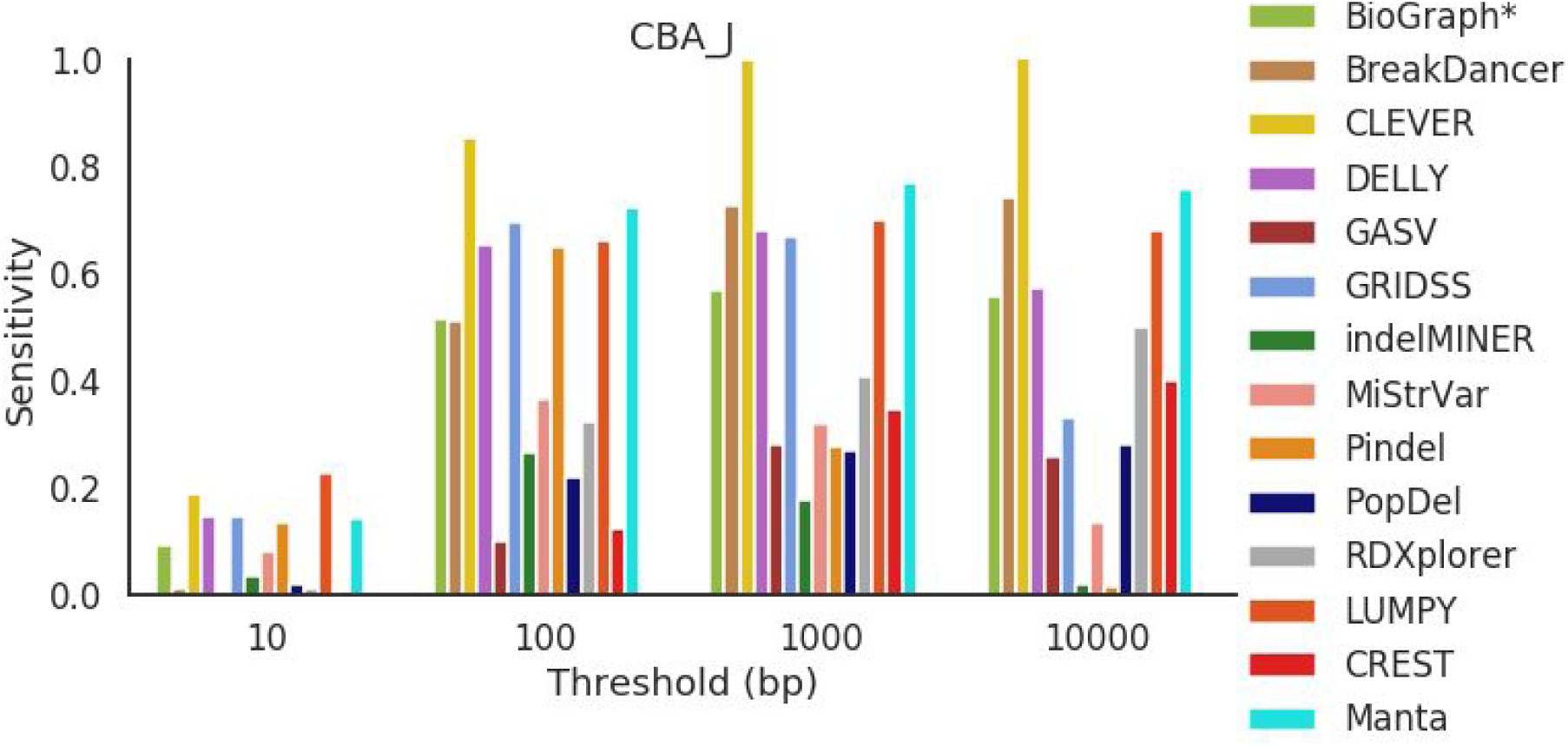
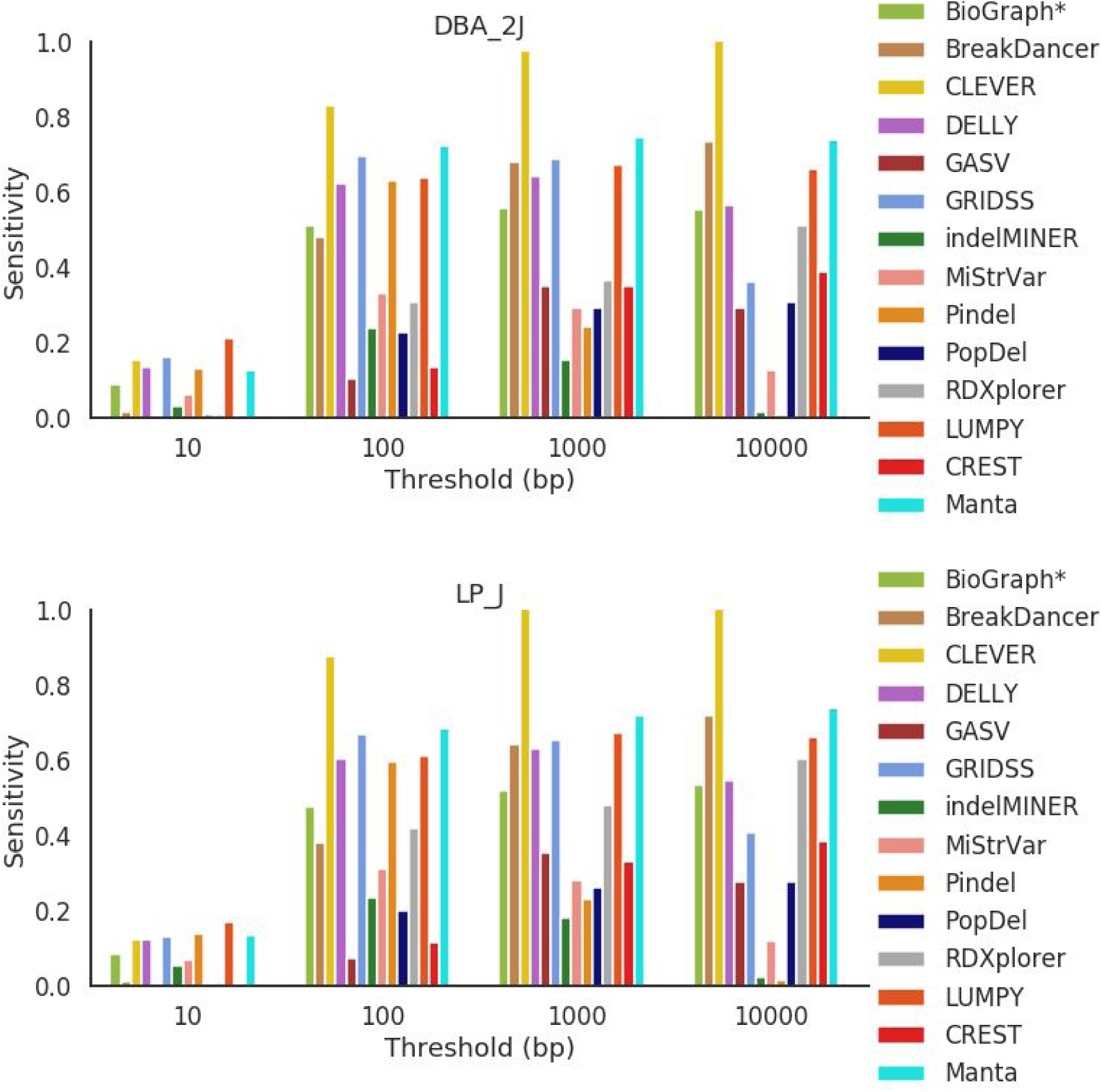
Sensitivity across all mouse strains for deletions between 50 bp and 100 bp in length.

**Figure S18.**
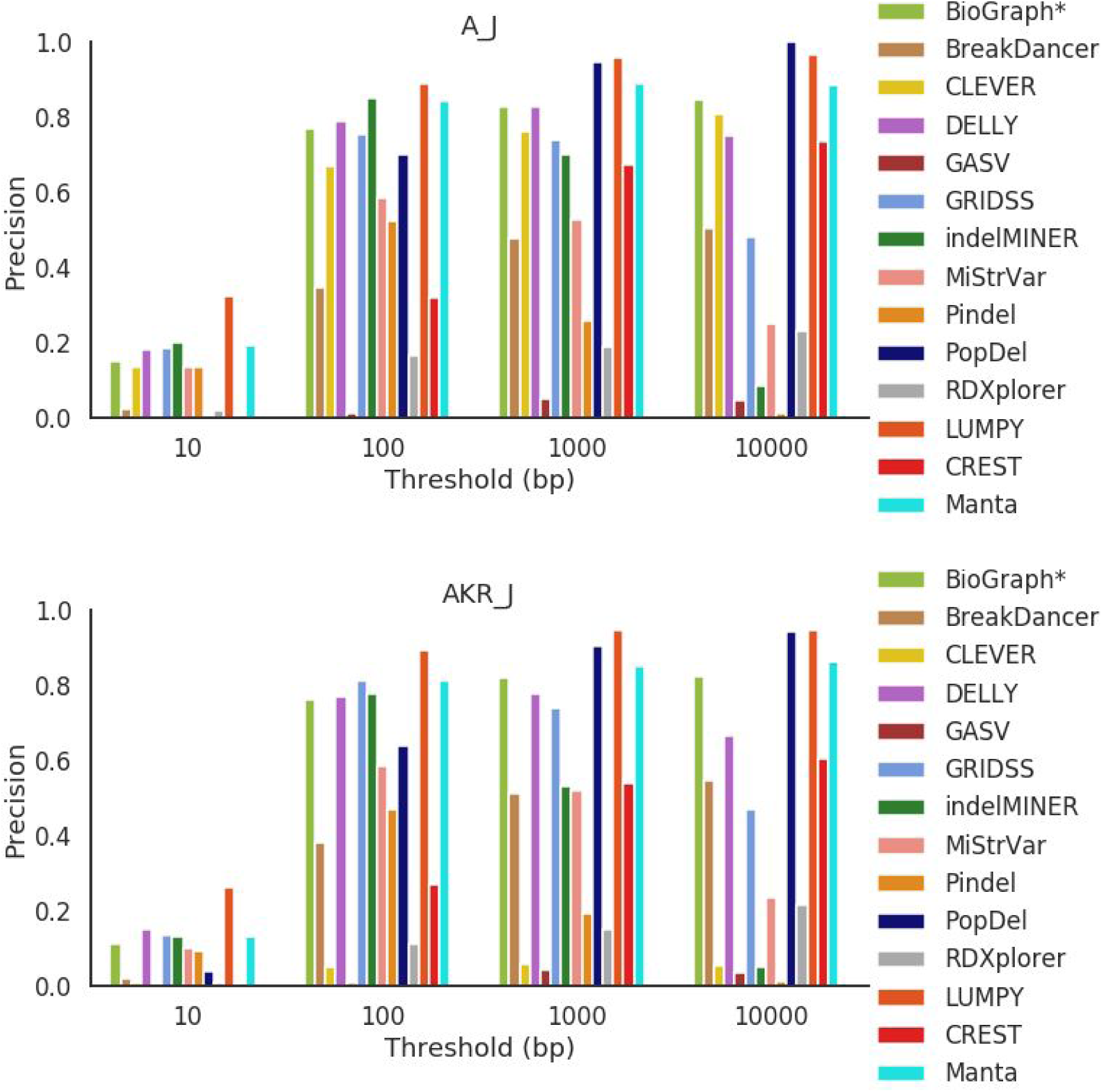
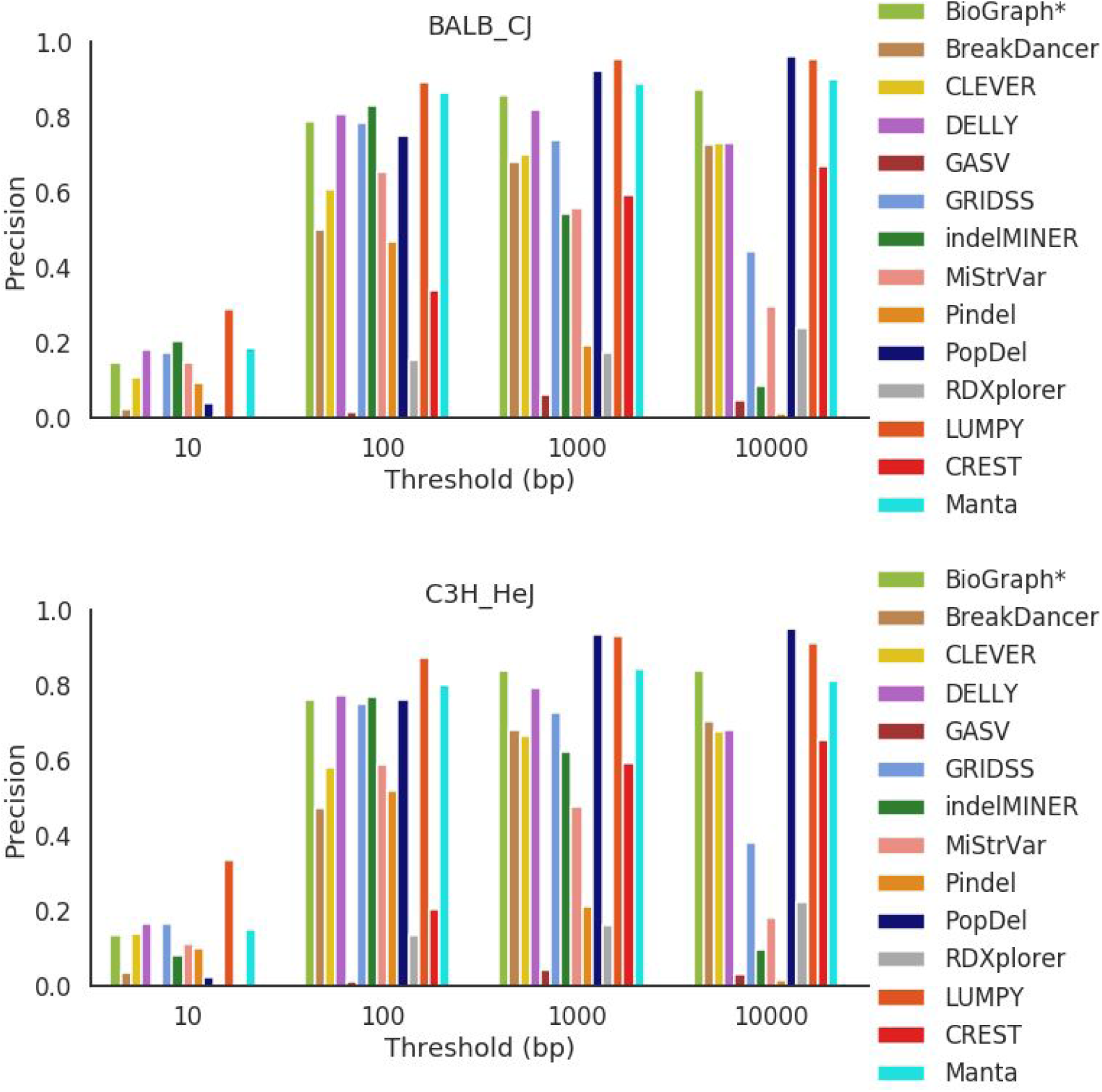
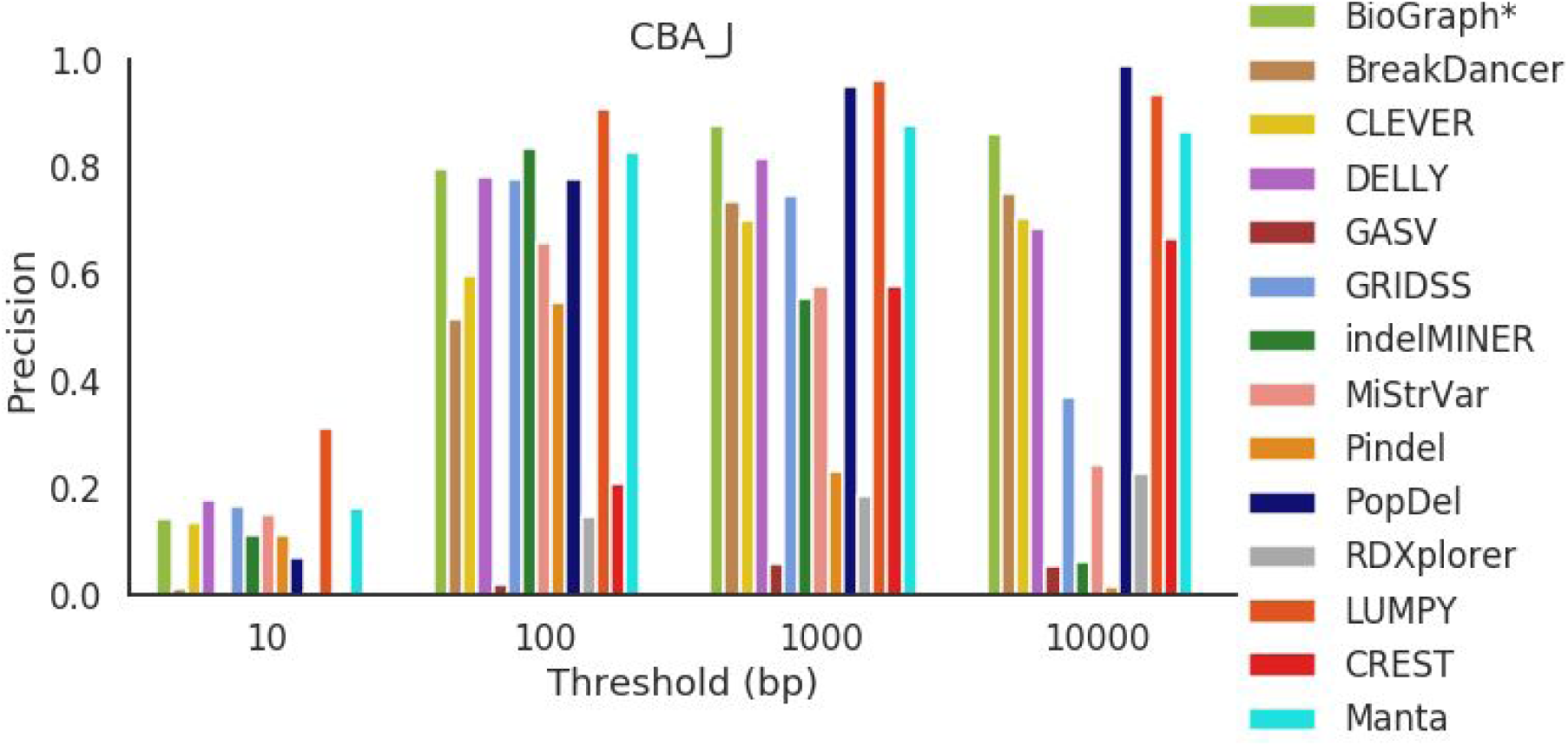
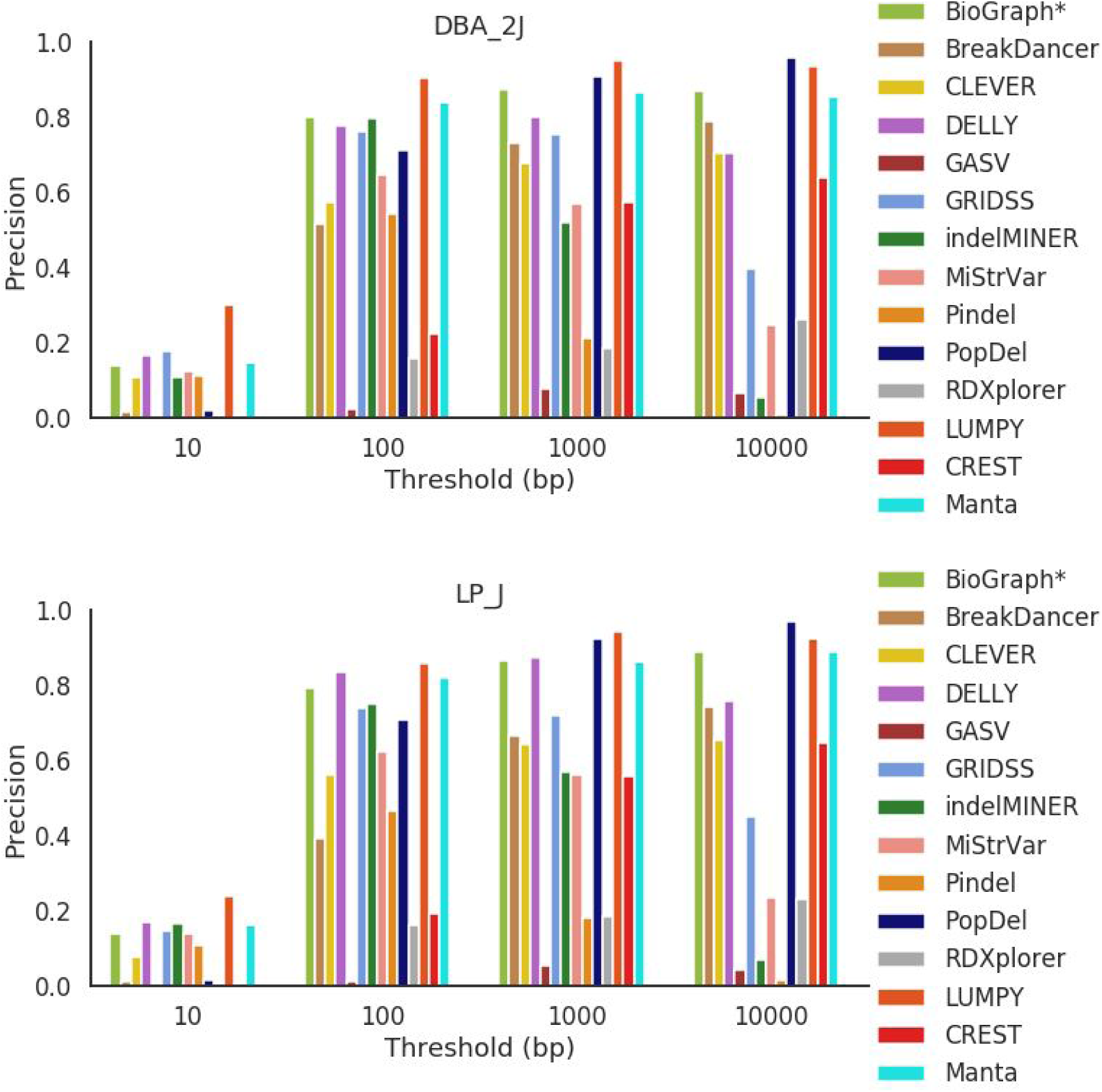
Precision across all mouse strains for deletions between 100 bp and 500 bp in length.

**Figure S19.**
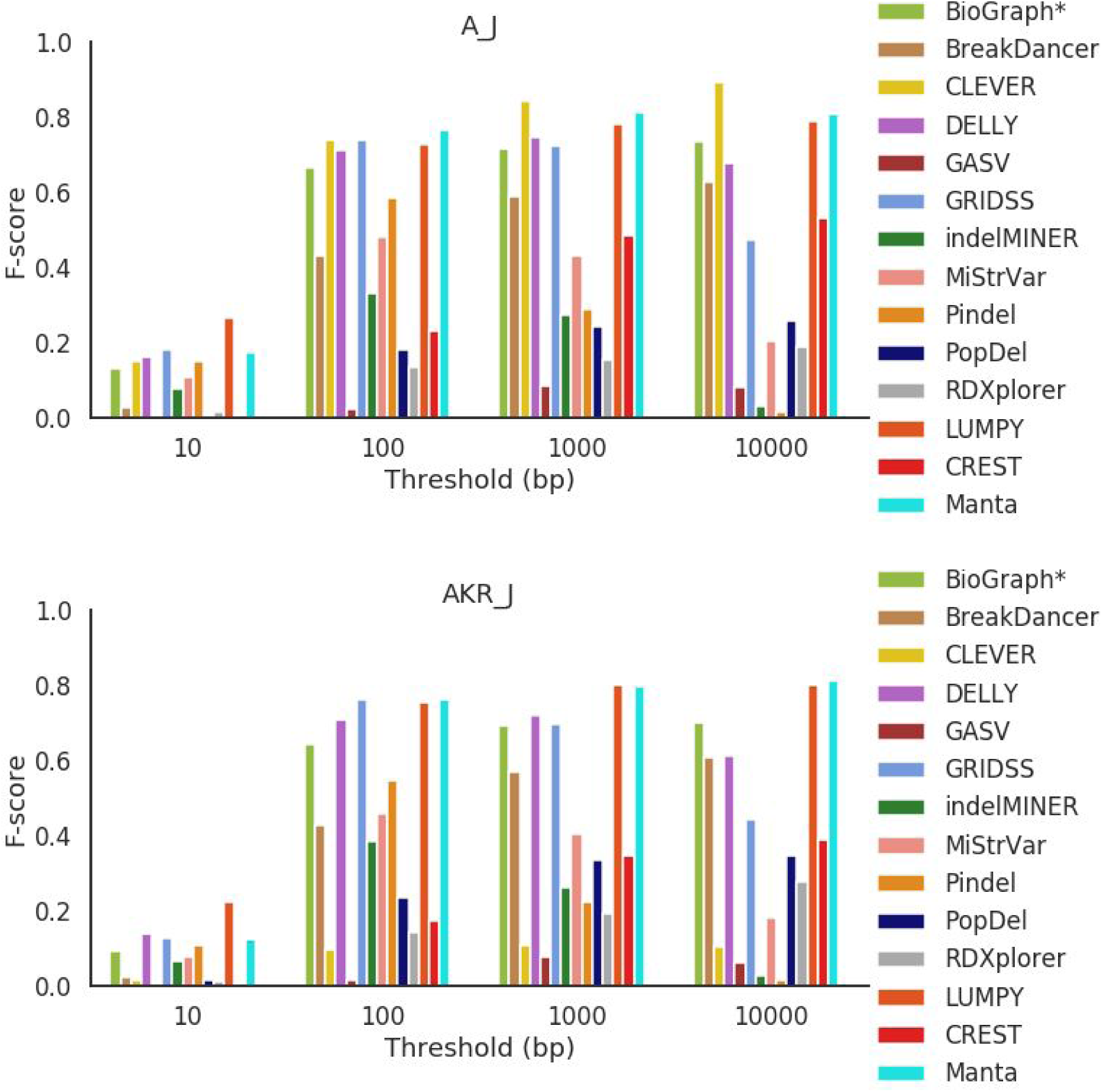
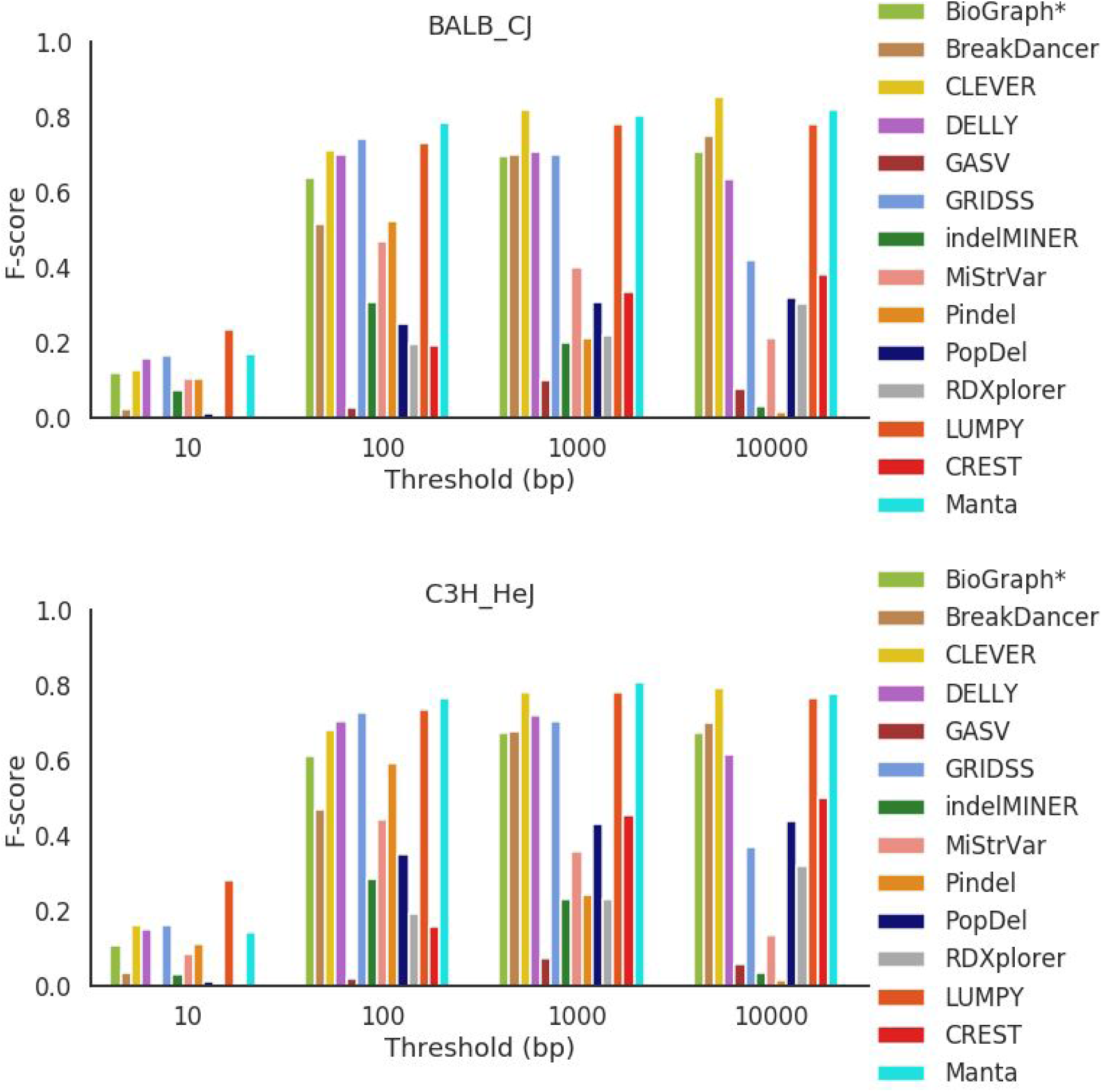
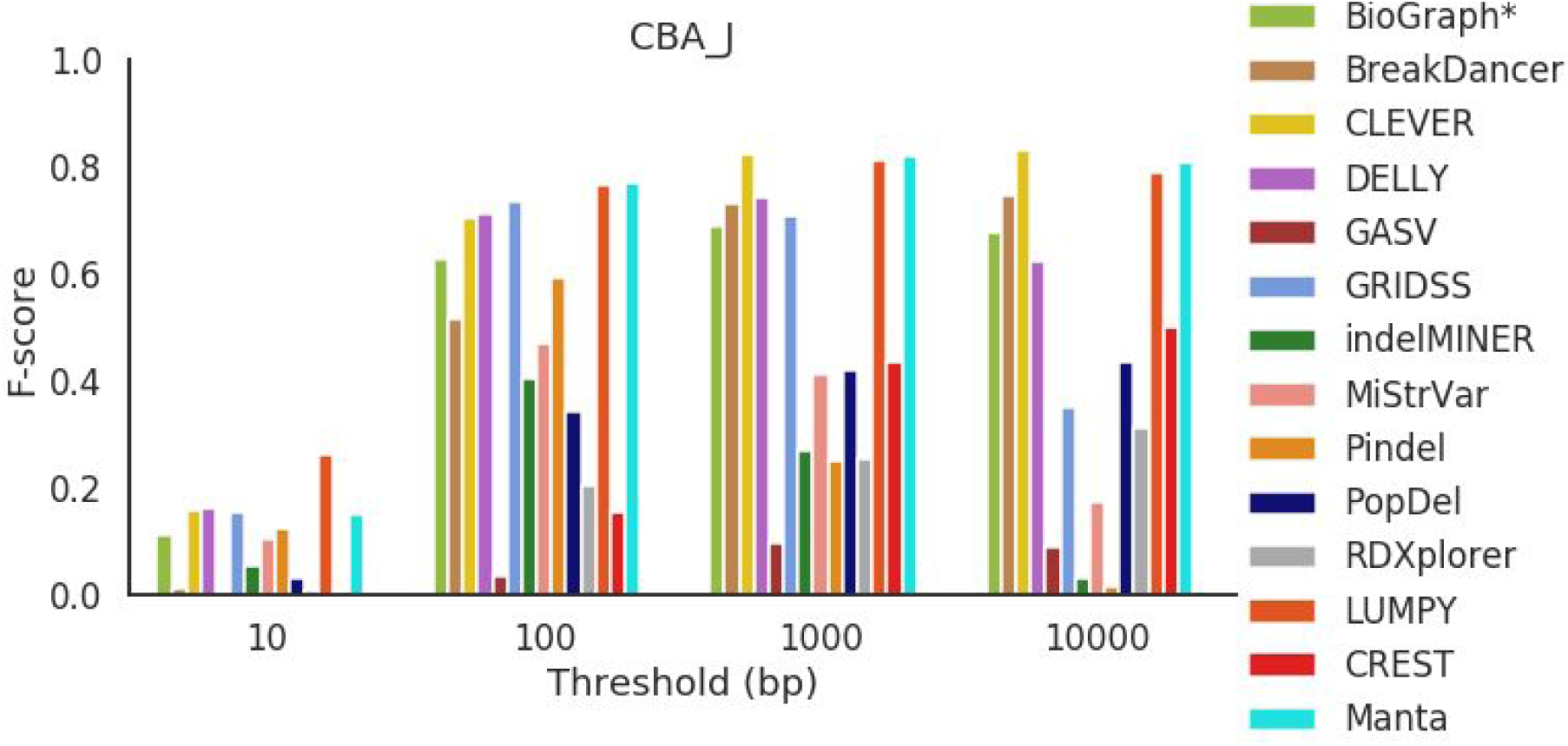
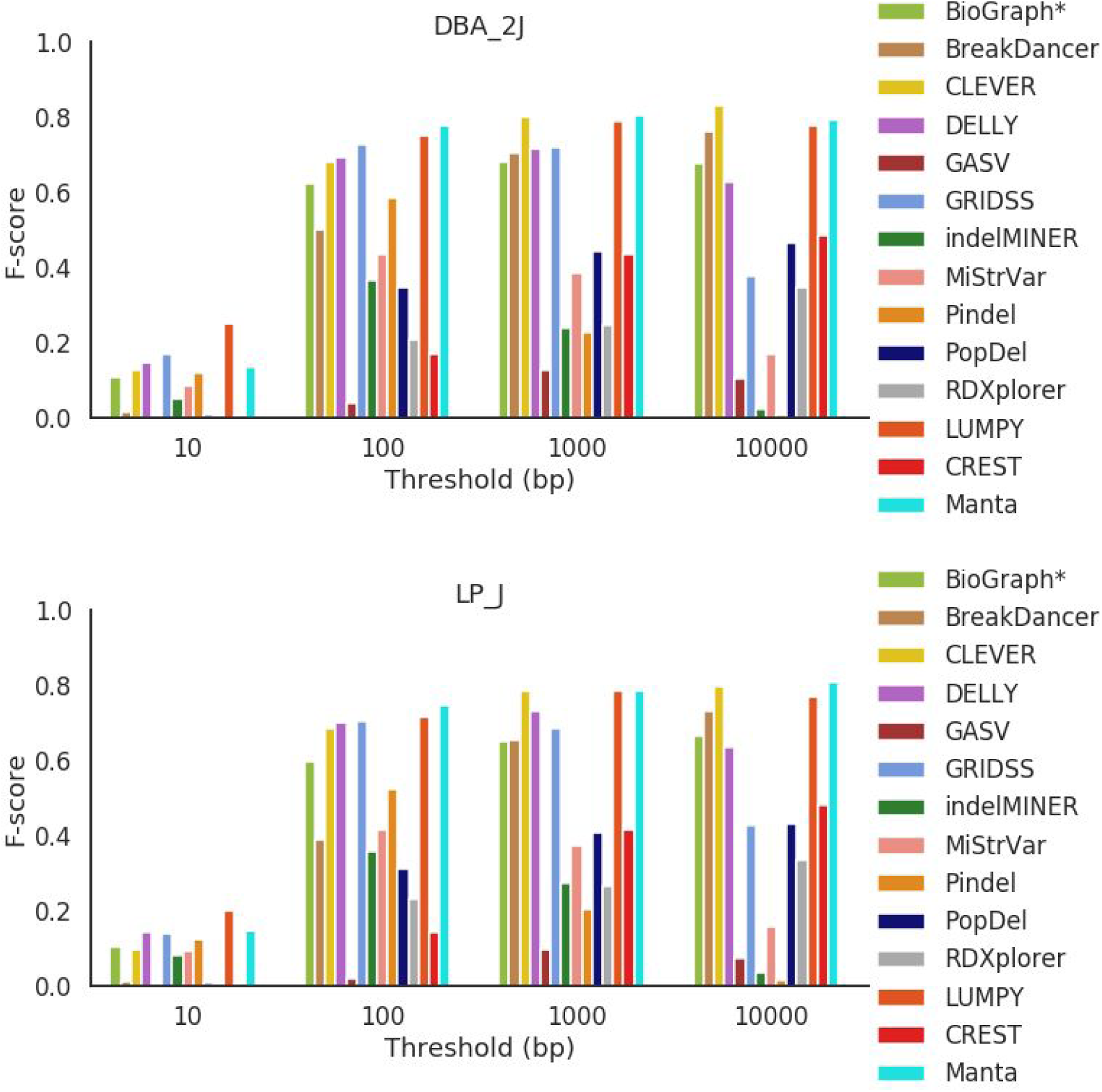
F-score across all mouse strains for deletions between 100 bp and 500 bp in length.

**Figure S20.**
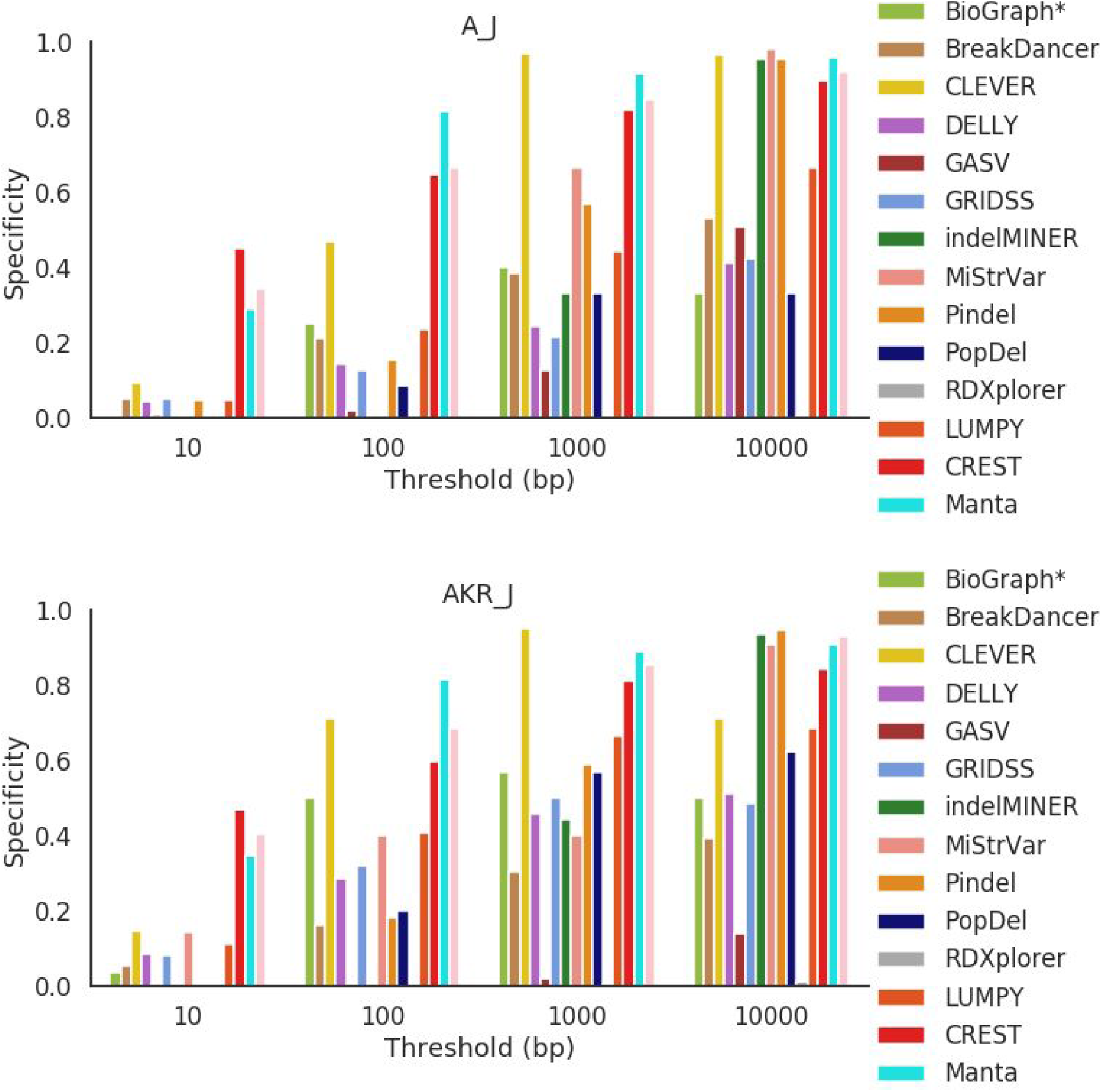
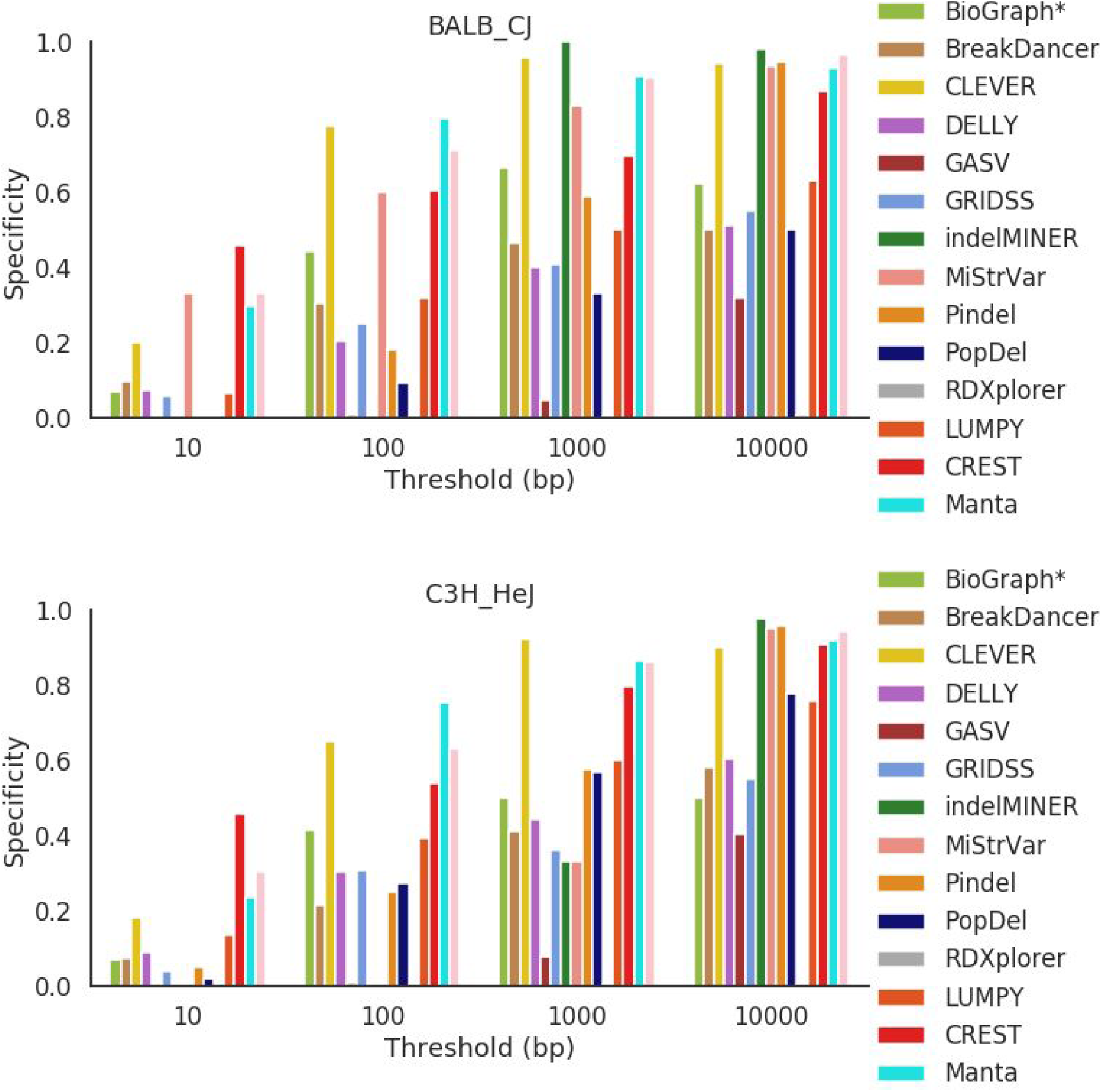
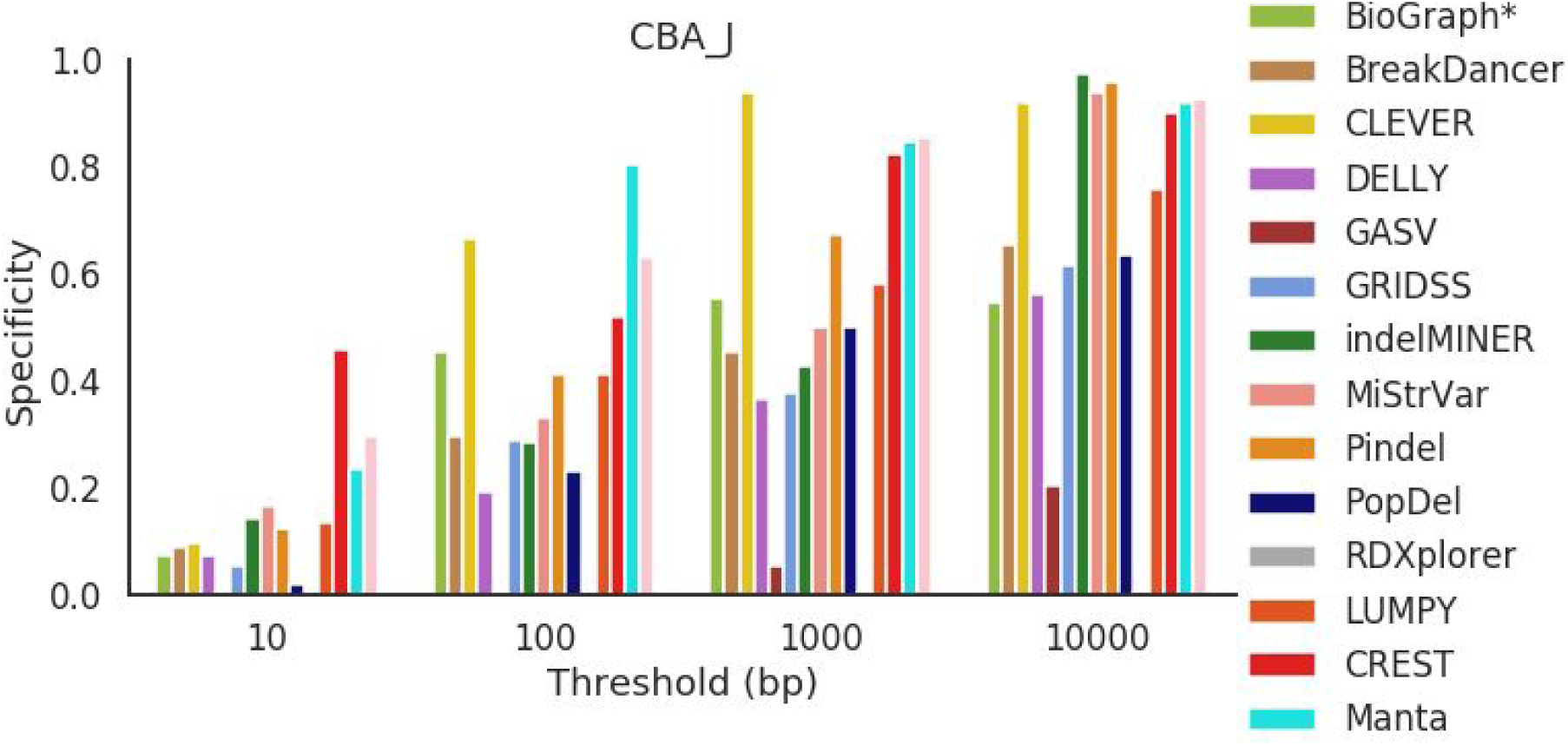
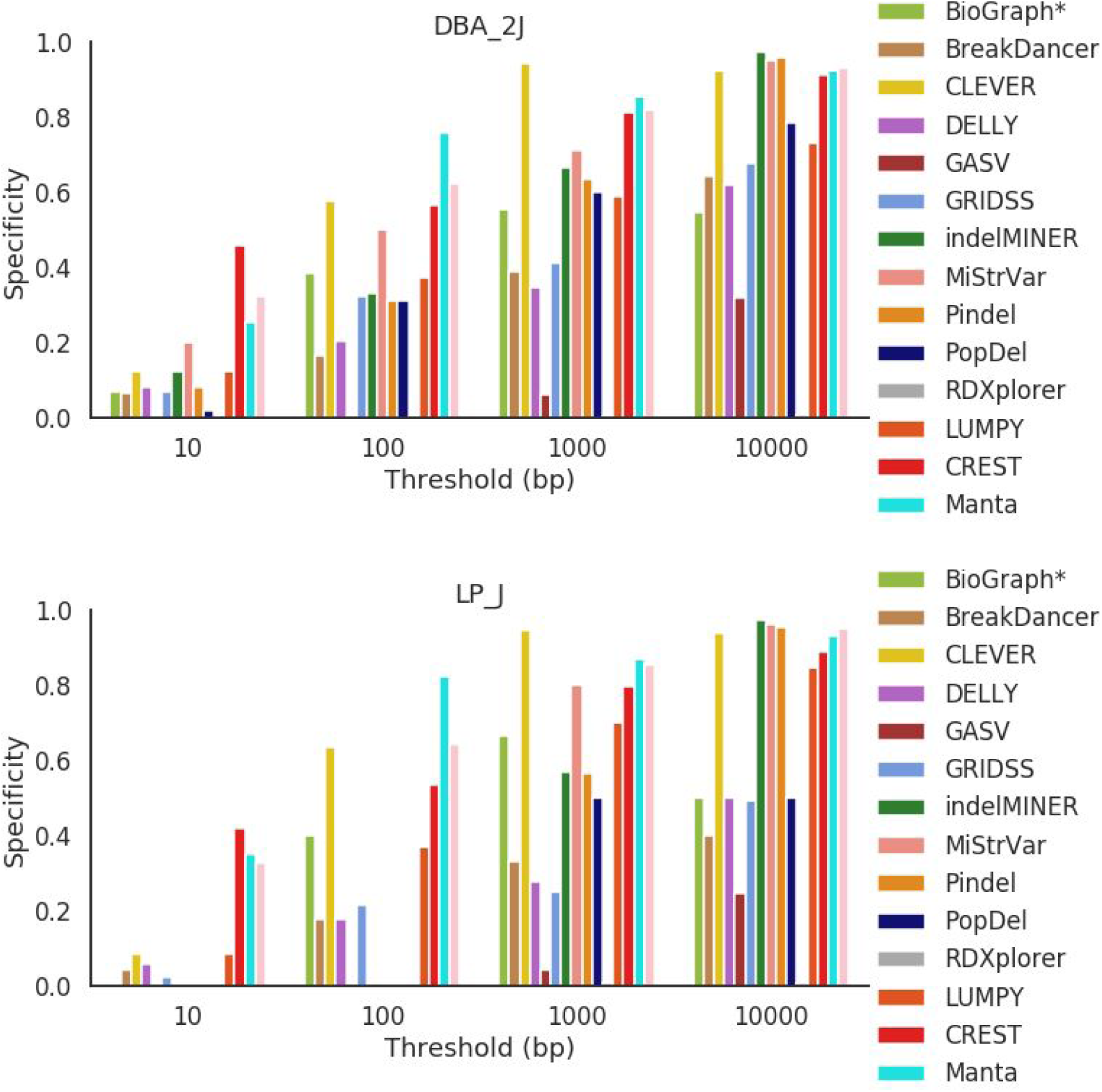
Specificity across all mouse strains for deletions between 500 bp and 1000 bp in length.

**Figure S21.**
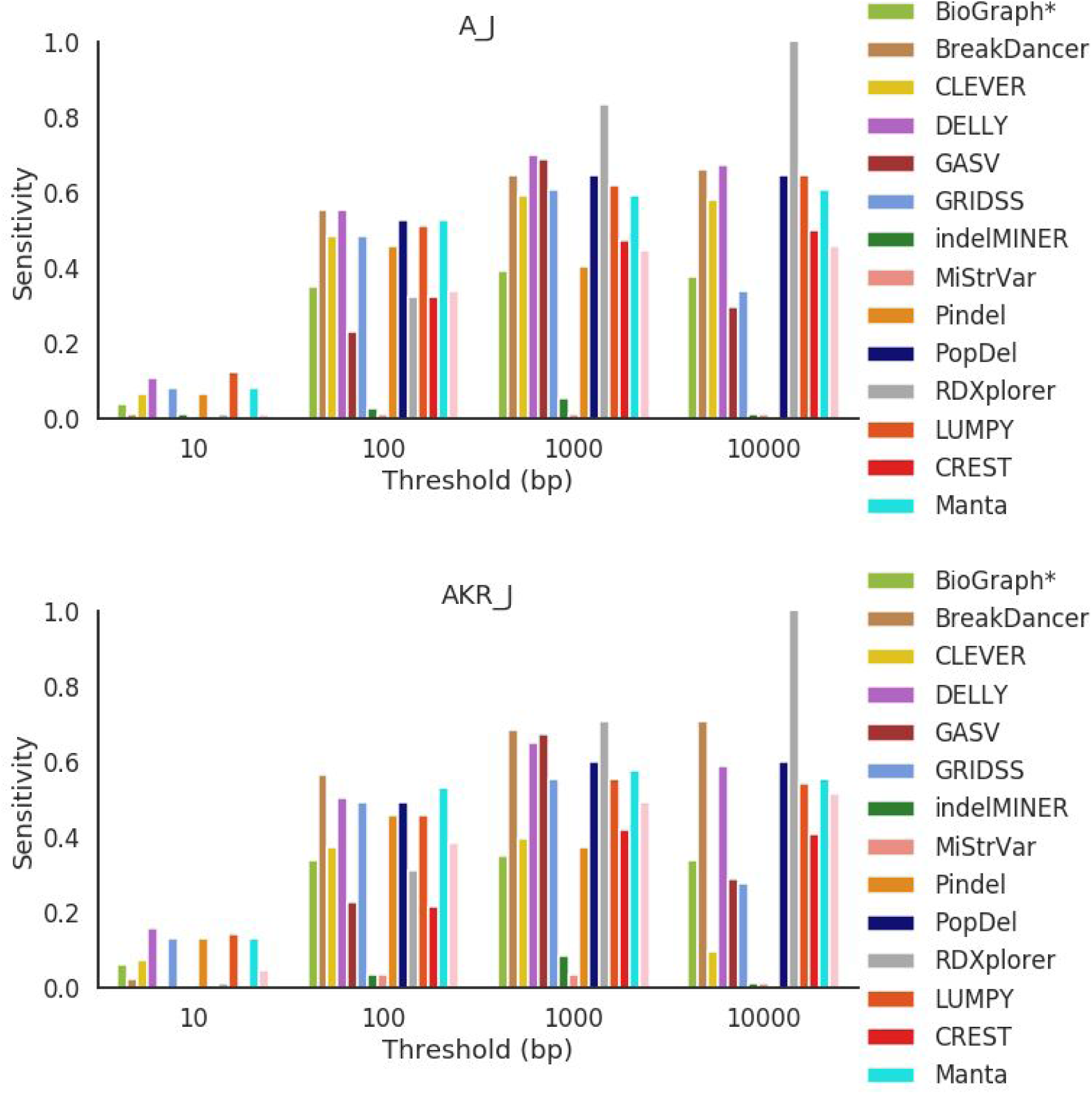
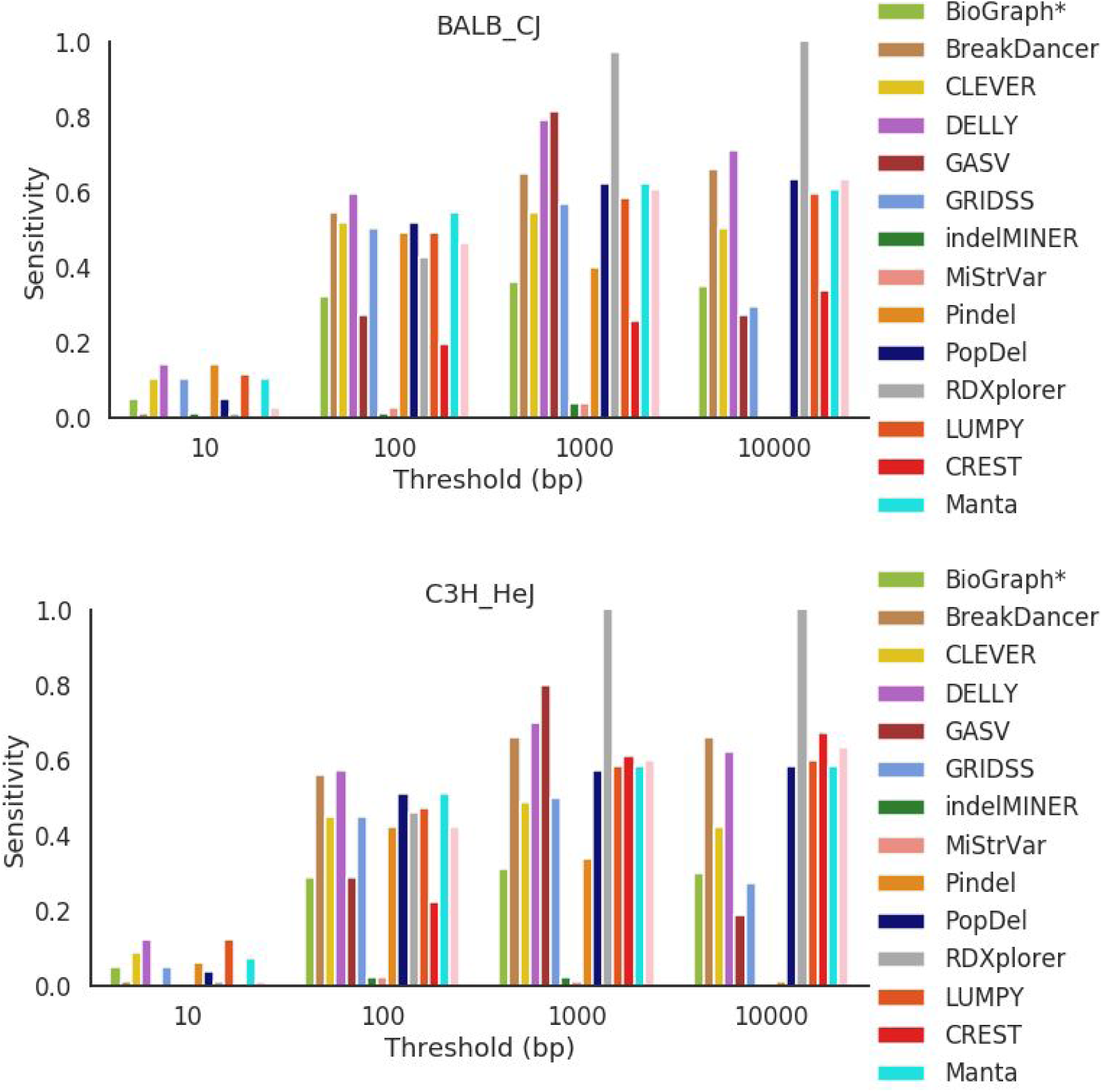
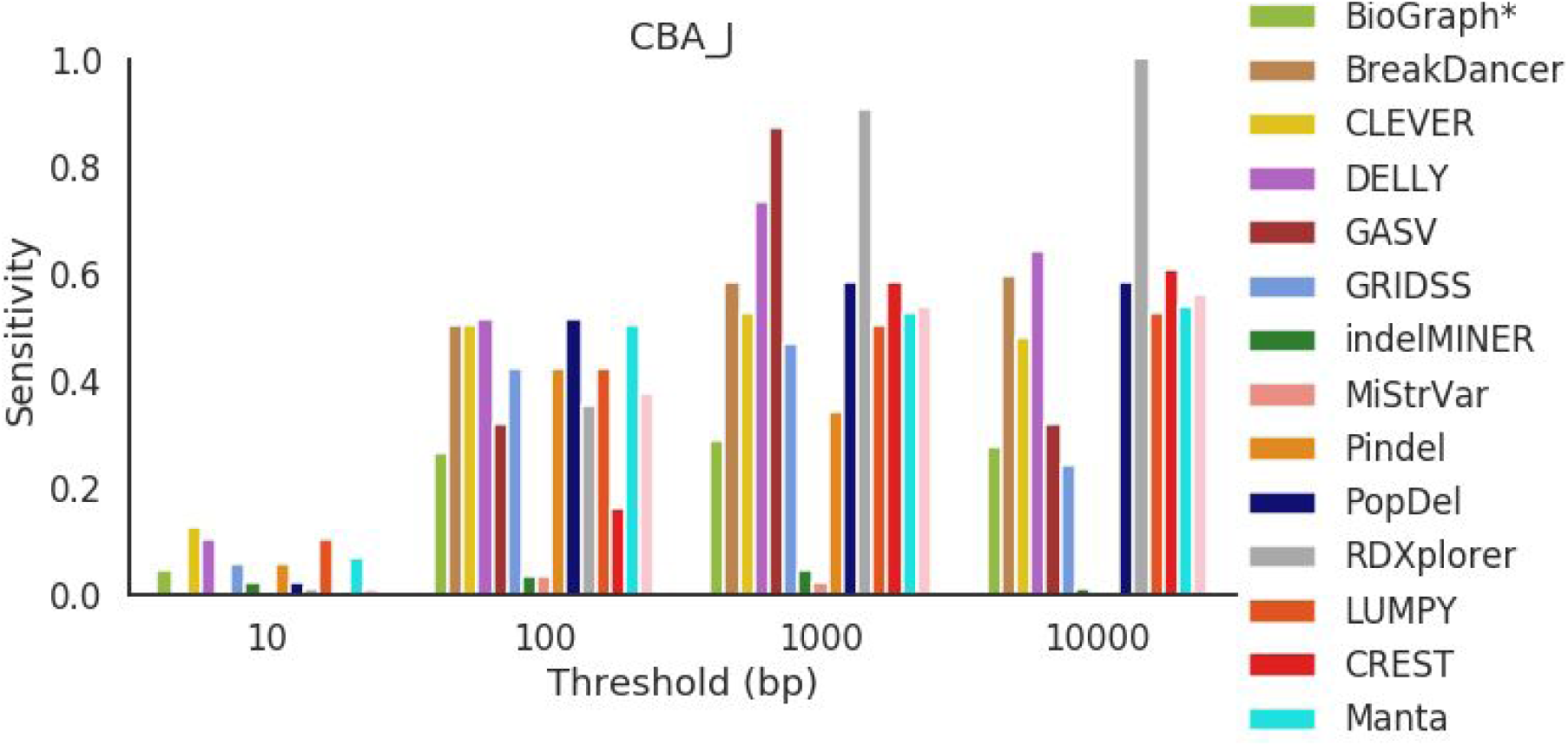
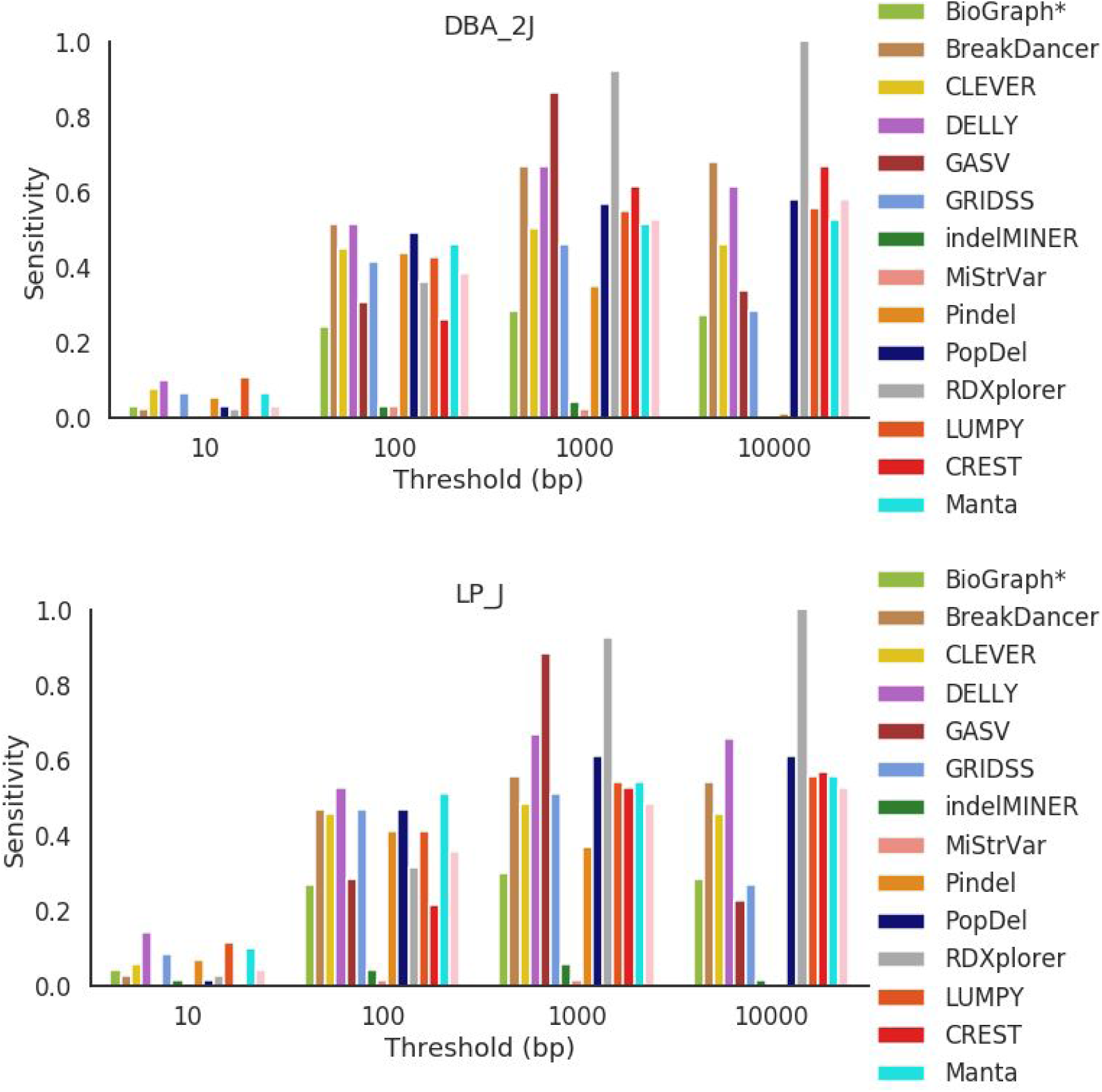
Sensitivity across all mouse strains for deletions between 500 bp and 1000 bp in length.

**Figure S22.**
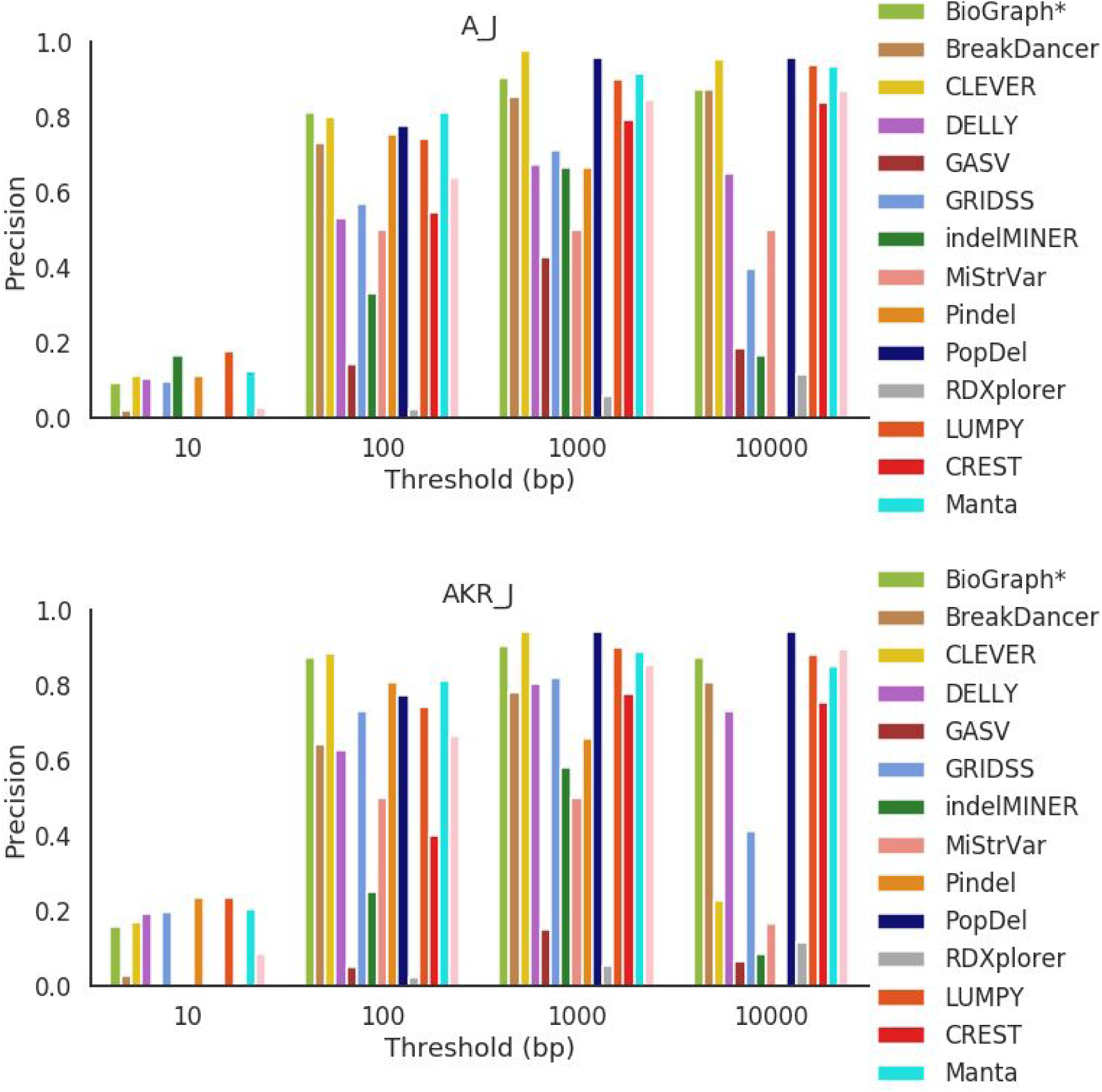
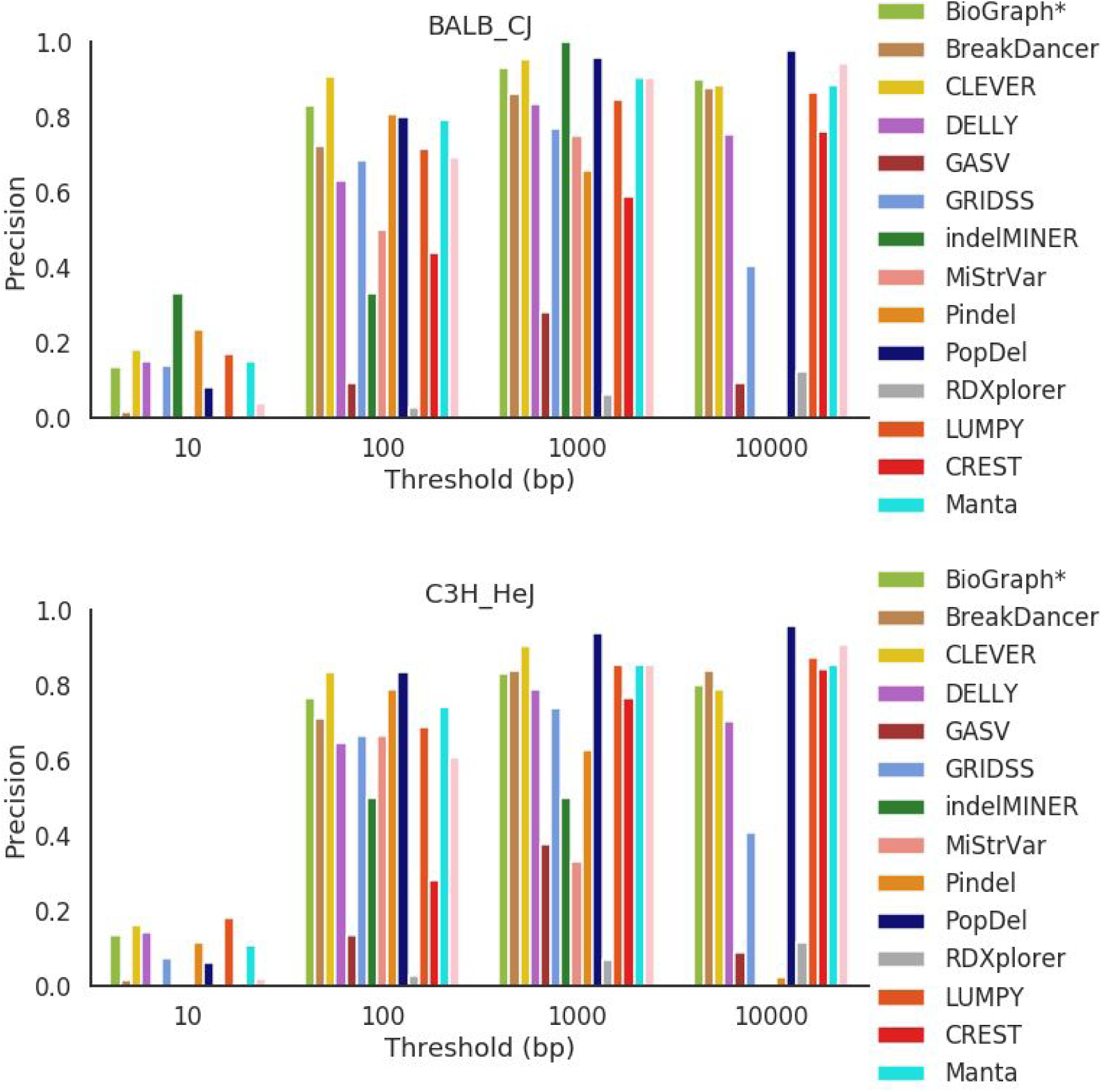
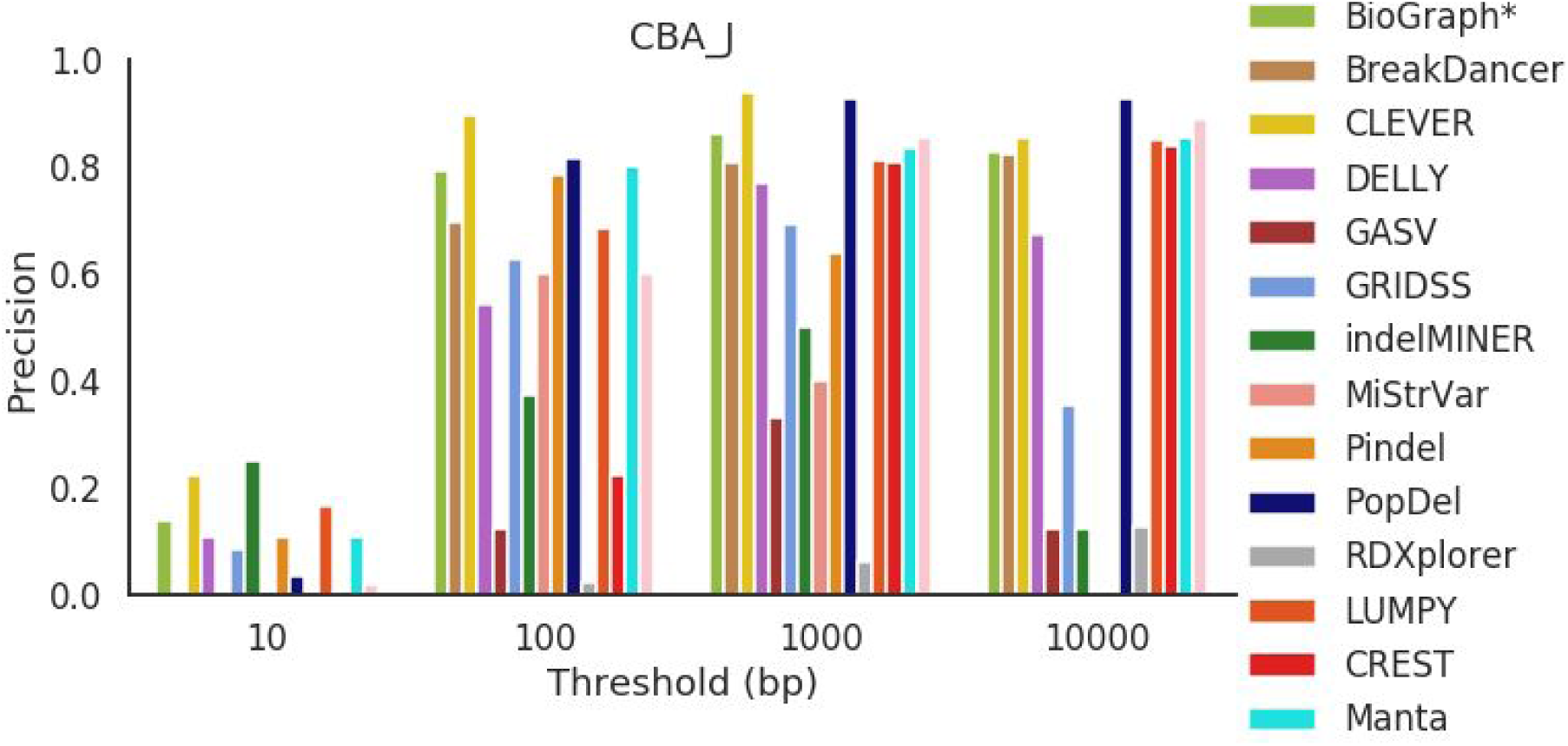
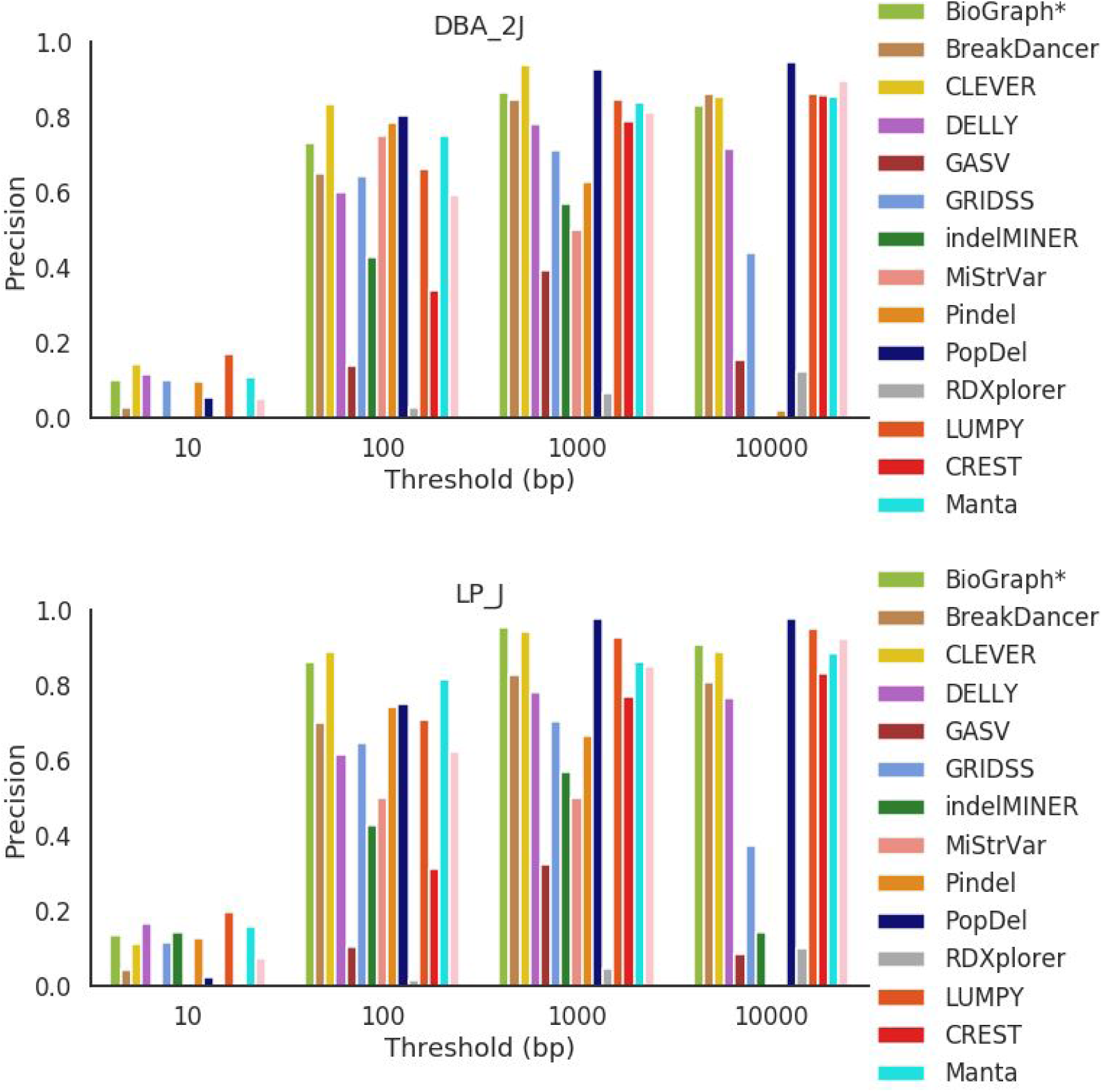
Precision across all mouse strains for deletions between 500 bp and 1000 bp in length.

**Figure S23.**
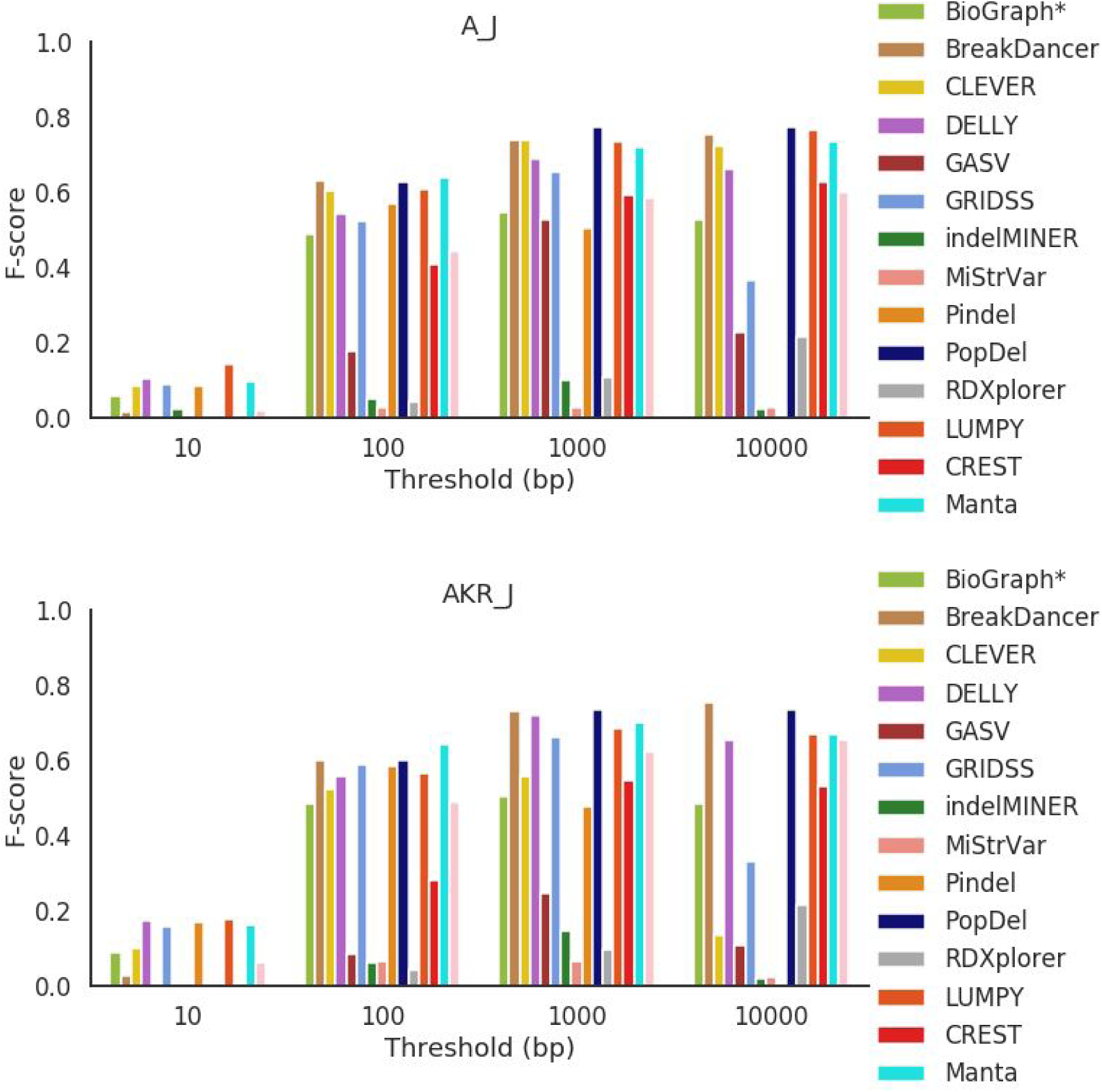
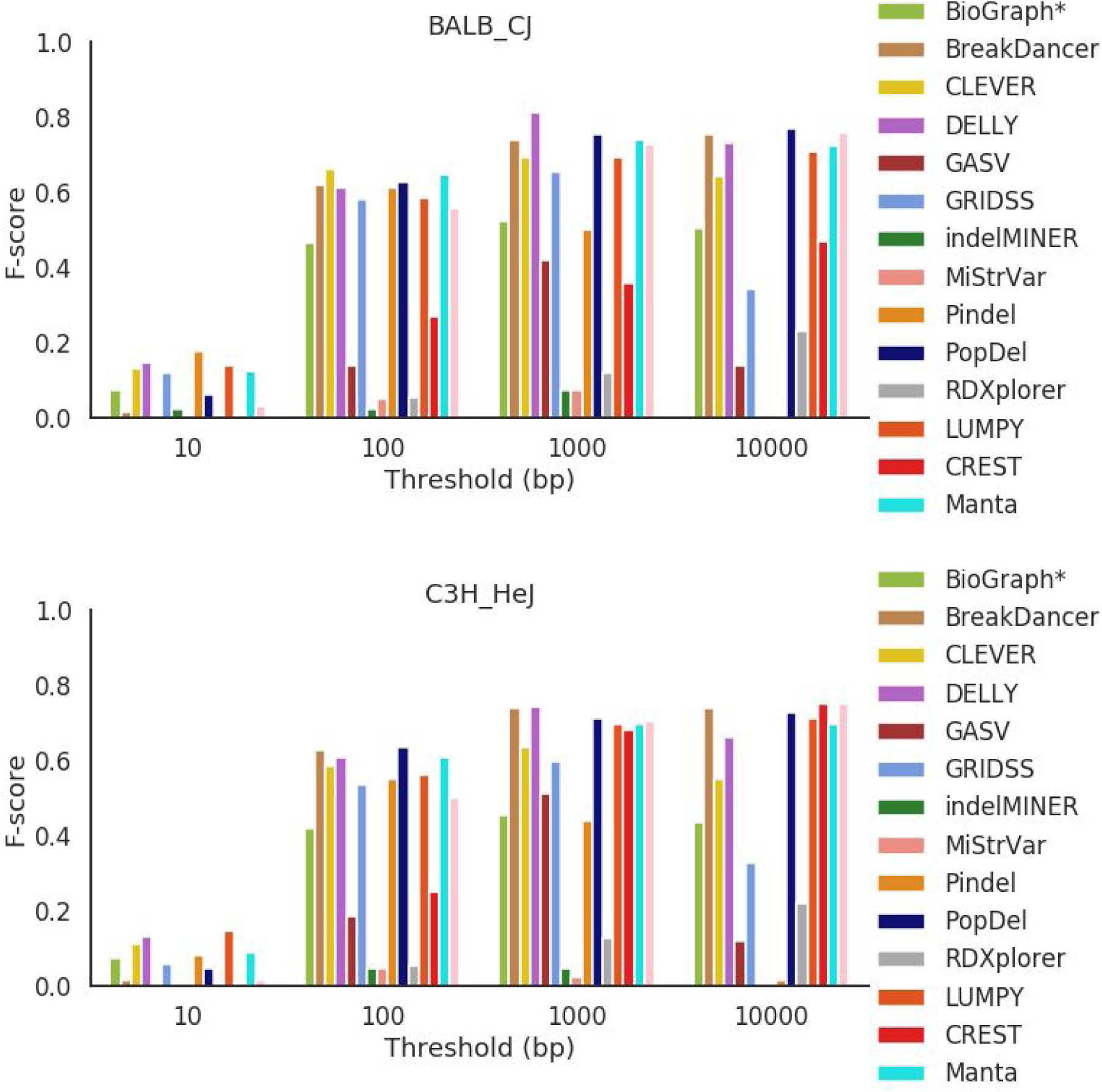
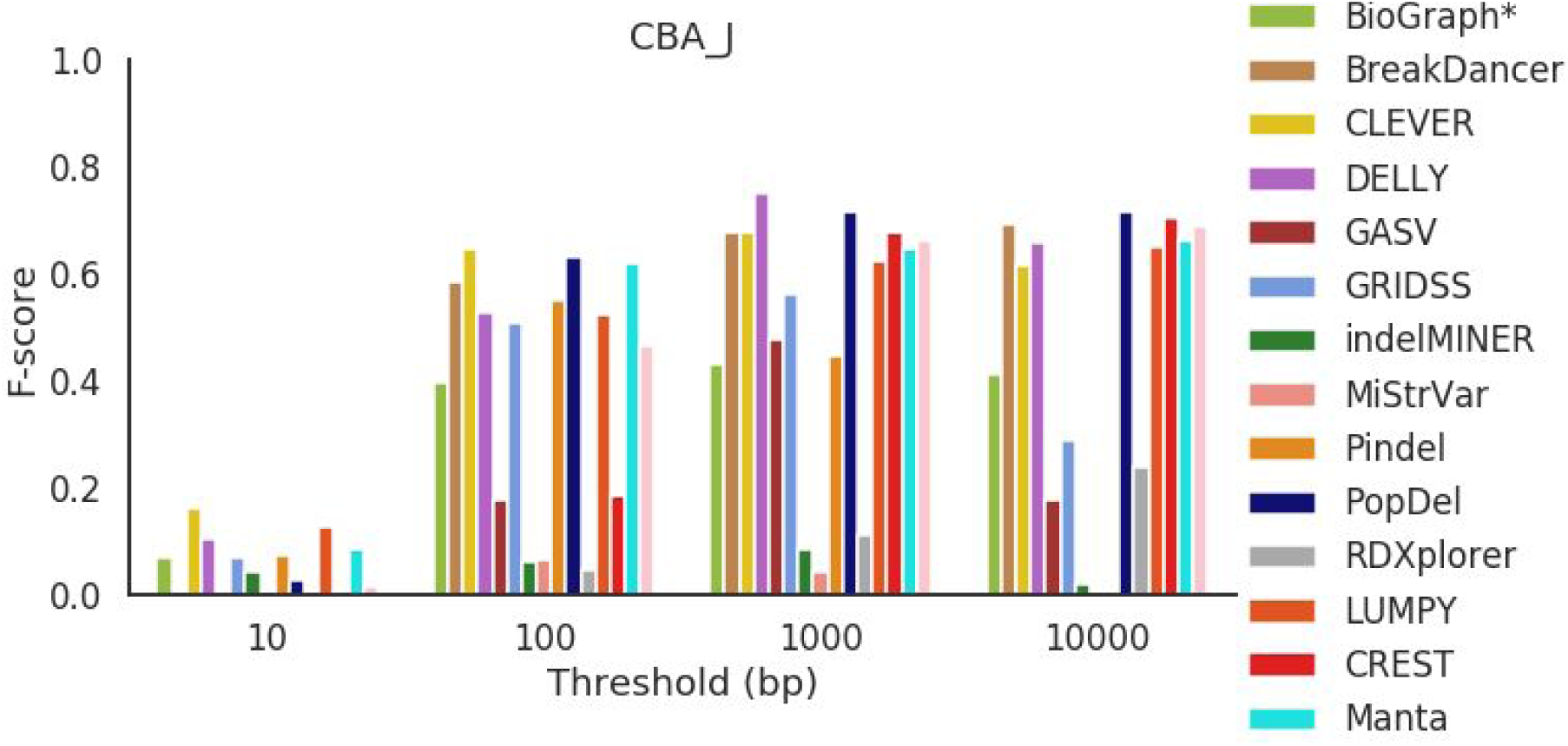
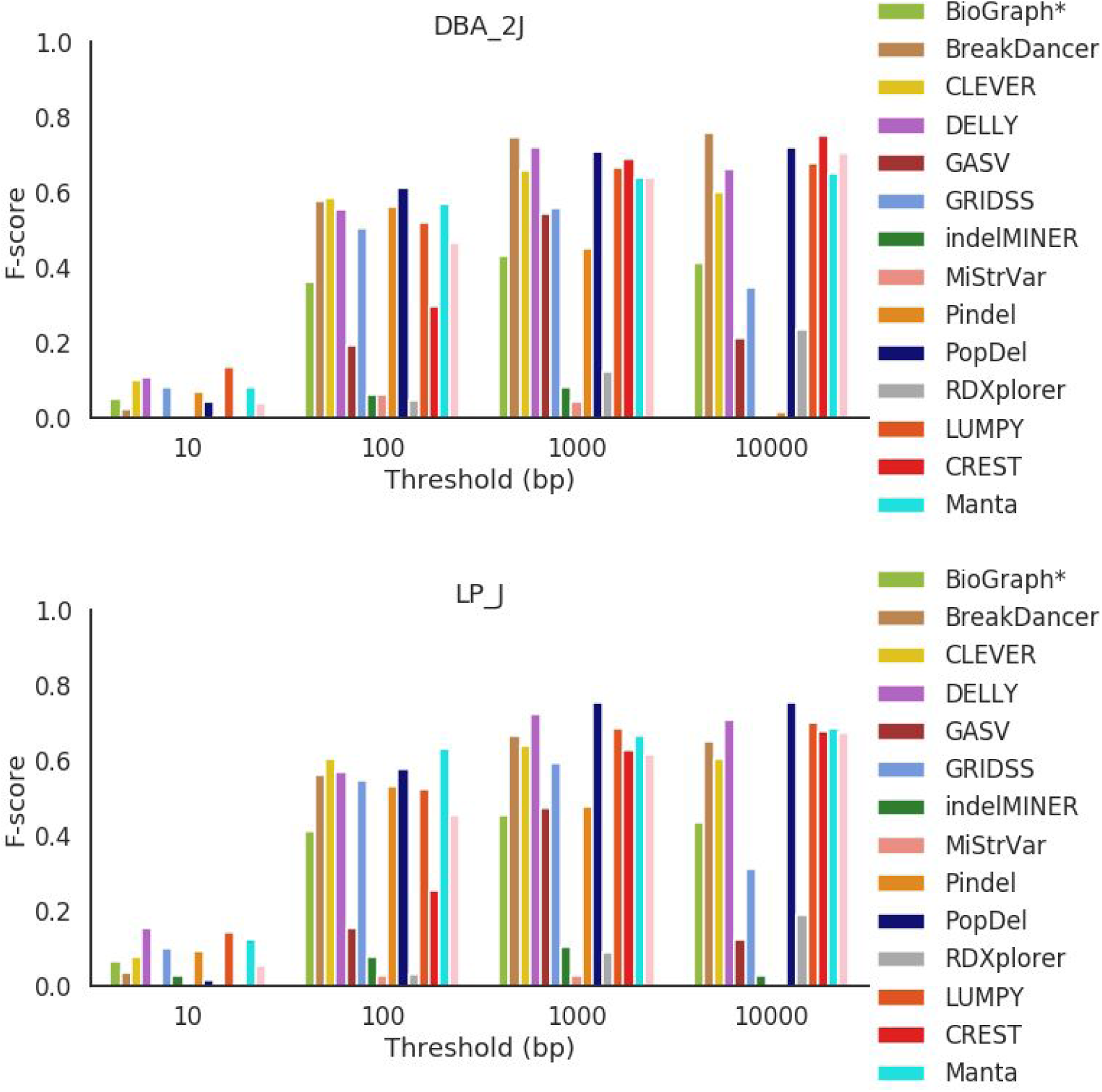
F-score across all mouse strains for deletions between 500 bp and 1000 bp in length.

**Figure S24.**
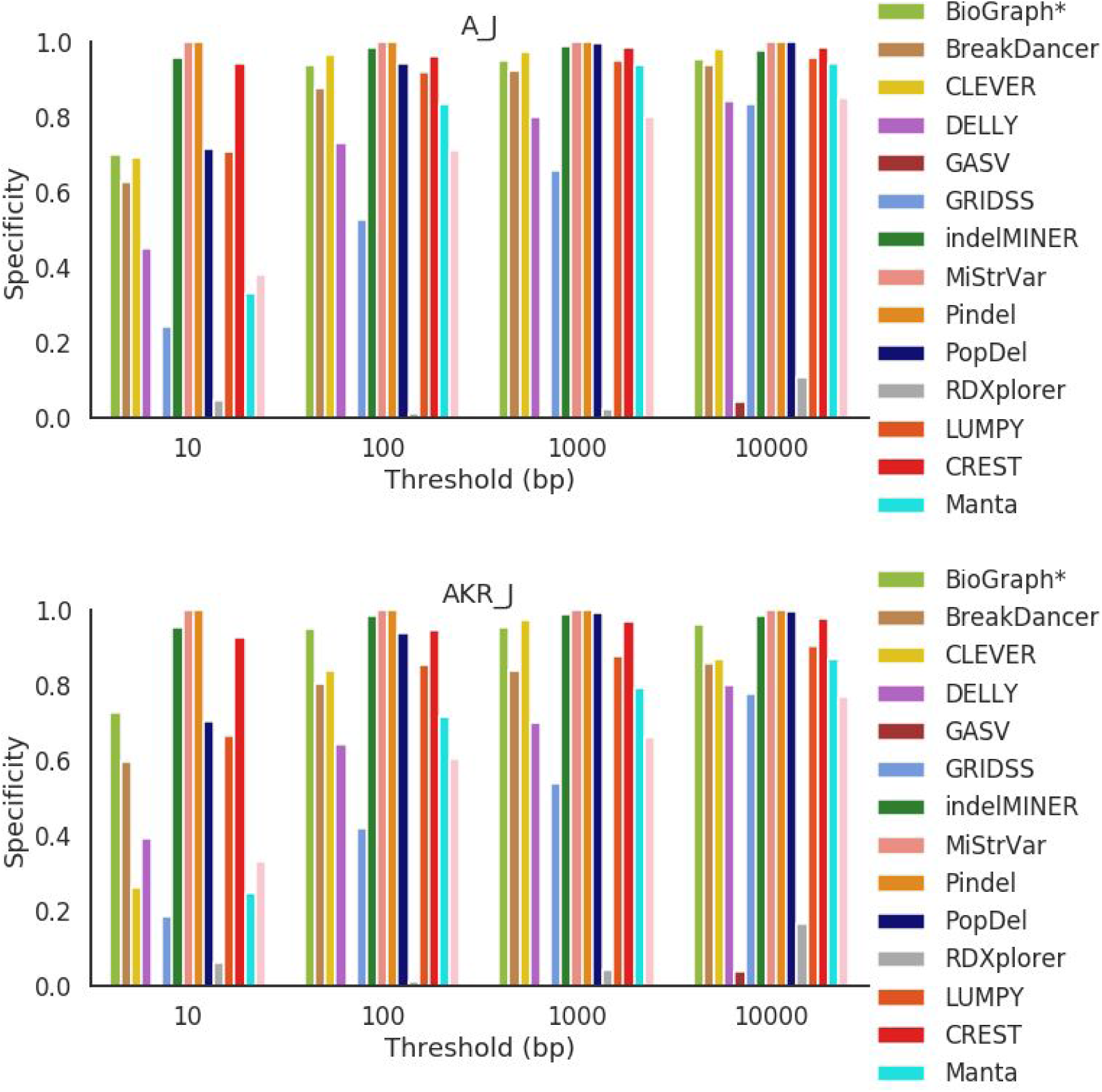
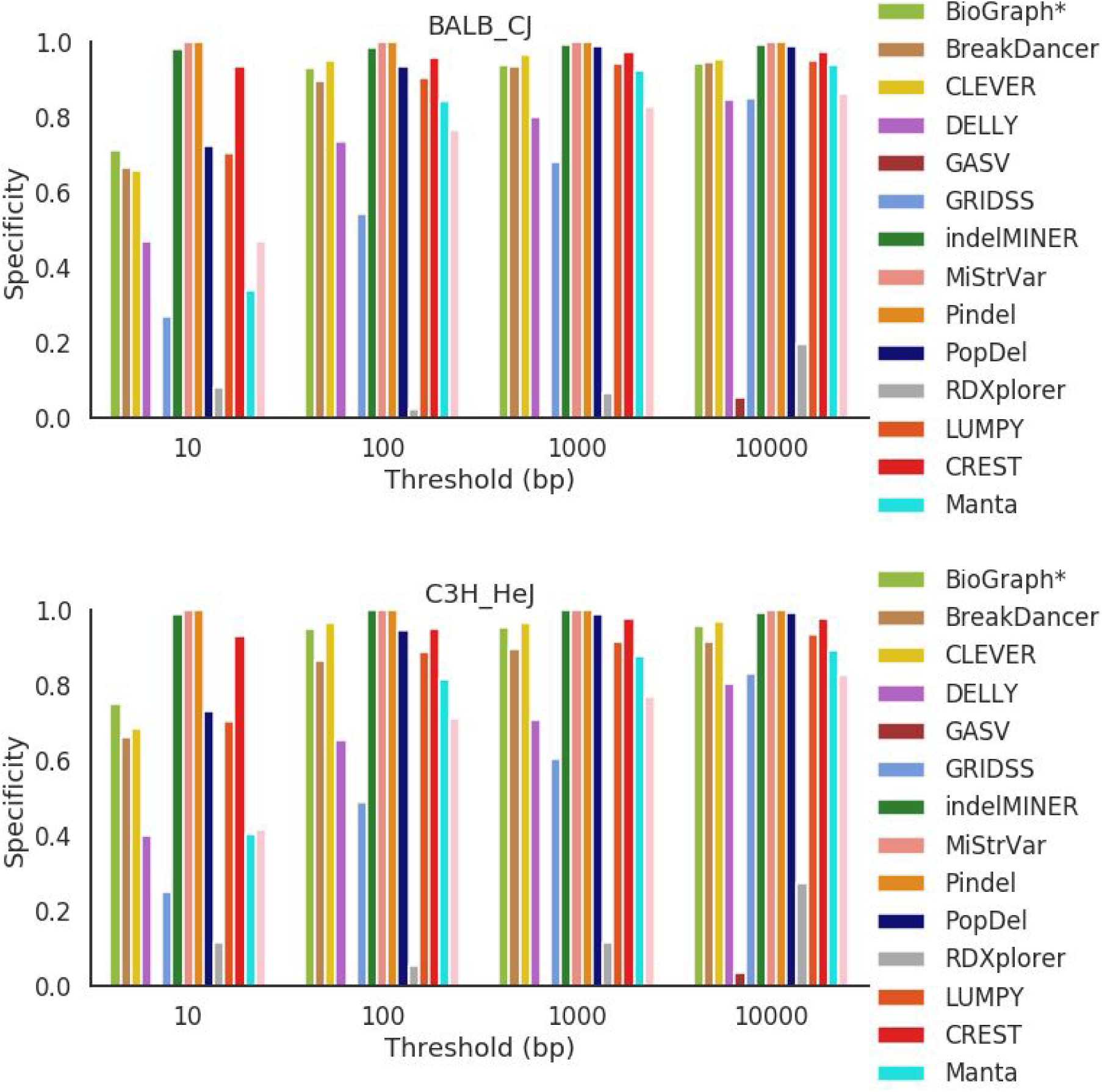
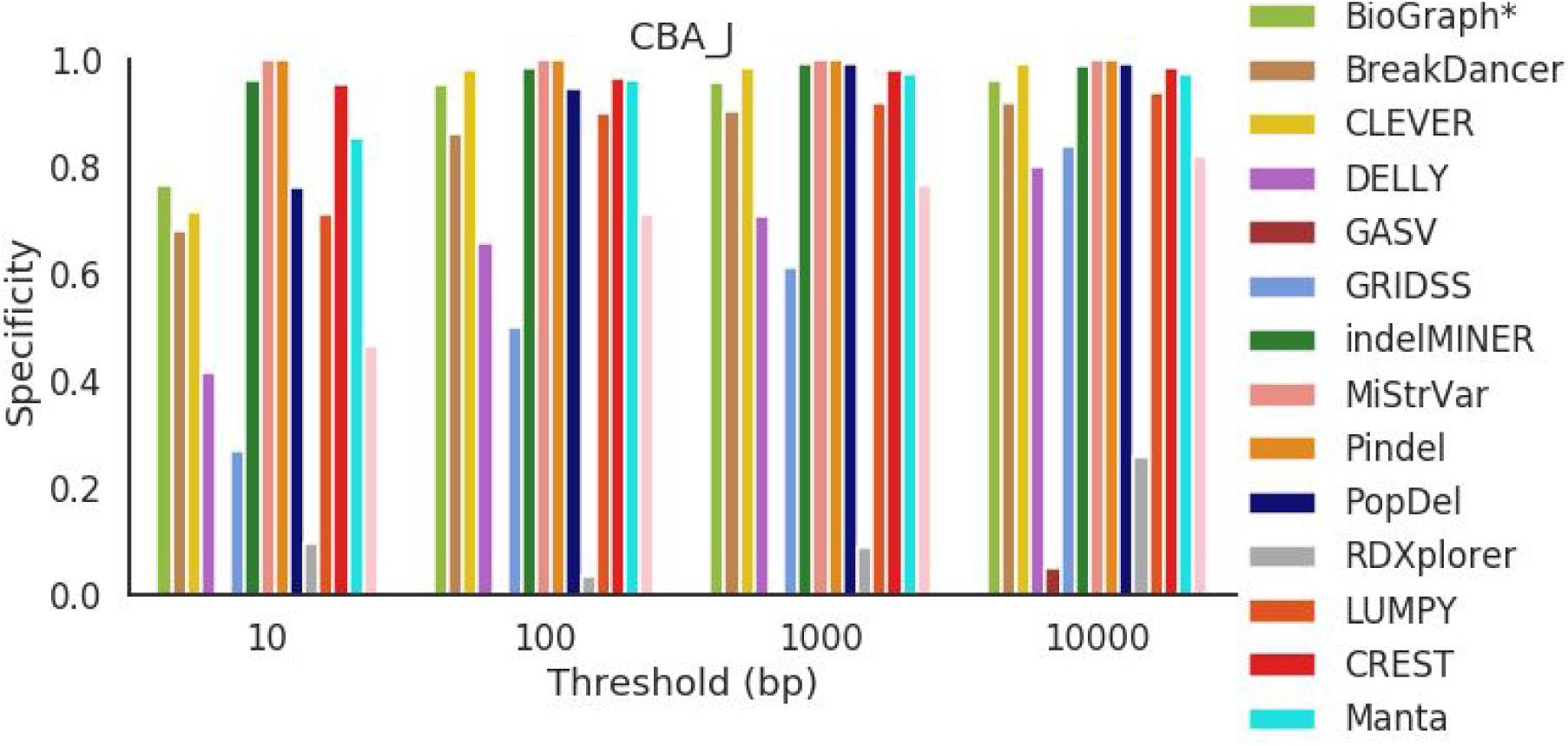
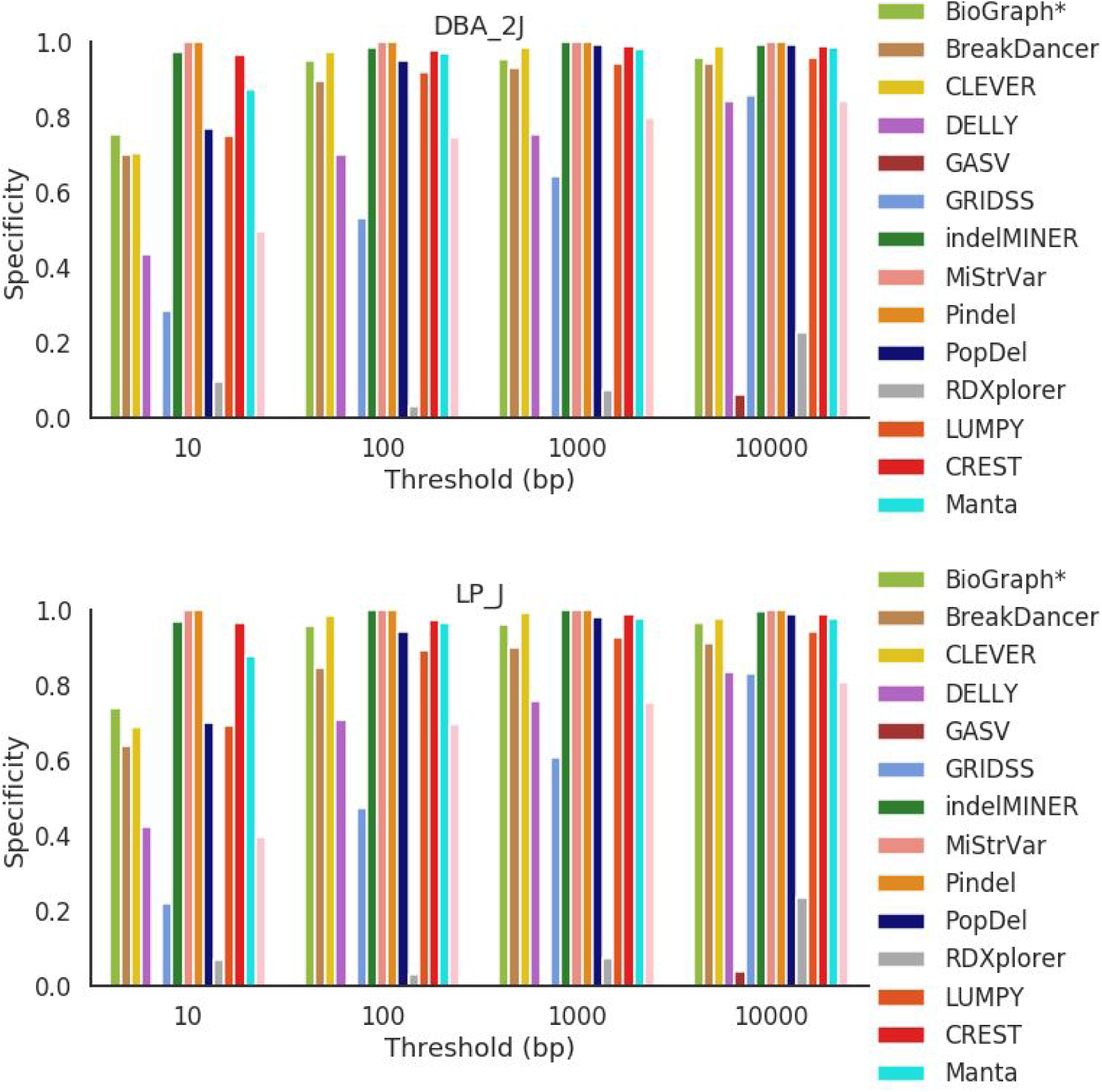
Specificity across all mouse strains for deletions 1000 bp and greater in length.

**Figure S25.**
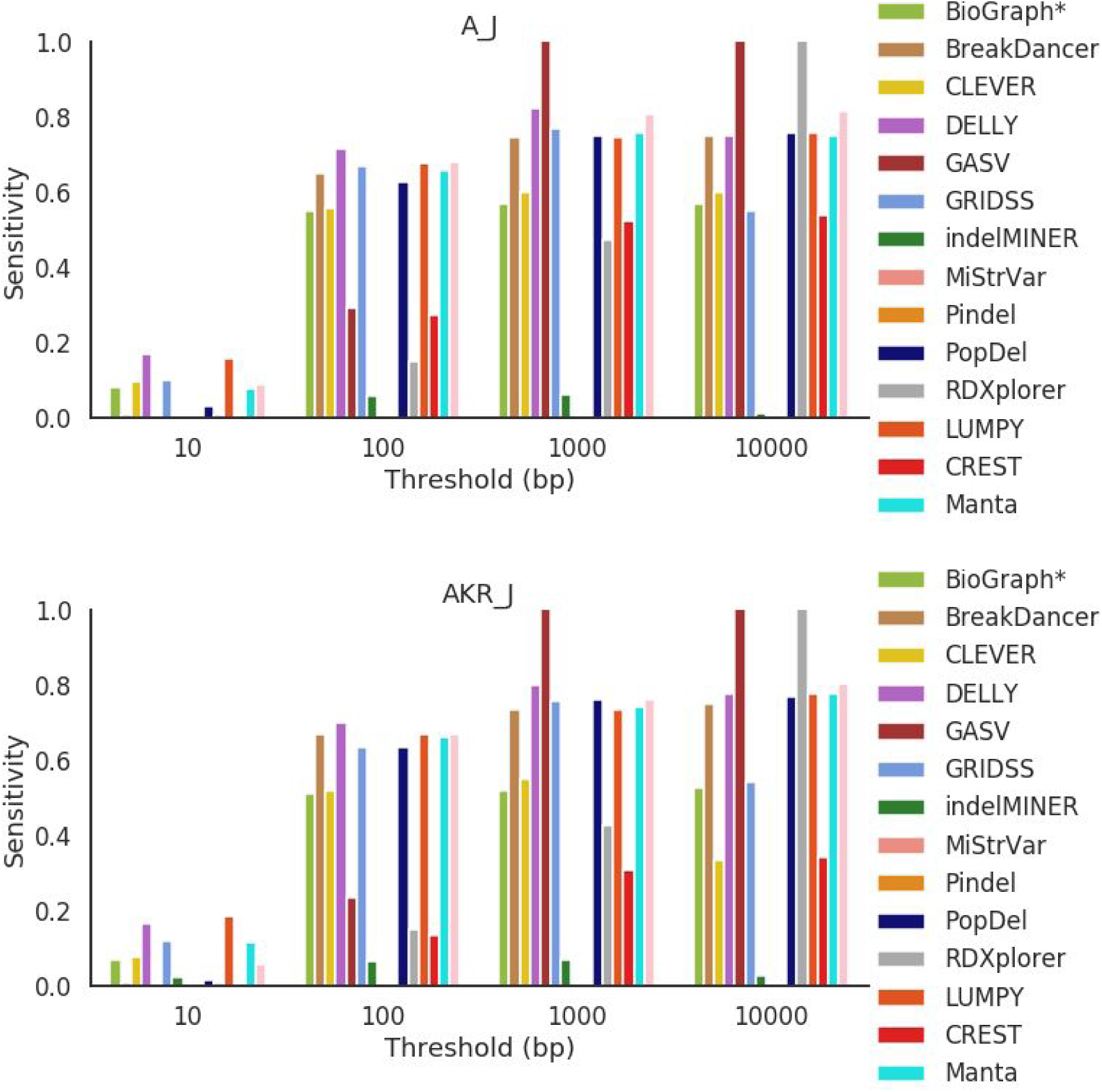
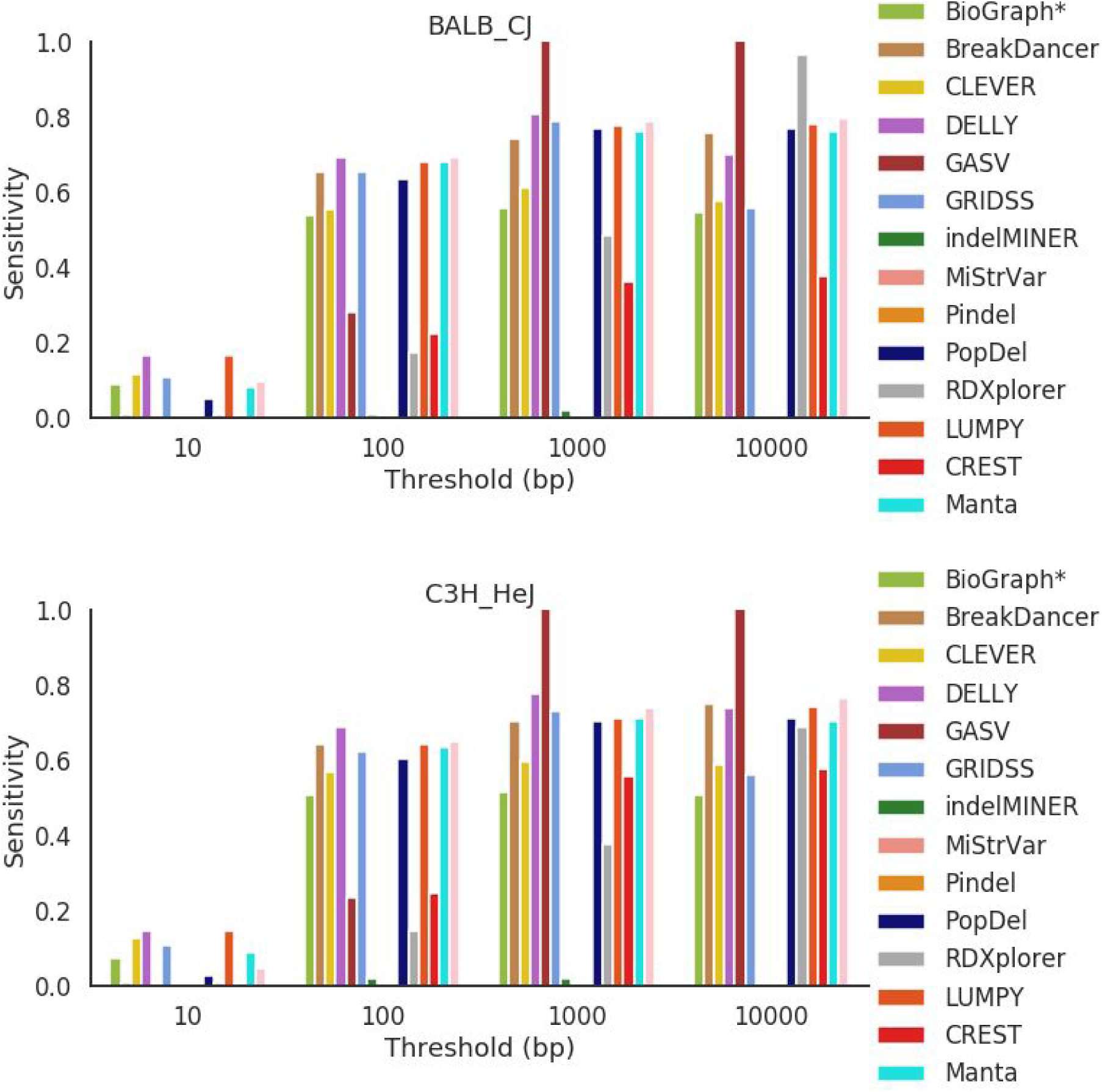
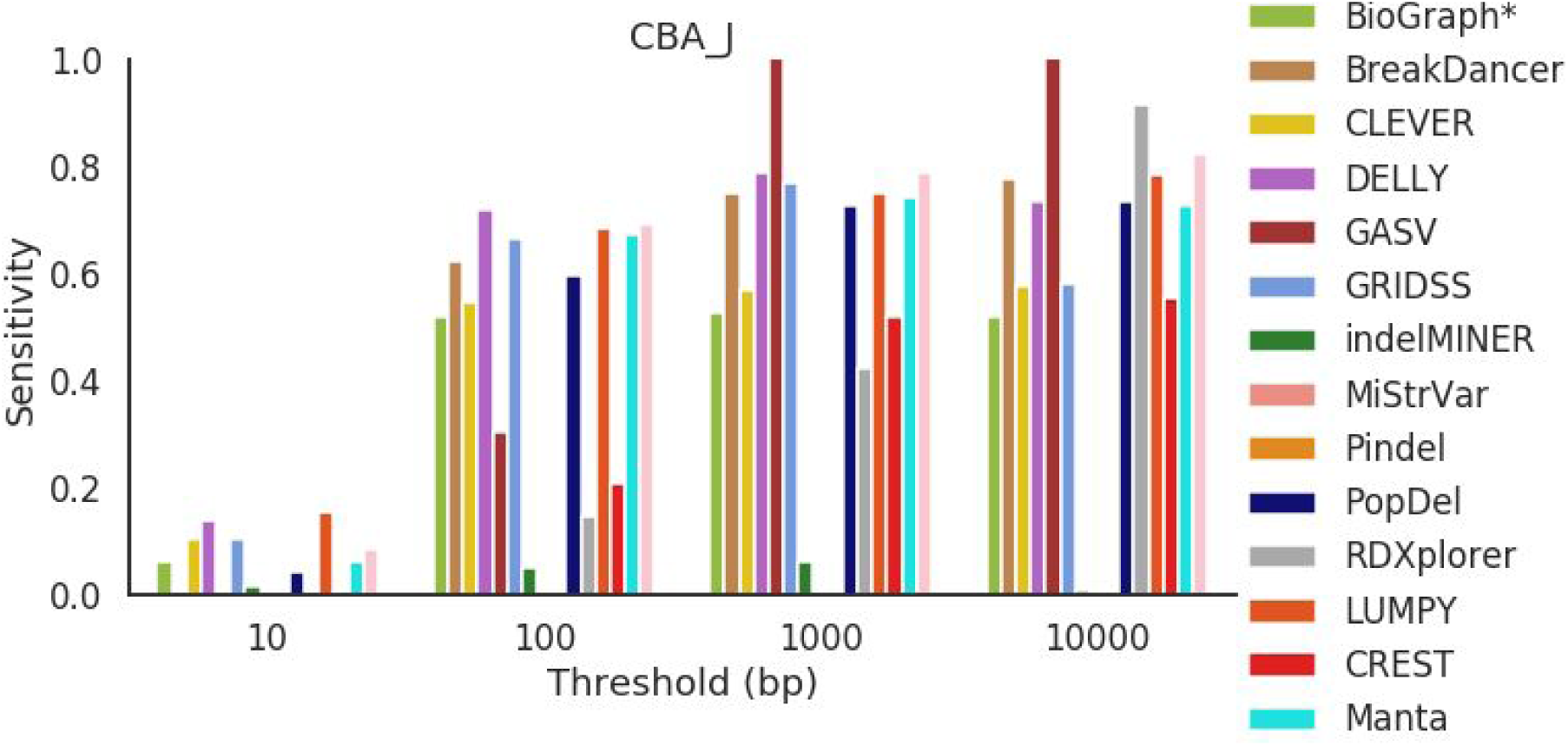
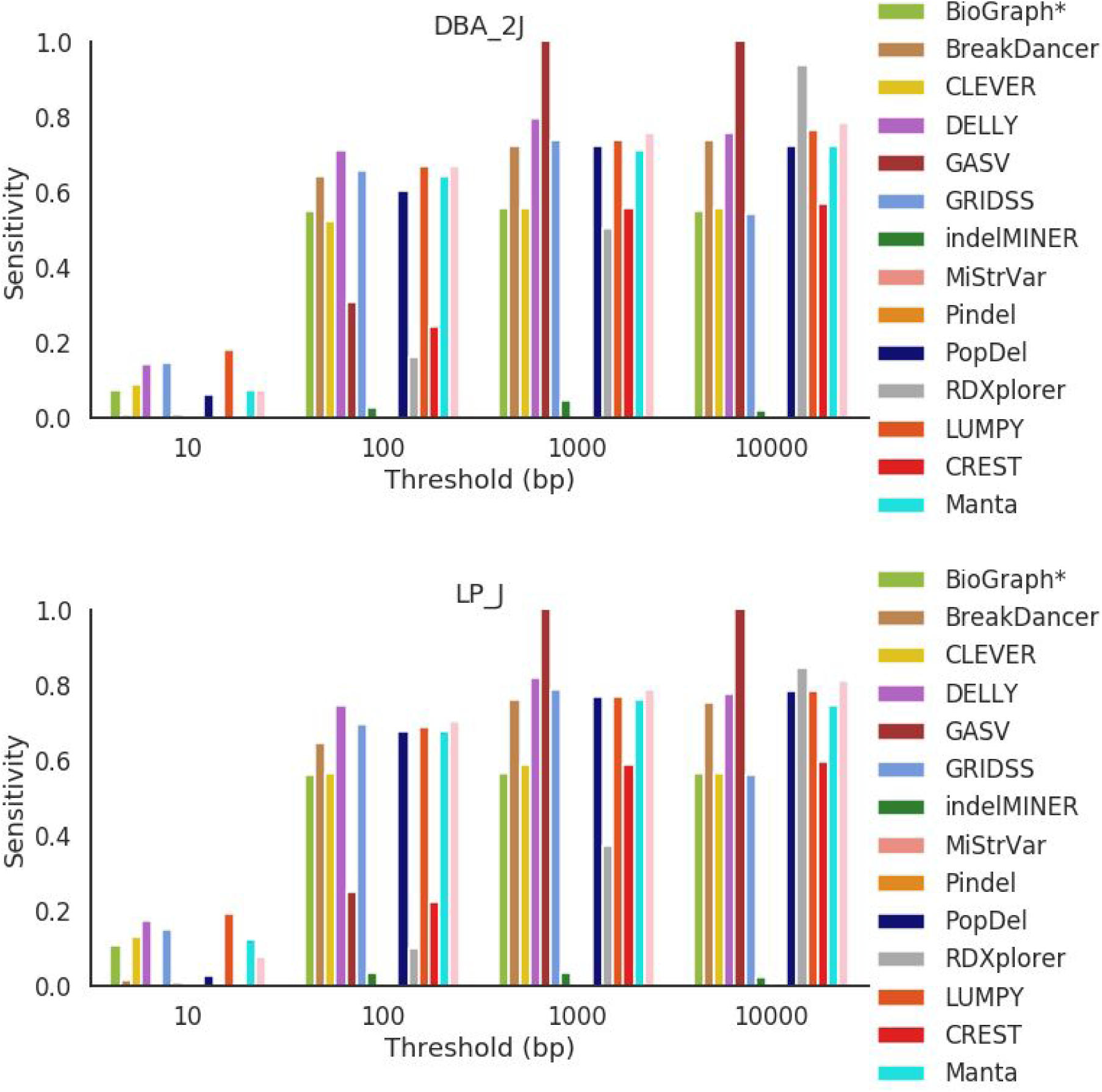
Sensitivity across all mouse strains for deletions 1000 bp and greater in length.

**Figure S26.**
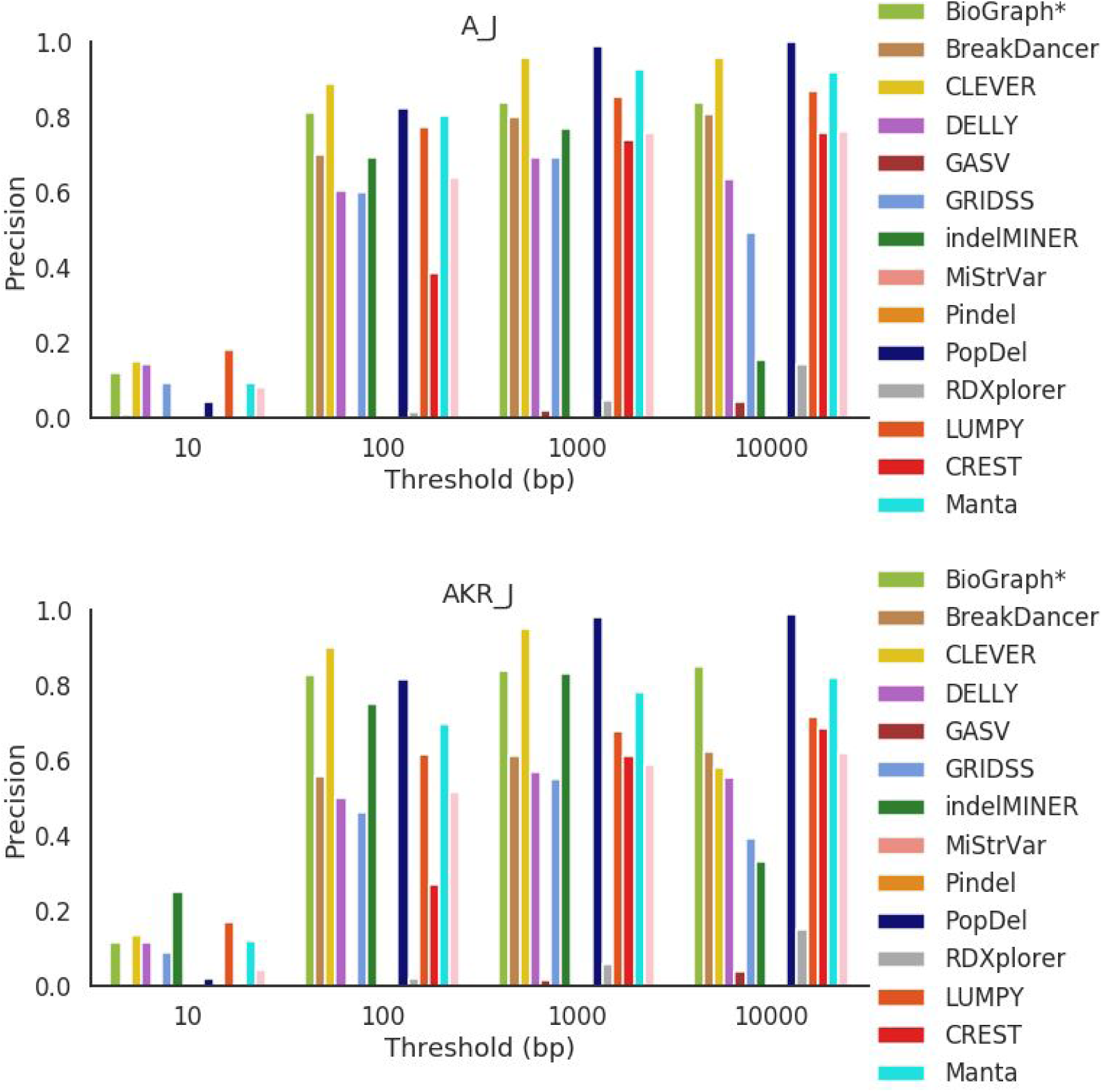
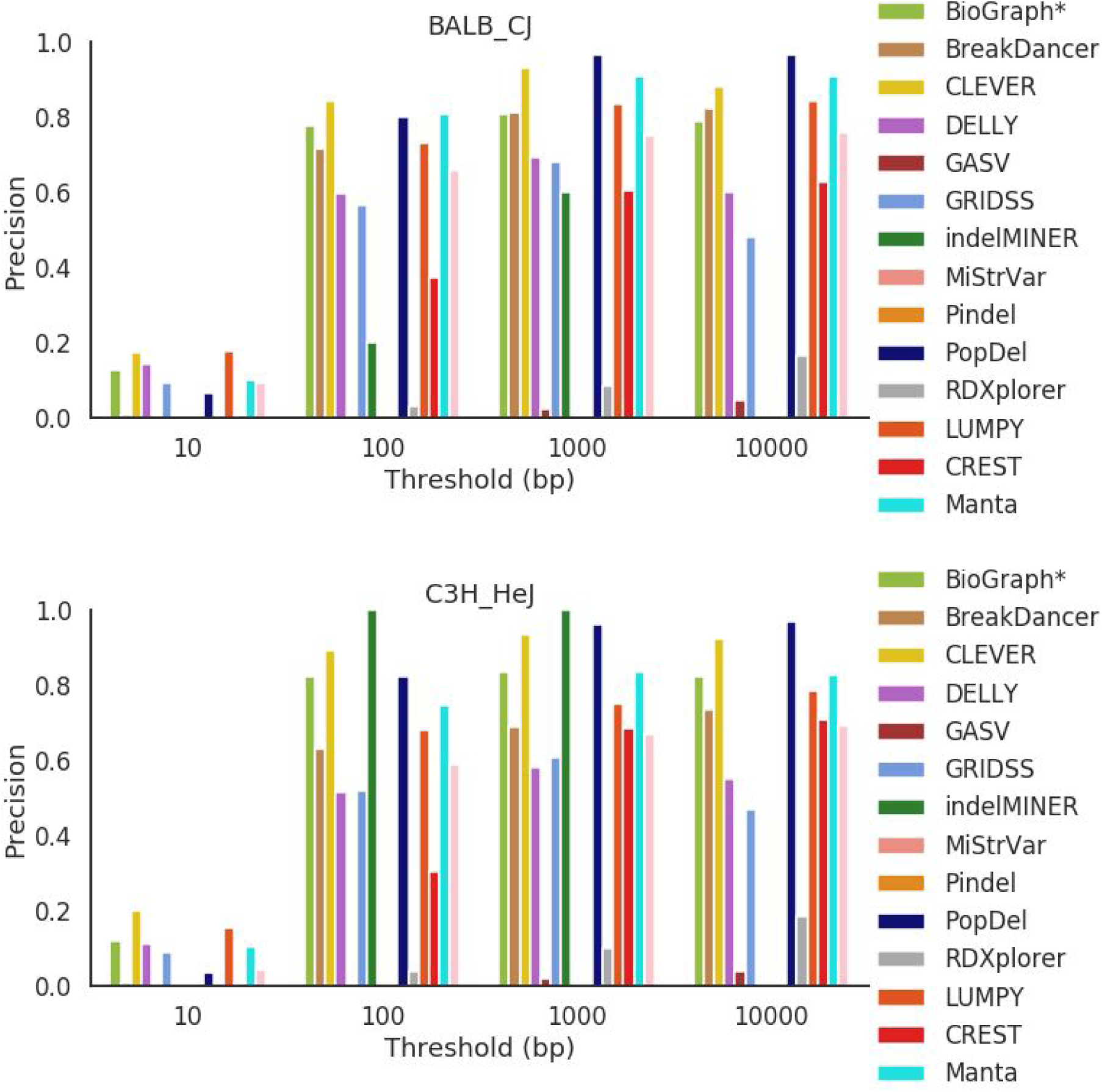
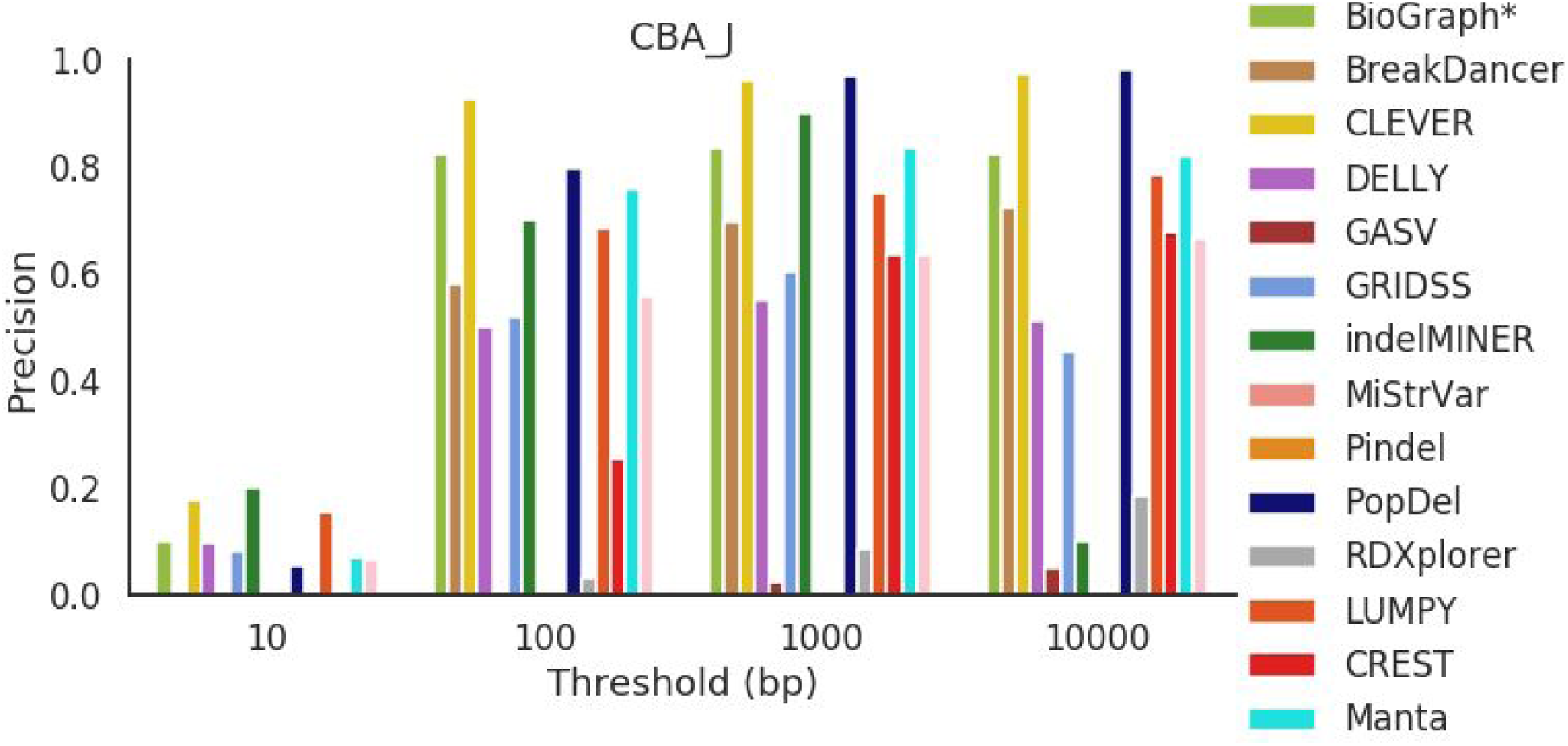
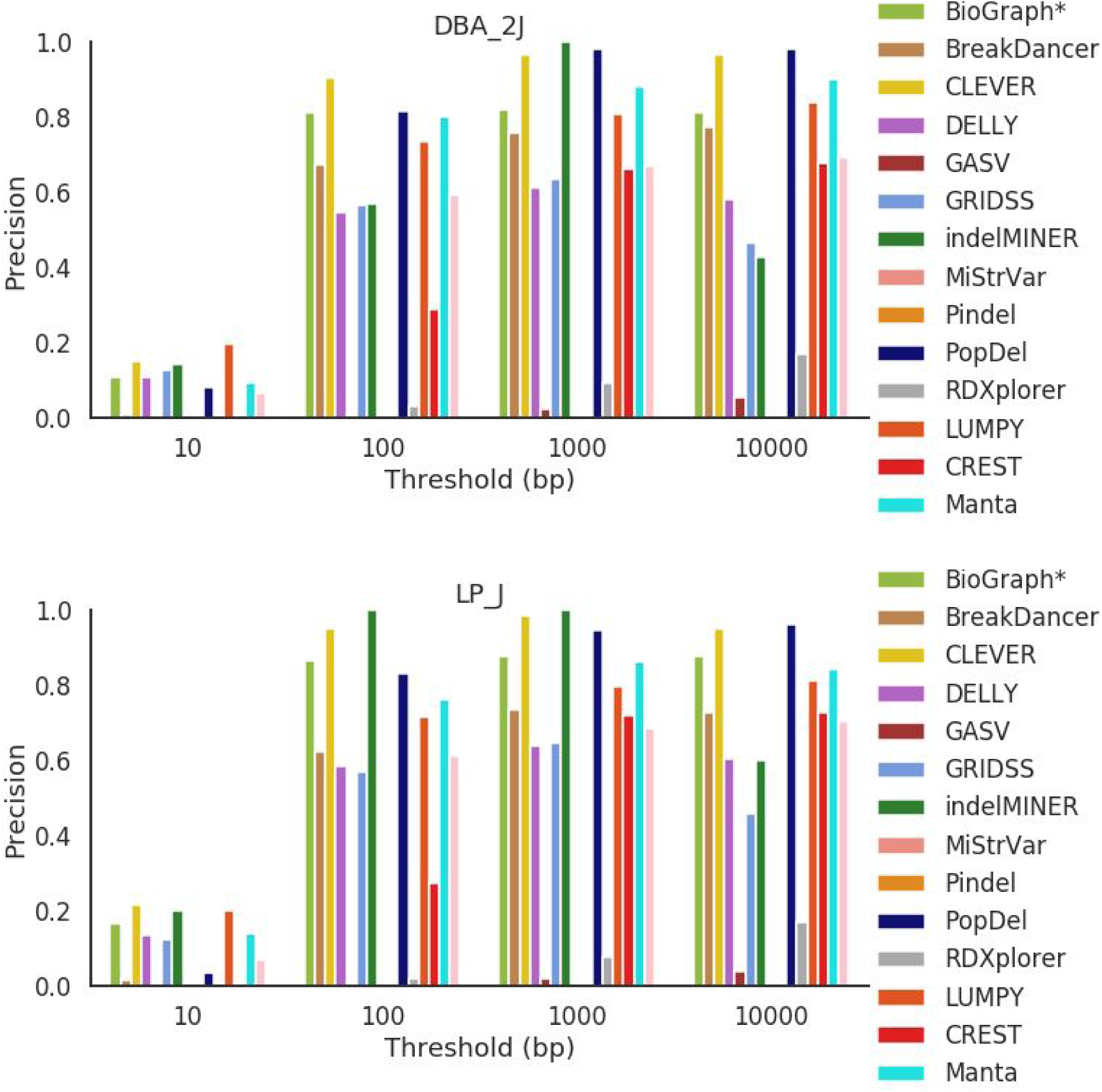
Precision across all mouse strains for deletions 1000 bp and greater in length.

**Figure S27.**
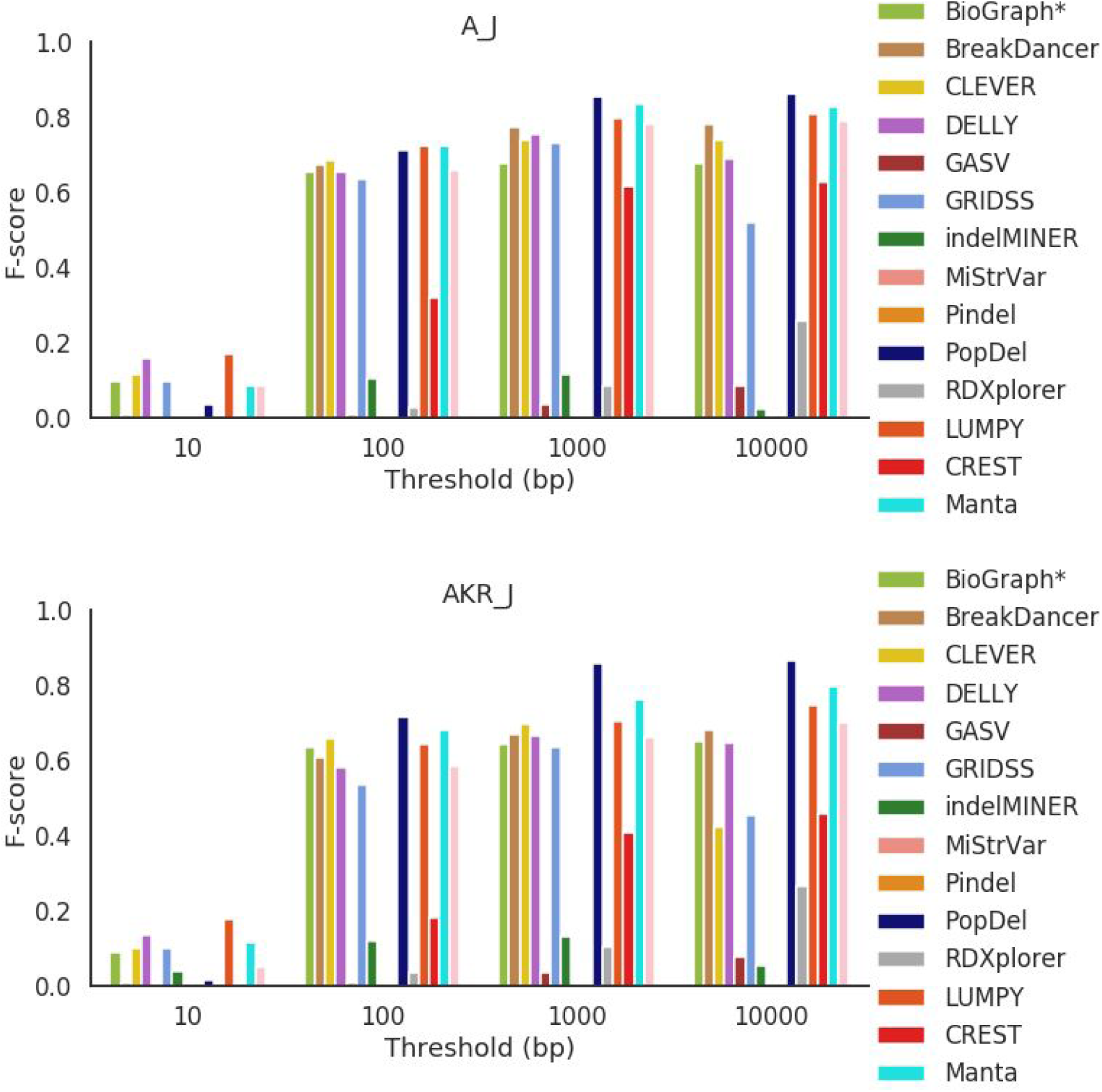
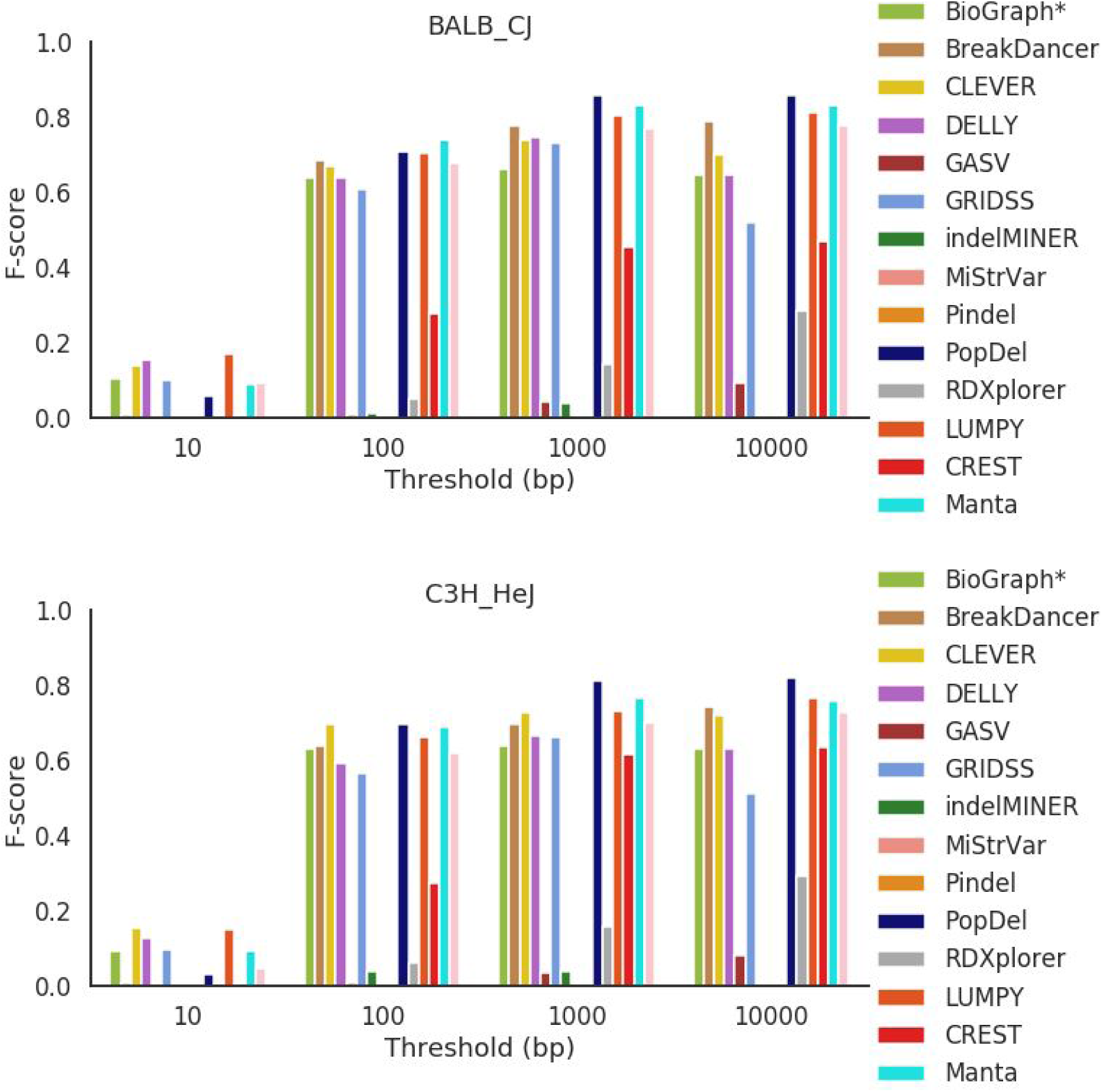
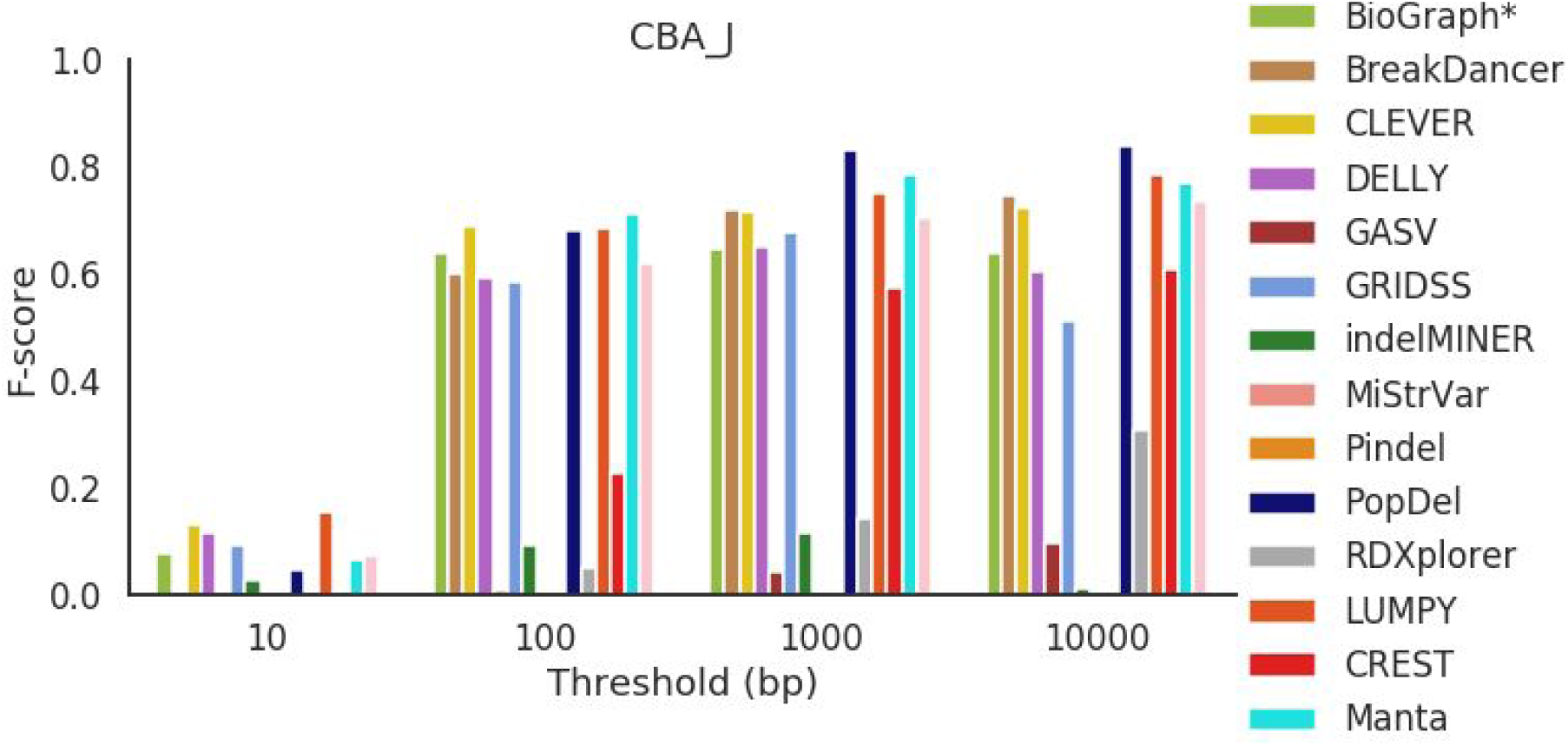
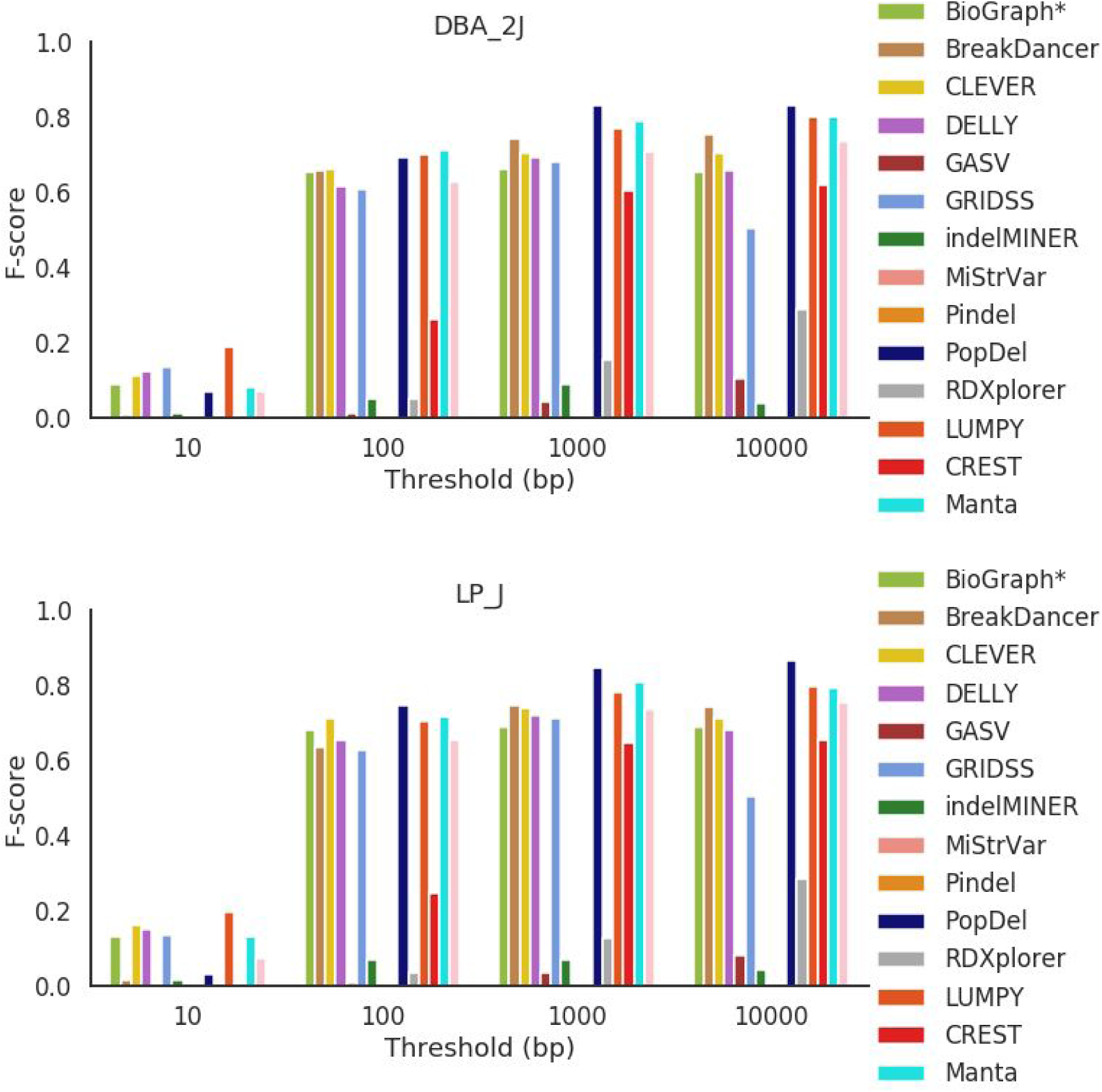
F-score across all mouse strains for deletions 1000 bp and greater in length.

